# Robotic perturbation proteomics and AI agents enable scalable drug mechanism discovery

**DOI:** 10.64898/2026.05.04.722718

**Authors:** Yuming Jiang, Cameron S. Movassaghi, Jesús Muñoz-Estrada, Niveda Sundararaman, Amanda Momenzadeh, Jesse G. Meyer

## Abstract

Large-scale mass spectrometry-based proteomic screening could reveal cellular mechanisms of drug action at systems resolution but remains limited by experimental complexity and the difficulty of extracting insight from high-dimensional datasets. Here, we describe an end-to-end platform that combines semi-automated sample preparation, rapid LC-MS/MS, and AI agent-based data analysis to enable scalable proteomic screening. In a screen of 172 compounds in HepG2 cells, we generated 1,232 proteomes with more than 8,700 quantified proteins in approximately three weeks. Agentic AI reduced data analysis and interpretation time to less than one day while translating proteomic measurements into structured mechanism-oriented summaries and experimentally testable hypotheses. Guided by this framework, we validated: (1) a cholesterol-lowering effect of methylene blue *in vitro* and (2) an association between loratadine exposure and increased circulating iron in matched electronic health record analyses. This work establishes a scalable platform for generating proteomic drug perturbation data and automatically converting that data into mechanistic insights and candidate translational hypotheses using AI.

## Introduction

Proteomics is critical for drug screening because conventional assays capture only limited aspects of drug action and often miss system-wide cellular responses. By directly measuring coordinated changes in protein abundance and activity, proteomics provides a comprehensive, mechanism-level view of drug effects.^1–4^ Recent advances in mass spectrometry have substantially improved the depth, throughput, and reproducibility of proteomic measurements, enabling profiling of thousands of proteins across increasingly large cohorts of samples.^5–12^ These developments have positioned proteomics as a powerful modality for drug screening and mechanism-of-action (MoA) inference.^13–17^

Despite these advances, two major challenges continue to limit the scalability and impact of proteomics-based drug discovery. First, experimental workflows remain complex and labor-intensive, requiring careful coordination of sample preparation, instrument operation, and quality control across multi-plate studies.^18,19^ Second, and more critically, as dataset size and dimensionality increase, extracting biologically meaningful insights from proteomic data has become the primary bottleneck. Large-scale screens generate high-dimensional matrices of protein-level changes across thousands of perturbations, making it difficult to systematically identify shared responses, reconstruct mechanisms of action, and prioritize hypotheses for downstream validation.

In parallel, recent advances in artificial intelligence (AI), particularly large language models (LLMs), have created new opportunities to address this interpretative challenge.^20,21^ Emerging studies suggest that AI systems can assist in organizing biological data, summarizing complex results, and generating hypotheses from heterogeneous sources.^22–26^ However, most existing efforts focus on isolated analytical tasks or literature-based reasoning and remain disconnected from experimental workflows.^27^ A unified framework that integrates large-scale data generation with AI-driven, end-to-end analytical reasoning remains largely unexplored.^28^ More importantly, it remains unclear whether AI systems can faithfully recapitulate the structured analytical logic traditionally applied by domain experts, spanning differential analysis, pathway interpretation, higher-order organization, and ultimately hypothesis generation.^29^

Here, we present an end-to-end proteomics platform that combines high-throughput experimental workflows with an AI-agent–driven analytical framework to enable scalable, mechanism-oriented interpretation of drug-induced proteomic responses. Using a HepG2-based screen of 172 compounds, we generated a deep and high-quality proteomic dataset comprising over 1,230 samples and 8,700 quantified proteins. We then applied agentic AI systems, Science Machine^30^, KOSMOS^31^ and GPT-based multi-agent systems, to transform raw quantitative outputs into structured biological interpretations under a predefined analytical logic. We demonstrate that, when guided by explicitly defined analytical frameworks, AI-assisted analysis can faithfully recapitulate conventional expert-driven workflows, including differential regulation analysis, pathway enrichment, and clustering-based organization of drug responses. Beyond reproducing known biology, the framework systematically identifies both shared and compound-specific proteomic signatures, generates structured molecular identity profiles for each drug, and prioritizes biologically coherent yet underexplored hypotheses. Finally, we validate AI-prioritized predictions of unexpected drug effects using orthogonal approaches, including biochemical assays and real-world electronic health record (EHR) data. Taken together, this work establishes a scalable “sample-to-insight” paradigm for proteomics-driven drug screening, demonstrating that AI systems can move beyond assisting with analysis to actively structuring biological interpretation and accelerating the generation of experimentally and clinically testable hypotheses.

## Results

### An end-to-end proteomics workflow enables large-scale drug screening

We established an end-to-end semi-automated workflow spanning cell-based drug perturbation, 96-well plate-based sample preparation, rapid LC–MS/MS acquisition, standardized data processing, and AI-assisted downstream interpretation (**Fig. 1a**). This workflow integrates a series of liquid handlers, including an automated platform for low-volume liquid handling, cell seeding and media washing tools, liquid pipetting and removing tools, and an automated platform for SP3-based proteomics sample preparation.^32^ The integrated use of these automation tools was designed to minimize manual intervention and time, thereby substantially improving efficiency while preserving the depth and consistency required for biologically meaningful analysis across multiple 96-well plates. We employed the Orbitrap Astral platform to rapidly collect deep proteomics data. We further integrated DIA-NN software^33^ with an HPC computing platform for raw data processing and used agentic AI tools to automate interpretive analysis starting from the raw proteomic data matrix.

**Figure 1.**
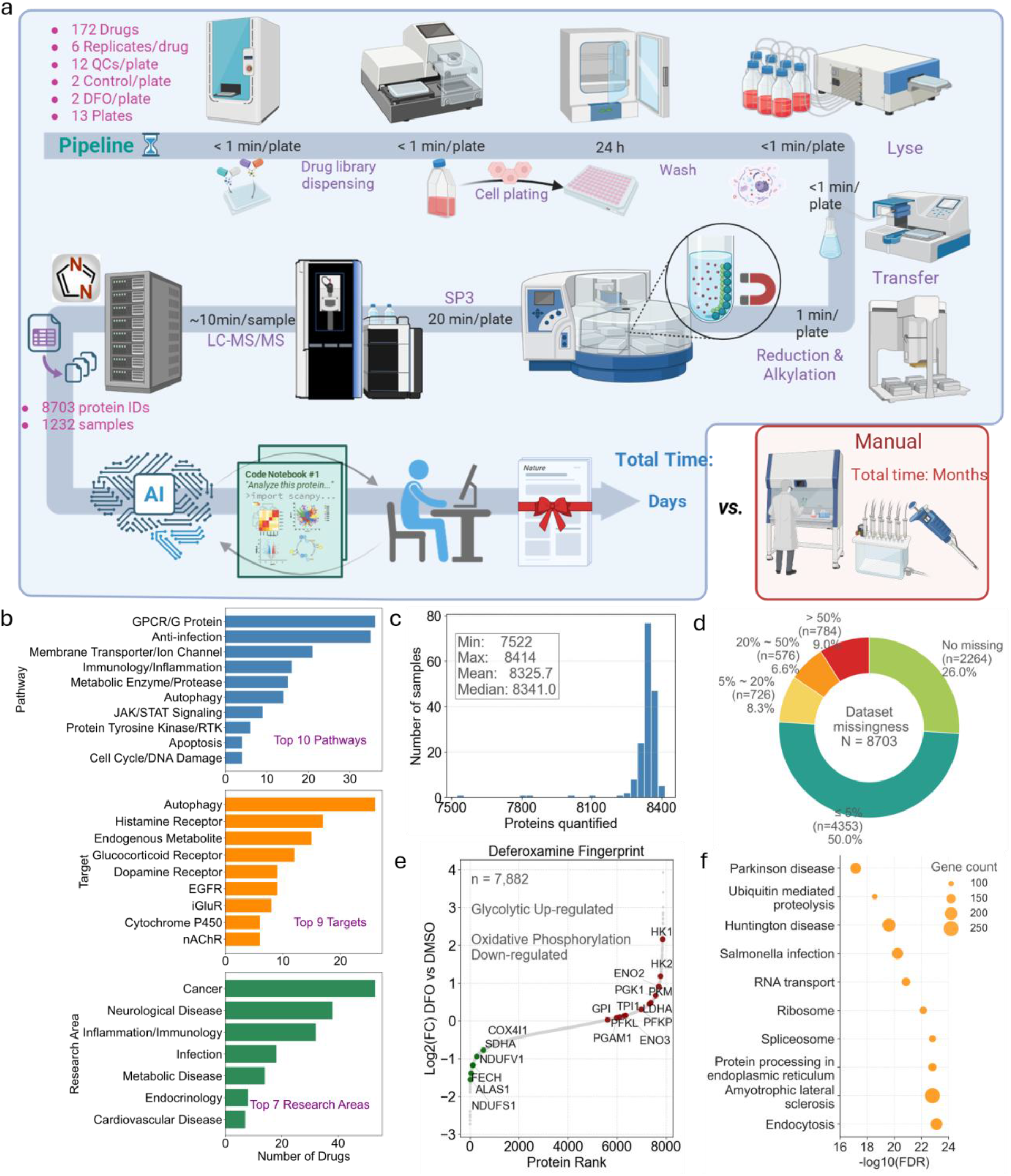
An end-to-end high-throughput deep proteomics workflow for large-scale drug screening. **a**, Overview of the end-to-end workflow from HepG2 cell culture in 96-well plates through rapid proteome acquisition, matrix generation, standardized downstream analysis, and AI-assisted interpretation/reporting. The screening comprised 172 compounds with six biological replicates per compound distributed across 13 96-well plates. Each plate included 12 QC injections for run-to-run calibration, 2 DMSO vehicle controls (negative controls), and 2 deferoxamine (DFO) positive controls to monitor assay performance. **b**, Composition of the screened compound collection summarized by annotated pathways (top), targets (middle), and research areas (bottom). **c**, Proteome depth per compound across the dataset, quantified as the number of proteins identified per compound (union across six biological replicates). **d**, Missingness landscape of the 8,703 quantified proteins across all samples, reporting the distribution of protein-level missing rates (fraction of missing values across samples). **e**, Representative proteome fingerprint of the positive control DFO highlighting increased abundance of glycolysis-associated proteins and decreased abundance of oxidative phosphorylation proteins, shown as log2 fold change (DFO vs DMSO). **f**, Pathway-level interpretation of the dataset by KEGG enrichment analysis performed on the identified proteome, summarizing the major biological processes captured by the screening.

In this study, we applied the platform to a large-scale HepG2 compound screen comprising 172 compounds, each measured in six biological replicates across thirteen 96-well plates. To support longitudinal quality control and assay calibration, each plate further included 12 QC samples, two DMSO vehicle controls, and two deferoxamine (DFO)-treated positive controls. The screened compound collection spanned a broad range of annotated biological pathways, molecular targets, and therapeutic research areas, indicating that the dataset captured chemically and functionally diverse perturbations rather than a narrowly focused compound class (**Fig. 1b, Fig. S1**). The workflow yielded a proteomic matrix of 1,232 samples and 8,703 quantified proteins (**Fig. 1a, Supplementary Table 1)**, totaling more than 10 million data points and enabling high-depth, scalable measurements across the screening library. At the compound level, when aggregated across the six biological drug replicates in each set, proteome coverage was consistently high, with a median of 8,341 proteins identified and a range of 7,522 to 8,414 proteins per compound group (**Fig. 1c**). Data completeness was likewise favorable: 76% of quantified proteins exhibited missing values in less than 5% of samples, whereas only a minority showed extensive missingness across the dataset (**Fig.1d**). Consistent with these global metrics, QC samples remained stable throughout the acquisition sequence, with limited drift in intensity and protein identifications (**Fig. S2a**). Batch-related structure was effectively reduced after correction by either sample type or loading plates (**Fig. S2b**). Reproducibility was additionally supported by the coefficient-of-variation distributions (∼10%) across sample classes, and protein identifications remained consistently high at the individual-sample level (**Fig. S2c, d**).

As an internal benchmark for biological sensitivity, the positive control DFO produced a characteristic and highly coherent proteomic response, including increased abundance of glycolysis-associated proteins and reduced abundance of oxidative phosphorylation components relative to DMSO controls (**Fig.1e**). Consistent with the breadth of the screened library, pathway-level annotation of the quantified proteome further highlighted enrichment of diverse cellular programs, including GPCR/G protein signaling, membrane transport, immune and inflammatory pathways, metabolic regulation, autophagy, and apoptosis-related processes (**Fig. 1f, Fig.S3a**). Collectively, these data establish that the platform can generate deep, high-quality, and biologically interpretable proteomic profiles in a format compatible with large-scale drug screening. Notably, by our estimation, completing a drug screening experiment of this depth and scale can be completed in only approximately 25 days from cell treatment to the generation of final experimental results and hypotheses, with the involvement of just two postdoctoral researchers. More specifically, the process required approximately 3 days for drug treatment and sample preparation, 14 days for MS acquisition, 7 days for MS raw data analysis and 1 day for interpretive data analysis with AI (**Fig. S3b**). Such throughput represents an unprecedented leap in efficiency compared to conventional workflows.

### AI analysis recapitulates manual interpretation and accelerating hypothesis generation

Having validated the dataset’s robustness, we first applied two AI-agent-driven frameworks, powered by the Science Machine and KOSMOS AI systems, to complement our automated sample preparation and high-throughput acquisition (**Fig. 2a–d**). In this architecture, Science Machine and KOSMOS were integrated as the primary engines for the end-to-end analytical pipeline, spanning raw proteomic matrix processing, result interpretation, and hypothesis generation. To guide analysis in a reproducible and biologically informed manner, we provided the systems with comprehensive single-shot prompts that included a detailed analytical logic chain, explicit task requirements, and stepwise instructions for data interpretation (**Supplementary File 2**), along with the raw proteomic dataset and sample annotations (**Supplementary Table 3**). This design enabled the AI platforms to perform data analysis and interpret results in a structured fashion, returning organized analytical reports that included differential-regulation summaries, mechanism-relevant interpretations, known drug-class associations, and candidate hypotheses for previously unrecognized effects. Rather than serving as isolated tools, these systems functioned as a unified analytical interface, translating complex proteomic outputs into coherent biological claims.

**Figure 2.**
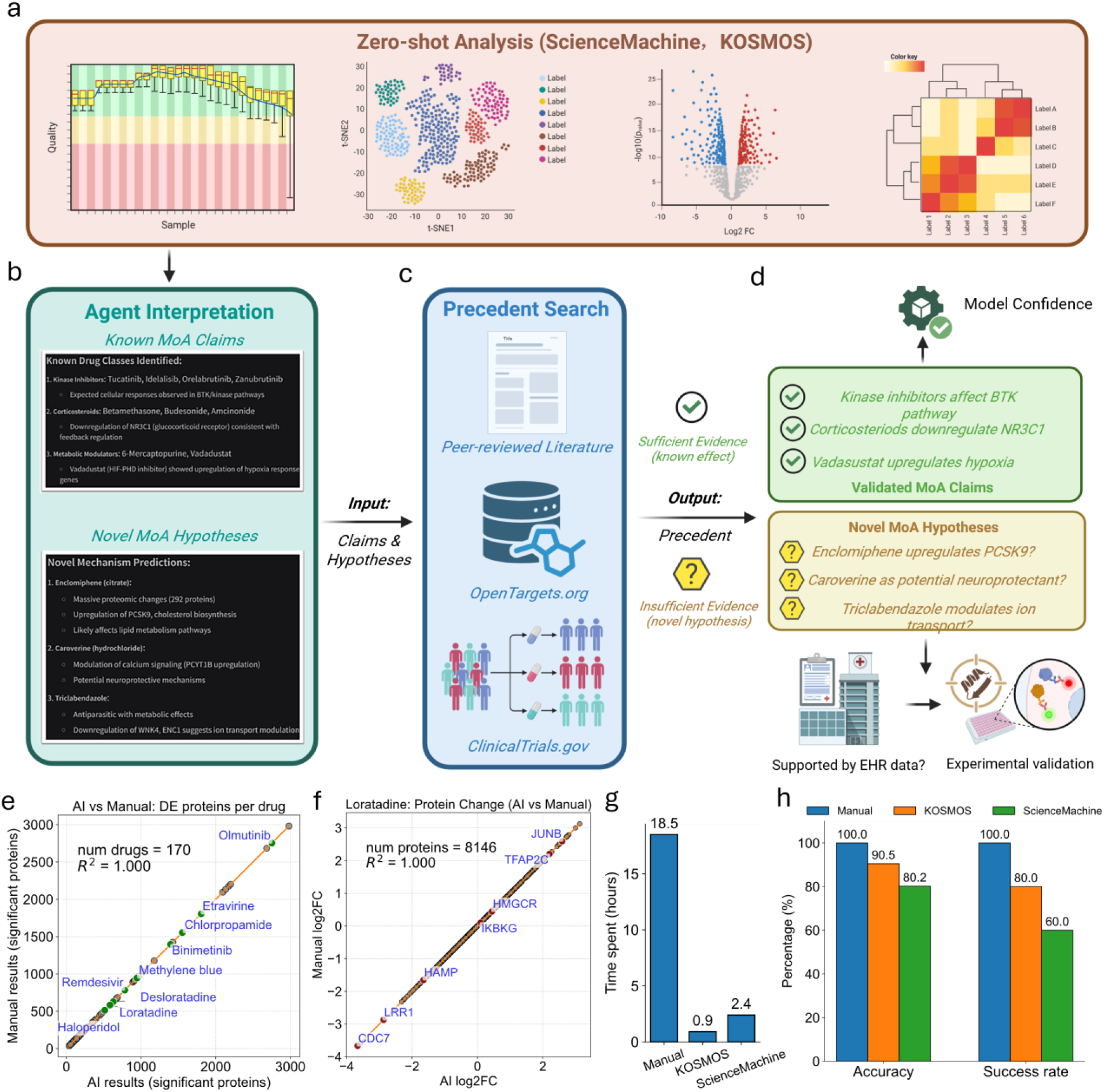
Schematic of the AI-agent analysis framework and concordance with manual results. **a**, AI-agent–assisted data analysis workflow, including zero-shot task instructions, raw data files, sample information and downstream analysis requests. **b**, AI analyzes and organizes the results and then provides the hypotheses. **c**, Rapid comparison and validation of the analysis results and AI-generated hypotheses against existing EHR data. **d**, Rapid validation of results considered novel by the AI. **e**, Comparison of AI-derived and manually derived differential-regulation results for each drug. The criteria for determining significant changes are an adjusted P-value < 0.05 and log2FC > 0.5. **f**, Comparison of AI-derived and manually derived log2FC values for representative drugs. **g**, Comparison of analysis time between manual analysis and AI-based platforms. **h**, Comparison of accuracy and success rate between manual analysis and AI platforms. Data is derived from five independent analyses conducted under an identical framework and set of instructions.

The agent workflow incorporated multiple stages of interpretation. First, the AI performed data cleaning and calibration of the raw proteomic matrix to ensure consistency and comparability across samples. It then extracted gene-level differential expression patterns and organized these into drug-level summaries. Building upon this, the analysis was extended to the pathway level through gene set enrichment analysis (GSEA) using Hallmark and Gene Ontology (GO) biological process gene sets, followed by clustering to identify shared and distinct pathway perturbation patterns across compounds (**Fig. S4**). Based on these multi-layered analyses, the AI generated integrated summaries of drug-induced proteomic responses. Subsequently, the AI separated observations into two categories: claims that were potentially consistent with established mechanisms of action and hypotheses that appeared novel or insufficiently characterized in the literature (**Fig. 2b**). To support rapid verification, these outputs were compared against precedent evidence from curated and external knowledge sources, which has been done with Edison Scientific Precedent search (**Fig. 2c**). Claims supported by sufficient prior evidence were categorized as validated mechanism-related observations, whereas outputs lacking strong precedent were prioritized as candidate novel hypotheses for downstream follow-up and experimental testing (**Fig. 2d**). This design allowed the AI framework to operate not only as an automated interpreter, but also as a triage system for distinguishing established biology from unexpected drug mechanisms.

To assess whether AI-assisted analysis faithfully reproduced conventional expert-driven analysis, we compared AI-derived and manually generated differential-regulation results across the screened compounds. At the drug level, the number of significantly altered proteins identified by the AI was the same as manual analysis across 170 drugs (2 drugs were filtered out due to poor proteome depth and not enough replicates) (**R**^**2**^**= 1.0; Fig. 2e**), indicating that the AI pipeline closely recapitulated human-generated prioritization of proteome perturbation magnitude. Concordance was similarly high at the protein level: for representative compounds such as methylene blue, remdesivir, and digitoxin (**Fig.S5**), AI-derived log2 fold-change estimates were identical to those obtained manually across 8,146 quantified proteins (**R**^**2**^**= 1.0; Fig. 2f**).

In addition to analytical concordance, we further evaluated efficiency and practical usability. Our practical assessment revealed that the transition from raw data processing to completing the t-test dysregulation analysis required more than 18 hours of work by an experienced proteomics data analyst. Importantly, this time estimate does not account for the additional cognitive effort required for biological interpretation and hypothesis generation—tasks that become increasingly formidable as data dimensionality grows, often requiring months or even years. In contrast, the two AI-agent platforms, Science Machine and KOSMOS, completed the entire commanded analytical workflow—including data processing, pathway analysis, and results interpretation—in ∼1 hour and 2.5 hours, respectively (**Fig. 2g**).

Subsequently, we performed five independent runs using an identical analytical framework and instruction set to evaluate the system’s consistency and reliability. The results showed that the AI platform achieved a very high level of accuracy in reproducing expected quantitative analytical outputs (e.g., dysregulation and correlation analyses) and maintained strong consistency across repeated executions (**Fig. 2h**). However, it is noteworthy that the AI performed less consistently in generating final hypotheses, with substantial variability observed across fully independent runs conducted at different time points. Although some conclusions were repeatedly identified, the overall output was not entirely consistent. Importantly, all conclusions were required to be supported by underlying data as evidence, suggesting that these differences do not reflect hallucinations but rather arise from alternative interpretations of the data. Given the high dimensionality of the dataset and the multitude of possible analytical perspectives, such variability is expected. Ultimately, we selected conclusions that were consistently observed across multiple independent reports as candidates for downstream experimental validation.

### AI-driven analysis of drug proteomic perturbations and cellular MoA

Building on this analytical framework, we next asked whether the AI-assisted pipeline could move beyond quantitative analysis to infer biologically meaningful mechanisms of action (MoA) from proteomic data at scale. By integrating protein-level differential regulation, pathway-level interpretation, and structured AI-guided reasoning, the framework generated a unified and interpretable view of both compound-specific and shared cellular responses across the screened library.

At the global level, the AI first quantified the extent of proteomic perturbation induced by each compound, measured as the number of significantly regulated proteins (**adjusted P value < 0.05 and** |**log2FC**| **> 0.5; Fig. 3a, Fig. S6**). This analysis revealed a broad spectrum of drug responses, ranging from compounds that induced extensive proteome-wide remodeling, such as mobocertinib, olmutinib, and clomiphene, to those with relatively modest effects, including cetirizine, pindolol, and diacerein. These differences suggest distinct modes of cellular engagement, from global metabolic and stress-associated remodeling to more selective and targeted perturbations.

**Figure 3.**
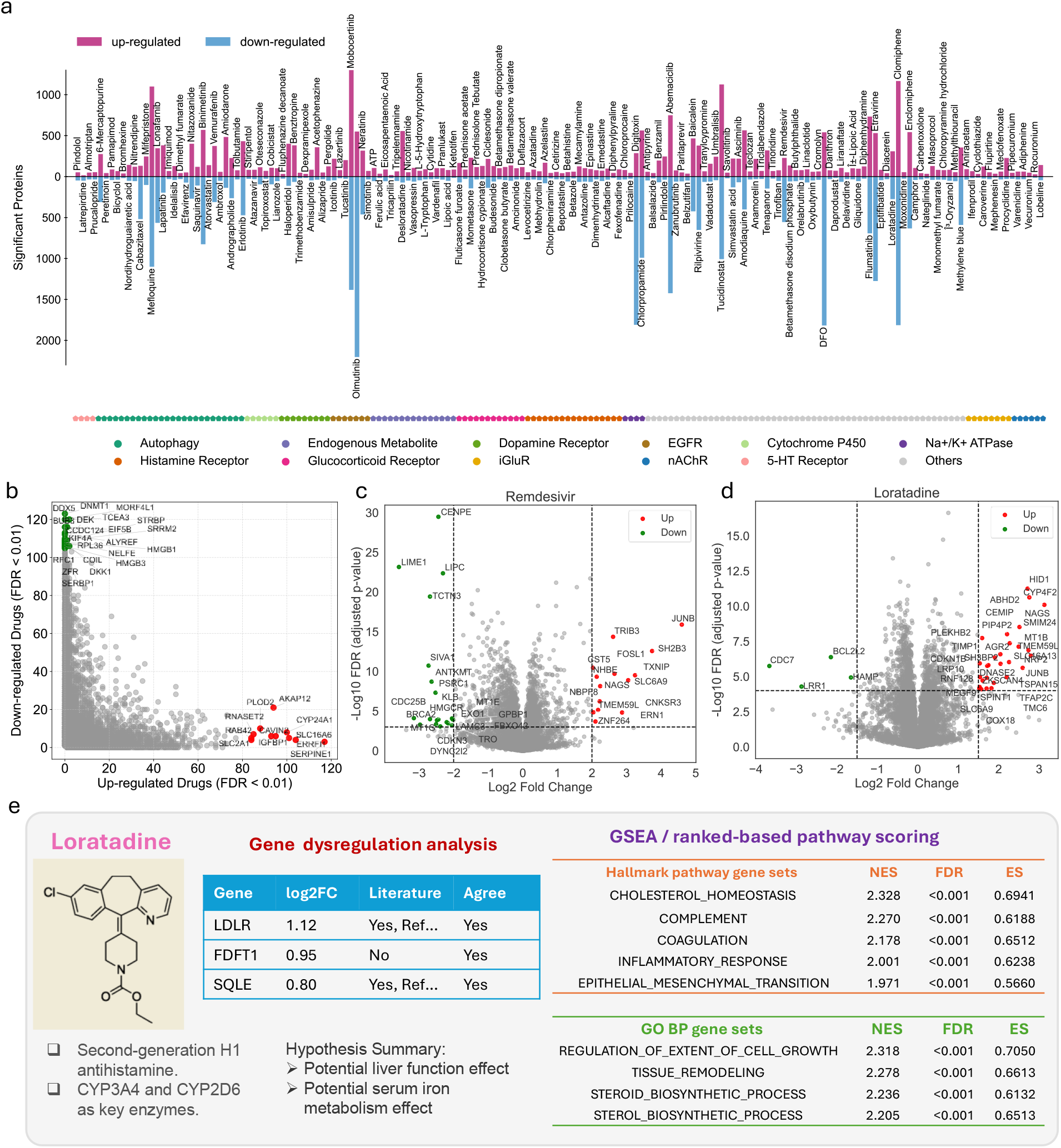
AI-assisted global proteomic perturbation and mechanism-of-action (MOA) analysis. **a**, Bar plot showing the number of significantly up- and downregulated proteins for each of the 170 drugs after data filtered. **b**, Compound-level waterfall plot showing the number of drugs regulating each protein (P < 0.01 and |log2FC| > 0.5). Genes that down-regulated in more than 105 drugs or up-regulated in more than 80 drugs were labeled. **c,d**, Representative volcano plots for selected drugs remdesivir (c) and Loratadine (d). **e**, Example of AI-generated molecular identity profiles for each drug, summarizing basic drug information, significantly dysregulated proteins, perturbed pathways (Hallmark pathways and GO Biological Process ontology gene sets), and potential novel findings.

To identify recurrent responses across compounds, the AI framework next quantified the frequency of significant regulation for each protein across all drugs (**Fig. 3b**). This analysis revealed a recurrently dysregulated protein module characterized by coordinated suppression of core biosynthetic and proliferative programs, including proteins involved in transcriptional regulation, RNA processing, and translation (e.g., DDX5, SRRM2, ALYREF, EIF5B, and RPL36). In contrast, a smaller subset of proteins, including PLOD2, SERPINE1, IGFBP2, CYP24A1, and SLC16A6, was consistently upregulated and enriched in pathways related to extracellular matrix remodeling, metabolic adaptation, and stress responses. Together, these patterns define a shared, system-level proteomic response landscape to pharmacological perturbation.

At the level of individual compounds, the AI framework further interpreted drug-specific proteomic signatures to recover mechanism-consistent observations aligned with prior knowledge. For example, remdesivir exhibited a focused perturbation profile marked by induction of stress-response and redox-regulatory proteins (e.g., SLC7A11, SESN2, and TXNIP),^34^ together with suppression of cell-cycle– associated factors (e.g., CENPE and CDC25B) (**Fig. 3c, Fig.S7**).^35^ Similarly, simvastatin showed strong and coherent activation of cholesterol and sterol biosynthesis pathways, accompanied by upregulation of key enzymes in these processes,^36–38^ consistent with its established mechanism as an HMG-CoA reductase inhibitor (**Fig. S8**). Likewise, haloperidol was associated with coordinated upregulation of cholesterol biosynthesis enzymes (e.g., SQLE, FDFT1, MSMO1, SCD, and LDLR) and enrichment of steroid biosynthesis pathways, suggesting activation of hepatic cholesterol synthesis programs that may underlie its clinically observed risk of dyslipidemia (**Fig. S9**).^39^ These results demonstrate that the AI framework can reliably recapitulate known biological mechanisms directly from high-dimensional proteomic data.

Importantly, beyond recovering established biology, the AI framework also identified observations that were strongly supported by coherent proteomic signatures yet insufficiently explained by existing annotations, thereby prioritizing candidate hypotheses for downstream validation. Notably, these inferences were derived directly from proteomic patterns rather than prior knowledge or literature annotation. For instance, Likewise, loratadine exhibited unexpected enrichment of cell-cycle, p53 signaling, and DNA repair–related pathways, together with marked downregulation of HAMP (hepcidin), a central regulator of systemic iron homeostasis (**Fig. 3d, Fig. S10a**). Moreover, Methylene blue exhibited coordinated suppression of cholesterol biosynthesis enzymes and fibrinogen components, together with induction of hypoxia-responsive proteins, suggesting a previously underappreciated hepatic response involving lipid metabolism and coagulation pathways (**Fig. S10b**). These findings highlight the ability of the AI framework to identify mechanistically coherent, data-driven patterns that extend beyond existing annotations and form the basis for hypothesis generation.

To systematically standardize and organize these multi-layered interpretations across all compounds, we further developed a role-based multi-agent system (MAS) framework with GPT as the backbone model. This architecture decomposed the interpretation process into coordinated roles, including an “Analyzer” for data mining and pattern recognition, a “Reviewer” for validating findings against existing biological knowledge, and a “Supervisor” for integrating outputs and ensuring logical consistency. Through this workflow, the system generated structured molecular identity profiles (**Fig. 3e, Supplementary File4**) and AI-summarized interpretations for each compound (**Fig.S11, Supplementary File5**). The molecular identity profiles integrate drug annotations, top dysregulated proteins, and pathway-level perturbations derived from pre-ranked gene set enrichment analysis using Hallmark and GO Biological Process gene sets. For the AI summaries, the MAS generates structured drug interpretation reports that integrate data-driven signatures, knowledge-based validation, and prioritized hypothesis generation, with explicit assessment of confidence and limitations. In this way, the AI framework transforms digitized, high-dimensional proteomic data that are difficult for humans to interpret into reports that are intuitive, interpretable, and hypothesis-oriented.

To further improve accessibility and enable interactive exploration of these results, we developed a publicly available web-based data portal (https://drug-ai-robotics-proteomics.streamlit.app/; the server may need to be reactivated upon first access) that extends these molecular identity profiles into a searchable and visual interface. The portal allows users to query individual AI-generated summaries, visualize differential protein regulation, and explore pathway-level perturbations across the screened library in HepG2 cells. In addition to static summaries, it supports interactive volcano plots, protein-centric views, cross-compound comparisons and data download, thereby converting the dataset from a static resource into an exploratory platform for AI-assisted biological interpretation and hypothesis generation.

### Groups of drugs with similar cellular mechanisms

In large-scale proteomics-based drug screening, mechanism-of-action (MoA) inference typically progresses from single-protein or single-drug perturbations to similarity-based clustering and ultimately pathway-level integration. Here, we recapitulated this analytical framework using an AI-led approach. The AI first constructed a drug–drug similarity network based on pairwise correlations of whole-proteome responses (log2FC across all quantified proteins). Both Pearson and Spearman analyses revealed structured relationships, with subsets of compounds forming tightly connected communities (**Fig. 4a and Fig. S12**). Restricting to strong correlations (Pearson r > 0.5 or Spearman r > 0.6) revealed clear clusters, indicating convergent proteomic remodeling among drugs with similar mechanisms. Notably, chemically distinct compounds often exhibited highly similar proteomic signatures, suggesting shared downstream effects or convergence on common pathways (**Fig. 4b, c**). In parallel, protein-level cross-drug correlation analysis revealed coordinated co-regulation among subsets of proteins. The AI further identified groups of proteins with shared pathway responses and potential cross-pathway coordination, indicating higher-order organization of proteomic responses (**Fig. S13, Supplementary Table 6**).

**Figure 4.**
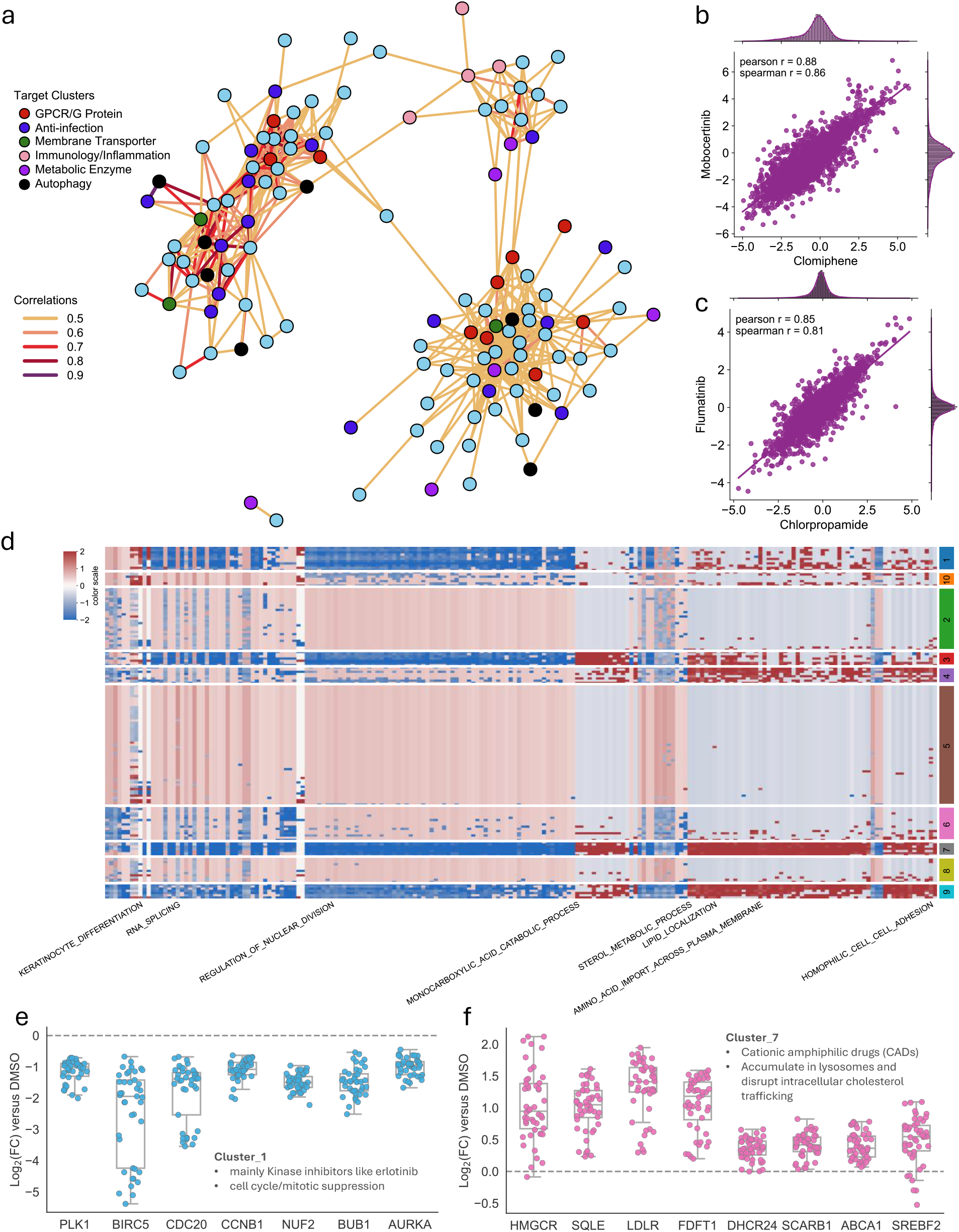
Groups of drugs with similar cellular MoA. **a**, Community plot built from a drug–drug correlation matrix of whole proteome. Typical drug clusters were labeled with different color. Pearson correlations are applied and filtered to only include edges with pearson r > 0.5. **b,c**, Pairwise correlation plot of two typical drug pairs: Clomiphene and Mobocertinib (b), Chlorpropamide and Flumatinib (b). **d**, Drugs were grouped into 10 clusters using fuzzy c-means clustering based on GO biological process–level pathway dysregulation profiles derived from GSEA normalized enrichment scores (NES). The heatmap displays standardized NES values, and for each cluster, the top positively and negatively enriched pathways (based on NES) are highlighted as representative functional signatures. **e**, Cluster 1 contains mainly kinase inhibitors (abemaciclib, binimetinib, erlotinib, neratinib etc.), suppressing genes (PLK1, BIRC5, CDC20, BUB1 etc.) in cell cycle/mitotic programs. **f**, Cluster 7 contains mainly cationic amphiphilic drugs (amiodarone, haloperidol, and amodiaquine etc.), perturbing sterol metabolism genes (HMGCR, SQLE, LDLR, FDFT1.etc).

To improve robustness and interpretability, we performed pathway-level clustering, which reduces noise from high-dimensional protein data and captures coordinated functional responses. Specifically, fuzzy c-means clustering was applied to GSEA-derived normalized enrichment scores (NES) based on GO Biological Process terms (**Supplementary Table 7**), complemented by hierarchical clustering of Hallmark gene sets (**Fig. S14**). These approaches grouped compounds by functional pathway dysregulation rather than individual protein changes. This analysis identified 10 clusters with coherent pathway signatures (**Fig. 4d, Fig. S15**). For example, Cluster 1 was enriched for kinase inhibitors (e.g., abemaciclib, binimetinib, erlotinib, neratinib) and showed consistent downregulation of cell cycle regulators (PLK1, BIRC5, CDC20, BUB1), reflecting anti-proliferative effects (**Fig. 4e**). In contrast, Cluster 7, dominated by cationic amphiphilic drugs (e.g., amiodarone, haloperidol, amodiaquine), exhibited coordinated perturbation of sterol metabolism and lipid homeostasis pathways, involving genes such as HMGCR, SQLE, LDLR, and FDFT1 (**Fig. 4f**).

As expected, not all clusters corresponded to well-characterized drug classes. Some contained chemically diverse compounds with shared but less understood proteomic signatures, suggesting convergence on incompletely characterized cellular processes. These clusters provide opportunities for further investigation. Collectively, these results demonstrate that the AI framework, under guided analytical design, can systematically perform large-scale proteomic clustering and pathway-level interpretation, enabling identification of drug groups with shared cellular mechanisms and providing a data-driven foundation for MoA inference, drug repurposing, and discovery of novel biological relationships.

### Validation of AI-prioritized novel findings

Once the AI framework completed the full analytical workflow, we next evaluated its ability to prioritize biologically meaningful and experimentally testable hypotheses from high-dimensional proteomic data. Within the same analytical prompt, the AI was instructed to identify candidate observations that were strongly supported by coherent proteomic signatures yet insufficiently explained by existing annotations or prior knowledge. From these AI-prioritized candidates, we selected representative hypotheses for orthogonal validation, focusing on mechanistically consistent patterns that lack strong precedent and are therefore suitable for testing AI-driven biological insight generation.

As a first case study, the AI framework highlighted methylene blue as a compound associated with coordinated suppression of hepatic cholesterol and lipid metabolic pathways (**Fig. 5a,b**). This inference was derived directly from the underlying proteomic data, rather than from prior knowledge or literature annotation. Specifically, the proteomic profile showed consistent downregulation of multiple key enzymes involved in cholesterol biosynthesis, including HMGCR, SQLE, FDFT1, and SCD (**Fig.S16**), together with broader proteomic changes indicative of altered hepatic metabolic function. These patterns define a coherent mechanistic signature linking suppression of sterol biosynthesis pathways to reduced lipid metabolic activity. Notably, this effect is not well characterized in existing literature. Based on this evidence, the AI framework nominated the hypothesis that methylene blue may exert a previously unrecognized cholesterol-lowering effect.

**Figure 5.**
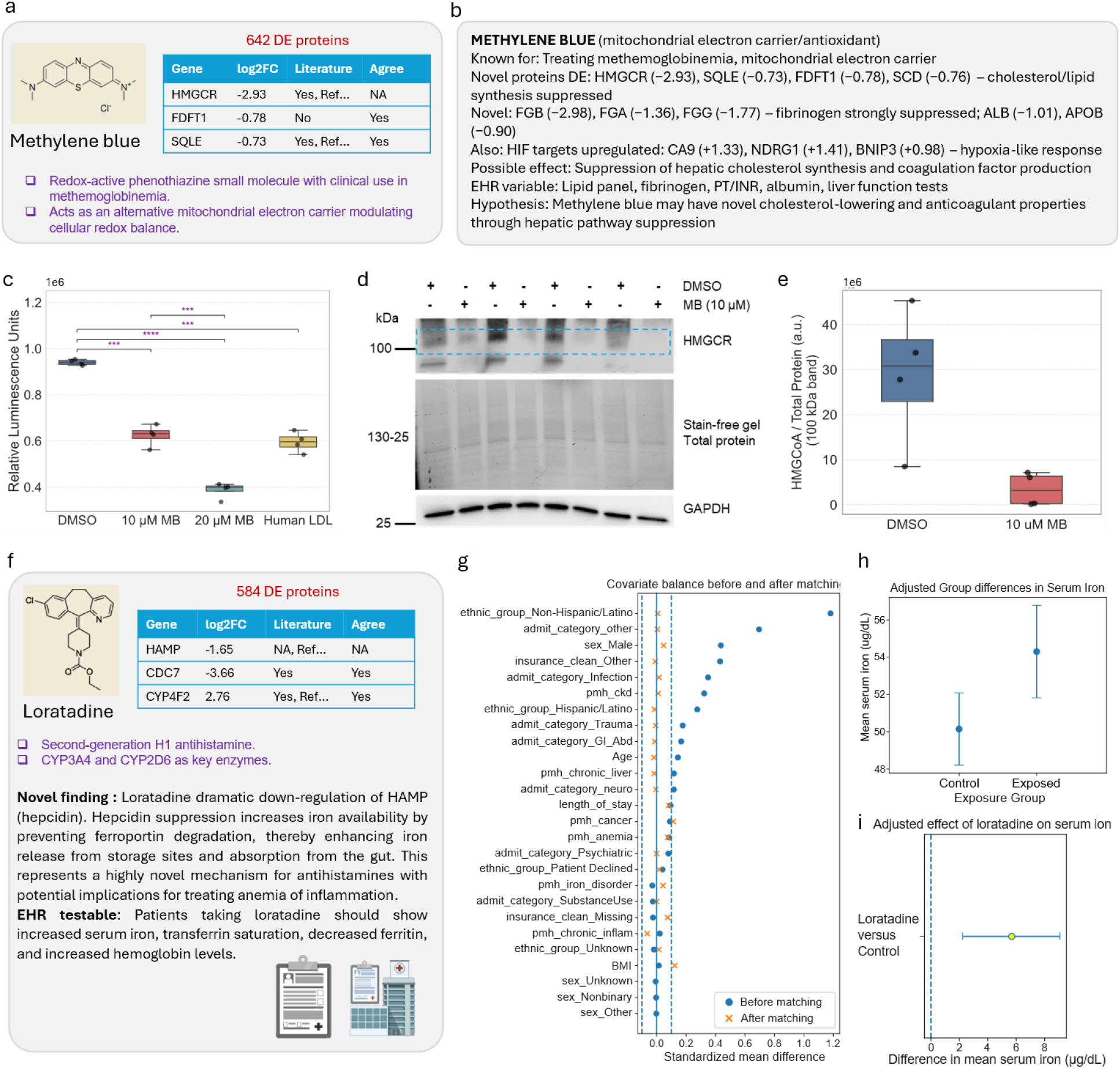
Validation of AI-proposed novel findings. **a**, A novel hypothesis generated by the AI based on the drug screening proteome dataset: Methylene Blue. **b**, Evidence supporting the AI agent’s conclusion derived from the experimental proteomic analysis results provided. **c**, Experimental validation demonstrates that the HepG2 cells treated with Methylene Blue significantly reduces cholesterol levels, confirming the AI-generated hypothesis. Free cholesterol levels were quantified using a luminescent assay, and levels are reported as Relative Luminescence Units (RLU). **d, e**, Western blot results showing a significant reduction in HMGCR levels upon Methylene Blue treatment (n=4 technical replicates). **f**, A novel AI-generated hypothesis from the drug screening proteomics dataset suggests that loratadine may reduce circulating free iron levels, supported by consistent evidence from experimental proteomic analysis. **g**, Covariate balance before and after matching from the analysis of real-world Electronic Health Record (EHR) data. **h**, Mean serum iron (ug/dL) between Loratadine-exposed group and the control group, error bars indicate the 95% CI for each group. **i**, Adjusted difference in serum iron (ug/dL) associated with Loratadine exposure, with positive values indicating higher serum iron in the Loratadine-exposued group vs. control.

Experimental follow-up supported this prediction. Treatment with methylene blue significantly reduced free cholesterol levels in HepG2 cells showing a dose-dependent effect relative to control conditions (**Fig. 5c**), consistent with the directionality inferred from the proteomic data. In agreement with this mechanism, immunoblot analysis, together with independently replicated mass spectrometry data from an additional methylene blue treatment, confirmed reduced HMGCR abundance following methylene blue exposure (**Fig. 5d,e; Fig. S17**), thereby validating suppression of a key rate-limiting enzyme in cholesterol biosynthesis. Together, these results provide orthogonal validation of the AI-derived hypothesis and demonstrate that proteome-level signatures can be translated into experimentally testable predictions.

We next examined a second AI-prioritized hypothesis centered on loratadine (**Fig. 5f**). In this case, the AI pointed a pronounced downregulation of hepcidin (HAMP) in the proteomic profile, a master regulator of systemic iron homeostasis. Given the central role of hepcidin in controlling iron export through regulation of ferroportin, its suppression is expected to increase circulating iron availability in humans. To evaluate this prediction in an independent real-world setting, we analyzed EHR data from Cedars-Sinai Medical Center spanning 2015-2025, using a matched comparison framework. Covariate balance between exposed and control groups was substantially improved after matching (**Fig. 5g**), supporting the validity of downstream comparisons. In the matched analysis, mean serum iron levels were higher among Loratadine-exposed patients compared to controls by ∼4 ug/dL (**Fig. 5h**) and after adjusting for clinical covariates using a regression model, Loratadine exposure was associated with an increase in serum iron by ∼6 ug/dL (**Fig. 5i**), consistent with the directionality predicted from the proteomic data. Although this observational analysis does not establish causality, it provides complementary real-world evidence supporting the hypothesis generated by the AI-assisted proteomic interpretation pipeline. Our AI-prioritized and validated hypotheses involving cholesterol biosynthesis and iron homeostasis pathways support HepG2 as a good cell model for identifying compounds that can modify hepatic function with potential therapeutic implications.

Together, these case studies establish a complete data-to-hypothesis-to-validation loop, demonstrating that AI-assisted analysis of large-scale proteomic screening data can move beyond descriptive interpretation to enable predictive and experimentally testable biological discovery. By systematically prioritizing mechanistically coherent yet underexplored patterns, this framework provides a scalable approach for translating high-dimensional proteomic data into actionable biological insights.

## Discussion

In this study, we present an end-to-end proteomics framework that integrates semi-automated experimental workflows with AI-led analytical interpretation, enabling scalable generation and interpretation of drug-induced proteomic data. By combining high-throughput sample preparation, rapid mass spectrometry acquisition, standardized data processing, and an AI-agent–driven analytical layer, this platform addresses two major bottlenecks in large-scale proteomics: data generation and data interpretation.

A key advance of this work is the simultaneous improvement in the efficiency of both data acquisition and knowledge production. Although recent technological advances have substantially increased the throughput and depth of proteomic measurements, the rapid, scalable generation of proteomic data—and the extraction of biologically meaningful insights from high-dimensional datasets—remain central challenges.^40–42^ Our results demonstrate that AI-led analysis can organize these complex outputs into structured, interpretable summaries, faithfully recapitulate established mechanisms of action, and prioritize hypotheses for further validation, along with proposed validation strategies.

We show that, when guided by clearly defined analytical logic and structured inputs, AI systems can reliably reproduce standard proteomics workflows, including differential analysis, pathway enrichment, and higher-order organization of drug responses. At the same time, the AI framework substantially reduces the time and effort required for downstream interpretation. In our experience, manual analysis of a dataset of this scale requires extensive coding, iterative validation, and domain-specific reasoning, often taking days of focused work for basic result generation alone, without including hypothesis exploration. In contrast, the AI systems were able to execute the full analytical workflow and generate structured biological interpretations within hours. These findings suggest that the primary bottleneck in large-scale proteomics is shifting from data acquisition to earlier and later in the workflow: at the beginning, defining clear scientific questions and systems, and at the end, validation of hypotheses.^43^

Beyond reproducing known biology, the framework’s important strength is its ability to highlight biologically coherent yet currently unknown patterns. By systematically comparing observed proteomic signatures with existing knowledge bases, the AI system can distinguish between well-supported mechanism-related observations and candidate hypotheses lacking strong precedent. This enables a form of computational triage, directing attention toward findings that are both biologically plausible and potentially novel. The subsequent validation of AI-prioritized hypotheses, including biochemical assays and real-world EHR analyses, demonstrates that such AI-generated insights can translate into experimentally and clinically relevant discoveries.^44,45^ At the systems level, our analysis reveals that drug-induced proteomic responses exhibit both shared and compound-specific features. Proteome-wide similarity analysis and pathway-level clustering identified groups of compounds with convergent cellular effects, including well-characterized classes such as kinase inhibitors and cationic amphiphilic drugs, as well as less-defined clusters that may represent underexplored biological mechanisms. These results highlight that proteomic responses capture functional organization beyond canonical target annotations, and further demonstrate that AI-led frameworks can recover this higher-order biological structure in a systematic and scalable manner.^46^

Despite these advances, several limitations should be considered. First, the performance of AI-led analysis depends critically on the clarity of task definition and the quality of input data. Ambiguity in analytical objectives or poorly specified instructions can introduce variability, particularly in higher-level interpretation and hypothesis generation. Second, although AI systems can rapidly generate biologically plausible hypotheses, these predictions require careful experimental and clinical validation, as demonstrated in this study. Third, current AI approaches remain limited in generating publication-ready visualizations that meet the standards of scientific communication, and human aesthetic Intuition remains essential for figure design and presentation.

More broadly, our results point to a shift in the utility of large-scale perturbation proteomics and in how large-scale biological datasets are interrogated. For complex datasets, there is rarely a single correct interpretation; instead, multiple analytical perspectives can yield complementary insights. In this context, AI systems provide a powerful means to systematically explore this interpretative space, rapidly generating and evaluating alternative hypotheses. As a result, the role of researchers may increasingly shift from performing analyses to defining analytical frameworks, guiding the exploration of data-driven hypotheses, and leveraging intuition to identify meaningful results. This trend suggests that researchers will increasingly be distinguished by their ability to clearly define scientific questions, and, to some extent, it places higher demands on innovation.

In conclusion, this work establishes a scalable sample-to-insight platform for proteomics-driven drug screening, demonstrating that the integration of automation and AI can transform both the generation and interpretation of high-dimensional biological data. More importantly, it suggests a broader paradigm in which clearly defined analytical frameworks, when coupled with AI systems, enable rapid, iterative hypothesis generation and validation, thereby accelerating the discovery cycle in biomedicine.

## Methods

### Drugs and Materials

The FDA-approved drug library was obtained from MedChemExpress LLC and supplied in 384-well plate format HeLa protein digest standards (88328), ulbecco’s Modified Eagle Medium ( MEM; 11995-065), phosphate-buffered saline (PBS; 14190144), dialyzed fetal bovine serum (FBS; A3382001), penicillin– streptomycin (15-140-122), radioimmunoprecipitation assay (RIPA) buffer (AAJ62524AE), ammonium bicarbonate (AC393210010), 0.25% trypsin–EDTA (25-200-056), 1 M Tris-HCl (pH 8; 15568025), and LC– MS-grade reagents including formic acid (A117–50) and 0.1% formic acid in water (LS118–1) were purchased from Thermo Fisher Scientific (Waltham, MA). Additional reagents, including protease inhibitor cocktail tablets (11836153001), HPLC-grade ethanol (59828-1L), anhydrous calcium chloride (C5670-100G), HPLC-grade acetonitrile (34851), Pierce HeLa protein digest standards (88329), dithiothreitol (DTT), and iodoacetamide (IAA), were obtained from MilliporeSigma (Burlington, MA). Sequencing-grade modified trypsin (V5113) was purchased from Promega, and silica magnetic beads (786-916) were obtained from G-Biosciences (St. Louis, MO).

### Cell treating and Sample preparation

HepG2 cells were cultured in 15 cm dishes in high-glucose DMEM supplemented with 10% fetal bovine serum (FBS) and antibiotics (100 U/mL penicillin and 100 μg/mL streptomycin) ells were subsequently transferred to 96-well plates and subjected to drug treatment for 24 h. Compounds were dispensed into 96-well plates using an Echo 50 Series liquid handler, with all drugs applied at a final concentration of 10 μM Following 24 h incubation, culture media were removed, and cells were washed twice with PBS. Cells were lysed in RIPA buffer supplemented with protease inhibitor cocktail (1 tablet per 10 mL) and vortexed for 5 min. All samples were then processed using an automated solid-phase enhanced sample preparation (SP3) protocol (17) on a KingFisher Flex system (Thermo Fisher Scientific). Proteins were reduced with dithiothreitol (DTT) for 30 min at room temperature, followed by alkylation with iodoacetamide (IAA) for 30 min at room temperature in the dark. Paramagnetic silica beads were added at a protein-to-bead ratio of 1:10 (w/w), and beads were pre-washed with Milli-Q water using the KingFisher Flex prior to sample addition. Acetonitrile (LC–MS grade) was then added to a final concentration of 70% (v/v), and samples were incubated at room temperature for 18 min with gentle agitation to induce protein binding. Bead-bound proteins were washed twice with 500 μL of 80% ethanol, followed by two washes with 500 μL of 100% acetonitrile. The beads were then resuspended in 50 mM Tris-HCl containing 10 mM CaCl_2_and digested with trypsin at a protein-to-enzyme ratio of 25:1 (w/w) at 37 °C overnight with gentle agitation. The following day, beads were removed, and digestion was quenched with formic acid. Samples were centrifuged and transferred to a new 96-well plate prior to mass spectrometry analysis.

### Mass spectrometry

Chromatographic separation was performed on a Vanquish Neo UHPLC system (Thermo Fisher Scientific) equipped with a epSep 18 column (15 cm × 150 μm, 1 5 μm particle size; Bruker) maintained at 55 °C. A 15 min linear gradient was applied using solvent A (0.1% formic acid in water) and solvent B (80% acetonitrile with 0.1% formic acid). The gradient was programmed as follows: 4% B at 0 min, increased to 9% B at 2 min, 25% B at 10 min, and 35% B at 13 min, followed by column washing and re-equilibration. The flow rate was set to 1 2 μL/min with a pump curve value of 5 MS analysis was performed on a Thermo Orbitrap Astral instrument operated in data-independent acquisition (DIA) mode. MS1 scans were acquired in the Orbitrap at a resolution of 240,000, with DIA isolation windows of 4 m/z covering a mass range of 380–980 m/z.

### Data analysis

Data analysis was primarily conducted using an AI-agent–guided workflow with human supervision and correction. The agent was directed by a rigorous and highly detailed prompt that specified the required analytical steps, quality-control criteria, and output formats, while also encouraging iterative evaluation of result quality and the inclusion of complementary analyses when necessary. At a minimum, the workflow included three stages. First, for data cleaning, detected proteins and missing-value rates were assessed; proteins detected in fewer than one-third of samples were removed, samples with fewer detected proteins than 80% of the cohort average were excluded, and drugs with fewer than three remaining replicates were not analyzed further. Second, missing values were imputed using shifted minimum per protein and batch effects were corrected on the filtered dataset after log2 transformation. ComBat was applied for batch correction, and UMAP visualizations were generated before and after correction, with samples annotated by control/QC/drug class and batch. Corrected data were used for all downstream analyses. Third, dysregulation analysis was performed using Welch’s t-test for each drug versus DMSO on the corrected dataset. P values were adjusted within each drug using the Benjamini–Hochberg procedure, and proteins were considered differentially regulated when the false discovery rate was <0.05 and the absolute log2 fold change was >0.5. A summary table containing dysregulation statistics for all proteins across all drugs was generated for downstream interpretation.

### Experimental validation

#### Cholesterol measurement by luminescence assay

Approximately 2.5 × 10^4 HepG2 cells were seeded into individual wells of a 96-well plate. After 48 h (∼60% confluence), cells were treated with either DMSO or Methylene Blue (10 or 20 µM) for an additional 24 h. Treatments were applied in culture medium supplemented with 10% FBS. Cellular cholesterol was quantified using the Cholesterol/Cholesterol Ester-Glo™ luminescent assay ( romega) according to the manufacturer’s instructions Briefly, after treatment, the culture medium was removed, and the cells were washed twice with BS 50 μL of the kit’s lysis buffer was added to the wells containing cells, and the plate was incubated at 37 ° for 30 minutes 50 μL of the cell lysates were transferred into individual wells of a white-walled 96-well plate (assay plate). The assay plate also contained 50 μL of cholesterol standard dilutions prepared in lysis solution (ranging from 0 to 80 µM) Then 50 μL of the cholesterol detection reagent was added to the tested wells in the assay plate, and samples were incubated for 1 hour at room temperature on a plate shaker. The detection reagent is enzymatically coupled to cholesterol to generate NADH, and luciferase activates a pro-luciferin to produce light. Luminescence was recorded using a BioTek Cytation 1 cell imaging multi-mode reader (Agilent) with an integration time of 1.0 s. Data were recorded as raw relative luminescence units (RLUs). As a positive control for cholesterol content, purified human low-density lipoprotein (LDL, 6.53mg/mL) was used and diluted in 50 µL the kit’s lysis buffer (1:500 dilution).

#### Western blot analysis

HepG2 cells were seeded, cultured, and treated similarly and in parallel with the cholesterol assay Following treatment of MSO or Methylene Blue (10 μM) for 2 h, cells were washed with ice-cold PBS and lysed on ice in RIPA buffer containing protease inhibitors (Roche). Micro BCA protein kit (Thermo Scientific) was used on cell supernatants to determine protein concentration (Pierce). Samples were prepared in Laemmli S S sample buffer containing β-mercaptoethanol reducing agent and heated at 95°C for 5 min prior to electrophoresis. Approximately 20 µg of protein per lane were resolved on 4-15% TGX stain-free gels (Bio-Rad) and transferred to V F membranes (0 22 μm) using Trans-Blot Turbo transfer system (25V/3min) (Bio-Rad). Total protein per lane was visualized using a stain-free imaging system to confirm uniform loading and enable normalization of total protein (ChemiDoc MP, Bio-Rad). Membranes were blocked in 5% nonfat milk in TBS-T (Tris-buffered saline, 0.1% Tween-20) for 1 h at room temperature and incubated with primary antibodies against HMG-CoA reductase (HMGCR; NovusBio, CL0259, 1:1000 dilution) and HRP anti-GAPDH (BioLegend, FF26A, 1:5000 dilution) overnight at 4°C. To detect HMGCR, after washing with TBS-T, membranes were incubated with HRP-conjugated secondary antibody for 1 h at room temperature (5450-0011, SeraCare). Bands were detected using Amersham ECL Select™ Western Blotting detection reagent (Cytiva, RPN2235) and imaged with a ChemiDoc MP imaging system (Bio-Rad). Band intensities were quantified using Image Lab software (v 6.1). HMGCR signal was normalized to total protein per lane (stain-free). Normalized values are reported in arbitrary units (a.u.).

#### Gene set enrichment analysis (GSEA)

To quantify pathway-level perturbations induced by each compound, we performed preranked gene set enrichment analysis (GSEA) using protein-level differential abundance estimates. For each drug, proteins were mapped to gene symbols and ranked according to their log2 fold change (log2FC) relative to DMSO. Enrichment analysis was conducted using the preranked GSEA algorithm implemented in the *gseapy* package. Gene sets were obtained from the Molecular Signatures Database (MSigDB), including the Hallmark collection and Gene Ontology (GO) Biological Process terms. Only gene sets containing between 20 and 300 genes were retained to reduce noise from overly small or broad categories.

For a given gene set, an enrichment score (ES) was calculated by traversing the ranked gene list and computing a running-sum statistic. At each position in the ranked list, the running sum was increased when a gene belonged to the gene set and decreased otherwise. The magnitude of the increment for genes within the set was proportional to the absolute value of the ranking metric (|log2FC|), thereby assigning greater weight to genes with stronger differential abundance. The ES was defined as the maximum deviation from zero encountered during this traversal, reflecting the degree to which genes from the set were concentrated at the top or bottom of the ranked list. To account for differences in gene set size and score distribution, the ES was normalized to yield a normalized enrichment score (NES). This was achieved by comparing the observed ES to a null distribution generated through permutation of the ranked gene labels. Specifically, for each gene set, the ranked list was randomly permuted multiple times (typically 100–1000 permutations), and ES values were recalculated to estimate the expected distribution under the null hypothesis of no enrichment. The observed ES was then divided by the mean of the corresponding null distribution (separately for positive and negative scores) to obtain the NES.

Statistical significance was assessed using empirical p-values derived from the permutation-based null distributions, followed by multiple hypothesis correction to control the false discovery rate (FDR). Positive NES values indicate that genes within a given pathway are preferentially enriched among upregulated proteins, whereas negative NES values indicate enrichment among downregulated proteins. The resulting pathway-level NES matrix (pathway × drug) was subsequently used for clustering, similarity analysis, and functional interpretation of drug-induced proteomic responses.

## Data Availability

The raw data from mass spectrometry are openly available from massive.ucsd.edu with reference number MSV000101671, FTP download link ftp://massive-ftp.ucsd.edu/v12/MSV000101671/. Interactive data and per drug summary can be check and downloaded in website https://drug-ai-robotics-proteomics.streamlit.app/

## Code Availability

Python version 3.13.9 was used and all code generated from AI agent and produced for manual check are provided open source via github-xomicsdatascience, https://github.com/xomicsdatascience/AI-robotics-proteomics. Versions and packages can be found in the requirements.txt file in the openly available link in Massive. Note that the manual check process does not imply that all functions and code were written line by line by humans. Some parts were assisted by AI; however, the overall logic and reliability of all functions were manually verified to ensure they meet expectations.

## Acknowledgements

This work was funded in part by the National Institute of General Medical Sciences (NIGMS R35GM142502).

## Notes

### Competing Interest Statement

The authors have declared no competing interest.

