## Supplementary Text and Figures for "Robotic perturbation proteomics and AI agents enable scalable drug mechanism discovery"

### for

#### Methods

##### Protein clustering

To improve robustness and interpretability, proteins were filtered prior to clustering based on data completeness, variability, and effect size. Proteins were retained if they satisfied the following criteria: (i) non-missing values in at least 70% of drug conditions, (ii) standard deviation across drug treatments $\geqslant 0.2$, and (iii) maximum absolute log2 fold-change $\geqslant 0.5$ across all drugs. Proteins failing any of these criteria were excluded from downstream analysis. Missing values were imputed using row-wise mean substitution, and protein response profiles were optionally standardized using row-wise z-score transformation. Pairwise distances between proteins were computed using correlation-based distance (1 − Pearson correlation), and hierarchical clustering was performed using average linkage. Cluster assignments were determined by cutting the dendrogram at a predefined number of clusters.

##### GO pathway–based fuzzy c-means clustering.

To classify compounds according to pathway-level response patterns, fuzzy c-means (FCM) clustering was performed on Gene Ontology Biological Process (GO BP) enrichment results derived from preranked gene set enrichment analysis (GSEA). GO terms with a false discovery rate (FDR) greater than 0.05 were excluded, and only pathways significantly enriched in at least three drugs were retained. Normalized enrichment scores (NES) were then organized into a drug-by-pathway matrix, with rows representing drugs and columns representing GO BP terms. Missing NES values were imputed as 0, indicating no detectable enrichment signal for a given drug–pathway pair. To reduce dimensionality and focus on the most informative pathway features, only the top 200 GO terms with the highest variance across drugs were retained.

Before clustering, the NES matrix was standardized across pathways using z-score scaling. Principal component analysis (PCA) was then applied, and the first 20 principal components were used as input for fuzzy c-means clustering. FCM was performed with 10 clusters and a fuzziness parameter of 1.7, using a fixed random seed of 0 for reproducibility. This approach assigned each drug a graded membership score across clusters rather than a single hard label. For downstream interpretation, each drug was assigned to its dominant cluster based on the highest membership value, and cluster-level pathway profiles were summarized by averaging NES values across drugs within each dominant cluster.

**Supplementary Figures**

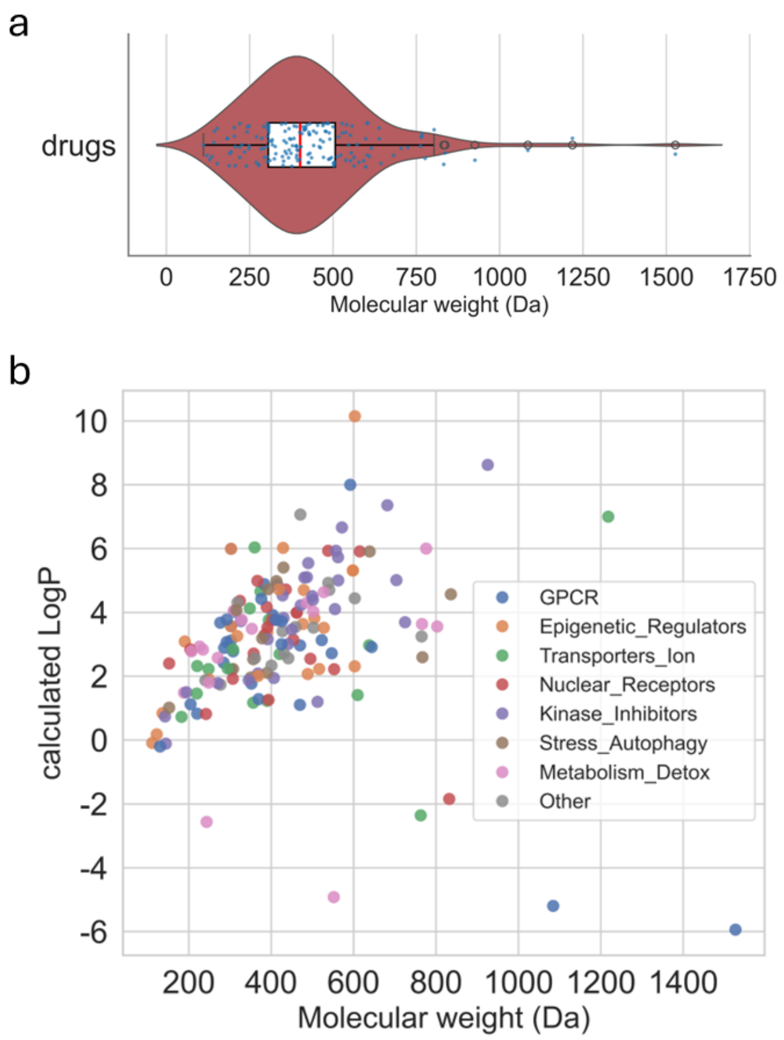


**Fig. S1 | Physicochemical landscape of the screened compound library. a,** Distribution of molecular weights for all screened compounds. **b,** Scatter plot of calculated logP (cLogP) versus molecular weight, illustrating the physicochemical diversity of the screened compound library across annotated classes.

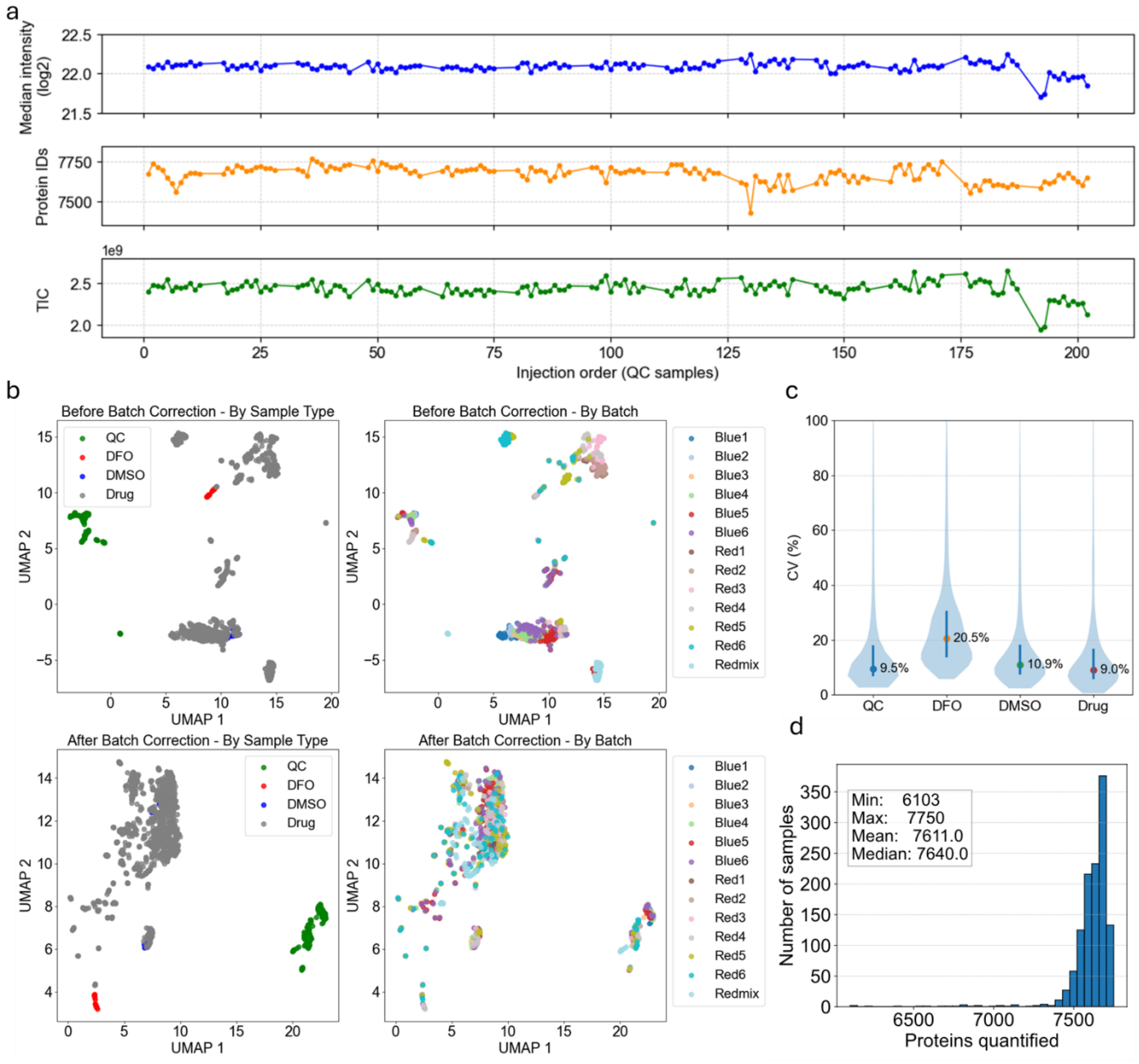


**Fig.S2 | Quality assessment of the proteomics dataset from the drug screening study. a,** Performance drift of QC samples across the entire acquisition process, including median signal intensity (top), number of identified proteins (middle), and total ion count (bottom). **b,** UMAP projections showing batch structure before and after batch correction. Projections colored by sample type are shown before (upper left) and after (lower left) correction, whereas projections colored by injection plate are shown before (upper right) and after (lower right) correction. Blue, red, and redmix represent the sample plate identifiers, while numbers 1–6 indicate the six biological replicates for each drug. **c,** Distribution of overall coefficients of variation across different sample types. **d,** Histogram showing the distribution of the number of identified proteins at the individual sample level.

a

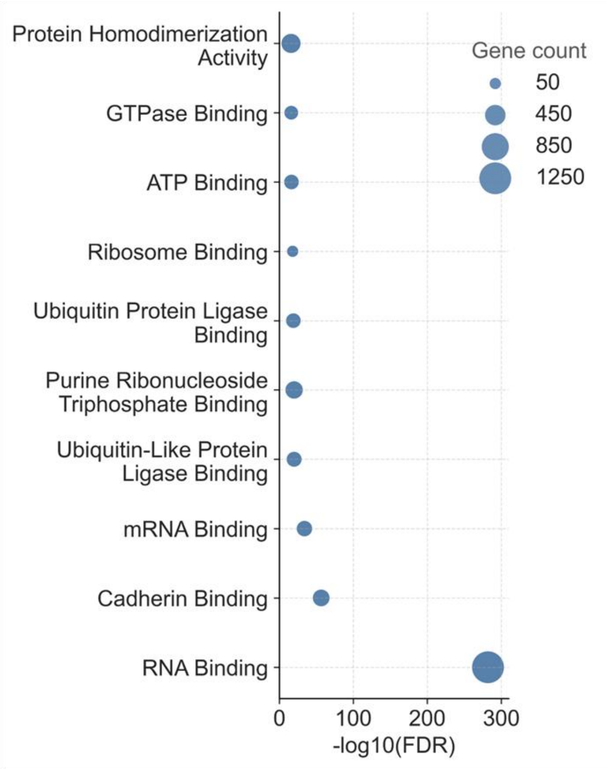


b

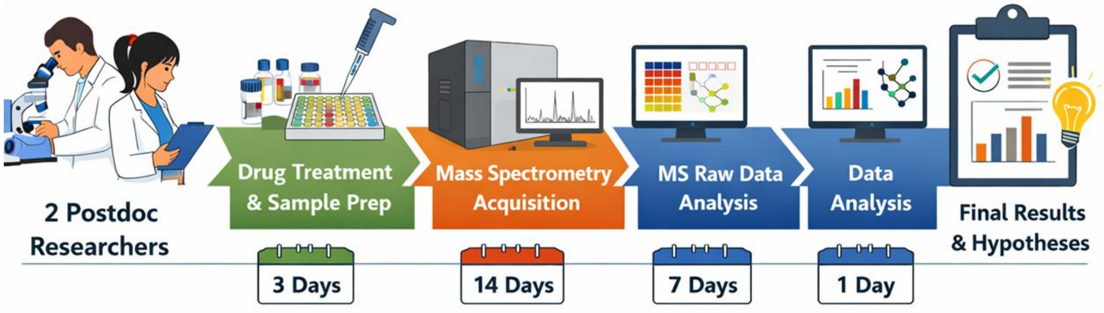


**Fig.S3 | Timeline of the semi-automated screening workflow and representative enrichment analysis. a,** Representative Gene Ontology molecular function (GO:MF) enrichment analysis shown as a bubble plot, where bubble size indicates gene count and the x axis represents enrichment significance as −log10(FDR). **b,** Time allocation across the major steps of the workflow under a two-postdoc operating setting, including 3 days for drug treatment and sample preparation, 14 days for mass spectrometry acquisition, and 1 day for data analysis, culminating in final results and hypothesis generation within 18 days.

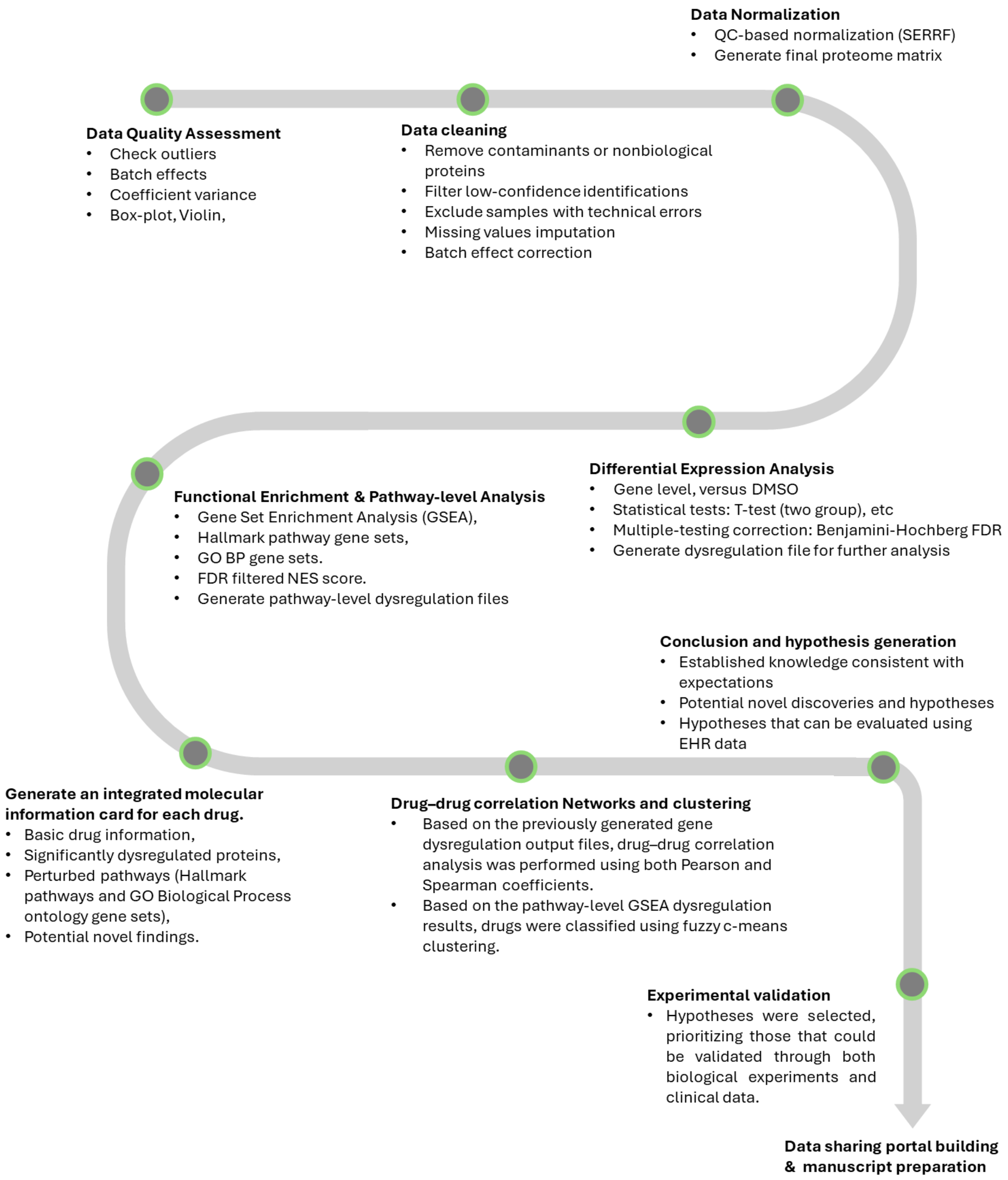


**Fig.S4| AI-driven analytical logic chain.** A structured, stepwise logic chain guiding AI-assisted proteomic data analysis. The workflow progresses from data input and preprocessing, through systematic result interpretation and knowledge extraction, to conclusion generation and hypothesis formulation, followed by experimental validation and manuscript preparation. Each node represents a key analytical stage under a unified framework with clearly defined instructions, enabling reproducible and interpretable outcomes.

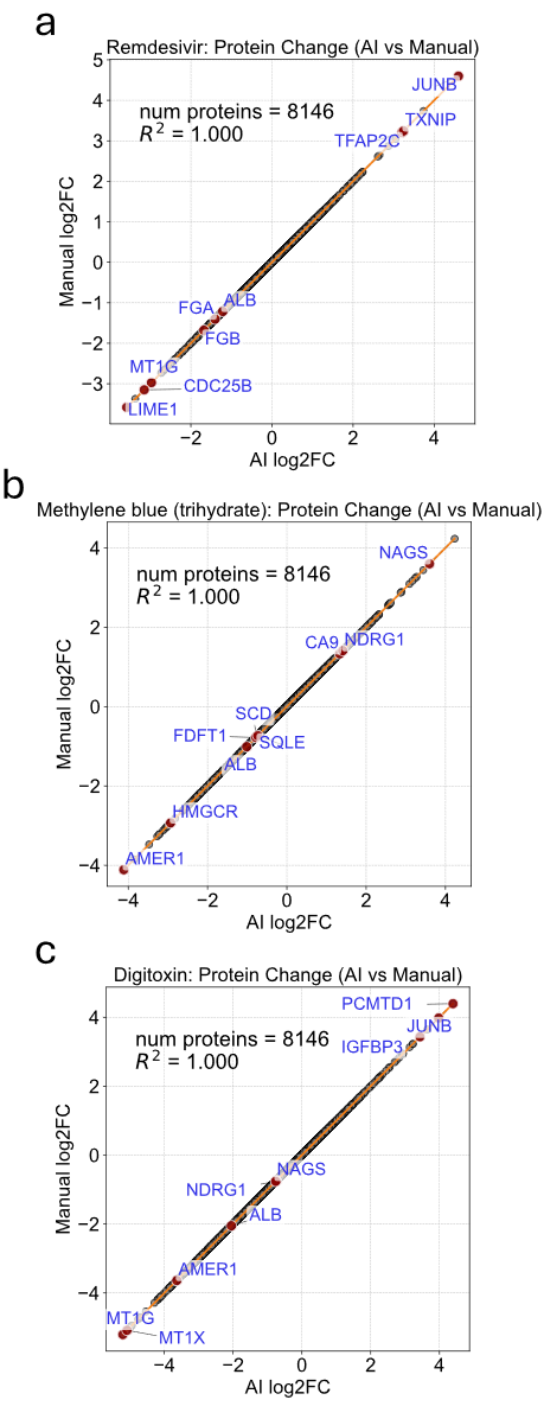


**Fig.S5 | Comparison of AI-derived and manually derived log2FC values for representative drugs. a,** Remdesivir. **b,** Methylene blue. **c,** Digitoxin

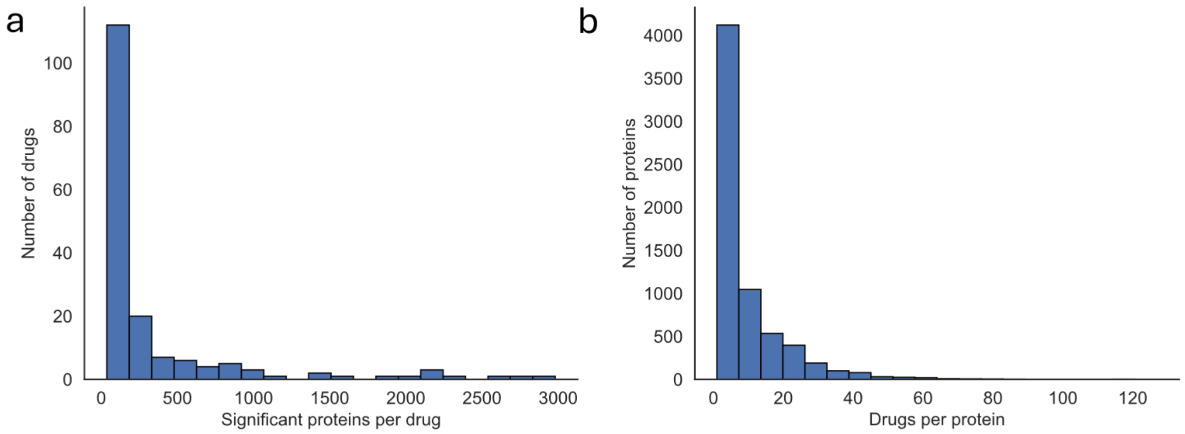


**Fig.S6 | Distribution of proteome-wide differential regulation across drugs and proteins. a,** Histogram showing the distribution of the number of significantly dysregulated proteins per drug, illustrating the variability in proteomic perturbation magnitude across compounds. **b,** Histogram showing the distribution of the number of drugs affecting each protein, highlighting a subset of proteins that are recurrently dysregulated across many treatments.

a

**Simvastatin – HMG-CoA Reductase Inhibitor**

•**327 DE proteins**

•**Same cholesterol pathway activation as atorvastatin**: HMGCR (log2FC=3.48), HMGCS1, MVK, MVD, FDFT1, SQLE, LDLR, PCSK9 — all significantly upregulated

•**Pathway enrichment**: Cholesterol Biosynthesis (Reactome FDR=4.5e-16), Steroid Biosynthesis (KEGG FDR=2.4e-9)

•**Validation**: VERY STRONG

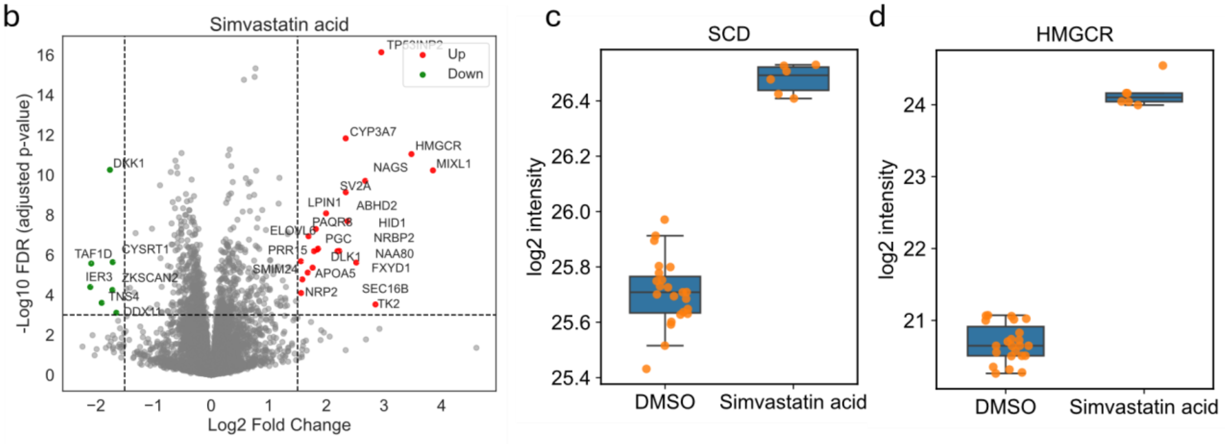


**Fig. S8 | Simvastatin induces coordinated activation of cholesterol biosynthesis pathways. a,** Representative example of AI-prioritized findings showing that simvastatin treatment is associated with activation of cholesterol biosynthesis programs. **b,** Volcano plot highlighting significantly dysregulated proteins, with key enzymes in cholesterol metabolism prominently upregulated. **c–d,** Boxplots of selected genes demonstrating consistent increases in protein abundance across replicates compared to DMSO controls.

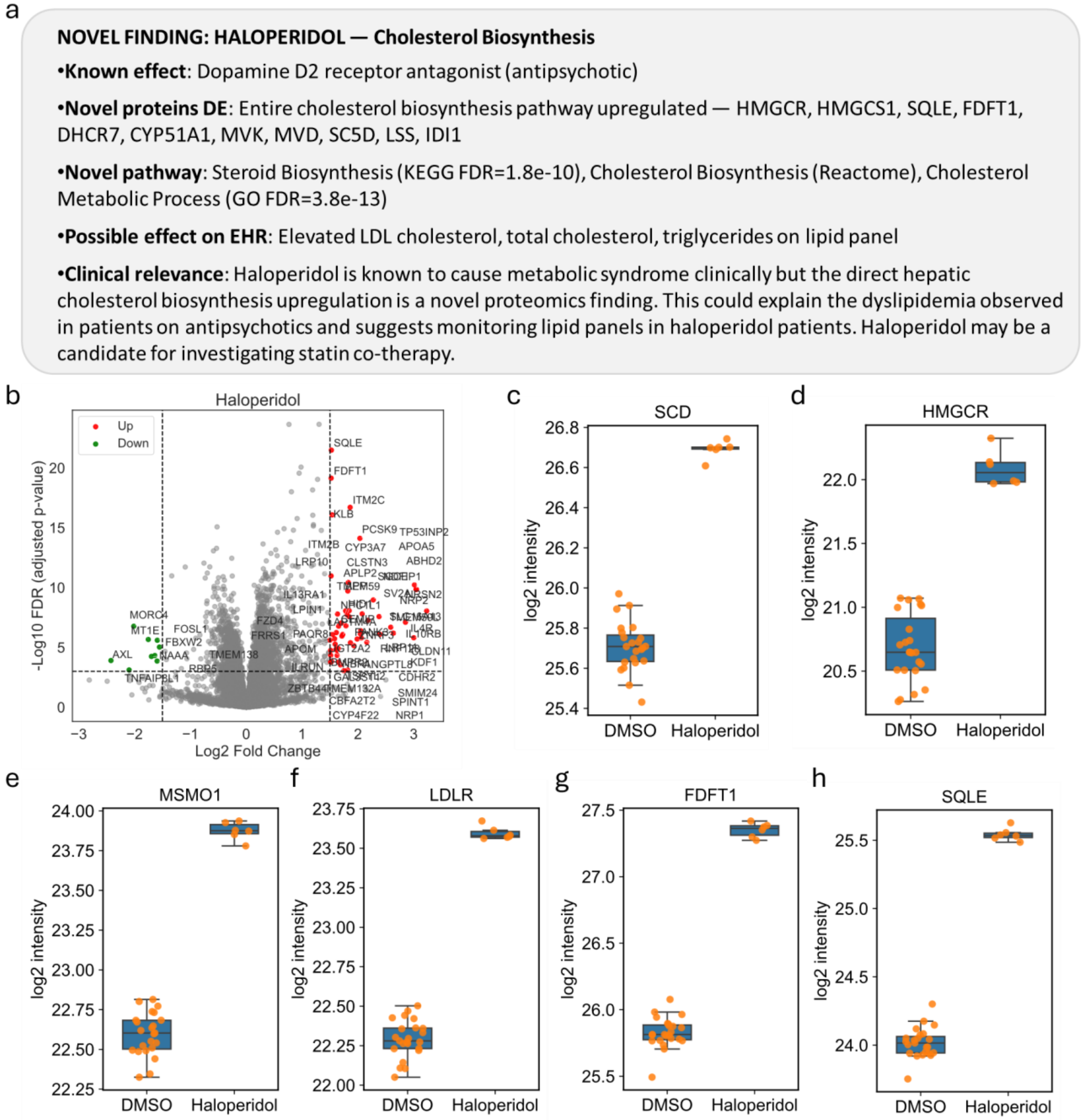


**Fig.S9 | Haloperidol induces coordinated upregulation of cholesterol biosynthesis pathways.** **a,** Representative example of AI-prioritized findings showing that haloperidol treatment is associated with activation of cholesterol biosynthesis programs. **b,** Volcano plot highlighting significantly dysregulated proteins, with key enzymes in cholesterol metabolism (e.g., SQLE, FDFT1, MSMO1, SCD, LDLR) prominently upregulated. **c-h,** Boxplots of selected genes increases in protein abundance across replicates compared to DMSO controls.

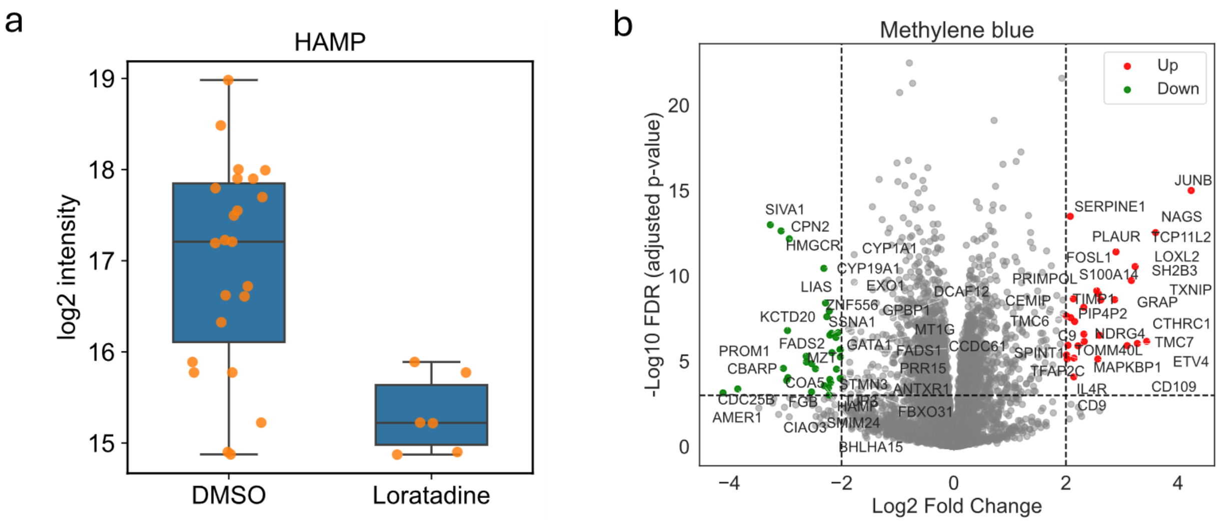


**Fig.S10 | Loratadine induces HAMP downregulation and Methylene blue triggers distinct proteomic perturbations. a,** Boxplot showing reduced HAMP (hepcidin) protein abundance in loratadine-treated samples compared to DMSO controls. **b,** Volcano plot of Methylene blue treatment highlighting significantly dysregulated proteins, with representative upregulated and downregulated targets labeled.

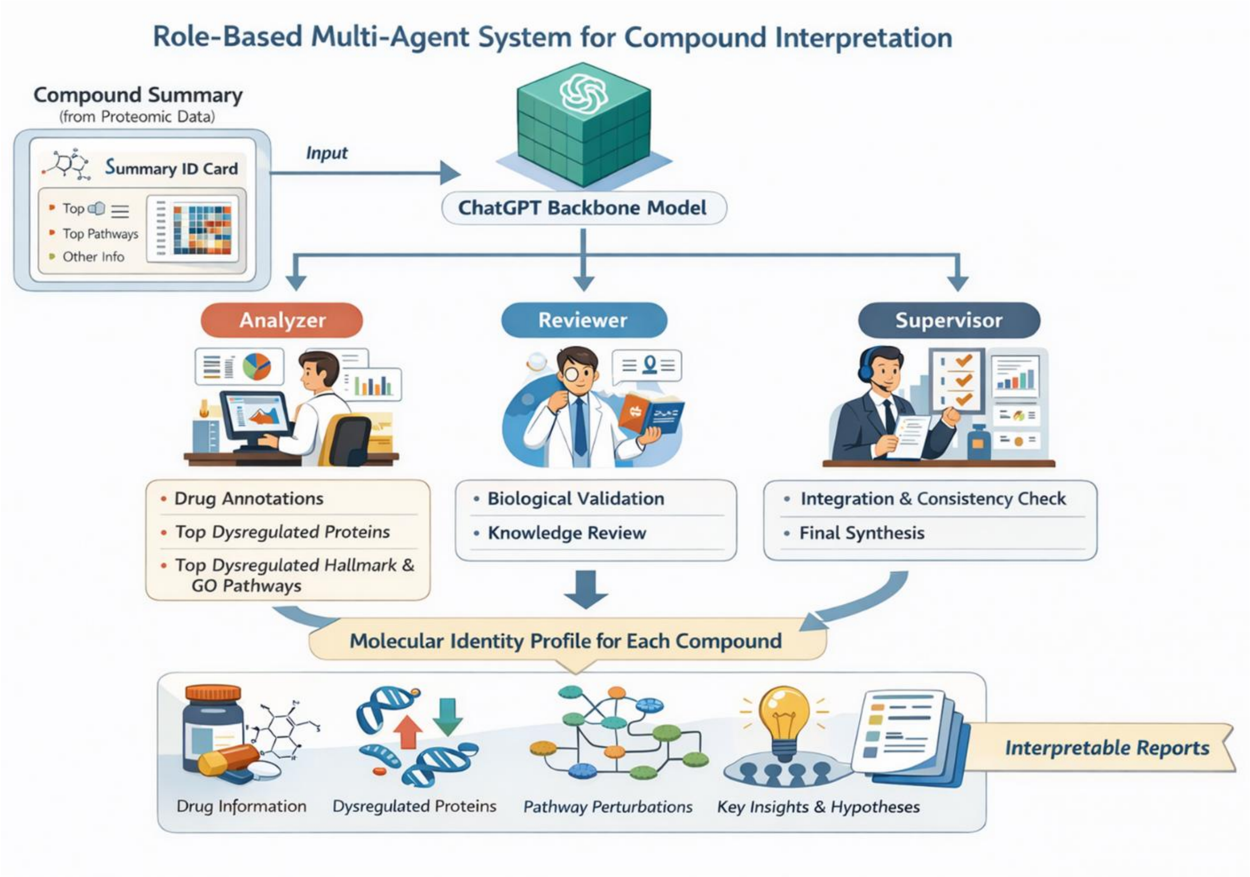


**Fig. S11 | Role-based multi-agent system for compound interpretation.** Schematic of the role-based multi-agent system (MAS) framework built on a ChatGPT backbone model. The workflow divides interpretation into three coordinated roles: an "Analyzer" for identifying dysregulated proteins and pathways, a "Reviewer" for biological validation, and a "Supervisor" for integration and final synthesis. This framework generates structured molecular identity profiles for each compound, integrating drug annotations, proteomic changes, pathway perturbations (Hallmark and GO BP), and key hypotheses into interpretable reports.

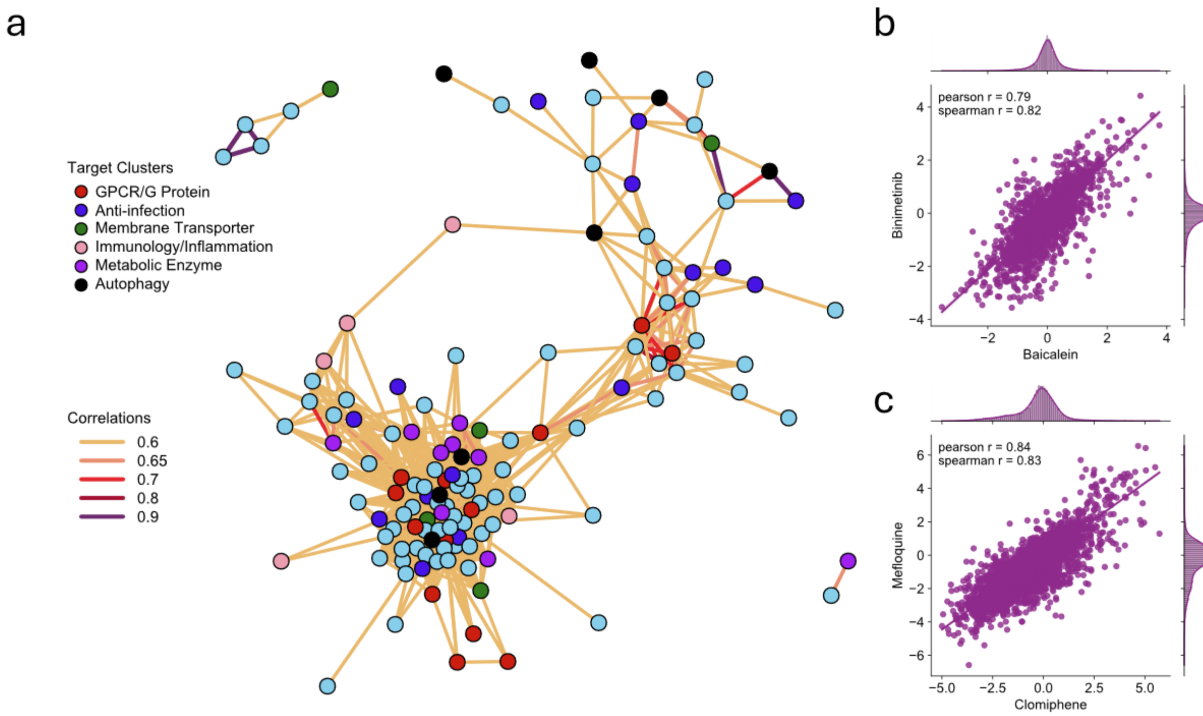


**Fig. S12 | Groups of drugs with proteome responses. a,** Community plot built from a drug–drug correlation matrix of whole proteome. Typical drug clusters were labeled with different color. Each dot represents a drug and the light blue dots represent other drugs which are not grouped into typical clusters. Spearman correlations were applied and filtered to only include edges with spearman r > 0.6. **b,c,** Pairwise correlation plot of two typical drug pairs: Baicalein and Binimetinib (b), Clomiphene and Mefloquine (b).

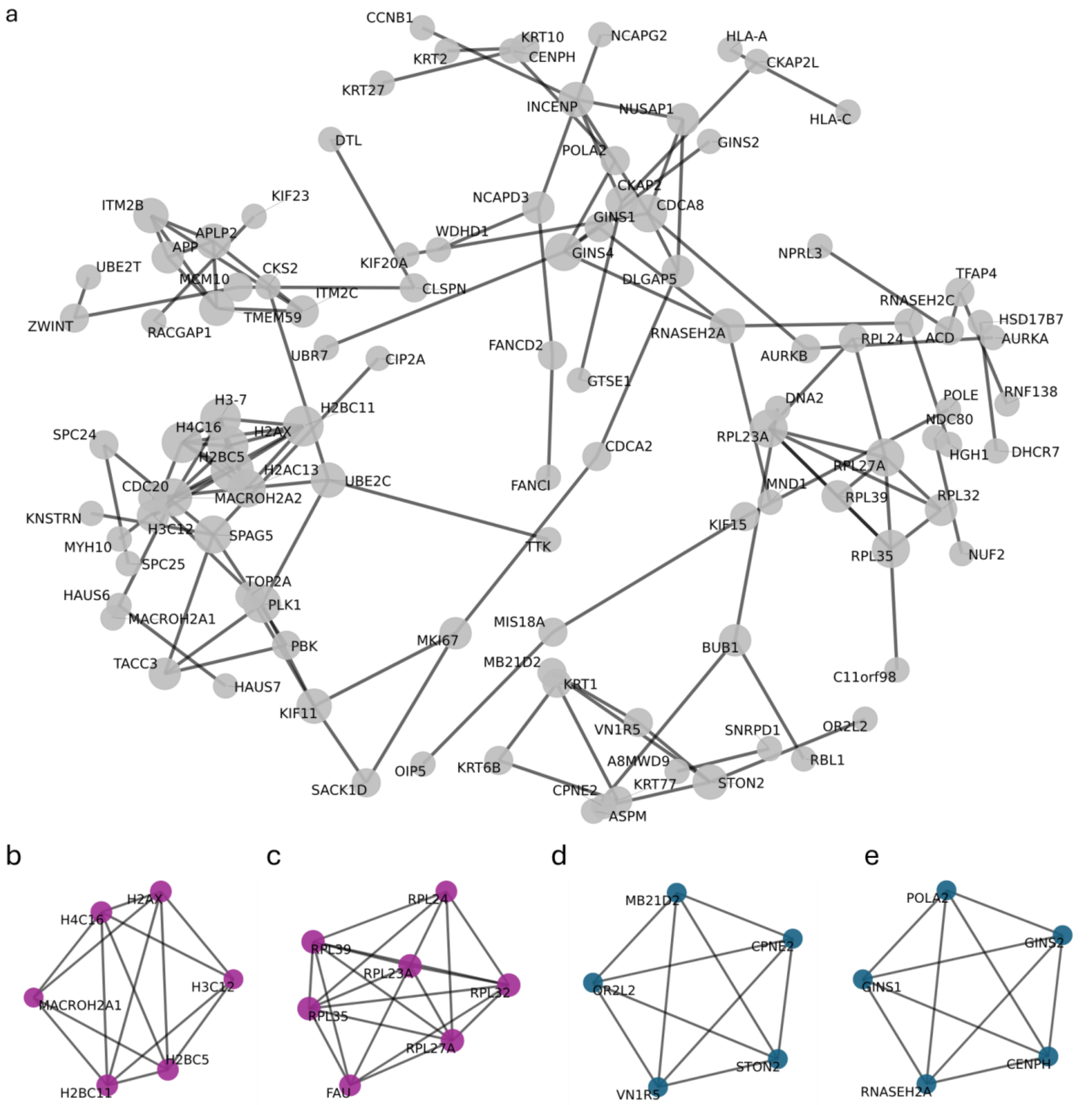


**Fig. S13 | Protein–protein correlations capture coordinated relationships within and across drug response pathways. a,** Protein–protein correlation network generated from pairwise correlations of protein response profiles across all drugs after protein-level filtering. Only the top correlated protein pairs (top 120) were retained for visualization. Edge width reflects correlation strength, whereas node size reflects the number of network connections for each protein. Labels denote gene symbols, and the force-directed layout emphasizes groups of proteins with coordinated responses across drug treatments. **b-e,** Subnetwork visualization of protein–protein correlation modules. (b,c) Representative clusters corresponding to known biological processes, including ribosome-associated proteins and chromatin-related proteins, showing strong co-regulation and validating the biological relevance of the network. (d,e) Highly correlated protein clusters lacking clear prior annotation, suggesting potential novel functional relationships or underexplored biological programs revealed by proteome-wide correlation analysis.

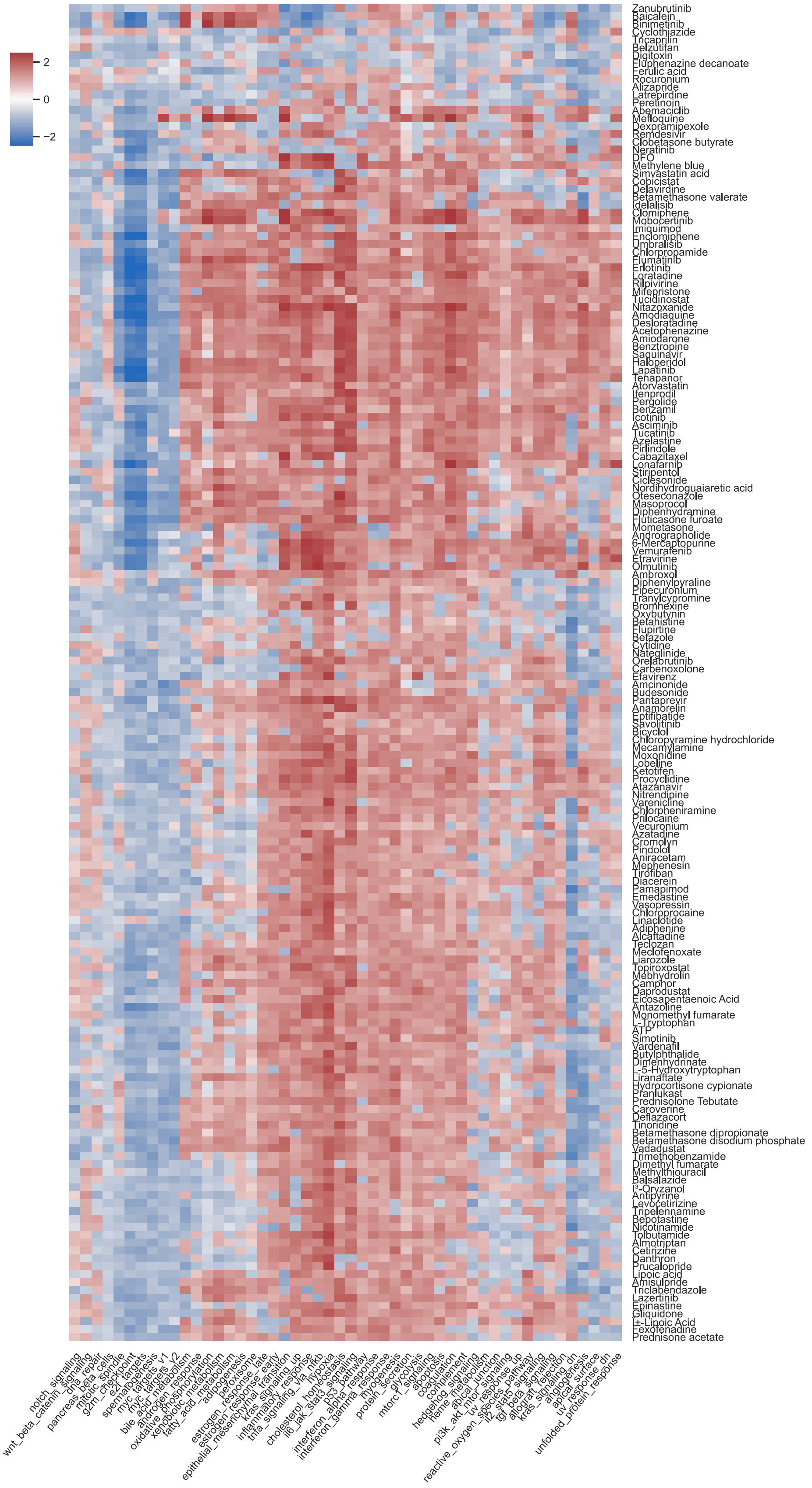

2
0
−2
Zanubrutinib
Baicalein
Binimetinib
Cyclothiazide
Tricaprilin
Belzutifan
Digitoxin
Fluphenazine decanoate
Ferulic acid
Rocuronium
Alizapride
Latrepirdine
Peretinoin
Abemaciclib
Mefloquine
Dexpramipexole
Remdesivir
Clobetasone butyrate
Neratinib
DFO
Methylene blue
Simvastatin acid
Cobicistat
Delavirdine
Betamethasone valerate
Idelalisib
Clomiphene
Mobocertinib
Imiquimod
Enclomiphene
Umbralisib
Chlorpropamide
Flumatinib
Erlotinib
Loratadine
Rilpivirine
Mifepristone
Tucidinostat
Nitazoxanide
Amodiaquine
Desloratadine
Acetophenazine
Amiodarone
Benztropine
Saquinavir
Haloperidol
Lapatinib
Tenapanor
Atorvastatin
Ifenprodil
Pergolide
Benzamil
Icotinib
Asciminib
Tucatinib
Azelastine
Pirlindole
Cabazitaxel
Lonafarnib
Stiripentol
Ciclesonide
Nordihydroguaiaretic acid
Oteseconazole
Masoprocol
Diphenhydramine
Fluticasone furoate
Mometasone
Andrographolide
6-Mercaptopurine
Vemurafenib
Etravirine
Olmutinib
Ambroxol
Diphenylpyraline
Pipecuronium
Tranylcypromine
Bromhexine
Oxybutynin
Betahistine
Flupirtine
Betazole
Cytidine
Nateglinide
Orelabrutinib
Carbenoxolone
Efavirenz
Amcinonide
Budesonide
Paritaprevir
Anamorelin
Eptifibatide
Savolitinib
Bicyclol
Chloropyramine hydrochloride
Mecamylamine
Moxonidine
Lobeline
Ketotifen
Procyclidine
Atazanavir
Nitrendipine
Varenicline
Chlorpheniramine
Prilocaine
Vecuronium
Azatadine
Cromolyn
Pindolol
Aniracetam
Mephenesin
Tirofiban
Diacerein
Pamapimod
Emedastine
Vasopressin
Chloroprocaine
Linaclotide
Adiphenine
Alcaftadine
Teclozan
Meclofenoxate
Liarozole
Topiroxostat
Mebhydrolin
Camphor
Daprodustat
Eicosapentaenoic Acid
Antazoline
Monomethyl fumarate
L-Tryptophan
ATP
Simotinib
Vardenafil
Butylphthalide
Dimenhydrinate
L-5-Hydroxytryptophan
Liranaftate
Hydrocortisone cypionate
Pranlukast
Prednisolone Tebutate
Caroverine
Deflazacort
Tinoridine
Betamethasone dipropionate
Betamethasone disodium phosphate
Vadadustat
Trimethobenzamide
Dimethyl fumarate
Methylthiouracil
Balsalazide
Î³-Oryzanol
Antipyrine
Levocetirizine
Tripelennamine
Bepotastine
Nicotinamide
Tolbutamide
Almotriptan
Cetirizine
Danthron
Prucalopride
Lipoic acid
Amisulpride
Triclabendazole
Lazertinib
Epinastine
Gliquidone
Î±-Lipoic Acid
Fexofenadine
Prednisone acetate
notch_signaling
wnt_beta_catenin_signaling
dna_repair
pancreas_beta_cells
mitotic_spindle
g2m_checkpoint
e2f_targets
spermatogenesis
myc_targets_v1
myc_targets_v2
bile_acid_metabolism
androgen_response
oxidative_phosphorylation
xenobiotic_metabolism
fatty_acid_metabolism
adipogenesis
peroxisome
estrogen_response_late
estrogen_response_early
epithelial_mesenchymal_transition
kras_signaling_up
inflammatory_response
tnfa_signaling_via_nfkb
hypoxia
cholesterol_homeostasis
il6_jak_stat3_signaling
p53_pathway
interferon_alpha_response
interferon_gamma_response
myogenesis
protein_secretion
glycolysis
mtorc1_signaling
apoptosis
coagulation
complement
hedgehog_signaling
heme_metabolism
apical_junction
pi3k_akt_mtor_signaling
uv_response_up
reactive_oxygen_species_pathway
il2_stat5_signaling
tgf_beta_signaling
allograft_rejection
kras_signaling_dn
angiogenesis
apical_surface
uv_response_dn
unfolded_protein_response

**Fig. S14 | Heatmap of pathway dysregulation across all drugs in the 50 Hallmark gene sets.** Data is derived from normalized enrichment scores (NES) of pre-ranked gene set enrichment analysis (GSEA).

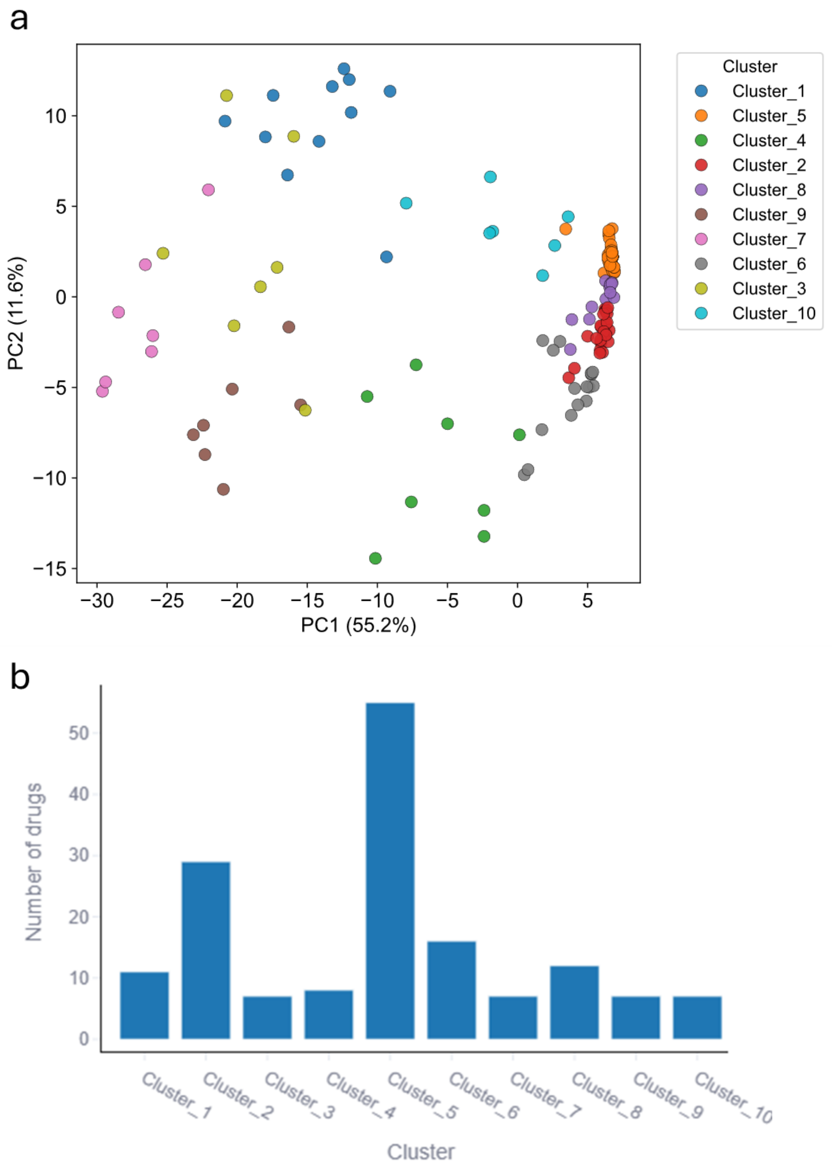


**Fig. S15 | GO pathway–based clustering of drugs reveals distinct functional groups. a,** Principal component analysis (PCA) of drugs based on pathway-level perturbation profiles derived from GO Biological Process enrichment scores, showing separation into 10 clusters identified by fuzzy c-means clustering. Each point represents a drug, colored by its assigned cluster. **b,** Bar plot showing the number of drugs in each cluster.

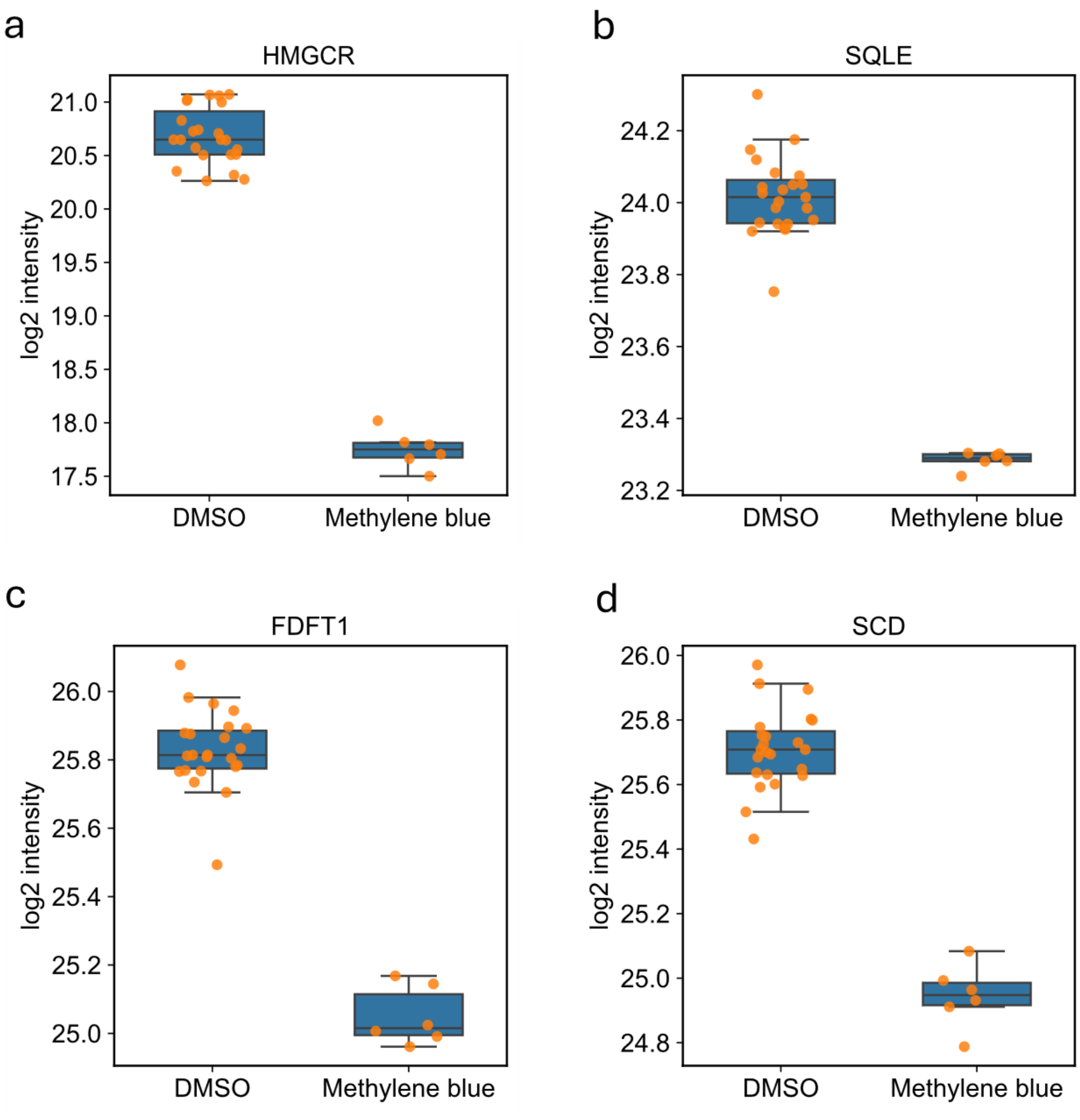


**Fig. S16 | Methylene blue downregulates cholesterol biosynthesis enzymes. a–d,** Boxplots showing reduced protein abundance of key cholesterol biosynthesis enzymes (HMGCR, SQLE, FDFT1, and SCD) in methylene blue–treated samples compared to DMSO controls, indicating suppression of cholesterol synthesis pathways.

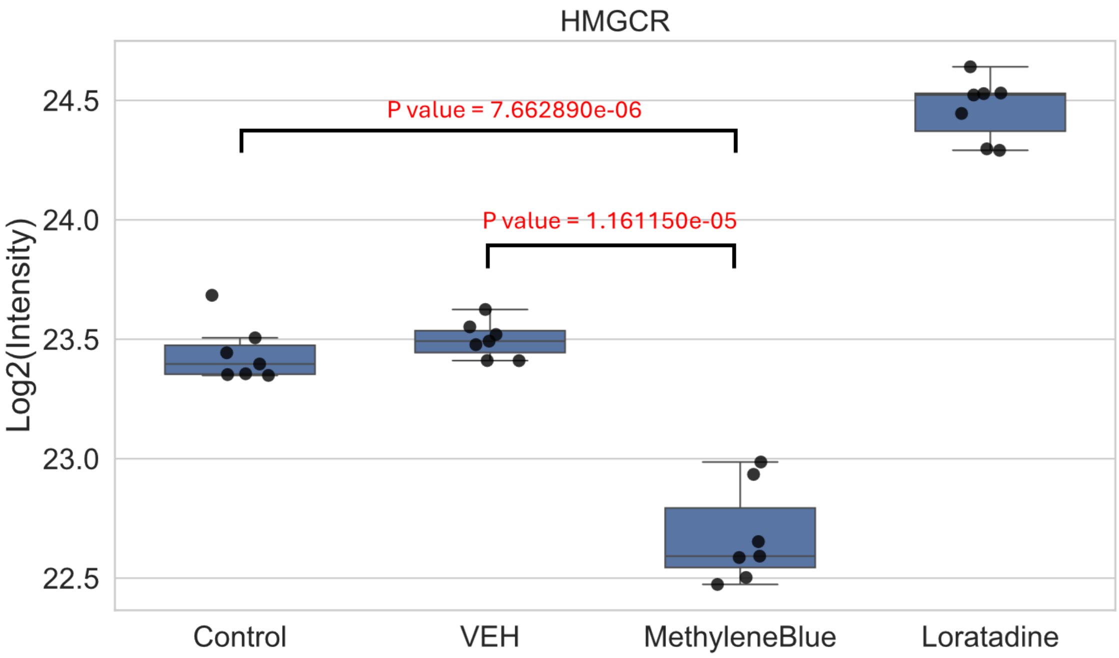


**Fig. S17 | Independent mass spectrometry validation of HMGCR regulation following methylene blue treatment.** Box-and-scatter plot showing log2-transformed HMGCR protein intensity across Control, VEH, Loratadine, and Methylene Blue treatment groups. The Control group represents untreated cells, while the VEH group corresponds to cells treated with the solvent control. Boxes indicate the interquartile range with median line. P values were calculated between the indicated groups.
