## Supplementary Tables and Files for "Robotic perturbation proteomics and AI agents enable scalable drug mechanism discovery": Supplementary File2.docx

### Zero-shot prompt

You are provided with a high-throughput drug screen proteomic dataset of HepG2 cells (file name: hepg2_drugscreen_prote.csv). Each column represents a sample. The column names represent drug names and sample type. Drug names are followed by an underscore and a number indicating the replicate numbers. Additional batch (plate) and sample type information was in (file name: map.csv). For example, Blue1 is a batch, Red1 is a separate batch, RedMix is a separate batch, etc. Column name with “QC” in it means QC samples, “DMSO” is control samples, and Deferoxamine(DFO) is a positive test to check if the whole experiment works fine. Additional details of tested drugs were included in a file (file name: drugs.csv), including mechanism of action (MoA), targets, categories, etc.

#### Analysis rules and goal

Assume fixed random seed = 42. Use corrected values (B-H FDR) for all statistical tests and visualizations. Organize into separate notebooks as needed and save the final copies of the fully executed notebooks. The overarching goals are to create reproducible, readable Jupyter notebooks-based bioinformatics analysis (QC, batch correction, differential expression (DE), gene ontology, etc) of the dataset provided; validate the workflow by ensuring known proteins and pathways are differentially expressed for well characterized drugs (e.g., DFO and iron chelation; search for novel insights into drug-protein-pathway responses based on the differential expression (DE) results, pathway enrichment analysis and literature search/drug metadata; create testable hypotheses for several drugs with unreported effects on certain proteins/pathways that might have downstream effects on EHR data variables; and lastly, plan targeted wet lab in vitro experiments to follow up on promising results.

#### Analyses to perform

Ensure your notebooks contain the following analyses at the bare minimum; perform additional analyses as you see fit. Think deeply about the outputs (i.e., do the DE results look correct? Are additional complementary analyses needed?) of your analyses. Within notebooks, use text cells followed by the outputs to explain your reasoning or insights as needed.

#### Step1. Data cleaning:

\# detected proteins, % missing; Drop proteins present in less than one third (1/3) of samples. Drop samples with detected proteins in less than 80% of cohort average. Exclude drugs with less than 3 replicates remaining.

#### Step2. Data missing value imputation, batch effect correction

Take log2 to the filtered dataset after **Step1**, Missing values imputed using shifted minimum per protein (min - 1 log2 unit). Perform ComBat batch correction and produce UMAP plots before and after batch correction (annotated by Control/QC/Drug and Batch). Color rules: QC green, positive controls red (DFO), negative controls blue (DMSO); other drugs colored by a distinct categorical palette. Use corrected data for all downstream analyses.

#### Step 3. Dysregulation analysis

Perform differential expression (DE) analysis (Welch t-test per protein for each Drug vs DMSO from the dataset generated after **Step2** above. Be sure to use corrected, filtered values. Perform multiple testing BH within drug; call DE if FDR<0.05 and |log2FC|≥0.5. Save the summary table of DE analysis of all drugs. The summary should include all proteins dysregulation values of all drugs.

#### Step 4. Drug validation study

From the drug DE summary above, select a subset (~10) of drugs that are commonly taken by human patients (we will later search real EHR data for patients taking these drugs and need sufficient sample size). From these common drugs, perform pathway enrichment analysis for separate drugs that have known, well-described pathways/proteins that would be expected to be enriched (utilizing drug.csv or web/literature searches). These are the "validation drugs”. For all gene ontology/pathway enrichment analyses, preferably use GSEApy; ensure correct dotplots or enrichplot plots are created. Use KEGG, Reactome, and GO BP as gene sets. For at least 5 of the validation drugs of highest confidence, write a short summary of validation results. For example, DFO should show iron chelation or related proteins. Ensure your summaries include the specific PROTEINS AND PATHWAYS for the DRUGS and their known MoAs. The goal is to verify that the drug screen data is reliable by demonstrating known drug responses are present, such that novel hypotheses generated in the follow steps from drugs affecting unexpected pathways/proteins can be reliably investigated further.

#### Step 5. Drug novelty analysis:

Also from these drugs, pathway enrichment analysis for drugs that have unique, novel, or unexpected proteins or pathways that were differentially abundant or enriched. These are the “novel drugs”. Again, you do not need to perform this analysis for all drugs, only a subset. Gather evidence either from the drug.csv or web/literature searches. For at least 10 of the validation drugs of highest confidence, write a short summary of the novelty of the proteins/pathways found versus proteins/pathways known or expected to be found. Propose plausible MoAs for drugs that do not have well characterized MoAs or potential drug repurposing pathways that have not been previously studied in the literature/clinical trials. Base your evidence on protein differential abundance signatures/GSEA/network, and web/literature/clinical trial searches. Suggest at least 10 novel drugs and their affected pathways/proteins. Focus on drugs that are commonly taken, but have unexpected protein/pathway responses in this dataset, and with measurable lab variables (i.e., electronic health data) that could verify/refute the hypothesis (e.g., things that you get measured at your annual physical, lipid panel, blood cell counts, comprehensive metabolic panel that includes lots of stuff, potentially thyroid and vitamin levels). Write summary results in a format such as: <DRUG> is known for <KNOWN EFFECT, PATHWAY OR PROTEINS>, but had the following <NOVEL PROTEINS FOR THIS DRUG> differentially expressed, which are related to <NOVEL PATHWAY FOR THIS DRUG>, possibly indicating an <EFFECT> on <EHR LAB VARIABLE>, making <DRUG> an indicator or treatment for <DISEASE AREA>. For context, your hypotheses will be used to search actual EHR data and plan actual wet-lab experiments to follow up on. Perform literature search to confirm these are novel findings. If it is not novel, try to find a different drug.

#### Step 6. Summary and final report.

The goal is to identify a few promising drugs and protein candidates to screen further to uncover novel mechanisms, pathways, or treatments (i.e., drug re-purposing; off-target effects), given that these are human HepG2 cells. Again, these should be drugs that are relatively commonly taken by patients with measurable variables present in EHR data so we can quickly verify or refute hypotheses. If your hypotheses require experiments outside of EHR data, include an experimental plan of how to test your hypothesis, including what exact materials (catalog numbers) to purchase if not part of a common proteomics lab inventory. End the session by creating a comprehensive PDF report of your figures and findings in the style of a high impact peer reviewed, scientific journal article, Include peer reviewed references where appropriate.

### Follow-up task1: Drug proteomic perturbations and cellular mechanism-of-action (MoA) analysis

#### Goal:

Generate quantitative, structured summaries of drug-induced proteomic perturbations that can be used as inputs for downstream AI-based interpretation (e.g., MAS system). Do NOT perform narrative interpretation, hypothesis generation, or per-drug summaries.

#### General rules:

- Use corrected, filtered, batch-corrected data only.

- Use BH-adjusted P values.

- Define significance as FDR < 0.05 and |log2FC| >= 0.5.

- Fixed random seed = 42.

- Do NOT generate long-form text interpretation.

- Focus on reproducible tables and figures.

#### Part A. Drug-level perturbation magnitude

1. For each drug:

- Count number of significantly upregulated proteins

- Count number of significantly downregulated proteins

- Count total number of significant proteins

2. Rank drugs by total perturbation magnitude.

3. Generate:

- Table: drug_perturbation_summary.csv

Columns:

drug, n_upregulated, n_downregulated, n_total_significant

- Figure:

Bar plot showing up/down counts per drug

#### Part B. Global protein response frequency

1. For each protein:

- Count number of drugs significantly upregulating it

- Count number of drugs significantly downregulating it

2. Generate:

- Table: protein_regulation_frequency.csv

Columns:

protein, gene_symbol, n_drugs_up, n_drugs_down

3. Identify:

- Top recurrently upregulated proteins

- Top recurrently downregulated proteins

4. Generate:

- Waterfall / ranked plot of protein regulation frequency

#### Part C. Per-drug structured feature extraction (NO interpretation)

For each drug, extract structured features ONLY:

1. Top dysregulated proteins:

- Top N upregulated proteins (sorted by FDR or log2FC)

- Top N downregulated proteins

2. Pathway-level features:

- Hallmark:

top positively enriched pathways

top negatively enriched pathways

- GO Biological Process:

top positively enriched terms

top negatively enriched terms

3. Save per-drug structured data (no narrative):

Folder: drug_feature_tables/

For each drug:

- <drug>_top_proteins.csv

- <drug>_hallmark_gsea.csv

- <drug>_go_bp_gsea.csv

#### Part D. Global summary matrices (for MAS input)

1. Construct:

- drug × protein log2FC matrix

- drug × protein significance matrix (binary: 0/1)

- drug × pathway NES matrices (Hallmark and GO BP)

2. Save:

- drug_log2fc_matrix.csv

- drug_significance_matrix.csv

- drug_hallmark_nes_matrix.csv

- drug_go_bp_nes_matrix.csv

#### Part E. Deliverables

Produce:

1. Drug perturbation summary table

2. Protein frequency table

3. Per-drug feature tables (NO interpretation)

4. Global matrices (for downstream MAS input)

5. Minimal figures (barplot + waterfall)

### Follow-up task2: Identification of groups of drugs with similar cellular mechanisms

#### Goal:

Identify and quantify similarity relationships between drugs based on proteomic responses. Generate clustering, network, and pathway-level structure outputs WITHOUT biological interpretation or narrative summaries.

#### General rules:

- Use corrected log2FC and GSEA NES values only.

- Fixed random seed = 42.

- Do NOT generate long-form interpretation or hypothesis text.

- Focus on quantitative similarity, clustering, and structured outputs.

#### Part A. Drug–drug similarity (protein level)

1. Use drug × protein log2FC matrix.

2. Compute:

- Pearson correlation matrix

- Spearman correlation matrix

3. Identify strong similarities:

- Pearson r > 0.5

- Spearman r > 0.6

4. Save:

- drug_pearson_corr.csv

- drug_spearman_corr.csv

- high_similarity_pairs.csv

Columns:

drug1, drug2, pearson_r, spearman_r

#### Part B. Drug similarity network

1. Build network:

- Nodes: drugs

- Edges: Pearson r > 0.5

2. Perform community detection:

- Louvain / Leiden / modularity-based method

3. Save:

- network_edges.csv

- network_nodes.csv

- drug_community_assignments.csv

Columns:

drug, community_id

4. Generate:

- network visualization

#### Part C. Protein co-regulation modules

1. Compute protein–protein correlation across drugs:

- based on log2FC profiles

2. Identify co-regulated modules:

- clustering or network-based grouping

3. Save:

- protein_correlation_matrix.csv

- protein_modules.csv

Columns:

protein, module_id

4. Generate:

- representative protein network plots

#### Part D. Pathway-level clustering (GO BP)

1. Use drug × GO BP NES matrix.

2. Filter:

- pathways present in ≥3 drugs (or similar threshold)

- optionally select top variable pathways

3. Perform fuzzy c-means clustering:

- n_clusters = 10

- fuzziness ~1.7

4. Save:

- drug_fcm_membership.csv

- drug_fcm_cluster_assignment.csv

Columns:

drug, cluster_id

5. Generate:

- GO BP NES heatmap grouped by cluster

#### Part E. Hallmark clustering

1. Use drug × Hallmark NES matrix.

2. Perform hierarchical clustering.

3. Save:

- hallmark_cluster_matrix.csv

4. Generate:

- hallmark clustering heatmap

#### Part F. Cluster feature extraction

For each cluster:

1. Extract:

- drugs in cluster

- top positively enriched GO BP pathways

- top negatively enriched GO BP pathways

- top hallmark pathways

- top representative proteins

2. Save:

- cluster_feature_summary.csv

Columns:

cluster_id,

drug_list,

top_up_pathways,

top_down_pathways,

top_hallmark_up,

top_hallmark_down,

representative_proteins

#### Part G. Deliverables

Produce:

1. Drug–drug correlation matrices

2. Similarity network + communities

3. Protein co-regulation modules

4. GO BP clustering (fuzzy c-means)

5. Hallmark clustering

6. Cluster-level structured features
