## Supplementary Tables and Files for "Robotic perturbation proteomics and AI agents enable scalable drug mechanism discovery": Supplementary File5.md

# All Drug AI Summaries


---

# 6-Mercaptopurine

# Drug summary: 6-Mercaptopurine

## Executive summary
6-Mercaptopurine elicits a clear antiproliferative phenotype consistent with a nucleoside antimetabolite: broad repression of E2F targets, G2/M checkpoint, DNA replication, and mitotic processes is accompanied by decreased CDC7, EXO1, E2F6/8, and STIL. Concomitantly, a strong inflammatory/TNFA–NFkB–like stress-secretory program and cellular stress responses (hypoxia, unfolded protein response, apoptosis) are induced, with marked increases in secreted/stress-associated proteins (GDF15, IGFBP1, PLAUR, TNFRSF12A, IL4R) and TXNIP. EMT-like tissue-remodeling and angiogenesis gene-set enrichments with LOXL2, PLAUR, CEMIP, NRP2, and TYRO3 up suggest a wound-healing–type adaptation rather than a primary mechanism. There is no proteomic evidence for autophagy activation in this dataset. Pathway-level signals also suggest potential adaptive IGF/PI3K–AKT and amino acid import programs and a tentative downshift in lipid/acetyl-CoA supply (HMGCR, SLC13A5 down), but these should be interpreted cautiously.

## Findings consistent with expected biology
- Strong suppression of proliferation programs (Hallmark E2F_TARGETS and G2M_CHECKPOINT down; GO DNA replication/mitotic processes down), consistent with 6-mercaptopurine’s antimetabolite activity.
- Concordant decreases in replication and cell-cycle regulators (CDC7, EXO1, E2F6, E2F8, STIL) supporting cell-cycle arrest.
- Robust inflammatory/TNFA–NFkB–like and stress-secretory signatures with increased GDF15, IGFBP1, PLAUR, TNFRSF12A, and IL4R.
- Induction of hypoxia, unfolded protein response, and apoptosis programs; TXNIP up-regulation consistent with metabolic/oxidative stress.
- No evidence of autophagy activation despite drug annotation.

## Unexpected but plausible findings
- EMT-like tissue-remodeling and angiogenesis signatures despite proliferation shutdown, with LOXL2, PLAUR, CEMIP, NRP2, and TYRO3 up; likely a stress/wound-healing phenotype.
- Pathway enrichment for IGF receptor and positive regulation of PI3K–AKT signaling during growth suppression; IGFBP1 is strongly up but may sequester IGFs, so receptor activation remains unproven.
- Down-regulation of HMGCR and the citrate transporter SLC13A5, suggesting a possible reduction in cholesterol/acetyl-CoA supply; not supported by lipid-metabolism pathway enrichment.

## Follow-up hypotheses
### Hypothesis 1
- **Hypothesis:** An NFkB-driven stress-secretory program contributes to cell survival and EMT-like remodeling under 6-mercaptopurine treatment, and its inhibition will enhance apoptosis.
- **Why interesting:** INFLAMMATORY_RESPONSE and TNFA_SIGNALING_VIA_NFKB are strongly enriched with increases in stress/secreted proteins (e.g., GDF15, PLAUR, TNFRSF12A), suggesting a pro-survival adaptive axis.
- **Suggested validation:** Measure NFkB activity (p65 nuclear translocation, NFkB reporter), secreted cytokines, and assess synergy of IKK/NFkB inhibitors with 6-mercaptopurine on viability and apoptosis (caspase activation, Annexin V).
- **Priority:** high

### Hypothesis 2
- **Hypothesis:** 6-mercaptopurine enforces G1/S arrest via repression of E2F programs and depletion of replication initiation signaling (e.g., CDC7), providing mechanistic biomarkers of response.
- **Why interesting:** Broad down-regulation of E2F/G2M pathways with decreased CDC7, EXO1, and E2F6/8 aligns with the drug’s antimetabolite class and suggests tractable pharmacodynamic readouts.
- **Suggested validation:** Cell-cycle profiling (PI/EdU), DNA synthesis assays (EdU/BrdU incorporation), and immunoblotting for CDC7 and MCM2 Ser40/41 phosphorylation alongside E2F target expression.
- **Priority:** high

### Hypothesis 3
- **Hypothesis:** The EMT-like/tissue-remodeling program is a stress-induced adaptation (e.g., TGF-β/LOXL2-driven) that can be mitigated to limit potential pro-invasive phenotypes.
- **Why interesting:** EMT and tissue-remodeling gene sets are up with LOXL2, PLAUR, and CEMIP increases, consistent with a wound-healing response that could affect long-term persistence.
- **Suggested validation:** Assess EMT marker panels (E-cadherin, Vimentin, ZEB1/SLUG), migration/invasion assays, and test TGF-β pathway or LOXL2 inhibitors in combination with 6-mercaptopurine.
- **Priority:** medium

### Hypothesis 4
- **Hypothesis:** PI3K–AKT/IGF signaling is an adaptive survival pathway engaged under 6-mercaptopurine treatment and may be co-targetable.
- **Why interesting:** GO enrichment suggests IGF receptor and PI3K–AKT pathway engagement during growth arrest, a common adaptive response in cytotoxic stress.
- **Suggested validation:** Quantify IGF1R and AKT phosphorylation, test IGF1 responsiveness, and evaluate synergy with IGF1R or PI3K inhibitors; assess the impact of IGFBP1 on ligand availability in conditioned media.
- **Priority:** medium

### Hypothesis 5
- **Hypothesis:** Reduced mevalonate/citrate input contributes to the arrested state, and sterol or citrate supplementation may partially rescue growth under 6-mercaptopurine.
- **Why interesting:** HMGCR and SLC13A5 are decreased, hinting at lowered lipid/acetyl-CoA supply during treatment.
- **Suggested validation:** Rescue assays with mevalonate/cholesterol or citrate, measurement of SREBP activation, and lipid biosynthesis flux analyses to determine functional contribution.
- **Priority:** low

## Confidence
high

## Caution notes
- Pathway activations (NFkB, IGF/PI3K–AKT, nutrient import) are inferred from proteome-based gene set enrichment; without phosphoproteomics or functional assays, treat these as signatures rather than proven activation.
- IGFBP1 up-regulation indicates IGF-axis modulation but does not by itself imply increased IGF receptor signaling; ligand sequestration is a plausible alternative.
- EMT/angiogenesis enrichments may reflect a generalized stress/wound-healing secretory state rather than bona fide EMT or angiogenic remodeling; confirm with canonical markers and phenotypic assays.
- The lipid/acetyl-CoA metabolism observation (HMGCR, SLC13A5 down) is based on few proteins and lacks pathway-level enrichment; interpret as tentative.
- No autophagy pathway enrichment or canonical autophagy protein changes were detected; autophagy should not be inferred despite drug annotations.
- Increased amino acid import is suggested by GO enrichment without strong transporter-level support among top hits; consider this provisional.


---

# Abemaciclib (methanesulfonate)

# Drug summary: Abemaciclib (methanesulfonate)

## Executive summary
Proteomics under Abemaciclib (CDK inhibitor) shows a robust cell-cycle shutdown with coordinated loss of E2F/G2–M programs and mitotic machinery, consistent with on‑target CDK pathway inhibition. Concurrently, a stress/p53–p21 axis is activated with apoptosis-related enrichment. Cells display contextual adaptations characteristic of a non‑proliferative state, including increased amino‑acid transport and enrichment of oxidative phosphorylation and lipid/cholesterol homeostasis gene sets. Enrichment of EMT/ECM remodeling, myogenesis, and angiogenesis hallmarks suggests stress-induced remodeling or differentiation-like programs rather than definitive lineage transition. A paradoxical mTORC1 hallmark enrichment co-occurs with SESN2/TRIB3 upregulation, arguing for nuanced regulation. Notably, metallothioneins are downregulated despite oxidative-stress-associated signatures. Reactome and KEGG sections are empty, but Hallmark and GO evidence are strong, supporting the core conclusions.

## Findings consistent with expected biology
- Broad suppression of cell-cycle progression: Hallmark G2M Checkpoint, Mitotic Spindle, and E2F Targets are down; GO terms for nuclear division/chromosome segregation decrease; key regulators (CDC25B, CDC7, CDT1, SKA1, SASS6, RAD54B, GEN1, CDCA7) are reduced.
- Stress/p53–p21 pathway engagement with apoptotic signaling: TP53, CDKN1A (p21), SESN2, TXNIP, TRIB3, IGFBP2/3 increase; Hallmark Apoptosis and GO regulation of cytochrome c release/hydrogen peroxide metabolism are enriched.

## Unexpected but plausible findings
- Metabolic and transport reprogramming: enrichment of oxidative phosphorylation, fatty-acid metabolism, and cholesterol homeostasis with increased amino-acid import/transport (LPIN1, PCSK9, GPX3 up; multiple amino-acid transport GO terms up).
- EMT/ECM remodeling, myogenesis, and angiogenesis hallmarks are enriched with increases in JUNB, FOSL1, BHLHE40, DLK1—suggesting stress-associated remodeling or quiescence/differentiation-like shifts rather than definitive EMT.
- Metallothioneins (MT1X, MT1E, MT1G) are strongly downregulated despite oxidative-stress-associated signatures.
- mTORC1 signaling hallmark is enriched despite SESN2/TRIB3 induction, indicating context-dependent or gene-set-driven behavior rather than clear pathway activation.

## Follow-up hypotheses
### Hypothesis 1
- **Hypothesis:** Abemaciclib induces a p53–p21–linked growth arrest with increased susceptibility to apoptosis.
- **Why interesting:** TP53 and CDKN1A upregulation with apoptosis-related enrichment suggests a cytostatic-to-cytotoxic transition potential in this context.
- **Suggested validation:** Confirm p53 functional status; quantify p21 and cleaved caspase-3/PARP; assess mitochondrial apoptosis markers (cytochrome c release); use BH3 profiling to measure apoptotic priming.
- **Priority:** high

### Hypothesis 2
- **Hypothesis:** Cells adapt to CDK blockade by shifting toward an oxidative and lipid/cholesterol-handling metabolic state and increasing amino-acid transport.
- **Why interesting:** OXPHOS, fatty-acid metabolism, and cholesterol homeostasis hallmarks are enriched with LPIN1/PCSK9/GPX3 up and strong amino-acid transport GO signatures, consistent with a non-proliferative metabolic program.
- **Suggested validation:** Measure mitochondrial respiration and ATP-linked OCR; monitor lipid/cholesterol flux (LDL uptake, lipidomics); quantify representative amino-acid transporters indicated by GO and uptake assays (e.g., radiolabeled amino acids).
- **Priority:** medium

### Hypothesis 3
- **Hypothesis:** EMT/ECM and myogenesis hallmark enrichments reflect stress-induced secretory/ECM remodeling rather than bona fide EMT.
- **Why interesting:** Hallmark EMT/angiogenesis/myogenesis up with JUNB/FOSL1/BHLHE40 increases, but canonical EMT markers are not presented.
- **Suggested validation:** Assess E-cadherin/Vimentin/FN1 protein levels, morphology, and secretome (ECM proteomics); perform qPCR/proteomics of canonical EMT markers to distinguish remodeling from lineage transition.
- **Priority:** medium

### Hypothesis 4
- **Hypothesis:** mTORC1 activity is restrained despite an mTORC1 hallmark signal, mediated by stress sensors (e.g., SESN2/TRIB3).
- **Why interesting:** Paradoxical co-occurrence suggests gene-set enrichment may not reflect functional mTORC1 activation.
- **Suggested validation:** Probe phospho-S6K, phospho-4EBP1, and phospho-S6 across time/dose; test response to amino-acid refeeding and ISR modulation (e.g., ATF4 pathway).
- **Priority:** medium

### Hypothesis 5
- **Hypothesis:** Downregulation of metallothioneins reflects altered metal-ion homeostasis rather than reduced oxidative stress.
- **Why interesting:** MT1X/MT1E/MT1G decrease despite oxidative-stress-associated pathways, suggesting metal-regulatory axis involvement.
- **Suggested validation:** Measure cellular zinc/copper levels; assess MT1/2 mRNA and MTF1 activity/nuclear localization; test whether metal supplementation rescues MT levels.
- **Priority:** low

### Hypothesis 6
- **Hypothesis:** PCSK9 upregulation modulates cholesterol uptake by reducing LDLR, contributing to the cholesterol homeostasis program.
- **Why interesting:** PCSK9 increase aligns with cholesterol homeostasis enrichment and may impact lipid handling in arrested cells.
- **Suggested validation:** Quantify LDLR protein and surface levels; measure DiI-LDL uptake; evaluate sensitivity to PCSK9 modulation (e.g., siRNA) on lipid uptake metrics.
- **Priority:** medium

## Confidence
medium

## Caution notes
- p53 pathway inference should rely on target induction (e.g., CDKN1A) rather than TP53 abundance alone; the cell-line p53 status is not provided.
- Interpret decreases in replication/repair-associated proteins as part of cell-cycle arrest rather than a global DNA damage response suppression; no DDR pathway-level downregulation is shown.
- mTORC1 hallmark enrichment should be treated cautiously; SESN2/TRIB3 upregulation is compatible with stress-mediated mTORC1 restraint.
- EMT/angiogenesis/myogenesis enrichments may reflect stress-induced ECM/secretory programs; canonical EMT marker changes were not reported.
- Functional metabolic shifts (e.g., to oxidative phosphorylation) are inferred from enrichment and a few proteins; orthogonal assays are needed for confirmation.
- Metallothionein downregulation may be sensitive to culture metal-ion conditions and may not generalize.
- Only 25 up and 25 down proteins reached significance; while Hallmark/GO enrichments are strong, secondary programs should be interpreted conservatively.
- Reactome and KEGG pathway sections are empty; conclusions are based on Hallmark and GO evidence.


---

# Acetophenazine (dimaleate)

# Drug summary: Acetophenazine (dimaleate)

## Executive summary
Proteomics profiling of acetophenazine (dimaleate), a dopamine receptor–targeting GPCR ligand, reveals a coherent program of membrane lipid remodeling, inflammatory signaling, and growth arrest. The most prominent change is robust activation of cholesterol/sterol biosynthesis and homeostasis (Hallmark CHOLESTEROL_HOMEOSTASIS NES 2.35, FDR 0; multiple sterol/steroid GO terms FDR 0) with increases in sterol/lipid-related proteins (e.g., SC5D, APOA5, PANK3, SPPL3, GAL3ST1). In parallel, cytokine/inflammatory pathways are induced (IL6–JAK/STAT3, interferon responses, complement/coagulation; elevated IL4R, RNF128, TNFRSF12A, MICA). Cell-cycle programs are suppressed (E2F targets, G2/M checkpoint down) with broad reductions in rRNA processing, ribosome biogenesis, and translation machinery, supporting growth arrest and a biosynthetic downshift. GPCR signaling is enriched upward and there is a neuronal/glial-leaning signature (GO glial cell differentiation; increased SV2A, NRSN2, TMEM59L, SGCE, GAL3ST1, WBP1L). Epithelial/intermediate filament programs are decreased (keratinocyte/IF GO terms down; KRT17, KRT79, TNS4 reduced). MTORC1 hallmark enrichment is observed but coincides with decreased translation-related processes. Together, the data support lipid anabolic and inflammatory stress responses with reduced proliferation and emergent neuronal/glial features. Mechanistic attributions (e.g., SREBP, STAT3, mTORC1) remain hypotheses pending phospho- or transcriptional validation.

## Findings consistent with expected biology
- Upregulation of GPCR signaling downstream of a dopamine receptor perturbation (GO G protein–coupled receptor signaling enriched; NES ~1.99, FDR ~0.0028).
- Emergence of neuronal/glial features consistent with the neuronal signaling context (GO glial cell differentiation up; NES ~2.02; increased SV2A, NRSN2, TMEM59L, SGCE, WBP1L).

## Unexpected but plausible findings
- Marked activation of cholesterol/sterol biosynthesis and homeostasis with broader lipid/lipoprotein metabolism (Hallmark CHOLESTEROL_HOMEOSTASIS and multiple sterol/steroid GO terms at FDR 0; up SC5D, APOA5, PANK3, SPPL3, GAL3ST1).
- Pronounced induction of cytokine/inflammatory programs including IL6–JAK/STAT3, interferon responses, complement/coagulation (hallmarks up; increased IL4R, RNF128, TNFRSF12A, MICA).
- Coordinated suppression of cell-cycle progression and core biosynthetic machinery (Hallmarks E2F_TARGETS and G2M_CHECKPOINT down; rRNA processing/biogenesis and translation GO terms down; decreased RSL24D1, TNRC6C, POLK, POLL).
- Loss of epithelial/intermediate filament programs (intermediate filament and keratinocyte/epidermal differentiation GO terms strongly negative; KRT17, KRT79, TNS4 down).
- MTORC1 hallmark enrichment occurs despite downregulation of translation and ribosome biogenesis, suggesting selective pathway engagement.

## Follow-up hypotheses
### Hypothesis 1
- **Hypothesis:** Sterol biosynthesis activation is driven by SREBP pathway engagement in response to membrane lipid/cholesterol demand.
- **Why interesting:** Would mechanistically link GPCR perturbation to membrane remodeling with potential impact on receptor trafficking and signaling.
- **Suggested validation:** Assess SREBF1/2 processing (nuclear SREBP), and abundance of canonical SREBP targets (e.g., HMGCR/HMGCS1, LDLR); perform lipidomics for cellular cholesterol/sterol intermediates; test attenuation with SREBP inhibitors (e.g., fatostatin) or statins and examine rescue of proteomic signatures.
- **Priority:** high

### Hypothesis 2
- **Hypothesis:** The inflammatory signature reflects activation of JAK/STAT pathways (particularly STAT3) downstream of cytokine signaling.
- **Why interesting:** Provides a tractable axis linking IL6/STAT3 hallmarks and immunomodulatory surface proteins to functional signaling.
- **Suggested validation:** Measure phospho-STAT3 (Y705) and STAT1 (Y701), quantify secreted cytokines (e.g., IL-6), and assess sensitivity to JAK inhibitors; verify surface increases of IL4R/TNFRSF12A/MICA by flow cytometry.
- **Priority:** high

### Hypothesis 3
- **Hypothesis:** Translation and ribosome biogenesis are suppressed despite MTORC1 hallmark up, indicating selective or context-specific decoupling of mTOR outputs.
- **Why interesting:** Explains the concurrent lipid anabolic signature with reduced protein synthesis and could reveal stress-adaptive signaling.
- **Suggested validation:** Profile mTORC1 activity (phospho-S6K, S6, 4EBP1), quantify global translation (puromycin/OPP incorporation), examine nucleolar markers for ribosome biogenesis, and test rapamycin/torin sensitivity of lipid and translation readouts.
- **Priority:** medium

### Hypothesis 4
- **Hypothesis:** Acetophenazine treatment promotes a glial/oligodendrocyte-leaning state in this system.
- **Why interesting:** Consistent with increases in GAL3ST1 and WBP1L and glial differentiation enrichment, with implications for neuronal/glial function and drug response.
- **Suggested validation:** Assess lineage markers (e.g., GFAP for astrocytes, OLIG2/MBP/PLP1 for oligodendrocytes) by proteomics/qPCR/immunostaining; evaluate morphology and, if feasible, single-cell transcriptomics to confirm cell-state shifts.
- **Priority:** medium

## Confidence
high

## Caution notes
- Cell type and baseline context are unspecified; neuronal/glial or epithelial program shifts may reflect inherent lineage or culture conditions rather than lineage conversion.
- Proteomics-only data limit pathway activation inferences (e.g., STAT3, mTORC1, SREBP); phosphoproteomics and transcriptional assays are needed for mechanistic confirmation.
- Mechanistic attributions (SREBP-driven lipid synthesis, STAT3 dependence, mTORC1 involvement, differentiation/EMT) are hypotheses based primarily on enrichment patterns and should be presented cautiously.
- Reactome and KEGG summaries are unavailable here; interpretations rely on Hallmark and GO concordance.
- Direct dopamine receptor engagement or specific GPCR adaptations are not demonstrated at the protein level; target engagement assays would strengthen conclusions.


---

# Adiphenine (hydrochloride)

# Drug summary: Adiphenine (hydrochloride)

## Executive summary
Proteomic profiling of Adiphenine (hydrochloride) exposure reveals a strong hypoxia-like and pro-inflammatory response, with significant enrichment of Hallmark HYPOXIA (NES ~2.07, FDR 0), TNFA_SIGNALING_VIA_NFKB (NES ~1.96, FDR 0), and IL6_JAK_STAT3_SIGNALING (NES ~1.75, FDR ~0.013). An epithelial–mesenchymal transition (EMT) program is also enriched (NES ~1.77, FDR ~0.014), accompanied by GO terms for homophilic cell–cell adhesion and migration (NES >2). Protein-level changes support these programs, including increases in IRAK4, IGFBP1, CEMIP, WASF1, and PRR5. A shift in the IGF axis is suggested by opposing regulation of IGFBP1 (up) and IGFBP2 (down). Several individual proteins involved in metabolism and protein translocation are decreased (e.g., NDUFA1, PHYH, SLC13A5, SEC61G, SPCS1), but no down-regulated pathway sets were significant; these trends should be viewed as hypothesis-generating rather than definitive pathway suppression. The data do not establish on-target nAChR engagement. Overall confidence is medium, driven by concordant pathway and protein evidence for the up-regulated programs and by limited support for broader metabolic or secretory interpretations.

## Findings consistent with expected biology
- Robust activation of a hypoxia-like program (Hallmark HYPOXIA; NES ~2.07, FDR 0).
- Pro-inflammatory signaling via TNFα/NF-κB and IL6/STAT3 (NES ~1.96 and ~1.75, FDR 0 and ~0.013), with protein-level support from IRAK4 and IGFBP1 up-regulation.
- Enrichment of EMT and adhesion/migration signatures (EMT NES ~1.77, FDR ~0.014; GO adhesion/migration NES >2), supported by increases in CEMIP, WASF1, and PRR5.

## Unexpected but plausible findings
- Opposing regulation of IGF-binding proteins: IGFBP1 up-regulated while IGFBP2 down-regulated, indicating a shift in IGF-axis balance under treatment.
- Up-regulation of the Wnt suppressor ZNRF3 despite an EMT signature, suggesting EMT may proceed via routes not requiring canonical Wnt activation (interpret with caution; no Wnt pathway analysis provided).
- Protein-level decreases in components linked to oxidative/peroxisomal metabolism (e.g., NDUFA1, PHYH) and citrate import (SLC13A5), consistent with but not conclusive for reduced oxidative metabolism.
- Decreases in ER translocation-associated proteins (SEC61G, SPCS1) without down pathway enrichment, raising a tentative possibility of reduced secretory capacity.

## Follow-up hypotheses
### Hypothesis 1
- **Hypothesis:** Adiphenine induces NF-κB/STAT3 activation through innate immune signaling independent of direct nAChR engagement.
- **Why interesting:** Strong enrichment of TNFα/NF-κB and IL6/STAT3 programs with IRAK4 up-regulation suggests a cytokine/IL-1R–TLR axis.
- **Suggested validation:** Measure NF-κB and STAT3 phosphorylation/nuclear translocation; quantify IL6/TNF secretion; test sensitivity to IRAK4 or MyD88 inhibitors or knockdown.
- **Priority:** high

### Hypothesis 2
- **Hypothesis:** The treatment elicits a hypoxia-like state that reduces oxidative metabolism.
- **Why interesting:** Hallmark HYPOXIA is strongly enriched; NDUFA1 and PHYH are decreased and SLC13A5 is down, consistent with reduced respiratory/TCA input.
- **Suggested validation:** Assess HIF1α stabilization and target gene induction; perform Seahorse OCR/ECAR assays; survey additional OXPHOS and TCA subunits at protein/transcript levels.
- **Priority:** medium

### Hypothesis 3
- **Hypothesis:** EMT and cell motility are functionally increased following treatment.
- **Why interesting:** EMT and migration/adhesion programs are enriched with supporting proteins (CEMIP, WASF1, PRR5) up.
- **Suggested validation:** Evaluate EMT markers (E-cadherin, vimentin, N-cadherin); perform wound-healing/transwell migration assays and live-cell motility analyses.
- **Priority:** high

### Hypothesis 4
- **Hypothesis:** The IGF signaling axis is reconfigured and may shift downstream PI3K/AKT dynamics.
- **Why interesting:** IGFBP1 up-regulation and IGFBP2 down-regulation indicate altered IGF ligand availability and signaling context.
- **Suggested validation:** Measure IGFBP1/2 secretion (ELISA); assess IGF1R phosphorylation and AKT signaling; test sensitivity to IGF pathway modulation.
- **Priority:** medium

### Hypothesis 5
- **Hypothesis:** Secretory/translocation capacity is modestly reduced.
- **Why interesting:** Decreased SEC61G and SPCS1 suggest possible attenuation of ER translocation machinery.
- **Suggested validation:** Quantify broader translocon/substrate components; use secreted reporter assays (e.g., Gaussia luciferase) and ER stress markers to assess functional impact.
- **Priority:** low

### Hypothesis 6
- **Hypothesis:** EMT occurs without canonical Wnt activation despite ZNRF3 up-regulation.
- **Why interesting:** ZNRF3 increase coincides with EMT enrichment, implying alternative EMT drivers.
- **Suggested validation:** Monitor β-catenin/TCF activity (TOPFlash) and Wnt target genes; perturb NF-κB/STAT3 and assess effects on EMT markers and migration.
- **Priority:** low

## Confidence
medium

## Caution notes
- No significant down-regulated pathway sets were detected; avoid inferring global pathway suppression from individual protein decreases.
- Mechanistic linkage to the stated nAChR target is not demonstrated by these data; observed signatures may reflect downstream or off-target stress responses.
- Metabolic interpretations (reduced mitochondrial/peroxisomal activity, citrate import) are based on a small number of proteins and require broader confirmation.
- Secretory/translocation conclusions from SEC61G/SPCS1 decreases are hypothesis-generating only and need functional validation.
- GO terms for forebrain/cortical migration likely reflect generic migration modules; do not overinterpret as neuron-specific effects without cell-type context.
- Key experimental context (cell type, dose, exposure time) is unspecified; pathway activation magnitude and direction can be context-dependent.
- Single-protein changes in tumor suppressors or signaling regulators (e.g., PTEN) should not be overinterpreted without corroborating pathway evidence.


---

# Alcaftadine

# Drug summary: Alcaftadine

## Executive summary
Proteomic profiling of alcaftadine indicates enrichment of hypoxia-like and TNFα/NF-κB–related programs alongside broad suppression of ribosome biogenesis, rRNA processing, and RNA splicing. Immune involvement is supported by increased IL4R, GRAP, CD68, and CD109. Cytoskeletal/ECM changes are suggested by reduced intermediate filament organization and downregulation of COL6A1, with protein-level increases in caveolar/transport and ECM-related proteins (CAVIN2, FXYD1, SLC16A6, LOXL2, CLEC3B), although no pathway-level upregulation corroborates this. A marked decrease in DKK1 may alter Wnt pathway tone but lacks pathway enrichment support. Overall, the profile is most consistent with a stress-adaptive, cytostatic state; mechanistic attribution to histamine receptor engagement remains uncertain.

## Findings consistent with expected biology
- Engagement of inflammatory/immune signaling, supported by Hallmark TNFA_SIGNALING_VIA_NFKB enrichment (NES ~1.83, FDR ~0.011).
- Upregulation of immune-associated proteins (IL4R, GRAP, CD68, CD109), consistent with the drug’s immunology focus.

## Unexpected but plausible findings
- Strong Hallmark HYPOXIA enrichment (NES ~1.98, FDR ~0.0016), indicating a hypoxia-like stress program.
- Broad downregulation of biosynthetic machinery (RIBOSOME_BIOGENESIS, rRNA processing, RNA splicing) suggestive of reduced translational capacity.
- Reduced intermediate filament organization (GO) with COL6A1 downregulation, consistent with cytoskeletal/ECM remodeling.
- Robust DKK1 decrease (adj p ~4.2e−10) without Wnt pathway enrichment; potential Wnt tone change remains speculative.
- Concordant downregulation of EEF1AKMT1/4, consistent with altered translation-elongation associated machinery, though functional impact is unproven.
- Protein-level hints of membrane/caveolae and ion-transport remodeling (elevated CAVIN2, FXYD1, SLC16A6, LOXL2, CLEC3B) without Reactome/KEGG/GO up corroboration.

## Follow-up hypotheses
### Hypothesis 1
- **Hypothesis:** Alcaftadine induces a stress-adaptive state with concurrent hypoxia-like and NF-κB–related responses that shift cells from growth to stress programs.
- **Why interesting:** Explains the co-occurrence of HYPOXIA and TNFA/NF-κB hallmark enrichments with biosynthetic downregulation.
- **Suggested validation:** Measure HIF-1α/2α stabilization and NF-κB p65 nuclear translocation; use HIF and NF-κB reporter assays; quantify canonical target transcripts and cytokines; assess cell proliferation/viability.
- **Priority:** high

### Hypothesis 2
- **Hypothesis:** Biosynthetic capacity is suppressed via reduced ribosome biogenesis and RNA processing, with potential contribution from lowered EEF1AKMT1/4.
- **Why interesting:** Strong GO downregulation and coordinated decreases in translation-associated modifiers suggest a cytostatic mechanism.
- **Suggested validation:** Assess rRNA transcription (5-EU/BrU incorporation, nucleolar markers), polysome profiling, global translation (SUnSET puromycin assay), and eEF1A methylation status; confirm EEF1AKMT1/4 protein changes by targeted proteomics.
- **Priority:** high

### Hypothesis 3
- **Hypothesis:** DKK1 reduction shifts Wnt/β-catenin signaling tone toward activation in this context.
- **Why interesting:** DKK1 is a key secreted Wnt antagonist; its decrease could impact pathway activity.
- **Suggested validation:** Measure β-catenin stabilization and TCF/LEF reporter activity; quantify Wnt target genes (e.g., AXIN2); measure secreted DKK1; perform rescue with recombinant DKK1 to test pathway dependence.
- **Priority:** medium

### Hypothesis 4
- **Hypothesis:** Caveolae/membrane transport and ECM remodeling are induced as part of the stress response.
- **Why interesting:** Increased CAVIN2, FXYD1, SLC16A6, LOXL2, and CLEC3B suggest membrane/ion transport and ECM reprogramming despite no pathway-level enrichments.
- **Suggested validation:** Immunostain/EM for caveolae (CAV1/CAVIN2), measure lactate export and membrane capacitance, and assay LOXL2 activity and ECM markers; query Reactome/GO with a broader protein set for corroboration.
- **Priority:** medium

### Hypothesis 5
- **Hypothesis:** Intermediate filament network remodeling contributes to altered cell architecture under treatment.
- **Why interesting:** GO downregulation of intermediate filament organization and reduced COL6A1 indicate structural changes.
- **Suggested validation:** Quantify vimentin/keratin isoforms and image IF organization; measure cell mechanics (e.g., AFM) and adhesion; test reversibility after washout.
- **Priority:** medium

## Confidence
medium

## Caution notes
- Conclusions are based on protein abundance and gene-set enrichment; direct activation of NF-κB or HIF is not shown without post-translational/localization evidence.
- Cell context, histamine receptor expression, and dose/time are not provided; antagonists can behave differently depending on baseline agonism and receptor activity.
- Hypoxia-like and biosynthetic suppression signatures may reflect generic stress or cytostatic/cytotoxic effects; viability and stress markers are not reported.
- Membrane/caveolae and ECM remodeling inferences rely on a limited set of upregulated proteins and lack pathway-level support.
- Wnt-related inference rests on DKK1 alone without pathway enrichment; additional Wnt components and functional assays are needed.
- Several effect sizes are modest (~±1 log2FC) despite strong FDRs; biological impact should be interpreted cautiously.
- Attribution to on-target histamine receptor mechanisms is not established; off-target or context-dependent effects remain possible.


---

# Alizapride (hydrochloride)

# Drug summary: Alizapride (hydrochloride)

## Executive summary
Alizapride exposure produced a coherent proteomic pattern centered on lipid pathway remodeling and a small DNA repair/replication-associated signal. Multiple lipid catabolic/turnover factors increased (ABHD5, DDHD1), while sterol and glycosphingolipid biosynthesis/transport components decreased (TM7SF2, UGCG, TSPO), consistent with reduced cholesterol/glycosphingolipid pathways and enhanced membrane lipid turnover. A limited set of DNA repair/replication-associated proteins (RAD51D, EXO1, ZNF365) increased, suggesting a replication stress/DNA repair response, though pathway-level activation is unconfirmed. ECM/cell-surface markers (LOXL2, LAMC3, PROM1) were up, indicating possible cell-state/adhesion changes. Select biosynthetic/processing nodes were reduced (TAF1D for rRNA transcription, SNRPC for splicing), and SEC16B decreased, tentatively pointing to reduced ER export (single-protein). Growth-related modulators (IGFBP2, DLK1) and TEAD4 were down, but concurrent upregulation of transcription/chromatin factors (BRF1, HDAC10, E4F1) argues against a broad proliferation shutdown. No significant pathway enrichments were detected, and on-target GPCR engagement is not evidenced; interpretations therefore rely on coherent protein-level patterns and are presented conservatively.

## Findings consistent with expected biology
- None listed.

## Unexpected but plausible findings
- Lipid remodeling: upregulation of lipolysis/phospholipase-associated proteins (ABHD5, DDHD1) concurrent with downregulation of sterol and glycosphingolipid biosynthesis/transport components (TM7SF2, UGCG, TSPO), indicating a shift away from lipid anabolism.
- Elevation of DNA repair/replication-associated proteins (RAD51D, EXO1, ZNF365), suggestive of a replication stress/DNA repair response despite the drug’s GPCR/neuronal target annotation.
- Upregulation of ECM/adhesion-linked proteins (LOXL2, LAMC3) and the surface marker PROM1, consistent with cell-surface/ECM state changes.
- Selective reduction in RNA production/processing components (TAF1D for rRNA transcription; SNRPC for splicing), compatible with targeted dampening of biosynthetic processes.
- Decreases in growth-associated modulators (IGFBP2, DLK1) and TEAD4, suggesting potential effects on proliferative signaling, though pathway-level conclusions (e.g., Hippo/YAP-TEAD suppression) are not supported.
- Possible ER export attenuation indicated by SEC16B downregulation (single-protein observation; hypothesis-level).

## Follow-up hypotheses
### Hypothesis 1
- **Hypothesis:** Alizapride shifts lipid metabolism toward increased lipolysis/membrane lipid turnover and reduced sterol/glycosphingolipid biosynthesis.
- **Why interesting:** Alters membrane composition and signaling, potentially impacting receptor function, drug uptake, and stress adaptation independently of GPCR engagement.
- **Suggested validation:** Untargeted and targeted lipidomics (cholesterol, desmosterol, GlcCer/ceramide); lipid droplet staining (BODIPY) and ATGL activity assays; rescue with mevalonate/cholesterol; quantify UGCG activity and TM7SF2-dependent sterol markers.
- **Priority:** high

### Hypothesis 2
- **Hypothesis:** Alizapride induces replication stress and activates DNA repair pathways, particularly homologous recombination.
- **Why interesting:** Could sensitize cells to ATR/PARP inhibition and inform combination strategies; indicates off-target genotoxic/replication stress liabilities.
- **Suggested validation:** Assess γH2AX, p-ATR/p-CHK1, RPA/PCNA foci; DNA fiber assays for fork dynamics; comet assay; test synergy/sensitization with ATR or PARP inhibitors.
- **Priority:** high

### Hypothesis 3
- **Hypothesis:** ECM and cell-surface remodeling via LOXL2 and LAMC3 upregulation alters adhesion/motility states.
- **Why interesting:** Changes in ECM interactions can modulate invasion, drug response, and differentiation state.
- **Suggested validation:** Measure LOXL2 activity and collagen crosslinking; adhesion and migration assays; surface PROM1 quantification by flow cytometry; evaluate sensitivity to LOXL2 inhibitors.
- **Priority:** medium

### Hypothesis 4
- **Hypothesis:** Selective suppression of rRNA transcription and pre-mRNA splicing reduces ribosome biogenesis and mRNA processing.
- **Why interesting:** Could reprogram translation and contribute to stress tolerance without globally suppressing proliferation.
- **Suggested validation:** 5-EU metabolic labeling of rRNA; pre-rRNA qPCR; nucleolar marker (fibrillarin) imaging; splicing reporter assays; snRNP component quantification.
- **Priority:** medium

### Hypothesis 5
- **Hypothesis:** ER exit-site function and secretory trafficking are attenuated via reduced SEC16B.
- **Why interesting:** Secretory pathway changes could influence surface receptor abundance and cytokine/ECM secretion.
- **Suggested validation:** VSV-G ts045 trafficking assay; secretion of reporter proteins; COPII component levels; rescue by SEC16B overexpression.
- **Priority:** low

## Confidence
medium

## Caution notes
- No significant pathway enrichments were detected; conclusions rely on coherent protein-level patterns and should be considered provisional.
- Program-level inferences sometimes rest on few proteins (e.g., SEC16B for ER export, TEAD4 for Hippo); treat these as hypotheses rather than established pathway changes.
- Replication stress/DNA repair activation is suggested by limited markers and lacks canonical DDR kinase/phospho evidence; orthogonal assays are needed.
- Lipid remodeling interpretations should be confirmed by lipidomics and functional assays before inferring broader metabolic consequences.
- Interpretation of IGFBP2 and DLK1 decreases as reduced growth signaling is context-dependent and not definitive.
- On-target GPCR/D2 engagement is not demonstrated in this system; observed changes may reflect off-target or generic stress responses.
- Single-omics, single-condition data (no phosphoproteomics, dose, or time course) limit mechanistic specificity and causality.


---

# Almotriptan (malate)

# Drug summary: Almotriptan (malate)

## Executive summary
Proteomics after Almotriptan exposure shows a stress-adaptive profile dominated by a hypoxia-like program (Hallmark HYPOXIA; NES 2.11, FDR 0), accompanied by activation of MAPK/AP-1–like signaling (Hallmark KRAS_SIGNALING_UP; NES 1.69, FDR ~0.037; FOSL1 up) and a modest IL6/JAK/STAT3 signature (NES 1.72, FDR ~0.042). A coordinated reduction in RNA splicing (GO: RNA_SPLICING; NES -1.997, FDR ~0.009) indicates post-transcriptional reprogramming. Protein-level changes include LOXL2 up (consistent with hypoxia-like remodeling), and increases in HLA-B and GRAP. The response comprises balanced up- and down-regulation (25 each), consistent with an adaptive rather than wholesale proteome shift. Mechanistic specificity is limited by absence of Reactome/KEGG enrichments and lack of phospho/target engagement data; several single-protein observations (e.g., RELB down, ciliary components) remain tentative.

## Findings consistent with expected biology
- MAPK/AP-1–like pathway engagement consistent with GPCR signaling (Hallmark KRAS_SIGNALING_UP enrichment; FOSL1 increased).
- A balanced, moderate proteomic response (25 up, 25 down) indicative of stress/adaptive signaling rather than broad proteome remodeling.

## Unexpected but plausible findings
- Robust hypoxia-like program induction (Hallmark HYPOXIA; NES 2.11, FDR 0) with supportive protein-level changes (e.g., LOXL2 up).
- Significant suppression of RNA splicing processes (GO: RNA_SPLICING; NES -1.997, FDR ~0.009), suggesting post-transcriptional adaptation.
- Enrichment of IL6/JAK/STAT3 signaling (NES 1.72, FDR ~0.042) with partial protein-level concordance (GRAP up, HLA-B up).
- Tentative indications of ciliary/centrosome component changes (CEP290, RPGR up; TMEM138 down) without pathway-level support.

## Follow-up hypotheses
### Hypothesis 1
- **Hypothesis:** Almotriptan triggers MAPK/ERK–AP-1 activation downstream of serotonergic GPCR signaling.
- **Why interesting:** Aligns with expected GPCR-to-ERK transactivation and explains KRAS_SIGNALING_UP enrichment and FOSL1 increase.
- **Suggested validation:** Measure pERK and AP-1 activity (FOSL1/c-FOS), perform time-course; test reversal with MEK inhibitors and 5-HT1B/1D antagonists; confirm receptor expression/engagement.
- **Priority:** high

### Hypothesis 2
- **Hypothesis:** The hypoxia-like signature reflects HIF pathway activation or metabolic stress rather than actual hypoxia.
- **Why interesting:** Explains strong Hallmark HYPOXIA and LOXL2 up-regulation; distinguishes true HIF activation from generalized stress.
- **Suggested validation:** Assess HIF-1α stabilization, CA9/VEGFA induction, and hypoxia reporter activity; profile OCR/ECAR for metabolic stress; test sensitivity to HIF inhibitors.
- **Priority:** medium

### Hypothesis 3
- **Hypothesis:** STAT3 activation arises via GPCR cross-talk or autocrine IL6 signaling.
- **Why interesting:** Accounts for IL6_JAK_STAT3 enrichment and increased GRAP/HLA-B; provides a mechanistic link to inflammatory tone.
- **Suggested validation:** Measure pSTAT3 (Y705), STAT3 target transcripts; quantify IL6 secretion; evaluate effects of JAK inhibitors or IL6 neutralization.
- **Priority:** medium

### Hypothesis 4
- **Hypothesis:** Almotriptan suppresses spliceosome/RNA processing as part of a stress-adaptive program.
- **Why interesting:** Explains significant RNA_SPLICING downregulation and may impact transcript isoform usage and translation.
- **Suggested validation:** RNA-seq to detect splicing alterations; immunoblot key spliceosomal components and phospho-SR proteins; test rescue with ISR modulators.
- **Priority:** medium

### Hypothesis 5
- **Hypothesis:** Almotriptan perturbs ciliary biology in this model.
- **Why interesting:** Protein-level changes in CEP290, RPGR (up) and TMEM138 (down) suggest ciliary/centrosomal remodeling.
- **Suggested validation:** Immunofluorescence for primary cilia (acetylated tubulin/ARL13B), quantify ciliation and ciliary length; assess cilia-dependent signaling readouts.
- **Priority:** low

## Confidence
medium

## Caution notes
- Proteomics reports protein abundance, not activation state; pathway activation (e.g., MAPK, STAT3) requires phospho/functional confirmation.
- Mechanistic specificity is limited by lack of Reactome/KEGG enrichments; conclusions rely on Hallmark and a single GO BP result.
- Several interpretations are based on single-protein changes (e.g., RELB, SEC16B, SUFU, ZNRF3, PTEN) and should be considered preliminary.
- The hypoxia hallmark may reflect generalized cellular stress rather than true hypoxia; orthogonal HIF/metabolic assays are needed.
- Cell model context and 5-HT receptor expression/engagement are not provided; off-target or indirect stress responses cannot be excluded.
- Empty pathway sections (e.g., Hallmark_down, GO_up) indicate no significant enrichment detected and should not be over-interpreted.


---

# Ambroxol

# Drug summary: Ambroxol

## Executive summary
Ambroxol exposure elicits a hypoxia-like metabolic rewiring characterized by increased glycolysis alongside activation of lipid/cholesterol and xenobiotic metabolism programs. Concurrently, there is robust innate immune activation with interferon (alpha/gamma) and TNF/NFkB signatures, and strong enrichment of intermediate filament/cytoskeletal remodeling programs that map to keratinocyte/epidermal differentiation gene sets. A broad suppression of RNA splicing/mRNA processing pathways is evident. Protein-level changes support these programs (e.g., LDHAL6B and TXNIP up; ISG20, CXCL16, IL17RC up), with decreases in select mitochondrial complex IV-associated components (MT-CO1, CMC1, DNAJC30), suggesting potential mitochondrial stress but not a coordinated OXPHOS down-regulation. Autophagy-lysosome involvement is plausible but provisional (RUBCN down; HPS1 and AP5M1 up) given the lack of explicit pathway enrichment. No KEGG/Reactome enrichments were available; conclusions are based on Hallmark/GO evidence and should be interpreted accordingly.

## Findings consistent with expected biology
- Upregulation of xenobiotic metabolism programs (Hallmark XENOBIOTIC_METABOLISM).
- Activation of lipid metabolic pathways, including fatty acid metabolism and cholesterol homeostasis (Hallmark FATTY_ACID_METABOLISM, CHOLESTEROL_HOMEOSTASIS).

## Unexpected but plausible findings
- Hypoxia-like response with increased glycolysis (Hallmark HYPOXIA and GLYCOLYSIS up) accompanied by reduced abundance of select mitochondrial complex IV–related proteins (MT-CO1, CMC1, DNAJC30).
- Innate immune and inflammatory activation (INTERFERON_ALPHA/GAMMA_RESPONSE, INFLAMMATORY_RESPONSE, TNFA_SIGNALING_VIA_NFKB up) with ISG20, CXCL16, and IL17RC increased.
- Strong intermediate filament/cytoskeletal remodeling mapped to keratinocyte/epidermal differentiation gene sets (multiple GO terms with high NES), potentially reflecting structural reorganization rather than true lineage differentiation.
- Global downregulation of RNA splicing and mRNA processing programs (multiple GO BP terms significantly decreased).
- KRAS_SIGNALING_DN gene set enrichment, indicating KRAS-off-like gene set behavior without direct evidence of KRAS pathway inhibition.

## Follow-up hypotheses
### Hypothesis 1
- **Hypothesis:** Ambroxol induces a pseudo-hypoxic state that shifts metabolism toward glycolysis and imposes mild mitochondrial stress.
- **Why interesting:** Aligns with strong HYPOXIA/GLYCOLYSIS enrichment and decreased abundance of selected complex IV–related proteins; metabolic shifts may be relevant to efficacy or stress tolerance.
- **Suggested validation:** Measure ECAR/OCR (Seahorse) for glycolytic/OXPHOS balance; quantify lactate; assess HIF1A stabilization; assay complex IV activity and broader respiratory chain protein levels.
- **Priority:** high

### Hypothesis 2
- **Hypothesis:** Ambroxol activates innate antiviral/inflammatory signaling (IFN and NFkB axes).
- **Why interesting:** Robust Hallmark enrichments with supporting ISG/chemokine increases suggest immunomodulatory effects that could impact safety and disease-relevant pathways.
- **Suggested validation:** qPCR or proteomics of ISGs and NFkB targets; STAT1/STAT2 and NFkB p65 phosphorylation; NFkB/ISRE reporter assays; cytokine/chemokine secretion profiling.
- **Priority:** high

### Hypothesis 3
- **Hypothesis:** Ambroxol engages lysosome–autophagy pathways by relieving RUBCN-mediated inhibition and altering endolysosomal trafficking.
- **Why interesting:** RUBCN down with HPS1 and AP5M1 up is consistent with autophagy/lysosome involvement and aligns with the drug’s putative lysosomal/autophagy relevance.
- **Suggested validation:** Assess LC3-II and p62/SQSTM1 turnover with/without bafilomycin; TFEB/TFE3 nuclear translocation; LAMP1/lysosomal protease levels; monitor autophagic flux by reporters.
- **Priority:** medium

### Hypothesis 4
- **Hypothesis:** Ambroxol induces cytoskeletal remodeling programs that map to keratinocyte/epidermal differentiation without actual lineage conversion.
- **Why interesting:** Multiple high-NES GO terms for intermediate filament and epidermal programs may reflect structural adaptation affecting adhesion and barrier-related functions.
- **Suggested validation:** Immunostaining/Western for intermediate filament proteins (e.g., keratins, vimentin); adhesion/spreading assays; quantification of integrin subunits and focal adhesion markers.
- **Priority:** medium

### Hypothesis 5
- **Hypothesis:** Ambroxol suppresses components of the RNA splicing machinery as part of a stress-adaptive program.
- **Why interesting:** Consistent, strong down-enrichment of RNA splicing/mRNA processing terms suggests altered transcript processing that could reshape gene expression.
- **Suggested validation:** Targeted proteomics for spliceosome components; RNA-seq to assess intron retention and alternative splicing; splicing reporter assays under drug exposure.
- **Priority:** medium

## Confidence
medium

## Caution notes
- Conclusions are derived from proteomics and Hallmark/GO enrichments; signaling activity is inferred indirectly and requires orthogonal validation (e.g., phosphoproteomics, reporters).
- No KEGG or Reactome enrichments were available; mechanistic specificity relies on Hallmark/GO sets.
- Evidence for reduced oxidative phosphorylation is limited to a few proteins without pathway-level down-enrichment; interpret mitochondrial effects cautiously.
- Autophagy engagement is provisional (RUBCN down; HPS1/AP5M1 up) without direct autophagy pathway enrichment or canonical flux markers.
- KRAS_SIGNALING_DN reflects gene set behavior and should not be interpreted as direct KRAS pathway inhibition.
- Cell model/context is unspecified; strong epidermal/keratinocyte signatures may represent cytoskeletal remodeling rather than true differentiation.


---

# Amcinonide

# Drug summary: Amcinonide

## Executive summary
Amcinonide (glucocorticoid receptor agonist) produces a proteomic profile marked by robust suppression of the MYC target program (Hallmark MYC_TARGETS_V1 down) and reduced apical surface/junction and pericellular matrix components (Hallmark APICAL_SURFACE down). These hallmark-level signals are supported by decreases in CDCA7, PATJ, NPNT, ITIH5, SULF2, and SPP1, consistent with an anti-proliferative state and epithelial barrier/junction remodeling. Protein-level induction of xenobiotic/drug-metabolizing enzymes and transporters (e.g., CYP1A1, CYP3A7, SLC16A6, SLCO4A1, ABHD2) suggests enhanced metabolic/transport capacity, though no up-regulated pathway sets were enriched. Decreased NR3C1 protein is consistent with ligand engagement. Additional coordinated decreases in PCSK9, APOB, and APOA5 point to reduced lipoprotein assembly/trafficking in this model, and HAMP reduction implies altered iron handling; these are plausible but context-dependent and not corroborated by pathway enrichment. Overall, the data support GR-driven anti-proliferative and junction/ECM remodeling effects with medium confidence, while several metabolic and secretory changes remain hypothesis-generating.

## Findings consistent with expected biology
- Strong suppression of the MYC target program (Hallmark MYC_TARGETS_V1 down; supported by decreased CDCA7), consistent with an anti-proliferative GR response.
- Reduction of apical surface/junction and ECM-related features (Hallmark APICAL_SURFACE down), supported by decreases in PATJ, NPNT, ITIH5, SULF2, and SPP1.
- Protein-level induction of xenobiotic metabolism and transport (CYP1A1, CYP3A7, SLC16A6, SLCO4A1, ABHD2), directionally consistent with nuclear receptor–mediated metabolic reprogramming.
- Decrease in glucocorticoid receptor (NR3C1) protein, consistent with ligand-induced receptor down-regulation (supportive but not standalone proof of target engagement).

## Unexpected but plausible findings
- Coordinated decreases in lipoprotein-related proteins (PCSK9, APOB, APOA5), suggesting dampened lipoprotein assembly/trafficking in this system (no pathway enrichment; cell-type dependent).
- Concurrent induction of CYP1A1 with CYP3A7, indicating activation of multiple xenobiotic axes (including a non-classical GR-linked enzyme), pointing to potential secondary receptor involvement.
- Marked reduction of HAMP (hepcidin), suggesting altered iron homeostasis (single-marker support).
- Decrease in DKK1 (Wnt antagonist) without Wnt pathway enrichment; the mechanistic significance is unclear.

## Follow-up hypotheses
### Hypothesis 1
- **Hypothesis:** GR activation represses MYC-driven programs to induce an anti-proliferative state.
- **Why interesting:** Explains the hallmark MYC_TARGETS_V1 suppression and could account for broad growth restraint under amcinonide.
- **Suggested validation:** Measure canonical MYC target transcripts and proliferation/cell-cycle markers; test rescue with GR antagonism or NR3C1 knockdown; assess viability/proliferation assays.
- **Priority:** high

### Hypothesis 2
- **Hypothesis:** GR signaling remodels epithelial junctions and pericellular matrix, reducing apical/adhesion components and modifying barrier properties.
- **Why interesting:** Aligns with APICAL_SURFACE hallmark reduction and decreased PATJ/NPNT/ITIH5/SULF2/SPP1, relevant for tissue integrity and inflammation.
- **Suggested validation:** Imaging of tight junctions and adhesion complexes; transepithelial electrical resistance (if applicable); quantify junction/ECM transcripts and proteins over time; GR dependence via antagonist or knockdown.
- **Priority:** medium

### Hypothesis 3
- **Hypothesis:** Amcinonide co-activates secondary xenobiotic receptors (e.g., AhR/PXR/CAR), driving CYP1A1 and CYP3A7 induction.
- **Why interesting:** Would explain simultaneous induction of enzymes typically controlled by distinct nuclear receptors and refine mechanistic attribution.
- **Suggested validation:** Receptor-specific reporter assays; pharmacologic antagonism or siRNA of candidate receptors; measure canonical downstream transcripts and enzyme activities.
- **Priority:** medium

### Hypothesis 4
- **Hypothesis:** GR exposure attenuates lipoprotein assembly/secretion in this cell context.
- **Why interesting:** Coordinated decreases in PCSK9/APOB/APOA5 suggest a shift in secretory lipid handling that could impact metabolic phenotypes.
- **Suggested validation:** Quantify intracellular and secreted ApoB and PCSK9 (ELISA/western); assess ApoB-containing particle secretion; lipidomics; test GR dependence and cell-type specificity.
- **Priority:** low

### Hypothesis 5
- **Hypothesis:** GR signaling lowers hepcidin and alters cellular iron handling.
- **Why interesting:** HAMP reduction implies a shift in iron regulatory balance with potential immunometabolic consequences.
- **Suggested validation:** Measure hepcidin transcripts/protein; assess ferritin, transferrin receptor, and ferroportin levels; perform iron uptake/export assays; evaluate GR dependence.
- **Priority:** low

## Confidence
medium

## Caution notes
- Up-regulated pathway enrichments are absent; several functional inferences rely on a small set of protein markers and should be considered hypothesis-generating.
- NR3C1 protein reduction is consistent with target engagement but is not definitive without orthogonal evidence (e.g., nuclear translocation or canonical GR target induction).
- Apparent effects on lipoprotein metabolism (PCSK9/APOB/APOA5) and iron handling (HAMP) may be highly cell-type specific and may reflect broader changes in secretory state.
- Induction of CYP1A1 suggests possible involvement of non-GR xenobiotic receptors; mechanistic attribution to GR alone should be made cautiously.
- Interpretation of GO term ‘negative regulation of mRNA metabolic process’ is ambiguous and lacks specific protein-level support; avoid overinterpretation.
- Protein-only data and unspecified cell/tissue context limit generalizability; presence of hepatic/secretory proteins and CYP3A7 suggests a context that may not translate to other systems.


---

# Amiodarone (hydrochloride)

# Drug summary: Amiodarone (hydrochloride)

## Executive summary
Amiodarone (hydrochloride) induces a coherent proteomic program characterized by: (1) strong activation of cholesterol/sterol biosynthesis and lipid homeostasis (Hallmark CHOLESTEROL_HOMEOSTASIS; multiple GO sterol processes, all FDR 0), supported by upregulation of SC5D, APOA5, ANGPTL8, GAL3ST1, and PCYOX1L; (2) robust enrichment of inflammatory/cytokine signaling (IL6–JAK–STAT3, TNFα–NFκB) with complement and coagulation programs, accompanied by increased immune modulators/receptors (IL4R, IL10RB, TNFRSF12A, RNF128, CEMIP); (3) broad suppression of proliferative and biosynthetic capacity (down E2F/MYC/G2M and mitotic processes; marked downregulation of ribosome biogenesis and both cytoplasmic and mitochondrial translation), consistent with a cytostatic, stress-adaptive state; and (4) membrane remodeling signatures, including enrichment for membrane protein ectodomain proteolysis and moderate pathway-level signals for angiogenesis/artery morphogenesis with increased NRP2 and DLK1. Moderate hormone-response (estrogen/androgen) and Hedgehog pathway enrichments are observed; autophagy involvement is suggested by TP53INP2 and selected ubiquitin/adaptor proteins but lacks pathway-level support. Decreases in several ciliogenesis-related proteins (IFT88, TMEM138, TRAF3IP1) raise the possibility of reduced ciliogenesis; this and the Hedgehog signal should be treated as tentative. Overall, the proteome points to lipid/cholesterol remodeling with an inflammatory secretory program and coordinated growth/translation repression.

## Findings consistent with expected biology
- Marked induction of cholesterol/sterol biosynthesis and lipid homeostasis programs with supporting protein increases (e.g., SC5D, APOA5, ANGPTL8, GAL3ST1, PCYOX1L).
- Robust enrichment of inflammatory/cytokine pathways (IL6–JAK–STAT3, TNFα–NFκB) together with complement and coagulation signatures, supported by increased immune receptors/modulators (IL4R, IL10RB, TNFRSF12A, RNF128, CEMIP).
- Coordinated suppression of proliferative programs (down E2F/MYC/G2M and mitotic processes) and global downshift of protein synthesis machinery (ribosome biogenesis, rRNA processing, cytoplasmic and mitochondrial translation).
- Membrane remodeling with enrichment for membrane protein ectodomain proteolysis; moderate enrichment for angiogenesis/artery morphogenesis at the pathway level.

## Unexpected but plausible findings
- Hormone-response pathway enrichments (estrogen early/late, androgen) with TFAP2C upregulation.
- Hedgehog signaling pathway enrichment despite decreased levels of several ciliogenesis-associated proteins (IFT88, TMEM138, TRAF3IP1), suggesting a potential cilia–Hedgehog disconnect.
- Limited autophagy-related protein signals (e.g., TP53INP2, NDFIP1, ZNRF3) without explicit autophagy pathway enrichment.

## Follow-up hypotheses
### Hypothesis 1
- **Hypothesis:** Amiodarone triggers SREBP-driven sterol/cholesterol biosynthesis as a compensatory response to membrane/lipid perturbation.
- **Why interesting:** Strong, concordant enrichment of sterol biosynthesis/homeostasis pathways (Hallmark and GO, FDR 0) and upregulation of multiple sterol/lipoprotein-related proteins indicate a regulated lipid compensation program.
- **Suggested validation:** Assess SREBP1/2 activation (precursor cleavage and nuclear forms), measure canonical target transcripts/proteins (e.g., HMGCR, FDFT1), perform lipidomics (cholesterol, sterol intermediates), and test suppression by 25-hydroxycholesterol or SREBP/S1P inhibitors.
- **Priority:** high

### Hypothesis 2
- **Hypothesis:** The drug induces a stress-associated secretory phenotype driven by IL6–JAK–STAT3 and NF-κB signaling.
- **Why interesting:** Highly significant enrichments for IL6–JAK–STAT3, TNFα–NFκB, inflammatory response, complement, and coagulation, with increased cytokine receptors/modulators, support pathway engagement.
- **Suggested validation:** Measure phospho-STAT3 (Y705) and NF-κB p65 nuclear translocation; quantify secreted cytokines (e.g., IL6, TNFα) and complement factors; test reversibility with JAK or IKK inhibitors and evaluate proteomic/phenotypic rescue.
- **Priority:** high

### Hypothesis 3
- **Hypothesis:** Amiodarone enforces a cytostatic program with repression of ribosome biogenesis and translation.
- **Why interesting:** Downregulation of E2F/MYC/G2M, mitotic processes, and strong suppression of ribosome biogenesis and translation (cytoplasmic and mitochondrial) indicate coordinated growth and biosynthesis arrest.
- **Suggested validation:** Perform cell-cycle profiling (flow cytometry), EdU incorporation for S-phase entry, and polysome or puromycin incorporation assays; monitor rRNA processing markers and MYC/E2F target expression.
- **Priority:** high

### Hypothesis 4
- **Hypothesis:** Amiodarone increases ectodomain shedding/surface receptor turnover via metalloprotease activity.
- **Why interesting:** Enrichment for membrane protein ectodomain proteolysis suggests enhanced shedding, potentially contributing to the inflammatory/secretory phenotype.
- **Suggested validation:** Quantify soluble ectodomains (e.g., IL6R, TNFRSF members) in conditioned media by ELISA or mass spectrometry; assess ADAM/TACE activity and test sheddase inhibitors; perform surfaceome proteomics.
- **Priority:** medium

### Hypothesis 5
- **Hypothesis:** The drug reduces ciliogenesis while engaging Hedgehog-related gene sets via non-canonical or context-specific mechanisms.
- **Why interesting:** Hedgehog pathway enrichment co-occurs with decreased ciliogenesis-associated proteins, an atypical combination that may reflect deciliation or alternative pathway wiring.
- **Suggested validation:** Image primary cilia (acetylated α-tubulin/ARL13B), monitor GLI-reporter activity, and examine SMO/GLI localization; test sensitivity to SMO inhibitors/agonists and assess additional ciliary markers.
- **Priority:** medium

### Hypothesis 6
- **Hypothesis:** Hormone-response pathway enrichment reflects cross-talk with nuclear receptor programs.
- **Why interesting:** Moderate estrogen/androgen hallmark enrichments and TFAP2C upregulation suggest context-dependent nuclear receptor signaling.
- **Suggested validation:** Determine ER/AR status and activity (reporter assays), test effects of ER/AR antagonists on the proteomic signature, and profile hormone-responsive transcripts.
- **Priority:** low

## Confidence
high

## Caution notes
- Pathway enrichments reflect ranked protein-level changes mapped to gene sets and indicate program-level shifts rather than definitive pathway activation.
- Despite the drug’s annotation including autophagy, there is no explicit autophagy pathway enrichment; avoid concluding autophagy induction without orthogonal evidence.
- Proteomics alone cannot assign a specific cell-cycle checkpoint arrest; validate phase-specific effects with cell-cycle assays.
- Inference of increased receptor shedding from ectodomain proteolysis enrichment is indirect and requires detection of soluble ectodomains or sheddase activation.
- Interpretation of inflammatory, complement, and coagulation signatures may depend on sample type (cell lysate vs secretome), which is not specified.
- Hedgehog–cilia observations may be cell-type specific; reduced ciliogenesis is inferred from a limited set of proteins and should be confirmed by imaging and additional markers.
- Hormone-response enrichments could reflect pathway cross-talk; confirm receptor status and dependency before inferring hormone-driven mechanisms.
- mTORC1 and EMT enrichments are moderate and should not be presented as primary mechanisms.


---

# Amisulpride

# Drug summary: Amisulpride

## Executive summary
Amisulpride elicited a coherent proteomic signature characterized by strong suppression of RNA maturation pathways (mRNA splicing/processing, rRNA processing) and ribosomal small subunit biogenesis, together with marked downregulation of intermediate filament organization and epithelial structural components. In parallel, a significant Hallmark cholesterol homeostasis program was upregulated with concordant increases in lipid-handling enzymes, indicating lipid/membrane remodeling. These conclusions are supported by multiple GO biological process down-enrichments and a Hallmark up signal with matching protein-level changes (e.g., POLR2K, RSL24D1, RPL36AL down; SMPDL3A, PANK3, CERK, ABHD2 up). Protein-only increases in ER/secretory stress-related markers (AGR2, HERPUD2), vesicle trafficking (VAMP4), and oxidative repair (MSRA) suggest, but do not prove, ER/secretory remodeling. No GPCR or neuronal pathway enrichments were detected, so direct dopamine receptor pathway engagement is not evident at the proteome level. Overall, the data support a shift toward lipid/cholesterol homeostasis with concurrent dampening of RNA processing and ribosome biogenesis and attenuation of epithelial structural programs, with additional remodeling/stress features remaining preliminary.

## Findings consistent with expected biology
- None listed.

## Unexpected but plausible findings
- Coordinated downregulation of RNA splicing/processing and rRNA processing with reduced ribosomal small subunit biogenesis (multiple GO BP down enrichments) and concordant protein decreases (e.g., POLR2K, RSL24D1, RPL36AL).
- Robust suppression of intermediate filament organization and related epithelial structural processes, consistent with decreases in KRT14, CLDN12, and ACTN2.
- Significant activation of cholesterol homeostasis (Hallmark NES 1.82, FDR ~0.034) with upregulation of lipid-handling enzymes (SMPDL3A, PANK3, CERK, ABHD2), indicating membrane/lipid remodeling.
- Divergent epithelial state markers (S100A7 up; S100A14 and KRT14 down) consistent with an epithelial differentiation/stress shift, albeit without pathway-level up-enrichment.
- Notable single-protein decreases in IGFBP2 and SLC13A5 that may have secretory/metabolic implications, but mechanistic interpretation remains uncertain.

## Follow-up hypotheses
### Hypothesis 1
- **Hypothesis:** Amisulpride suppresses RNA processing and ribosome biogenesis via activation of a translational stress axis (e.g., nucleolar stress and/or integrated stress response), reducing effective translational capacity.
- **Why interesting:** Explains the broad downregulation of RNA/ribosome biogenesis pathways and could underlie functional phenotypes independent of proliferation changes.
- **Suggested validation:** Assess pre-rRNA processing by northern blot (ITS1/ITS2), nucleolar morphology and fibrillarin levels; measure eIF2α phosphorylation and ATF4 induction; perform polysome profiling to quantify global translation.
- **Priority:** high

### Hypothesis 2
- **Hypothesis:** Cholesterol homeostasis upregulation reflects SREBP pathway activation that drives membrane lipid remodeling under amisulpride exposure.
- **Why interesting:** Provides a mechanistic link for the Hallmark cholesterol signal and lipid enzyme increases, with potential consequences for membrane protein trafficking and signaling.
- **Suggested validation:** Monitor SREBF1/2 cleavage and nuclear localization; quantify canonical SREBP target transcripts/proteins (e.g., HMGCR, LDLR, FDFT1); measure cellular free cholesterol (filipin staining) and test sensitivity to 25-hydroxycholesterol.
- **Priority:** medium

### Hypothesis 3
- **Hypothesis:** Loss of intermediate filament and tight junction components reduces epithelial adhesion/barrier function, indicating structural remodeling.
- **Why interesting:** Aligns with strong GO downregulation of intermediate filament organization and protein-level decreases in KRT14 and CLDN12; functional consequences would be biologically meaningful.
- **Suggested validation:** Immunofluorescence for keratins and claudins, cell morphology and adhesion assays, and transepithelial electrical resistance (TEER) where applicable.
- **Priority:** medium

### Hypothesis 4
- **Hypothesis:** Downregulation of SLC13A5 limits citrate entry, shifting metabolism away from de novo lipogenesis toward lipid remodeling/uptake.
- **Why interesting:** Integrates reduced citrate import with increased lipid-handling proteins and the cholesterol hallmark, offering a metabolic explanation for lipid remodeling.
- **Suggested validation:** Measure citrate uptake and cytosolic acetyl-CoA; perform 13C-glucose/13C-acetate tracing to assess de novo lipogenesis; untargeted lipidomics; rescue with exogenous acetate or citrate.
- **Priority:** low

## Confidence
medium

## Caution notes
- ER/secretory remodeling and adhesion/secretion changes are suggested by a limited set of protein increases (AGR2, HERPUD2, VAMP4, MSRA) without pathway-level up-enrichment and should be treated as hypotheses.
- Inferences about a less proliferative state are not supported by direct proliferation or cell-cycle signatures (e.g., E2F/MYC hallmarks); avoid overinterpreting RNA/ribosome suppression in this context.
- Interpret GO categories labeled as 'negative regulation' cautiously; primary support should rely on core process terms (e.g., RNA splicing, rRNA processing) showing downregulation.
- No GPCR or neuronal pathway enrichments were detected; direct dopamine receptor pathway engagement is not evidenced at the proteome level, and effects may be downstream or off-target.
- The absence of GO BP up and Hallmark down signals limits confident assertions about additional upregulated programs beyond cholesterol homeostasis.
- Cell type/context is not provided; interpretation of epithelial markers (e.g., S100A7, KRT14, CLDN12) may be context dependent.
- Prominent single-protein changes (e.g., IGFBP2) may reflect altered secretion or extracellular protein levels; confirm with orthogonal assays (e.g., ELISA, conditioned media analyses).


---

# Amodiaquine (dihydrochloride)

# Drug summary: Amodiaquine (dihydrochloride)

## Executive summary
Proteomic profiling of Amodiaquine (dihydrochloride) reveals a robust anti-proliferative response characterized by broad suppression of E2F/MYC programs, G2/M checkpoint, and mitotic spindle components, with concordant decreases in key mitotic regulators. In parallel, there is strong activation of inflammatory/stress signaling (IL6–JAK–STAT3, TNFA via NFKB) and context-dependent TGF-β/EMT/angiogenesis modules. Lipid and cholesterol homeostasis pathways (sterol/lipoprotein metabolism, transport) are upregulated, alongside xenobiotic metabolism, heme metabolism, and coagulation, consistent with a generalized drug/stress adaptation and membrane remodeling. Increased abundance of cytokine receptors and ECM/adhesion-related proteins, together with signatures of membrane protein ectodomain proteolysis and APP catabolic processes, suggests enhanced membrane protein processing. Unexpectedly, neuronal projection guidance annotations and neuronal membrane-associated proteins are induced, and several chromatin/epigenetic-associated factors decrease. Overall, the data support an anti-proliferative, pro-inflammatory, and lipid/membrane remodeling response to amodiaquine; direct linkage to histone methyltransferase inhibition remains unproven.

## Findings consistent with expected biology
- Broad suppression of proliferation and cell-cycle programs (Hallmark E2F_TARGETS, G2M_CHECKPOINT, MITOTIC_SPINDLE down) with decreased mitotic regulators (e.g., CDC7, CDC25B, PTTG1, ESPL1, KIF18A), consistent with anti-proliferative activity.
- Activation of inflammatory cytokine signaling (IL6–JAK–STAT3, INFLAMMATORY_RESPONSE, TNFA_SIGNALING_VIA_NFKB up) with increased cytokine receptors (IL4R, IL10RB, TNFRSF12A).
- Upregulation of cholesterol/lipid metabolism and transport pathways (Hallmark CHOLESTEROL_HOMEOSTASIS; GO sterol/lipoprotein processes), consistent with membrane/lipid remodeling under drug exposure.
- Enrichment of xenobiotic metabolism, heme metabolism, and coagulation pathways, indicating a generalized stress/injury response.
- Elevated membrane protein ectodomain proteolysis and APP catabolic process signatures with increased trafficking/proteostasis-associated proteins (e.g., NDFIP1, TP53INP2, RNF128, SPINT1, LAPTM4A), indicating enhanced membrane protein processing.

## Unexpected but plausible findings
- Enrichment of neuron projection guidance and APP catabolic process with strong induction of neuronal membrane-associated proteins (TMEM59L, NRSN2, CELSR2, NRP2) in a non-neuronal context.
- Concurrent upregulation of both pro- and anti-inflammatory receptors (IL4R, IL10RB, TNFRSF12A), suggesting a mixed immunomodulatory state.
- Downregulation of chromatin/epigenetic-associated factors (PHF19, ANTKMT, ZNHIT3) alongside proliferation suppression, without direct evidence of on-target epigenetic inhibition.
- Induction of EMT/TGF-β/angiogenesis signatures consistent with tissue remodeling; interpretation as a wound-healing–like program is plausible but cell-context dependent.

## Follow-up hypotheses
### Hypothesis 1
- **Hypothesis:** Amodiaquine slows proliferation by suppressing DNA replication/mitotic entry programs, leading to reduced S–G2/M progression.
- **Why interesting:** Clarifies whether the strong anti-proliferative proteomic signature reflects functional cell-cycle slowdown relevant to efficacy and safety.
- **Suggested validation:** Time-course cell-cycle profiling by flow cytometry (PI or DAPI with EdU/BrdU), mitotic index (pH3-Ser10), and immunoblotting for cyclins/CDK substrates (Cyclin B1, p21, p53); assess dose–response.
- **Priority:** high

### Hypothesis 2
- **Hypothesis:** Amodiaquine activates IL6–JAK–STAT3 and TNF–NFκB signaling through stress-responsive pathways, driving the observed cytokine receptor upregulation.
- **Why interesting:** Defines inflammatory signaling consequences that may influence survival, differentiation, or toxicity pathways.
- **Suggested validation:** Measure STAT3 Tyr705 phosphorylation and p65 nuclear translocation; reporter assays for STAT3/NFκB; multiplex cytokine secretion; test reversibility with JAK or IKK inhibitors.
- **Priority:** high

### Hypothesis 3
- **Hypothesis:** Amodiaquine perturbs cellular cholesterol flux and sterol regulatory networks, contributing to membrane/lipid remodeling.
- **Why interesting:** Lipid remodeling may modulate membrane protein processing, trafficking, and drug disposition.
- **Suggested validation:** Assess SREBP1/2 processing and LXR/SREBP target expression; filipin staining for free cholesterol; LDL uptake assays; lysosomal pH (LysoTracker) and cholesterol trafficking markers; quantify APOA5 secretion.
- **Priority:** medium

### Hypothesis 4
- **Hypothesis:** Increased ectodomain shedding and APP processing reflect heightened metalloprotease/secretase activity and membrane protein turnover.
- **Why interesting:** Alters surface receptor signaling and may explain concurrent changes in cytokine/adhesion receptors.
- **Suggested validation:** Measure ADAM10/17 activity and sAPPα/β release; monitor shedding of IL4R/TNFRSF12A; track endocytosis/trafficking by live-cell imaging; optionally assess autophagic flux (LC3-II turnover with bafilomycin) as supportive evidence.
- **Priority:** medium

### Hypothesis 5
- **Hypothesis:** Amodiaquine modulates epigenetic complexes (e.g., Polycomb-associated) indirectly, contributing to downregulation of chromatin factors.
- **Why interesting:** Links proteomic changes (e.g., PHF19 decrease) to chromatin state and may relate to the putative histone methyltransferase axis.
- **Suggested validation:** Quantify H3K27me3 and other histone marks; assess PRC2 component abundance and occupancy (ChIP-qPCR/ChIP-seq); evaluate expression/activity of ANTKMT and related chromatin factors.
- **Priority:** medium

## Confidence
high

## Caution notes
- Conclusions are based on steady-state protein abundance and enrichment analyses; pathway activation and cell-cycle arrest require functional and phospho-signaling validation.
- Cell model, dose, and timepoint are unspecified; off-target or lysosomotropic effects may contribute to the observed signatures.
- Inferring increased autophagy/lysosomal turnover from limited markers is tentative; autophagic flux assays are needed for confirmation.
- EMT/angiogenesis and neuronal annotation enrichments may reflect generic adhesion/trafficking remodeling and are highly cell-context dependent.
- No direct evidence links the proteomic changes to histone methyltransferase inhibition; epigenetic factor downregulation is suggestive but not proof of on-target activity.
- Hallmark ‘Apoptosis’ enrichment indicates a stress program but does not establish apoptotic cell death without orthogonal assays.


---

# Anamorelin

# Drug summary: Anamorelin

## Executive summary
Proteomic profiling of the GHSR-targeting GPCR compound Anamorelin reveals a coherent activation of cytokine/JAK–STAT and inflammatory programs (IL6–JAK–STAT3, IL2–STAT5, interferon-α), with concordant increases in cytokine receptor/regulator proteins (IL4R, IL10RB, RNF128, NDFIP1). Sterol/cholesterol and broader steroid/isoprenoid biosynthetic processes are strongly induced, accompanied by increases in lipid trafficking/membrane-associated proteins (LRP10, LRP11) and tight junction organization (e.g., CLDN12). TGF-β/BMP-associated signals are enriched with supportive increases in BMPR2, DLK1, and ZNRF3. In contrast, rRNA processing, ribosome biogenesis, and RNA splicing pathways are markedly suppressed, and intermediate filament/keratin components are reduced (e.g., KRT14, KRT79), indicating cytoskeletal remodeling. These signatures are pathway-level and protein-concordant, but no direct evidence of proximal GHSR engagement or the immediate upstream cause of STAT activation is provided in this dataset. Downregulated programs are inferred from GO enrichments despite an empty Hallmark-down section, and some pathway labels (e.g., osteoclast differentiation) likely reflect shared signaling modules rather than lineage transitions.

## Findings consistent with expected biology
- None listed.

## Unexpected but plausible findings
- Robust activation of cytokine/JAK–STAT and inflammatory programs (IL6–JAK–STAT3, IL2–STAT5, interferon-α) with increased cytokine receptors/regulators (IL4R, IL10RB, RNF128, NDFIP1).
- Pronounced upregulation of sterol/cholesterol/steroid biosynthesis and lipid homeostasis, with increased LRP10/LRP11 and tight junction organization (CLDN12).
- Engagement of TGF-β/BMP-associated programs with increases in BMPR2, DLK1, and ZNRF3.
- Strong suppression of rRNA processing, ribosome biogenesis and RNA splicing, consistent with reduced biosynthetic/translation capacity.
- Downregulation of keratin/intermediate filament networks (e.g., KRT14, KRT79), indicating cytoskeletal remodeling.

## Follow-up hypotheses
### Hypothesis 1
- **Hypothesis:** Anamorelin induces secondary autocrine/paracrine cytokine signaling that drives JAK–STAT activation.
- **Why interesting:** Explains the strong IL6–JAK–STAT3/IL2–STAT5 and interferon-α signatures and the upregulation of IL4R and IL10RB without direct evidence of proximal GPCR-to-STAT coupling.
- **Suggested validation:** Measure secreted cytokines (e.g., IL6, IFN-α) and STAT3/STAT5 phosphorylation; test sensitivity to JAK inhibitors; include secretome profiling or multiplex cytokine assays.
- **Priority:** high

### Hypothesis 2
- **Hypothesis:** Anamorelin shifts cellular lipid metabolism toward cholesterol/sterol biosynthesis, altering membrane composition and junctional organization.
- **Why interesting:** Integrates strong Hallmark/GO enrichments with increases in LRP10/LRP11 and CLDN12, potentially influencing signaling platforms and barrier properties.
- **Suggested validation:** Perform lipidomics and SREBP target analyses; assess membrane order (e.g., Laurdan), tight junction integrity (TEER or permeability), and sensitivity to statins/SREBP inhibitors.
- **Priority:** medium

### Hypothesis 3
- **Hypothesis:** Global translational output is reduced despite activation of selected signaling pathways.
- **Why interesting:** Multiple high-confidence down-enrichments in rRNA processing, ribosome biogenesis, and splicing suggest a coordinated decrease in protein synthesis capacity.
- **Suggested validation:** Quantify translation by puromycin incorporation (SUnSET) or polysome profiling; assess 18S/28S rRNA levels and rRNA transcription; evaluate eIF2α phosphorylation.
- **Priority:** high

### Hypothesis 4
- **Hypothesis:** TGF-β/BMP pathway engagement contributes to the observed remodeling state.
- **Why interesting:** Hallmark TGF_BETA_SIGNALING enrichment with BMPR2, DLK1, and ZNRF3 upregulation suggests pathway activation that could interact with inflammatory and metabolic reprogramming.
- **Suggested validation:** Measure SMAD2/3 and SMAD1/5/8 phosphorylation; quantify TGF-β/BMP ligands; test pathway inhibition (e.g., receptor kinase inhibitors) for impact on proteomic signatures.
- **Priority:** medium

## Confidence
medium

## Caution notes
- No direct evidence of GHSR target engagement is provided by this proteomic dataset; proximal GPCR signaling events are not measured.
- Upstream cause of JAK–STAT activation remains unresolved; both secondary cytokine signaling and indirect crosstalk are plausible.
- Pathway enrichments (e.g., osteoclast differentiation, morphogenesis) may reflect shared signaling modules rather than true lineage changes in this system.
- Reduced ribosome biogenesis/rRNA processing and splicing imply, but do not prove, decreased translation; functional assays are needed for confirmation.
- Inferences about membrane remodeling and trafficking are based on lipid biosynthesis/tight junction signatures; direct functional evidence is not provided here.
- Hallmark down-regulated sets were not available; conclusions on suppressed programs rely on GO BP down enrichments and protein-level decreases.


---

# Andrographolide

# Drug summary: Andrographolide

## Executive summary
Proteomics after Andrographolide exposure shows a robust stress–inflammation state and a cytostatic program. Hallmarks for inflammatory response and TNFα–NF-κB signaling are strongly enriched alongside AP-1 components (JUNB, FOSL1) and ECM regulators (SERPINE1, TIMP1, PLAUR). A pronounced oxidative stress/antioxidant response is evident (ROS hallmark up) with increased SRXN1 and SLC7A11. Cell-cycle programs are broadly suppressed (E2F targets, G2/M checkpoint, mitotic spindle), and intermediate filament organization is strongly down, indicating cytoskeletal reorganization. Modest enrichment of EMT, TGF-β, and hypoxia hallmarks together with ECM regulators suggests stress-associated matrix remodeling, but not definitive EMT. A divergent xenobiotic response is observed: the xenobiotic metabolism hallmark is up while select P450s (CYP1A1, CYP3A7) and lipid/xenobiotic-related proteins (LIPC, KLB, HAMP) decrease, consistent with a shift toward antioxidant/transport-based detoxification; this is cell-context dependent and inferential. Notably, NF-κB appears activated under these conditions, which contrasts with the drug’s target annotation. No autophagy-related signatures are detected in this dataset.

## Findings consistent with expected biology
- Strong oxidative stress/antioxidant program activation (REACTIVE_OXYGEN_SPECIES_PATHWAY up) with increased SRXN1 and SLC7A11.
- Pro-inflammatory signaling with TNF–NF-κB/AP-1 features (INFLAMMATORY_RESPONSE and TNFA_SIGNALING_VIA_NFKB up; JUNB, FOSL1, SERPINE1, TIMP1, PLAUR elevated).
- Cytostatic effect with broad suppression of proliferation programs (E2F targets, G2/M checkpoint, mitotic spindle hallmarks down; mitosis/segregation GO terms down).
- Cytoskeletal reorganization marked by strong downregulation of intermediate filament organization.
- Stress-associated extracellular matrix remodeling signals (SERPINE1, TIMP1, PLAUR, PALLD up) with modest EMT hallmark enrichment.

## Unexpected but plausible findings
- NF-κB pathway appears activated (TNFA_SIGNALING_VIA_NFKB up with inflammatory mediators elevated) despite NF-κB being listed as a drug target, suggesting activation under these assay conditions.
- Divergent xenobiotic response: xenobiotic metabolism hallmark up while CYP1A1 and CYP3A7 decrease and antioxidant/transport systems (e.g., SLC7A11, SRXN1) increase, indicating a potential shift away from P450-centric metabolism.

## Follow-up hypotheses
### Hypothesis 1
- **Hypothesis:** Andrographolide induces a combined NF-κB/AP-1 inflammatory program and antioxidant response that contributes to cytostasis.
- **Why interesting:** Explains the coexistence of strong inflammatory signaling and proliferation arrest; clarifies directionality relative to the drug’s annotated NF-κB targeting.
- **Suggested validation:** Assess NF-κB and AP-1 activation (p65 nuclear translocation, IκBα degradation, NF-κB/AP-1 reporter assays), confirm target gene induction (e.g., IL6, CXCL8, SERPINE1), and measure cell-cycle arrest (EdU/BrdU incorporation, pRB phosphorylation, cyclin/CDK levels) across dose/time.
- **Priority:** high

### Hypothesis 2
- **Hypothesis:** An Nrf2-like antioxidant pathway is engaged downstream of oxidative stress.
- **Why interesting:** Elevated SRXN1 and SLC7A11 and ROS hallmark up suggest Nrf2 involvement, with therapeutic implications for redox buffering and ferroptosis sensitivity.
- **Suggested validation:** Measure Nrf2 stabilization and nuclear localization; quantify canonical Nrf2 targets (NQO1, HMOX1/HO-1, GCLM) by proteomics/qPCR; assess intracellular ROS and glutathione levels; test sensitivity to Nrf2 inhibition or SLC7A11 blockade.
- **Priority:** medium

### Hypothesis 3
- **Hypothesis:** Andrographolide triggers stress-associated ECM remodeling without full EMT.
- **Why interesting:** Upregulation of SERPINE1/TIMP1/PLAUR and modest EMT hallmark enrichment suggest matrix remodeling that may affect invasiveness and adhesion.
- **Suggested validation:** Profile canonical EMT markers (E-cadherin, keratins, vimentin, SNAI/ZEB), assess cell migration/invasion, uPA/uPAR activity, MMP activity, and matrix deposition; include time course to distinguish transient stress responses from stable state changes.
- **Priority:** medium

### Hypothesis 4
- **Hypothesis:** Xenobiotic handling shifts from P450-dependent metabolism toward antioxidant/transport-mediated detoxification in a cell-type–dependent manner.
- **Why interesting:** Could impact drug–drug interactions and metabolic liability, especially if liver-like functions are affected.
- **Suggested validation:** Determine cell model identity; measure CYP1A1/3A activity and expression, glutathione synthesis and export, and reliance on SLC7A11; include Reactome/KEGG pathway analyses and metabolite profiling to map detox routes.
- **Priority:** low

## Confidence
medium

## Caution notes
- Pathway activity inferences (NF-κB, Nrf2, TGF-β, hypoxia) are based on Hallmark enrichments and marker proteins; they are not direct measurements of pathway activation.
- Interpretations of upregulated programs rely on Hallmark signatures with no significant GO BP up or Reactome/KEGG enrichments provided.
- EMT-related conclusions should be considered tentative without consistent shifts in canonical epithelial/mesenchymal markers.
- The xenobiotic/P450 interpretation is context dependent and requires knowledge of the cell model; avoid generalizing to hepatic function without confirmation.
- Autophagy is listed as a target but no autophagy markers or pathways are evident here; no conclusions about autophagy should be drawn from this dataset.
- Some individual protein changes are near significance thresholds; dose and timepoint are unspecified, so responses may be transient.
- Proteomics is the only data layer shown; orthogonal assays (e.g., phosphoproteomics, transcriptomics, reporter assays) are recommended to confirm pathway directionality.


---

# Aniracetam

# Drug summary: Aniracetam

## Executive summary
Proteomic profiling of Aniracetam exposure shows a stress-adaptive signature characterized by robust enrichment of inflammatory/TNFA–NF-κB and hypoxia-like hallmarks, alongside strong, coordinated suppression of RNA splicing/mRNA processing and rRNA processing/ribosome biogenesis. Concurrent decreases in intermediate filament organization and specific junctional/structural proteins (e.g., KRT9, CLDN11/12) are observed. Selected proteins associated with membrane lipid metabolism/trafficking (CERK, DDHD1, GDPD1, TSPAN13, RNF167) and actin remodeling (WASF1) are increased, but no corresponding pathway enrichments were detected. The overall pattern indicates a shift away from growth/biogenesis toward stress response, which is unexpected for a neuromodulator and may reflect generalized cellular stress in this model rather than canonical iGluR/nAChR target engagement.

## Findings consistent with expected biology
- None listed.

## Unexpected but plausible findings
- Strong up-regulation of inflammatory and TNFA–NF-κB-associated programs together with a hypoxia-like response (Hallmark INFLAMMATORY_RESPONSE NES ~2.04, HYPOXIA NES ~1.98; TNFA_SIGNALING_VIA_NFKB NES ~1.75), with protein-level support (e.g., FOSL1, LOXL2, MSRA up).
- Marked suppression of RNA splicing/mRNA processing and rRNA processing/ribosome biogenesis (multiple GO BP terms with large negative NES), with decreases in RSL24D1, RPL36AL, and histones, consistent with reduced RNP biogenesis.
- Downregulation of intermediate filament organization (GO BP NES ~-2.26) with KRT9 decreased; tight junction components CLDN11 and CLDN12 reduced, suggesting junctional/structural program attenuation in this context.
- Protein-level signals suggestive of membrane/lipid remodeling and actin dynamics (CERK, DDHD1, GDPD1, TSPAN13, RNF167, WASF1 up) despite absence of corresponding enriched up pathways.
- PTEN down with FOSL1 up and Hallmark KRAS_SIGNALING_DN decreased, hinting at possible signaling rewiring (e.g., MAPK–AP-1 or PI3K/AKT involvement), but this remains tentative without phospho-pathway evidence.

## Follow-up hypotheses
### Hypothesis 1
- **Hypothesis:** Aniracetam induces NF-κB- and HIF-associated stress programs in this cell context.
- **Why interesting:** Explains the robust inflammatory and hypoxia Hallmark enrichments with supporting protein increases and could indicate off-target or context-dependent stress responses.
- **Suggested validation:** Assess NF-κB activation (p65 nuclear translocation, phospho-IKK, NF-κB target transcripts/proteins), HIF1α stabilization and HIF target readouts, cytokine secretion profiling, and time-course confirmation.
- **Priority:** high

### Hypothesis 2
- **Hypothesis:** Global downregulation of RNA splicing and ribosome biogenesis reflects nucleolar stress and growth arrest.
- **Why interesting:** Multiple strong GO BP decreases and reduced RNP/ri bosome-associated proteins suggest a shift away from biogenesis that could underlie functional outcomes.
- **Suggested validation:** Measure nascent RNA synthesis (5-EU incorporation), pre-rRNA processing (ITS probe assays), nucleolar integrity (fibrillarin/NPM1 staining), cell cycle/viability metrics, and dose–time dependence.
- **Priority:** high

### Hypothesis 3
- **Hypothesis:** Aniracetam perturbs cytoskeletal and junctional architecture in this model.
- **Why interesting:** Consistent decreases in intermediate filament organization and specific junctional/structural proteins indicate potential remodeling that could affect cell barrier properties.
- **Suggested validation:** Immunofluorescence for keratins and claudins, TEER or permeability assays (if epithelial), and cytoskeletal imaging (phalloidin) with morphology quantification.
- **Priority:** medium

### Hypothesis 4
- **Hypothesis:** Secondary activation of MAPK–AP-1 occurs under treatment-induced stress.
- **Why interesting:** FOSL1 up and KRAS_SIGNALING_DN hallmark changes suggest AP-1/MAPK involvement, which could coordinate stress-adaptive transcriptional programs.
- **Suggested validation:** Phospho-ERK/cJUN immunoblots, AP-1 reporter assays, and inhibitor challenge (e.g., MEK inhibitors) to test dependence; broaden AP-1 component profiling.
- **Priority:** medium

### Hypothesis 5
- **Hypothesis:** Membrane lipid metabolism and endolysosomal trafficking are increased as adaptive remodeling.
- **Why interesting:** Upregulation of CERK, DDHD1, GDPD1, TSPAN13, and RNF167 suggests altered lipid signaling and trafficking that may influence receptor and membrane dynamics.
- **Suggested validation:** Targeted lipidomics (ceramide/phosphatidic acid/lyso-PL), endolysosomal trafficking assays (e.g., transferrin or LDL uptake/recycling), surface proteome analysis, and live-cell imaging of endosomes.
- **Priority:** low

## Confidence
medium

## Caution notes
- Cell type/model and receptor expression are unspecified; lack of clear iGluR/nAChR target engagement markers may reflect context rather than drug action.
- Conclusions are based on steady-state proteomics; pathway activation (NF-κB, HIF, MAPK, PI3K/AKT) is not confirmed without phosphoproteomics or transcript-level data.
- Interpretation of KRAS_SIGNALING_DN as evidence of RAS/MAPK activation is context-dependent and not definitive.
- Inferences about membrane/lipid remodeling and actin dynamics derive from a small set of proteins without enriched up pathways and should be treated as hypothesis-generating.
- Decreases in claudins/keratin and claims of barrier weakening are cell-type dependent and require functional validation.
- Broad suppression of RNA processing and ribosome biogenesis could reflect generalized stress or reduced proliferation; relate findings to dose, exposure time, and viability.
- PTEN downregulation is a single-protein observation and does not establish pathway rewiring on its own.
- Some pathway sections are empty (e.g., GO BP up); absence of enrichment there does not negate protein-level observations but limits pathway-level interpretation.


---

# Antazoline (hydrochloride)

# Drug summary: Antazoline (hydrochloride)

## Executive summary
Proteomic profiling of Antazoline (hydrochloride) exposure reveals a stress–inflammation signature with a pronounced hypoxia-like program (HYPOXIA hallmark) and activation of pro-inflammatory modules (INFLAMMATORY_RESPONSE, TNF/NFkB, IL6/STAT3), alongside a shift toward protein secretion. Concordantly, several secreted and matrix-modifying proteins increase (e.g., TXNIP, LOXL2, CEMIP, CTHRC1, SPINT1, CLEC3B, TYRO3), while some structural ECM components decrease (e.g., COL6A1, EFEMP1), indicating mixed ECM remodeling. Anti-proliferative features are evident: E2F targets and G2/M checkpoint hallmarks are down, and GO processes for chromatin/chromosome functions, RNA splicing, and ribosome biogenesis are broadly suppressed. These changes are consistent with reduced biosynthetic and cell-cycle activity and a stress-adaptive, more secretory state. A decrease in selected mitochondrial/Complex I–related proteins (e.g., NDUFA1, NDUFAF6, DNAJC30) may indicate mitochondrial contribution to the stress phenotype, but this remains tentative without pathway-level enrichment. The interferon-alpha response is present but more modest than the TNF/NFkB and IL6/STAT3 signals. No significant GO biological processes were enriched on the upregulated side, and Reactome/KEGG summaries were not detected; conclusions are therefore driven by Hallmark and GO-down evidence plus protein-level changes.

## Findings consistent with expected biology
- Activation of inflammatory signaling programs, including TNF/NFkB and IL6/STAT3, consistent with the drug’s immunology annotation.
- Increased secretory tendency (Hallmark PROTEIN_SECRETION) with multiple secreted/matrix-associated proteins up.

## Unexpected but plausible findings
- A strong hypoxia-like program (HYPOXIA hallmark) in response to a histamine receptor–targeting agent.
- Broad suppression of proliferation-linked programs (E2F targets, G2/M) and chromatin/RNA processing/ribosome biogenesis pathways, consistent with reduced biosynthetic and cell-cycle activity.
- Mixed ECM remodeling: upregulation of matrix-modifying/secreted proteins (LOXL2, CEMIP, CTHRC1, SPINT1, CLEC3B, TYRO3) with concurrent decreases in some ECM components (COL6A1, EFEMP1).
- Modest interferon-alpha response alongside stronger TNF/NFkB-driven inflammation.
- Tentative evidence of reduced mitochondrial/Complex I function based on a limited protein set (e.g., NDUFA1, NDUFAF6, DNAJC30) without pathway-level enrichment.

## Follow-up hypotheses
### Hypothesis 1
- **Hypothesis:** Antazoline induces a hypoxia-like stress program independent of oxygen deprivation, contributing to inflammatory signaling and secretory shift.
- **Why interesting:** Explains the strong HYPOXIA hallmark alongside TNF/NFkB and IL6/STAT3 activation and increased secreted/ECM-modifying proteins.
- **Suggested validation:** Assess HIF pathway activation (HIF1A stabilization, CA9/VEGFA targets), hypoxia reporters, and OCR/ECAR measurements across dose/time; test whether HIF inhibitors attenuate the signature.
- **Priority:** high

### Hypothesis 2
- **Hypothesis:** NFkB/STAT3 activation drives a stress-adaptive secretory phenotype.
- **Why interesting:** Aligns with upregulated inflammatory hallmarks and the PROTEIN_SECRETION hallmark plus increased secreted proteins.
- **Suggested validation:** Measure NFkB p65 nuclear translocation and STAT3 Y705 phosphorylation; quantify IL6/TNF secretion; use NFkB/STAT3 inhibitors to test dependency of the secretory/ECM protein increases.
- **Priority:** high

### Hypothesis 3
- **Hypothesis:** Suppression of rRNA biogenesis and RNA splicing contributes to reduced proliferation.
- **Why interesting:** Strong GO downregulation (rRNA processing, ribosome biogenesis, splicing) co-occurs with E2F/G2M repression, suggesting a coordinated anti-proliferative state.
- **Suggested validation:** Monitor rRNA synthesis (5-EU/FUrd incorporation), nucleolar markers (fibrillarin), and cell-cycle profiles (EdU/PI); evaluate proliferation/viability metrics across dose/time.
- **Priority:** high

### Hypothesis 4
- **Hypothesis:** Mitochondrial Complex I impairment contributes to the stress phenotype and hypoxia-like signaling.
- **Why interesting:** Decreases in NDUFA1/NDUFAF6/DNAJC30 suggest mitochondrial involvement that could reinforce hypoxia-like and inflammatory responses.
- **Suggested validation:** Perform complex I–driven respirometry (OCR with pyruvate/malate), assess mitochondrial membrane potential and ROS, and test sensitivity to rotenone; include time-course and rescue experiments.
- **Priority:** medium

### Hypothesis 5
- **Hypothesis:** Antazoline promotes ECM remodeling characterized by increased matrix-modifying enzymes and altered ECM composition.
- **Why interesting:** Protein-level increases in LOXL2, CEMIP, CTHRC1, SPINT1, CLEC3B and decreases in COL6A1/EFEMP1 indicate a remodeling trajectory.
- **Suggested validation:** Secretome profiling, collagen crosslinking/activity assays (LOXL2), matrix deposition imaging, and targeted ECM pathway analyses over time.
- **Priority:** medium

## Confidence
medium

## Caution notes
- Pathway enrichments are derived from proteomic rankings using Hallmark/GO sets curated largely from transcriptomics; protein-level regulation and stoichiometry may differ.
- No dose, timepoint, or cell-type details are provided; hypoxia-like and inflammatory signatures could be context- or concentration-dependent or reflect generalized stress/off-target effects.
- Claims of reduced proliferation should be framed as ‘consistent with’ and confirmed by functional assays (cell cycle, proliferation/viability).
- Mitochondrial/Complex I impairment is based on a limited set of proteins and lacks pathway-level enrichment; treat as tentative pending functional validation.
- ECM remodeling interpretation is mixed (both increases and decreases) and should be corroborated with targeted ECM/secretome analyses.
- The absence of significant GO_BP_up enrichments indicates limited detectable coordinated upregulated GO processes, not an absence of upregulated biology overall.


---

# Antipyrine

# Drug summary: Antipyrine

## Executive summary
Proteomic profiling of Antipyrine exposure shows robust activation of pro‑inflammatory programs with TNF–NFkB features (Hallmark INFLAMMATORY_RESPONSE and TNFA_SIGNALING_VIA_NFKB up) and a hypoxia‑like stress signature (Hallmark HYPOXIA up). In parallel, intermediate filament/keratin organization is strongly suppressed (GO down), with decreases in keratins (e.g., KRT2, KRT79), indicating cytoskeletal remodeling. A striking single‑protein change is the marked downregulation of CYP1A1, suggesting a selective impact on this xenobiotic enzyme without pathway‑level corroboration. Additional increases in proteins related to cell communication and transcriptional control (e.g., LOXL2, EFNA1, CEMIP, several zinc‑finger factors) are observed but lack pathway‑level enrichment. Overall, the data support an inflammatory/hypoxic stress response with cytoskeletal reorganization and a notable CYP1A1 decrease; mechanistic attribution beyond these core findings remains tentative.

## Findings consistent with expected biology
- Pro‑inflammatory signaling activation with TNF–NFkB features (Hallmark INFLAMMATORY_RESPONSE NES ~1.97, FDR ~0.0067; TNFA_SIGNALING_VIA_NFKB NES ~1.78, FDR ~0.0157)
- Induction of a hypoxia‑like stress program (Hallmark HYPOXIA NES ~1.86, FDR ~0.009)

## Unexpected but plausible findings
- Strong suppression of intermediate filament/keratin organization (GO down: INTERMEDIATE_FILAMENT_BASED_PROCESS and INTERMEDIATE_FILAMENT_ORGANIZATION; decreased keratins KRT2 and KRT79)
- Marked repression of CYP1A1 (log2FC ~−2.32; adj P ~7.9e−09) without xenobiotic pathway‑level enrichment
- Decreased IL17RA protein (log2FC ~−1.17; adj P ~0.025) despite overall inflammatory Hallmark induction

## Follow-up hypotheses
### Hypothesis 1
- **Hypothesis:** Antipyrine‑induced inflammatory and hypoxia‑like responses are mediated by NFkB and HIF signaling, respectively.
- **Why interesting:** Validates the dominant Hallmark signals and clarifies the proximal transcriptional regulators driving the proteomic changes.
- **Suggested validation:** Assess NFkB activation (p65 nuclear translocation, NFkB reporter activity) and HIF pathway engagement (HIF1α stabilization, target gene induction such as CA9/VEGFA) with and without pharmacologic inhibitors; confirm cytokine/chemokine protein outputs (e.g., IL‑8).
- **Priority:** high

### Hypothesis 2
- **Hypothesis:** Antipyrine selectively dampens CYP1A1 through reduced activity of its upstream regulatory axis.
- **Why interesting:** A selective decrease in a key xenobiotic enzyme may have implications for metabolic capacity and potential drug–drug interactions.
- **Suggested validation:** Measure CYP1A1 mRNA and enzyme activity (e.g., EROD assay), evaluate upstream transcriptional activity using a responsive reporter, and test rescue by pathway agonists; survey additional enzymes in the same axis (e.g., CYP1B1) for coordinated changes.
- **Priority:** medium

### Hypothesis 3
- **Hypothesis:** Antipyrine exposure reduces keratin/intermediate filament network integrity in this model.
- **Why interesting:** Cytoskeletal remodeling can impact cell shape, mechanics, and barrier‑related functions, aligning with strong GO down enrichments.
- **Suggested validation:** Immunofluorescence imaging and immunoblotting for a broader keratin panel (e.g., KRT2, KRT79 and lineage‑appropriate keratins), quantification of filament organization, and time‑course analysis to define onset and reversibility.
- **Priority:** high

### Hypothesis 4
- **Hypothesis:** Inflammatory activation occurs with reduced IL‑17 signaling capacity due to decreased IL17RA.
- **Why interesting:** Indicates potential bias toward TNF/NFkB‑driven inflammation over IL‑17‑driven inputs, which may shape downstream effector profiles.
- **Suggested validation:** Measure cellular responses to IL‑17A versus TNF (target gene induction, signaling readouts), quantify IL17RA by flow cytometry/western, and test whether restoring IL17RA rescues IL‑17 responsiveness.
- **Priority:** medium

## Confidence
medium

## Caution notes
- Mechanistic inferences beyond core Hallmark signals should be considered provisional; single‑protein changes (e.g., CYP1A1) do not establish pathway‑level suppression.
- Avoid interpreting KRAS_SIGNALING_DN directionality without complementary evidence; its semantics are non‑intuitive.
- Do not infer epithelial identity changes from keratin decreases without cell‑type context; the finding supports intermediate filament downregulation but not lineage shifts.
- No significant GO BP up, Reactome, or KEGG enrichments were detected; upregulated biology is primarily captured by Hallmark sets in this dataset.
- Autophagy/MTOR or mitochondrial involvement is not supported at pathway level; isolated proteins (e.g., ATG12 up, NDUFA3 down) should not be over‑interpreted.
- Dose, timepoint, and cell‑type metadata are not provided; generalization of mechanism and physiological implications requires additional context and orthogonal validation.


---

# Asciminib

# Drug summary: Asciminib

## Executive summary
Asciminib exposure yields a coherent antiproliferative proteomic signature characterized by strong suppression of E2F/G2M programs and mitotic/chromosome-segregation machinery, accompanied by broad downregulation of RNA splicing/mRNA processing. In parallel, there is robust induction of xenobiotic/detox responses (notably CYP1A1) and marked upregulation of sterol/cholesterol/steroid biosynthesis and transport programs. Enrichment for inflammatory/IL6–JAK–STAT3 signatures is observed alongside increased amino/organic acid transporters. Additional enrichments (Hedgehog signaling, apical junction, coagulation/angiogenesis) are present but less substantiated at the individual-protein level. Overall, the data support growth arrest consistent with BCR-ABL pathway inhibition, coupled with metabolic and stress-adaptive remodeling; some pathway activations should be interpreted as signature enrichments rather than proven functional activation.

## Findings consistent with expected biology
- Antiproliferative program with strong downregulation of cell-cycle/mitotic machinery: Hallmark E2F_TARGETS and G2M_CHECKPOINT are suppressed, with decreases in BIRC5 (survivin), CENPM/CENPX, and SKA2.
- Broad reduction in RNA splicing and mRNA processing pathways consistent with growth arrest (multiple GO terms with strong negative NES).
- Robust xenobiotic/detox response with marked CYP1A1 induction and enrichment of Hallmark XENOBIOTIC_METABOLISM.

## Unexpected but plausible findings
- Marked induction of sterol/cholesterol/steroid biosynthesis and lipid/organic acid transport (Hallmark CHOLESTEROL_HOMEOSTASIS; multiple GO sterol/steroid terms), with protein-level support including CYP19A1 and monocarboxylate transporters SLC16A6/SLC16A13.
- Enrichment of inflammatory/IL6–JAK–STAT3 signaling and INFLAMMATORY_RESPONSE, with increases in IL4R and JUNB, suggesting cytokine-driven adaptation during arrest.
- Hedgehog signaling gene-set enrichment, indicating engagement of developmental signaling, though lacking corroborating effector proteins in this dataset.
- Downregulation of keratin/intermediate filament programs with modest enrichment of apical junction, consistent with cell-state/adhesion remodeling during arrest.
- Evidence consistent with a pro-survival shift: enrichment for positive regulation of PI3K–AKT signaling and decreases in DDIT4 and TXNIP; interpretation remains tentative without phospho-readouts.
- Potential Wnt-pathway tuning signaled by decreased DKK1 and increased ZNRF3; pathway-level enrichment is absent, so this remains a hypothesis-generating observation.

## Follow-up hypotheses
### Hypothesis 1
- **Hypothesis:** Asciminib triggers SREBP-driven activation of cholesterol/sterol biosynthesis as an adaptive survival program.
- **Why interesting:** Sterol/steroid biosynthetic processes are among the strongest enrichments (Hallmark and multiple GO terms), with supporting protein-level changes; targeting this axis may enhance antiproliferative efficacy.
- **Suggested validation:** Assess SREBP1/2 cleavage and nuclear localization; perform lipidomics and de novo cholesterol synthesis assays; test synergy with statins or squalene synthase inhibitors under Asciminib treatment.
- **Priority:** high

### Hypothesis 2
- **Hypothesis:** Cytokine/IL6–JAK–STAT3 signaling is engaged as a compensatory survival pathway during proliferation arrest.
- **Why interesting:** Hallmark IL6_JAK_STAT3 and INFLAMMATORY_RESPONSE are enriched, with IL4R and JUNB upregulated, suggesting actionable inflammatory signaling.
- **Suggested validation:** Measure STAT3 Tyr705 phosphorylation and nuclear translocation; profile secreted cytokines (e.g., IL6, IL8); evaluate combination effects with JAK inhibitors (e.g., ruxolitinib) on viability.
- **Priority:** medium

### Hypothesis 3
- **Hypothesis:** A PI3K–AKT pro-survival tone emerges under Asciminib, contributing to persistence despite growth arrest.
- **Why interesting:** GO enrichment for positive regulation of PI3K–AKT coupled with reduced DDIT4 and TXNIP suggests potential AKT relief from negative regulators.
- **Suggested validation:** Probe pAKT (Ser473/Thr308) and downstream effectors (pS6, pGSK3β); test sensitivity to PI3K/AKT/mTOR inhibitors in combination with Asciminib.
- **Priority:** medium

### Hypothesis 4
- **Hypothesis:** Hedgehog pathway contributes to cell-state remodeling after Asciminib treatment.
- **Why interesting:** Hallmark HEDGEHOG_SIGNALING is significantly enriched, indicating possible involvement of developmental signaling.
- **Suggested validation:** Quantify GLI1/GLI2 protein and transcriptional reporter activity; test SMO or GLI inhibitors for functional impact on cell-state markers and viability.
- **Priority:** low

## Confidence
high

## Caution notes
- Pathway ‘activation’ here reflects enrichment of protein abundance signatures; functional activation (e.g., phosphorylation, ligand levels) was not measured.
- Several secondary inferences (e.g., Hedgehog, PI3K–AKT compensation, Wnt tuning) rely on gene-set enrichment or few proteins and should be viewed as hypothesis-generating.
- Coagulation/angiogenesis and apical junction enrichments are near FDR thresholds and lack specific marker corroboration; avoid strong mechanistic conclusions.
- Induction of sterol/cholesterol programs may represent a general adaptive response to stress/drug exposure rather than a direct on-target effect; causality requires orthogonal validation.
- Wnt pathway status should not be inferred from DKK1 and ZNRF3 alone; broader pathway evidence is needed.
- Absence of Reactome/KEGG outputs does not negate the Hallmark/GO findings but limits pathway-level cross-validation.


---

# Atazanavir (sulfate)

# Drug summary: Atazanavir (sulfate)

## Executive summary
Across proteome and pathway analyses, Atazanavir (sulfate) induces a stress- and cytokine-driven state characterized by robust enrichment of IL6–JAK–STAT3, TNFA–NFkB, interferon-alpha, and inflammatory-response hallmarks, supported by increased IL4R, TNFRSF12A (Fn14), FOSL1, TXNIP, and IGFBP3. A hypoxia-like program is evident alongside mixed mitochondrial remodeling (TOMM40L/TOMM5 and PET100 up; CHCHD4 and NDUFV3 down), consistent with mitochondrial proteostasis/respiratory strain. Cells show strong pathway-level suppression of RNA splicing/mRNA processing and ribosome biogenesis (multiple GO sets down; RSL24D1 decreased). Protein secretion and lipid/cholesterol homeostasis hallmarks are also enriched with supporting increases in HID1, HPS1, GAL3ST1, and CYP4F22. An apoptosis hallmark is up, aligning with stress signaling. EMT is enriched at the pathway level, although per-protein evidence is limited. Overall, the signature supports inflammatory and hypoxia-like stress with coordinated downshift of RNA processing/biogenesis and adjustments in secretory/lipid programs; several mechanistic inferences are pathway-level and warrant targeted validation.

## Findings consistent with expected biology
- Activation of pro-inflammatory cytokine programs (IL6–JAK–STAT3, TNFA–NFkB, interferon-alpha), supported by increases in IL4R, TNFRSF12A, FOSL1, TXNIP, and IGFBP3
- Hypoxia-like/mitochondrial stress signature with mixed remodeling of mitochondrial import and respiratory components (TOMM40L/TOMM5 and PET100 up; CHCHD4 and NDUFV3 down)
- Strong pathway-level suppression of RNA splicing/mRNA processing and ribosome biogenesis (multiple GO BP sets down; RSL24D1 decreased)
- Enrichment of protein secretion and lipid/cholesterol homeostasis pathways with supporting increases in HID1, HPS1, GAL3ST1, and CYP4F22
- Apoptosis hallmark up, consistent with a cellular stress response

## Unexpected but plausible findings
- Epithelial–mesenchymal transition (EMT) pathway enrichment (NES ~1.83) with limited canonical per-protein support (e.g., LOXL2, FOSL1 up)
- Concomitant decreases in PTEN and PHLPP2 despite stress/apoptosis and inflammatory program activation; no direct PI3K/AKT activation evidence at the pathway level
- KRAS_SIGNALING_UP enrichment without direct KRAS protein changes, likely reflecting a secondary stress-associated signature
- Mixed mitochondrial directionality (import and assembly factors up with other mitochondrial components down) rather than uniform mitochondrial up- or downregulation

## Follow-up hypotheses
### Hypothesis 1
- **Hypothesis:** Atazanavir induces STAT3 and NFkB pathway activation secondary to cellular stress.
- **Why interesting:** Proteomic abundance patterns show strong enrichment of IL6–JAK–STAT3, TNFA–NFkB, and interferon programs with concordant receptor/transcription factor changes.
- **Suggested validation:** Measure phospho-STAT3 (Y705) and NFkB p65 nuclear translocation/IKBα degradation; quantify cytokine secretion (e.g., IL6, TNFα) by ELISA.
- **Priority:** high

### Hypothesis 2
- **Hypothesis:** Atazanavir perturbs mitochondrial proteostasis/respiration, leading to hypoxia-like signaling.
- **Why interesting:** HYPOXIA hallmark up with TOMM40L/TOMM5 and PET100 increases and CHCHD4/NDUFV3 decreases suggests import/respiratory strain.
- **Suggested validation:** Assess mitochondrial respiration (Seahorse OCR), HIF stabilization/target gene induction, and mitochondrial protein import assays; evaluate CHCHD4-dependent oxidative folding.
- **Priority:** high

### Hypothesis 3
- **Hypothesis:** Global downregulation of RNA splicing and ribosome biogenesis reflects a stress-adaptive reduction in biosynthetic capacity.
- **Why interesting:** Multiple splicing and ribosome biogenesis gene sets are strongly downregulated.
- **Suggested validation:** Measure splicing efficiency using splicing reporters or intron retention assays; assess pre-rRNA processing and nascent translation rates (e.g., SUnSET).
- **Priority:** medium

### Hypothesis 4
- **Hypothesis:** EMT-like reprogramming and increased secretory/lipid metabolism are secondary adaptations to inflammatory/hypoxic stress rather than direct EMT induction.
- **Why interesting:** EMT and protein secretion/cholesterol homeostasis hallmarks are enriched, but canonical EMT protein shifts are limited.
- **Suggested validation:** Profile canonical EMT markers (E-cadherin, Vimentin, ZEB1) and cell motility; perform lipidomics and secretion assays to confirm pathway activation.
- **Priority:** medium

### Hypothesis 5
- **Hypothesis:** Partial relief of AKT negative regulation occurs (PTEN/PHLPP2 down) without full PI3K/AKT pathway activation.
- **Why interesting:** PTEN and PHLPP2 are decreased despite apoptotic/inflammatory signaling; pathway-level PI3K evidence is absent.
- **Suggested validation:** Measure AKT phosphorylation (T308/S473) and downstream targets (e.g., PRAS40, GSK3β) to determine functional impact.
- **Priority:** low

## Confidence
medium

## Caution notes
- Pathway enrichments are inferred from protein abundance and do not directly demonstrate signaling activation; phospho-based assays (e.g., STAT3, NFkB) are needed.
- Mitochondrial oxidative folding/respiration impairment is suggested by a small set of proteins (e.g., CHCHD4 down) and requires functional validation.
- EMT inference is primarily pathway-level; classical EMT marker shifts are not prominent among the most changed proteins.
- Cholesterol homeostasis interpretation should be cautious; canonical regulators (e.g., HMGCR, LDLR, SREBF2) are not highlighted among top changes.
- Suppression of splicing and ribosome biogenesis is strong at the pathway level but needs validation with RNA processing and translation rate assays.
- Cell type, dose, and time are not specified; the observed inflammatory/hypoxia-like signatures may be context-dependent stress responses rather than primary on-target effects.
- Do not infer PI3K/AKT activation from PTEN/PHLPP2 decreases alone; pathway-level PI3K/AKT activation signatures are not observed.


---

# Atorvastatin (hemicalcium salt)

# Drug summary: Atorvastatin (hemicalcium salt)

## Executive summary
Proteomics after atorvastatin exposure shows a canonical, on-target mevalonate/cholesterol feedback response. Cholesterol/sterol and isoprenoid biosynthetic programs are strongly enriched, with increased HMGCR, SC5D, and ACSS2, alongside broader lipid handling (fatty acid, lipoprotein, bile acid metabolism) and xenobiotic/detoxification signatures. A pronounced metal-ion response is evident, driven by marked metallothionein induction (MT1B, MT1H) and enrichment for zinc/copper/cadmium response terms, consistent with a secondary metal-handling/oxidative stress adaptation. Complement and coagulation Hallmark sets are enriched with an increase in C9, but broader protein-level support is limited; some immune receptors (TLR1, TAPBPL) are decreased, indicating selective immune protein changes rather than definitive pathway activation. Ribosome biogenesis, rRNA processing, and RNA splicing pathways are coherently downregulated, suggesting reduced biosynthetic capacity, though functional translation was not directly measured. Decreases in COQ4 and TOMM40L point to possible mitochondrial involvement, but no mitochondrial pathway enrichment is observed, so this remains tentative. The absence of Hallmark_down sets reflects input structure; suppression is captured by GO_BP_down terms.

## Findings consistent with expected biology
- Robust compensatory engagement of the mevalonate/cholesterol axis: strong enrichment of cholesterol/sterol and isoprenoid biosynthesis with increased HMGCR and sterol enzymes (e.g., SC5D, ACSS2).
- Broader lipid metabolism rewiring consistent with mevalonate pathway perturbation: enrichment of fatty acid, lipoprotein, and bile acid metabolism with supporting proteins (e.g., APOA5, ANGPTL8).
- Induction of xenobiotic/detoxification programs accompanying drug exposure.

## Unexpected but plausible findings
- Strong metallothionein/metal-ion response (MT1B, MT1H up) with enrichment for zinc/copper/cadmium response and intracellular zinc homeostasis.
- Enrichment of complement and coagulation Hallmarks with increased C9, occurring alongside reduced abundance of select immune receptors (TLR1, TAPBPL); interpret as pathway-level enrichment rather than confirmed functional activation.
- Coherent suppression of ribosome biogenesis, rRNA processing, and RNA splicing pathways, suggesting reduced biosynthetic capacity without direct evidence of decreased translation.
- Potential mitochondrial involvement indicated by decreased COQ4 and TOMM40L, though not supported by pathway-level enrichment (tentative).

## Follow-up hypotheses
### Hypothesis 1
- **Hypothesis:** Atorvastatin triggers SREBP-driven compensatory upregulation of cholesterol/sterol and isoprenoid biosynthesis.
- **Why interesting:** Explains the coordinated increase of HMGCR and sterol pathway proteins and the strong enrichment of sterol/mevalonate programs.
- **Suggested validation:** Assess SREBP2 activation (cleavage and nuclear translocation) and downstream transcript levels; test mevalonate add-back rescue of sterol biosynthetic protein induction.
- **Priority:** high

### Hypothesis 2
- **Hypothesis:** Atorvastatin induces a metal-handling/oxidative stress adaptation reflected by metallothionein upregulation.
- **Why interesting:** Accounts for large MT1B/MT1H increases and metal-ion response enrichments; may influence redox balance and metal homeostasis.
- **Suggested validation:** Measure intracellular Zn/Cu levels and ROS; confirm metallothionein induction by orthogonal assays; evaluate antioxidant response markers.
- **Priority:** high

### Hypothesis 3
- **Hypothesis:** Complement/coagulation pathway components are modulated at the secreted protein level despite limited intracellular protein changes.
- **Why interesting:** Hallmark enrichment with increased C9 suggests pathway involvement that may be more evident in the secretome.
- **Suggested validation:** Secretome proteomics or ELISA for complement components and activation products (e.g., C3/C5 fragments, C9); coagulation factor assays as appropriate to cell model.
- **Priority:** medium

### Hypothesis 4
- **Hypothesis:** Suppression of ribosome biogenesis and RNA processing reduces global translational output under atorvastatin treatment.
- **Why interesting:** Pathway-level downregulation indicates potential dampening of protein synthesis as an adaptive response to metabolic stress.
- **Suggested validation:** Quantify protein synthesis by puromycin incorporation (SUnSET) or polysome profiling; survey ribosomal protein abundances; monitor rRNA maturation markers.
- **Priority:** medium

### Hypothesis 5
- **Hypothesis:** Mevalonate-pathway perturbation secondarily impacts mitochondrial function via reduced coenzyme Q biosynthesis.
- **Why interesting:** Decreased COQ4 and TOMM40L hint at mitochondrial consequences that could affect respiration and redox status.
- **Suggested validation:** Measure CoQ levels and mitochondrial respiration (OCR), membrane potential, and ROS; test rescue with mevalonate pathway intermediates.
- **Priority:** low

## Confidence
medium

## Caution notes
- Complement/coagulation findings are based on Hallmark enrichment with limited protein-level support (C9 increase); treat as enrichment rather than functional activation and validate in the secretome.
- mTORC1 signaling Hallmark enrichment does not establish pathway activity; phospho-markers (e.g., S6K, 4EBP1) are required for inference.
- Downregulation of ribosome biogenesis/splicing does not by itself prove reduced translation; functional assays are needed.
- Metallothionein induction supports altered metal homeostasis but the cause (oxidative stress versus metal flux) is not established.
- Mitochondrial effects are tentative, based on few proteins (COQ4, TOMM40L) without pathway enrichment; interpret cautiously.
- Cell type/context is unspecified; lipid, complement/coagulation, and bile acid-related enrichments can be tissue-specific.
- Empty Hallmark_down reflects the input set; downregulation is captured in GO_BP_down and should not be considered absent.
- Proteomic coverage and intracellular focus may underrepresent secreted factors and low-abundance regulators.


---

# ATP (dimagnesium)

# Drug summary: ATP (dimagnesium)

## Executive summary
Proteomic profiling after ATP (dimagnesium) exposure shows a robust hypoxia-like stress program with hallmark HYPOXIA enrichment and concordant up-regulation of stress/HIF-associated proteins (e.g., DDIT4, SERPINE1, LOXL2, FSTL3, JUNB). In parallel, multiple biosynthetic growth programs are suppressed (negative enrichment for ribosome biogenesis/rRNA processing, ribonucleoprotein complex biogenesis, and regulation of DNA replication), indicating reduced anabolic/proliferative activity. This pattern is consistent with attenuation of mTORC1-dependent translation, though direct mTOR readouts are not available. Immune signals are divergent: hallmark COMPLEMENT is enriched, whereas GO terms linked to immunoglobulin production and B‑cell–mediated immunity are down, suggesting an innate-skewed, context-dependent immune footprint. Additional increases in a caveolar component (CAVIN2) and several mitochondrial import/assembly factors (TOMM5, PET100, BOLA1), together with transport regulators (SLC16A6, FXYD1), point to structural and metabolic adaptation under stress; these latter features are preliminary due to limited marker breadth.

## Findings consistent with expected biology
- Hypoxia-like stress program activation: Hallmark HYPOXIA enriched (NES ~1.96, FDR ~0.0038) with concordant up-regulation of DDIT4, SERPINE1, LOXL2, FSTL3, and JUNB.
- Suppression of biosynthetic growth programs: significant negative enrichment for ribosome biogenesis/rRNA processing and regulation of DNA replication (NES ~−1.8 to −1.9, FDR <0.05).
- Innate immune cue up-regulation: Hallmark COMPLEMENT enriched (NES ~1.78, FDR ~0.033).
- Changes consistent with attenuated anabolic translation/mTORC1 activity: DDIT4 up alongside broad down-regulation of ribosome-related programs (inference only).

## Unexpected but plausible findings
- Innate–adaptive divergence: complement-associated signaling up while immunoglobulin production and B‑cell–mediated immunity programs are down (GO), indicating a decoupled immune signature dependent on cellular context.
- Caveolar/membrane architecture signal: marked up-regulation of CAVIN2 suggests membrane remodeling under stress (preliminary; lacks broader caveolae markers).
- Mitochondrial adaptation: increased TOMM5, PET100, and BOLA1 imply adjustment of mitochondrial import/respiratory chain assembly (preliminary; no pathway-level enrichment).
- Single-protein decreases in NR3C1 (glucocorticoid receptor) and RELB raise the possibility of reduced GR/NF‑κB-associated signaling, but this remains tentative without network-level support.

## Follow-up hypotheses
### Hypothesis 1
- **Hypothesis:** ATP (dimagnesium) induces a HIF–DDIT4–linked stress response that attenuates mTORC1-dependent anabolic programs, reducing ribosome biogenesis and DNA replication.
- **Why interesting:** Connects energy/stress signaling with broad growth repression and may explain coordinated proteome shifts across metabolism and immunity.
- **Suggested validation:** Assess phospho-S6 and phospho-4EBP1, global translation (puromycin incorporation), HIF1α stabilization; modulate DDIT4 (KD/OE) to test rescue of ribosome biogenesis markers.
- **Priority:** high

### Hypothesis 2
- **Hypothesis:** Treatment biases immune signaling toward innate/complement pathways while dampening humoral/B‑cell programs.
- **Why interesting:** Reveals selective immunomodulation that could affect inflammatory tone and therapeutic responses in relevant cell contexts.
- **Suggested validation:** Measure complement components (e.g., C3/C4/CFB) by proteomics/ELISA and functional hemolytic assays; verify B‑cell program markers where applicable; transcriptomic profiling to corroborate pathway shifts.
- **Priority:** medium

### Hypothesis 3
- **Hypothesis:** Membrane/caveolar remodeling occurs under stress, driven by increased CAVIN2, potentially altering endocytosis and mechanotransduction.
- **Why interesting:** Membrane architecture changes can influence receptor signaling, nutrient uptake, and cellular stress resilience.
- **Suggested validation:** Quantify CAV1/CAV2 and additional cavin proteins; visualize caveolae by EM; membrane fractionation and cholesterol perturbation to test caveolae dependence.
- **Priority:** low

### Hypothesis 4
- **Hypothesis:** Mitochondrial import and respiratory chain assembly are adaptively upregulated to meet altered energy demands.
- **Why interesting:** Mitochondrial remodeling may underpin survival under stress and shape downstream signaling.
- **Suggested validation:** Measure mitochondrial respiration (OCR), membrane potential, and complex assembly (BN-PAGE); assess TOM import efficiency and broader mitochondrial proteome changes.
- **Priority:** medium

## Confidence
medium

## Caution notes
- mTORC1/translation control inference is indirect (no phospho-S6/4EBP1/eIF2α data); interpret as ‘consistent with’ rather than definitive.
- Hallmark HYPOXIA supports a hypoxia-like program, but HIF activation/stabilization is not directly demonstrated.
- Immune GO terms for immunoglobulin/B‑cell processes may not reflect functional humoral changes unless relevant cell types are present; treat as motif-level enrichment.
- Membrane/caveolae remodeling rests primarily on CAVIN2; absence of broader caveolar markers or pathway enrichment weakens this claim.
- Mitochondrial remodeling is suggested by a small set of proteins without pathway-level enrichment; consider it preliminary.
- Single-protein decreases in NR3C1 and RELB should not be overinterpreted as GR/NF‑κB pathway suppression without corroborating evidence.
- Some GO down enrichments have modest FDR values (~0.03–0.046); replication or orthogonal assays will increase confidence.
- Empty Hallmark_down and GO_bp_up sections do not imply lack of effects; conclusions should be grounded in observed significant enrichments and protein changes.


---

# Azatadine (dimaleate)

# Drug summary: Azatadine (dimaleate)

## Executive summary
Proteomics following Azatadine (dimaleate) exposure shows a dominant activation of inflammatory/cytokine programs, with strong enrichment of Hallmark INFLAMMATORY_RESPONSE and IL6–JAK–STAT3 signaling, supported by increased IGFBP1 and the immune marker CD68. Integrin-mediated cell adhesion is also enriched, consistent with adhesion/cytoskeletal remodeling (with modest Hallmark MYOGENESIS). A hypoxia hallmark is enriched and coincides with limited mitochondrial protein changes (COQ4, TOMM5 up; ATP5IF1 down), suggesting a hypoxia-like or metabolic stress response, but metabolic pathway-level enrichment is not observed. No significantly enriched downregulated pathways were detected; decreases in selected innate/acute-phase or UPR-related proteins (e.g., ITIH4, TRIM22, SSC4D, ERN1) outline tentative down themes that require caution. Overall, the most coherent and expected signal is IL6–STAT3–driven inflammation with accompanying adhesion remodeling; hypoxia/metabolic and UPR features remain provisional.

## Findings consistent with expected biology
- Robust inflammatory and IL6–JAK–STAT3 signature enrichment (Hallmark INFLAMMATORY_RESPONSE NES ~2.16; IL6_JAK_STAT3_SIGNALING NES ~2.06; FDR ~0), supported by upregulated IGFBP1 and CD68.
- Upregulation of integrin-mediated cell adhesion (GO BP NES ~2.01) with modest cytoskeletal/myogenesis signal (Hallmark MYOGENESIS NES ~1.64), consistent with adhesion/morphology remodeling.

## Unexpected but plausible findings
- Enrichment of a hypoxia hallmark (NES ~1.84) with limited mitochondrial protein shifts (COQ4 and TOMM5 up; ATP5IF1 down), suggesting hypoxia-like/metabolic stress rather than confirmed hypoxia.
- Mixed immune profile: strong IL6–STAT3 activation coinciding with decreases in selected innate/acute-phase or immune-related proteins (e.g., ITIH4, TRIM22, SSC4D).
- Downregulation of the UPR sensor ERN1 (IRE1) without corroborating pathway-level UPR suppression; interpretation is uncertain.
- Upregulation of signaling/transcriptional regulators (e.g., RIC8B, TYRO3, ZKSCAN3, TFAP2C) suggests broader signaling changes, but no pathway enrichment supports a defined reprogramming.

## Follow-up hypotheses
### Hypothesis 1
- **Hypothesis:** Azatadine activates the IL6–JAK–STAT3 axis, driving an acute inflammatory response.
- **Why interesting:** Strong, coherent hallmark enrichment and IGFBP1 upregulation indicate functional STAT3 activity; validating this would clarify the drug’s immunomodulatory impact in this context.
- **Suggested validation:** Measure STAT3 Tyr705 phosphorylation and nuclear translocation; quantify STAT3 target transcripts and IL6 secretion; test reversal with JAK inhibitors or IL6 neutralization across relevant cell models.
- **Priority:** high

### Hypothesis 2
- **Hypothesis:** Azatadine enhances integrin-dependent adhesion and focal adhesion signaling, leading to cytoskeletal remodeling.
- **Why interesting:** GO enrichment for integrin-mediated adhesion with cytoskeletal signature suggests functional changes in cell–matrix interactions.
- **Suggested validation:** Perform adhesion and cell spreading assays; assess focal adhesion markers (FAK/SRC phosphorylation, vinculin/paxillin imaging); probe dependence on integrins with blocking antibodies.
- **Priority:** medium

### Hypothesis 3
- **Hypothesis:** The hypoxia-like signature is secondary to inflammatory/cytokine stress rather than true hypoxia.
- **Why interesting:** Hypoxia hallmark co-occurs with strong IL6–STAT3 enrichment and limited mitochondrial protein changes, pointing to stress signaling cross-talk.
- **Suggested validation:** Assess HIF1α stabilization and HRE-reporter activity; measure VEGFA/CA9 transcripts; profile oxygen consumption and glycolysis (Seahorse); test whether IL6–JAK pathway inhibition attenuates the hypoxia signature.
- **Priority:** medium

### Hypothesis 4
- **Hypothesis:** Azatadine modulates mitochondrial bioenergetics via ATP5IF1 downregulation and selective mitochondrial protein increases.
- **Why interesting:** ATP5IF1 decrease with COQ4/TOMM5 increases hints at OXPHOS regulation changes, but current support is limited.
- **Suggested validation:** Measure ATP synthase activity, respiratory control ratio, ATP/ADP levels, and mitochondrial membrane potential; confirm broader mitochondrial protein changes; perturb ATP5IF1 to test causality.
- **Priority:** low

### Hypothesis 5
- **Hypothesis:** Azatadine attenuates the IRE1 arm of the UPR under these conditions.
- **Why interesting:** ERN1 (IRE1) downregulation suggests possible UPR modulation, though evidence is single-protein based.
- **Suggested validation:** Evaluate XBP1 mRNA splicing and IRE1 phosphorylation; monitor BiP/CHOP levels and UPR reporter activity; benchmark against tunicamycin-induced UPR.
- **Priority:** low

## Confidence
medium

## Caution notes
- No significantly enriched downregulated pathways were detected; themes inferred from a few downregulated proteins should be interpreted cautiously.
- Hallmark and GO gene sets are transcriptome-derived; proteomic enrichment reflects signature-level concordance rather than direct pathway activation.
- Mechanistic assertions based on single or few proteins (e.g., ERN1 for UPR, ATP5IF1 for OXPHOS) require orthogonal validation.
- Hypoxia hallmark enrichment may reflect inflammatory/metabolic stress, not necessarily true hypoxic conditions; dedicated hypoxia assays are needed.
- Cell type/context and histamine receptor expression are not specified; context-dependent or off-target effects may contribute to the observed signatures.
- Inference of IL6–JAK–STAT3 activity beyond enrichment (e.g., STAT3 phosphorylation) is not established without phosphoproteomic or functional assays.


---

# Azelastine (hydrochloride)

# Drug summary: Azelastine (hydrochloride)

## Executive summary
Azelastine (HCl) produced a coherent proteomic response marked by strong induction of cholesterol/sterol metabolism and transport, enhanced receptor catabolism and endocytosis/endo-lysosomal trafficking, and broad suppression of ribosome biogenesis, RNA processing, and both cytoplasmic and mitochondrial translation. Hallmark E2F targets and G2/M checkpoint programs were decreased, consistent with reduced proliferative drive, and intermediate filament–related processes were down, indicating cytoskeletal remodeling. Inflammatory/cytokine-associated hallmarks (IL6–JAK–STAT3, IL2–STAT5, complement/coagulation) were enriched at the pathway level. Notable single-protein changes included marked FURIN upregulation and increases in proteins annotated to neuronal/amyloid processing (TMEM59L, TMEM59, ITM2C, SV2A), concordant with a gene-set enrichment for amyloid beta clearance. An mTORC1 hallmark increase co-occurred with translational repression, suggesting a stress-associated/noncanonical mTORC1 signature rather than increased protein synthesis. Overall, the data support membrane lipid remodeling and heightened receptor trafficking with a coordinated translational and proliferative brake; mechanistic attributions to specific signaling axes should be considered provisional without phospho-functional validation.

## Findings consistent with expected biology
- Modulation of inflammatory/cytokine programs (IL6–JAK–STAT3, IL2–STAT5, complement/coagulation hallmarks up)
- Enhanced receptor turnover and endo-lysosomal trafficking (receptor catabolic process and positive regulation of endocytosis up)
- Suppression of proliferative programs (E2F targets and G2/M checkpoint down)

## Unexpected but plausible findings
- Robust induction of cholesterol/sterol metabolic and transport pathways (e.g., SC5D and NPC1L1 up)
- Broad repression of ribosome biogenesis, rRNA processing, RNA splicing, and global translation
- Cytoskeletal remodeling with decreased intermediate filament–related processes
- Marked FURIN upregulation in the context of increased receptor trafficking
- Amyloid beta clearance gene-set enrichment with increased TMEM59L/TMEM59/ITM2C/SV2A
- mTORC1 hallmark up despite global translation-related programs down (consistent with a stress-associated signature)

## Follow-up hypotheses
### Hypothesis 1
- **Hypothesis:** Azelastine engages an SREBP-driven cholesterol homeostasis program that increases cholesterol biosynthesis and uptake.
- **Why interesting:** Membrane lipid remodeling can alter receptor trafficking, signaling competence, and endo-lysosomal function and may contribute to the observed receptor catabolism and inflammatory signatures.
- **Suggested validation:** Assess SREBP1/2 activation (nuclear SREBP fragments), cellular/free cholesterol (filipin staining), LDL uptake, and expression of canonical SREBP targets; test dependence using SREBP pathway inhibitors.
- **Priority:** high

### Hypothesis 2
- **Hypothesis:** Azelastine increases endo-lysosomal flux leading to enhanced receptor internalization and degradation, reducing cell-surface receptor abundance.
- **Why interesting:** This could mechanistically link GPCR/inflammatory pathway modulation to reduced signaling and explain receptor catabolic and amyloid/vesicular signatures.
- **Suggested validation:** Surface proteomics or flow cytometry for representative receptors (e.g., transferrin receptor and GPCRs of interest), pulse–chase with lysosomal inhibition, lysosomal activity assays, and interrogation of NDFIP1-dependent ubiquitination.
- **Priority:** high

### Hypothesis 3
- **Hypothesis:** A stress-associated translational brake (e.g., integrated stress response) operates alongside a noncanonical mTORC1-like signature.
- **Why interesting:** It explains the coexistence of an mTORC1 hallmark with broad repression of translation and cell-cycle programs and may underlie growth restraint.
- **Suggested validation:** Measure phospho-eIF2α and ATF4, global translation by puromycin incorporation (SUnSET), and mTORC1 readouts (phospho-S6K, phospho-4E-BP1); probe ISR dependence using ISRIB.
- **Priority:** medium

### Hypothesis 4
- **Hypothesis:** FURIN upregulation increases proprotein processing capacity in the secretory pathway, impacting maturation of multiple receptors/ligands.
- **Why interesting:** Augmented processing could reshape signaling and trafficking networks, reinforcing receptor turnover and pathway remodeling.
- **Suggested validation:** Quantify FURIN enzymatic activity, assess cleavage status of canonical substrates (e.g., pro-TGFβ family members, Notch precursors), and test sensitivity to FURIN inhibition.
- **Priority:** medium

## Confidence
high

## Caution notes
- Cell model/context is not specified; neuronal/amyloid and developmental annotations may reflect gene set overlap rather than lineage programs.
- Cytokine/STAT and mTORC1 statements are based on pathway enrichments; phosphorylation-based confirmation is needed to claim pathway activation.
- FURIN upregulation alone does not demonstrate increased substrate processing; functional assays are required.
- Amyloid beta clearance enrichment may reflect generalized vesicle/lysosomal trafficking and should not be over-attributed to amyloid biology without APP/Aβ measurements.
- Direct engagement of histamine receptors is not evidenced in these proteomic data; links to known GPCR MoA are indirect.
- Mechanistic interpretation relies primarily on Hallmark/GO enrichments; absence of KEGG/Reactome enrichments warrants conservative wording.
- Avoid overinterpretation of specific GO terms (e.g., syncytium-related terms) without corroborating protein-level evidence.


---

# Baicalein

# Drug summary: Baicalein

## Executive summary
Proteomics profiling of Baicalein shows a coordinated metabolic rewiring toward lipid catabolism and oxidative metabolism, with strong enrichment of fatty acid β-oxidation, peroxisomal pathways, oxidative phosphorylation, and bile acid/xenobiotic processing. Concordantly, several liver-associated/secretory proteins (e.g., APOA5, ITIH3, INHBE, SLCO2B1, NUDT7) are increased. In parallel, there is robust suppression of proliferation and biosynthetic programs (G2/M checkpoint, E2F/MYC targets, rRNA processing, ribosome biogenesis, cytoplasmic translation), consistent with cytostatic effects and reduced anabolic capacity. Pro-inflammatory and EMT/invasive signaling modules (TNFα/NF-κB, TGF-β, WNT/β-catenin) are attenuated with decreases in AXL, TYRO3, PLAUR, LOXL2, CCR6/CCL20 and related markers. Upregulation of DDIT4 and TP53INP2 is consistent with a shift toward catabolism and possible autophagy, but mTOR/autophagy status is not directly demonstrated. KRAS hallmark suppression likely reflects global cell-cycle arrest rather than direct KRAS pathway inhibition. No explicit ferroptosis signature is evident. Overall, Baicalein elicits cytostatic and anti-inflammatory effects coupled to enhanced lipid oxidative metabolism; hepatic-like metabolic/secretory features are observed but should be interpreted as adaptive reprogramming pending further validation.

## Findings consistent with expected biology
- Suppression of proliferation and biosynthetic machinery (down of G2M/E2F programs, MYC targets, rRNA processing, ribosome biogenesis, and translation), consistent with cytostatic effects
- Anti-inflammatory attenuation (Hallmark INFLAMMATORY_RESPONSE and TNFA_SIGNALING_VIA_NFKB down) with concordant decreases in CCR6 and CCL20
- Induction of xenobiotic/biotransformation pathways (Hallmark XENOBIOTIC_METABOLISM up), aligning with a metabolic enzyme modulation profile

## Unexpected but plausible findings
- Robust activation of fatty acid β-oxidation, peroxisome and oxidative phosphorylation, plus bile acid metabolism, with increased liver-associated proteins (APOA5, ITIH3, INHBE, SLCO2B1, NUDT7), suggesting a hepatic-like metabolic program
- Attenuation of EMT/invasive signaling (down of TGF-β, WNT/β-catenin hallmarks; decreases in AXL, TYRO3, PLAUR, LOXL2, TNS4, PROM1)
- Suppression of KRAS hallmark (KRAS_SIGNALING_UP down with KRAS_SIGNALING_DN up) despite no KRAS-directed mechanism in the package
- Increase in coagulation/secretory features (Hallmark COAGULATION up; APOA5, ITIH3, INHBE elevated)

## Follow-up hypotheses
### Hypothesis 1
- **Hypothesis:** Baicalein shifts cellular energetics toward fatty acid β-oxidation and peroxisomal/mitochondrial oxidation, supporting growth arrest.
- **Why interesting:** Strong enrichment for FAO/peroxisome/OXPHOS pathways with upregulation of NUDT7, SLC25A42, TK2 and related metabolic proteins.
- **Suggested validation:** Measure fatty acid oxidation rates (e.g., palmitate oxidation), Seahorse OCR/ECAR including FAO dependency, peroxisomal β-oxidation assays, acyl-carnitine profiling; test sensitivity to etomoxir or peroxisomal inhibitors.
- **Priority:** high

### Hypothesis 2
- **Hypothesis:** Baicalein dampens TNFα/NF-κB, TGF-β, and WNT signaling, reducing EMT and invasive behavior.
- **Why interesting:** Downregulation of corresponding Hallmark programs and decreased levels of invasion-associated proteins (AXL, LOXL2, PLAUR, CCR6/CCL20).
- **Suggested validation:** Assess p-p65/IκBα, p-SMAD2/3, and β-catenin reporter activity; quantify EMT markers (E-cadherin, vimentin); perform migration/invasion assays and cytokine/chemokine profiling.
- **Priority:** high

### Hypothesis 3
- **Hypothesis:** Baicalein restrains mTORC1-driven anabolism and promotes autophagy.
- **Why interesting:** Upregulation of DDIT4 (REDD1) and TP53INP2 with reduced translation and ribosome biogenesis suggests mTORC1 suppression and autophagy bias.
- **Suggested validation:** Immunoblot p-S6K and p-4EBP1; puromycin incorporation for protein synthesis; LC3-II flux with bafilomycin and p62 turnover to confirm autophagic flux.
- **Priority:** medium

### Hypothesis 4
- **Hypothesis:** The hepatic-like metabolic/secretory program reflects activation of HNF-driven transcriptional networks.
- **Why interesting:** Increases in HNF1B and liver-associated secretory proteins (APOA5, ITIH3, INHBE) accompany bile acid/coagulation hallmark enrichment.
- **Suggested validation:** Quantify HNF1B/HNF4A/CEBPA levels and activity (reporters/ChIP-qPCR); measure secreted APOA5/INHBE; test dependence using siRNA/CRISPR of HNF factors.
- **Priority:** medium

### Hypothesis 5
- **Hypothesis:** Under these conditions Baicalein does not induce canonical ferroptosis; enhanced FAO may buffer lipid peroxidation.
- **Why interesting:** No ferroptosis pathway signature observed despite target listing; lipid catabolism pathways are strongly induced.
- **Suggested validation:** Assess lipid ROS (C11-BODIPY), cell death rescue by ferrostatin-1/deferoxamine, and effects of GPX4 inhibition; measure ACSL4, SLC7A11, GPX4; test etomoxir co-treatment for altered sensitivity.
- **Priority:** low

## Confidence
high

## Caution notes
- Pathway inferences are based on proteomics-mapped enrichments of gene sets largely curated from transcriptomics; coverage and directionality may differ.
- Cell type and experimental context are unspecified; hepatic-like signatures (bile acid/coagulation, APOA5/ITIH3/INHBE) may be context-dependent and require corroboration with additional markers and functional assays.
- KRAS hallmark suppression may reflect global proliferation arrest rather than direct KRAS pathway inhibition; avoid inferring upstream blockade without signaling measurements (e.g., p-ERK).
- mTOR/autophagy interpretations are tentative; confirm with phospho-readouts (p-S6K/4EBP1) and autophagic flux assays.
- Increased OXPHOS gene-set signal could represent a stress-adaptive response; functional respiration measurements are needed.
- No explicit ferroptosis signature is detected here; ferroptotic mechanisms cannot be concluded from this dataset.
- Reactome/KEGG sections are empty; conclusions are drawn from consistent Hallmark and GO BP enrichments and protein changes.


---

# Balsalazide

# Drug summary: Balsalazide

## Executive summary
Balsalazide exposure produced a coherent proteomic and pathway response dominated by activation of IL6/JAK/STAT3 signaling (Hallmark NES ~2.00, FDR ~0.0014), consistent with the drug’s annotated pathway space. Concomitantly, a hypoxia-like program was up-regulated (Hallmark HYPOXIA NES ~1.79, FDR ~0.015), alongside decreases in multiple mitochondrial respiratory/import/translation components (e.g., NDUFA1, CHCHD4, MRPL52, MTRES1), indicating a shift away from oxidative metabolism. A pathway-level reduction in intermediate filament–based processes (GO BP NES ~−1.91, FDR ~0.041) suggests cytoskeletal remodeling, though this signal is borderline. At the protein level, several secreted/growth-regulatory factors increased (IGFBP3, EFNA1, SPINT1), and markers of metabolic stress/energy handling changed (TXNIP, ABHD5 up; AMACR, ACBD7 down), consistent with metabolic reprogramming but without pathway corroboration. Reactome/KEGG enrichments and GO BP up terms were absent, and total significant proteins were modest (n=47; 25 up/22 down), so interpretations beyond the core JAK/STAT and mitochondrial themes should be treated as tentative. Overall, the dominant biology is inflammatory/JAK–STAT engagement with associated metabolic stress and attenuation of mitochondrial function.

## Findings consistent with expected biology
- Robust enrichment of IL6/JAK/STAT3 signaling (Hallmark NES ~2.0, FDR ~0.001), aligning with the drug’s annotated interleukin/STAT pathway space.
- Increased abundance of select secreted/growth-regulatory proteins (e.g., IGFBP3, EFNA1, SPINT1) consistent with cytokine/STAT pathway engagement.

## Unexpected but plausible findings
- Up-regulation of a hypoxia-like program (Hallmark HYPOXIA NES ~1.79, FDR ~0.015).
- Coherent decreases in mitochondrial respiratory/import/translation machinery (e.g., NDUFA1, CHCHD4, MRPL52, MTRES1), suggesting reduced oxidative phosphorylation capacity.
- Pathway-level down-regulation of intermediate filament–based processes (GO BP NES ~−1.91), indicative of cytoskeletal remodeling (borderline significance).
- Shifts in lipid/energy handling markers (ABHD5 and TXNIP up; AMACR and ACBD7 down), compatible with reduced β-oxidation/peroxisomal activity and altered lipid droplet dynamics, but currently supported by few proteins (tentative).
- ZNRF3 up-regulation, potentially implying Wnt pathway modulation, based on a single protein (highly tentative).

## Follow-up hypotheses
### Hypothesis 1
- **Hypothesis:** Balsalazide induces metabolic and mitochondrial stress that attenuates oxidative phosphorylation and secondarily engages hypoxia-like and IL6/STAT3 programs.
- **Why interesting:** Integrates the strongest pathway signals (HYPOXIA up; IL6/JAK/STAT3 up) with coherent down-regulation of mitochondrial components, providing a unifying mechanism for the observed proteomic state.
- **Suggested validation:** Measure mitochondrial respiration and glycolysis (OCR/ECAR, spare respiratory capacity) by Seahorse; assess HIF-1α stabilization and target gene expression; quantify STAT3 activation (pY705/pS727 and nuclear localization); test mitochondrial import integrity (CHCHD4 substrates) and ROS levels.
- **Priority:** high

### Hypothesis 2
- **Hypothesis:** Balsalazide modestly remodels intermediate filament networks and cell architecture.
- **Why interesting:** GO BP down-regulation of intermediate filament–based processes suggests cytoskeletal effects that could influence adhesion and motility.
- **Suggested validation:** Immunofluorescence for vimentin/keratins and filament organization; cell shape and migration assays (wound healing/Transwell); transcriptomic GO/Cytoskeleton panels for corroboration.
- **Priority:** medium

### Hypothesis 3
- **Hypothesis:** Balsalazide alters lipid metabolism with reduced peroxisomal/β-oxidation and changes in lipid droplet handling.
- **Why interesting:** AMACR and ACBD7 decreases with ABHD5 increase point to reprogrammed lipid handling, which could couple to the observed metabolic stress.
- **Suggested validation:** Peroxisomal β-oxidation assays (e.g., pristanic/phytanic acid oxidation); lipid droplet quantification (BODIPY) and lipolysis assays; acylcarnitine/CoA profiling; examine peroxisome abundance (PEX markers).
- **Priority:** medium

### Hypothesis 4
- **Hypothesis:** Balsalazide dampens Wnt/β-catenin signaling via up-regulation of the negative regulator ZNRF3.
- **Why interesting:** If confirmed, Wnt modulation could contribute to anti-proliferative or differentiation-like effects in certain contexts.
- **Suggested validation:** TOPFlash/TCF reporter assays; β-catenin nuclear levels and target gene expression (e.g., AXIN2, MYC); test sensitivity to exogenous Wnt ligands or RSPOs.
- **Priority:** low

## Confidence
medium

## Caution notes
- Pathway enrichments are limited to Hallmark and a single GO BP down term; Reactome/KEGG and GO BP up are empty, constraining multi-pathway conclusions.
- Proteomic abundance does not directly report signaling activity; lack of phosphoproteomics/transcriptional data limits claims about STAT3 and HIF pathway activation.
- Several findings rely on a modest number of significant proteins (n=47) with generally moderate fold changes.
- The GO ‘INTERMEDIATE_FILAMENT_BASED_PROCESS’ signal is borderline (FDR ~0.041); cytoskeletal conclusions should be qualified.
- Single-protein-driven inferences (e.g., ZNRF3 for Wnt; ACBD7 for β-oxidation) are tentative and require orthogonal validation.
- TXNIP elevation can reflect metabolic stress independent of hypoxia and should not be over-interpreted as a HIF readout.
- Directionality of JAK/STAT engagement can be context-dependent; the proteomic signature suggests activation but requires confirmation by STAT phosphorylation/localization.


---

# Belzutifan

# Drug summary: Belzutifan

## Executive summary
Belzutifan treatment produced a robust proteomic signature of suppressed intermediate filament organization, supported by down-enrichment of GO biological processes (INTERMEDIATE_FILAMENT_ORGANIZATION and INTERMEDIATE_FILAMENT_BASED_PROCESS; NES ≈ -2, FDR < 0.01) and decreased KRT14. Several actin/cytoskeleton regulators (CDC42EP2, MICALL2, GAS2L3) were also reduced, consistent with broader cytoskeletal modulation. At the protein level, select extracellular matrix/adhesion-associated proteins (LAMA4, LAMC3, CD109, CD164, CEMIP) increased, but without pathway-level corroboration and thus best considered a trend. Additional protein-level decreases were observed in two DNA repair components (OGG1, POLK) and three lipid-metabolic enzymes (AMACR, LPIN1, ABHD4). IGFBP1 was among the most downregulated proteins. No hallmark or Reactome/KEGG pathway enrichments were detected, and no direct proteomic readouts of HIF target engagement were evident in this dataset. Overall, the most confident conclusion is downregulation of intermediate filament programs; other themes are plausible but remain tentative.

## Findings consistent with expected biology
- None listed.

## Unexpected but plausible findings
- Strong suppression of intermediate filament organization/processes (GO BP negative enrichment; decreased KRT14)
- Concomitant downregulation of two DNA repair–related proteins (OGG1, POLK)
- Downregulation of lipid-metabolic enzymes (AMACR, LPIN1, ABHD4)
- Protein-level uptrend in select ECM/adhesion-associated proteins (LAMA4, LAMC3, CD109, CD164, CEMIP) without pathway enrichment
- Marked decrease of IGFBP1 among top down-regulated proteins

## Follow-up hypotheses
### Hypothesis 1
- **Hypothesis:** Belzutifan reduces intermediate filament abundance/organization, altering cellular structural architecture in this context.
- **Why interesting:** Supported by two IF-related GO BP down-enrichments (NES ≈ -2, FDR < 0.01) and decreased KRT14; cytoskeletal changes could influence cell mechanics and stress responses.
- **Suggested validation:** Immunofluorescence imaging and fractionation for keratin IFs (e.g., KRT14 and other keratins), quantify filament integrity; assess cell mechanics (e.g., AFM) or resilience to shear/strain.
- **Priority:** high

### Hypothesis 2
- **Hypothesis:** Belzutifan attenuates select DNA repair activities (base excision and/or translesion synthesis) via reduced OGG1 and POLK.
- **Why interesting:** Coordinated decreases in OGG1 and POLK suggest potential impact on oxidative lesion processing and lesion bypass.
- **Suggested validation:** Expose cells to oxidative or alkylating damage (e.g., H2O2, KBrO3), measure 8-oxoG and comet assay tail moments; assess POLK-dependent bypass (e.g., UV or BPDE lesions); test rescue by overexpression.
- **Priority:** medium

### Hypothesis 3
- **Hypothesis:** Belzutifan shifts lipid metabolism through downregulation of AMACR, LPIN1, and ABHD4, altering cellular lipid composition.
- **Why interesting:** Concurrent decreases in three lipid enzymes imply reduced flux through specific lipid processing steps.
- **Suggested validation:** Targeted proteomics and enzymatic activity assays for AMACR/LPIN1/ABHD4; lipidomics to quantify acyl-CoA species, DAG/PA balance, and peroxisomal β-oxidation readouts.
- **Priority:** medium

### Hypothesis 4
- **Hypothesis:** Upregulation of laminin subunits and adhesion modulators reflects increased matrix engagement or secretion rather than broad ECM remodeling.
- **Why interesting:** Protein-level increases (LAMA4, LAMC3, CD109, CD164, CEMIP) suggest adhesion/ECM trends, though pathway support is lacking.
- **Suggested validation:** Secretome/matrisome proteomics and immunoblotting of conditioned media; adhesion/spreading assays on laminin; integrin activation/readouts (e.g., FAK phosphorylation) with careful controls.
- **Priority:** low

### Hypothesis 5
- **Hypothesis:** Observed proteomic changes occur with effective HIF pathway inhibition or are HIF-independent in this model.
- **Why interesting:** Belzutifan targets HIF/HIF prolyl-hydroxylase, yet no clear proteomic HIF target readouts are detected here.
- **Suggested validation:** Measure canonical HIF targets (e.g., CA9, GLUT1, VEGFA) at mRNA/protein level or via HRE-reporter assays under hypoxia; confirm on-target engagement in this cell context.
- **Priority:** high

### Hypothesis 6
- **Hypothesis:** Increased PET100 reflects enhanced assembly of mitochondrial complex IV in this context.
- **Why interesting:** Single-protein increase could indicate respiratory chain adjustment but lacks broader support.
- **Suggested validation:** Blue native PAGE and in-gel activity for complex IV; measure oxygen consumption rate and complex IV–specific respiration; replicate across doses/time.
- **Priority:** low

## Confidence
medium

## Caution notes
- Positive-direction trends (ECM/adhesion, contractile markers) lack pathway enrichment and are driven by small protein sets; interpret as trends, not broad programs.
- Several themes rest on few proteins (e.g., DNA repair n=2; lipid enzymes n=3); avoid inferring global capacity changes without additional evidence.
- Only proteomic data are provided; functional or multi-omic assays are needed to ascribe pathway activity or causality.
- Cell type/context is unspecified; cytoskeletal and ECM responses may be context-dependent.
- No clear proteomic readouts of HIF target engagement are evident; avoid attributing changes to HIF biology without corroboration.
- IGFBP1 is a secreted protein; intracellular levels may not mirror secretion dynamics or extracellular abundance.
- Inference of vesicle trafficking/endocytic downtrend (e.g., CLTCL1, SNAP47, HPS1) is tentative without pathway-level support.


---

# Benzamil (hydrochloride)

# Drug summary: Benzamil (hydrochloride)

## Executive summary
Benzamil (hydrochloride) elicited a robust, coherent proteomic signature characterized by: (1) a strong xenobiotic/AhR-like response (hallmark XENOBIOTIC_METABOLISM; top protein CYP1A1 log2FC ~7); (2) broad inflammatory/stress activation (IL6/JAK/STAT3, TNFA/NFkB, UV responses); (3) EMT/TGF-beta–associated remodeling with auxiliary morphogen features (hedgehog hallmark up; angiogenesis/hypoxia modestly up); (4) coordinated induction of membrane transport programs, especially amino acid import and monocarboxylate transport (e.g., SLC16A6); and (5) marked suppression of proliferative programs (E2F targets, G2/M checkpoint, and chromosome segregation processes). Several ECM/EMT-associated proteins (LOXL2, CEMIP, NRP1, AGR2, JUNB) and adaptors/regulators (SPRY2, SPRED1, GRAP, SOS1, NDFIP1) increase, while ESPL1 and multiple mitosis-associated processes decrease, consistent with an anti-proliferative state. Decreases in HMGCR and FADS2 suggest dampened lipid/cholesterol synthesis, though no dedicated pathway enrichment supports a global lipid program shift. DKK1 is reduced without evidence of canonical WNT activation (SP5 down; no WNT pathway enrichment), indicating a nuanced morphogen balance rather than overt WNT upregulation. Overall, the data support a non-proliferative, stress/inflammation and remodeling phenotype with a prominent xenobiotic signature; causal linkage to ion homeostasis remains plausible but unproven in this dataset. Absence of Reactome/KEGG enrichments does not detract from the strong Hallmark/GO evidence.

## Findings consistent with expected biology
- Suppression of proliferative cell-cycle programs (Hallmark E2F_TARGETS and G2M_CHECKPOINT down; broad GO chromosome segregation terms down), indicating an anti-proliferative proteomic state
- Activation of inflammatory and generalized stress signaling (Hallmark IL6_JAK_STAT3_SIGNALING, TNFA_SIGNALING_VIA_NFKB, UV_RESPONSE_UP/DN up), consistent with cellular stress under perturbation

## Unexpected but plausible findings
- Pronounced xenobiotic/AhR-like signature with massive CYP1A1 induction and Hallmark XENOBIOTIC_METABOLISM enrichment from an ion-channel modulator
- EMT/TGF-beta program activation with ECM/matrix remodeling proteins up (LOXL2, CEMIP, NRP1, AGR2, JUNB) and concurrent hypoxia/angiogenesis hallmarks (modest), suggesting a remodeling phenotype
- HEDGEHOG_SIGNALING hallmark enrichment without clear core pathway protein confirmation, implying possible morphogen pathway engagement that requires validation
- Broad upregulation of amino acid import/transport programs (multiple GO terms with high NES) and monocarboxylate transport (SLC16A6), concurrent with mitotic repression, consistent with non-proliferative nutrient adaptation
- Reduced DKK1 (a WNT antagonist) without evidence of WNT pathway activation (SP5 down; no WNT enrichment), indicating a nuanced morphogen shift rather than canonical WNT activation
- Downregulation of lipid/cholesterol biosynthesis enzymes (HMGCR, FADS2) without pathway-level support, suggesting a potential but unconfirmed decrease in lipogenesis

## Follow-up hypotheses
### Hypothesis 1
- **Hypothesis:** Benzamil engages the AhR/xenobiotic response program, driving CYP1A1 induction and contributing to downstream stress signaling.
- **Why interesting:** If confirmed, this off-target pathway could influence metabolic liabilities, drug–drug interactions, and safety profiles.
- **Suggested validation:** Assess a broader AhR target panel (CYP1B1, AHRR, NQO1, UGTs), AHR nuclear translocation, and sensitivity to AhR blockade (e.g., CH223191) or AHR knockdown/knockout to determine dependence.
- **Priority:** high

### Hypothesis 2
- **Hypothesis:** Perturbation of Na+/Ca2+ homeostasis by benzamil contributes to activation of inflammatory/stress pathways and anti-proliferative programs.
- **Why interesting:** Links the drug’s target class to the observed proteomic phenotype and could clarify mechanism-based liabilities or combination strategies.
- **Suggested validation:** Measure intracellular Na+ and Ca2+ dynamics (live-cell imaging), NCX activity/ion fluxes, and test rescue or phenocopy with ion manipulations and comparator NCX/channel inhibitors; evaluate proliferation markers (EdU/Ki-67) across dose/time.
- **Priority:** high

### Hypothesis 3
- **Hypothesis:** Benzamil induces a TGF-beta–driven EMT state with hedgehog/GLI crosstalk in the absence of canonical WNT activation.
- **Why interesting:** Points to pro-fibrotic/stromal remodeling risk and identifies potential pathway liabilities or therapeutic combination targets.
- **Suggested validation:** Monitor phospho-SMAD2/3 and EMT markers (e.g., Vimentin, LOXL2, CEMIP), assess GLI1/GLI2 and PTCH1, and test pathway dependence using TGF-beta receptor and SMO inhibitors; include cell migration/invasion assays.
- **Priority:** medium

### Hypothesis 4
- **Hypothesis:** Upregulated amino acid and monocarboxylate transport supports a survival-oriented metabolic adaptation during growth arrest.
- **Why interesting:** Reveals a potential nutrient-dependence that may be exploitable with transporter inhibitors.
- **Suggested validation:** Quantify amino acid and lactate/pyruvate uptake, perform targeted metabolomics and mTORC1 readouts, and test sensitivity to LAT1/MCT inhibitors under benzamil treatment.
- **Priority:** medium

### Hypothesis 5
- **Hypothesis:** Benzamil reduces de novo lipid/cholesterol synthesis as part of the anti-proliferative response.
- **Why interesting:** Could identify additional metabolic vulnerabilities or safety considerations related to membrane/lipid homeostasis.
- **Suggested validation:** Measure SREBP target activation and de novo lipogenesis/cholesterol synthesis (e.g., isotope tracing), assess cholesterol levels, and probe rescue with mevalonate pathway intermediates.
- **Priority:** low

## Confidence
high

## Caution notes
- Interpret the anti-proliferative signature as proteomic evidence of reduced proliferation, not definitive cytostasis without functional assays.
- Treat the AhR conclusion as AhR-like; confirm with expanded marker panels and pharmacological/genetic perturbation before asserting pathway engagement.
- Hedgehog and angiogenesis/hypoxia signals derive primarily from Hallmark enrichment; validate with pathway-specific effectors (e.g., GLI1/PTCH1, HIF1A/VEGF).
- The link to NCX/ion homeostasis is inferential; direct measurements of Na+/Ca2+ fluxes are needed to support causality.
- Do not overinterpret metabolic rewiring from transport signatures alone; complement with metabolomics/phosphoproteomics.
- DKK1 reduction should not be taken as WNT activation in light of SP5 down and absent WNT enrichment; confirm with beta-catenin/TCF readouts.
- Replicate across cell models and confirm dose–response/time dependence to exclude cell line–specific or solvent-driven effects.
- Absence of Reactome/KEGG enrichments does not diminish the strong Hallmark/GO signals but limits pathway granularity.


---

# Benztropine (mesylate)

# Drug summary: Benztropine (mesylate)

## Executive summary
Benztropine (mesylate) exposure yields a coherent proteomic profile marked by: (1) strong activation of sterol/cholesterol and broader lipid metabolism (multiple Hallmark/GO enrichments; increased SC5D, NPC1L1, ANGPTL8, APOA5); (2) robust induction of IL6–JAK–STAT3 and inflammatory modules with complement/coagulation and angiogenesis enrichments; and (3) pronounced suppression of proliferation programs (E2F/MYC, G2/M) alongside broad repression of ribosome biogenesis/translation. Neuronal projection/vesicle-associated features are enriched (e.g., SV2A, TMEM59L, NRSN2), aligning with a neuropharmacologic context. Developmental signaling changes are suggested, with Hedgehog enrichment and mixed Wnt regulator shifts (ZNRF3 up; DKK1 down). Evidence for increased membrane protein ectodomain proteolysis and a possible tilt toward mTORC1/glycolytic metabolism is present at the enrichment level but remains preliminary. Reactome and KEGG yielded no significant terms in this run; interpretations rely on Hallmark and GO. Overall, the data indicate inflammatory stress and lipid remodeling concurrent with cytostatic growth arrest.

## Findings consistent with expected biology
- Induction of IL6–JAK–STAT3 and broader inflammatory modules, with complement/coagulation enrichment.
- Enrichment of neuronal projection/vesicle-associated features consistent with neuronal signaling context.

## Unexpected but plausible findings
- Robust activation of cholesterol/sterol and lipoprotein metabolic programs with increased SC5D, NPC1L1, ANGPTL8, and APOA5.
- Suppression of cell-cycle programs (E2F/MYC, G2/M) and marked downregulation of ribosome biogenesis/translation.
- Upregulation of angiogenesis and epithelial–mesenchymal transition signatures, suggesting tissue/ECM remodeling.
- Hedgehog signaling enrichment with concurrent Wnt regulator shifts (ZNRF3 up; DKK1 down), indicating altered developmental signaling.
- Enrichment of membrane protein ectodomain proteolysis without direct sheddase markers.
- Preliminary indication of reduced autophagy components (ULK1, WDFY3 down) with mTORC1/glycolysis enrichments.

## Follow-up hypotheses
### Hypothesis 1
- **Hypothesis:** Lipid/cholesterol metabolic remodeling is driven by SREBP-dependent and/or JAK–STAT–linked mechanisms under benztropine.
- **Why interesting:** Explains coordinated upregulation of sterol biosynthesis/transport and lipid regulators (SC5D, NPC1L1, ANGPTL8, APOA5) together with IL6–JAK–STAT3 enrichment.
- **Suggested validation:** Assess SREBP1/2 cleavage and nuclear translocation; measure STAT3 phosphorylation; perform lipidomics and cellular cholesterol assays; test dependence on JAK inhibition and mTORC1 inhibition for the lipid signature.
- **Priority:** high

### Hypothesis 2
- **Hypothesis:** Benztropine induces cytostatic growth arrest with translational downregulation rather than overt cytotoxicity.
- **Why interesting:** Consistent with E2F/MYC and G2/M suppression and strong repression of ribosome biogenesis/translation.
- **Suggested validation:** Time- and dose-dependent EdU incorporation and phospho-H3 assays; MYC protein assessment; pre-rRNA (47S) and polysome profiling; viability/apoptosis metrics to distinguish cytostatic vs cytotoxic effects.
- **Priority:** high

### Hypothesis 3
- **Hypothesis:** Hedgehog pathway activity increases and Wnt signaling is perturbed following benztropine treatment.
- **Why interesting:** Hedgehog hallmark is enriched and key Wnt regulators shift (ZNRF3 up; DKK1 down), suggesting altered extracellular signaling.
- **Suggested validation:** Measure GLI1/2 transcript/protein as Hedgehog readouts; TCF/LEF reporter assays for Wnt activity; quantify secreted DKK1; probe sensitivity to Hedgehog (e.g., SMO) or Wnt pathway modulators.
- **Priority:** medium

### Hypothesis 4
- **Hypothesis:** Surface receptor ectodomain shedding increases, modulating cytokine responsiveness.
- **Why interesting:** GO enrichment for membrane protein ectodomain proteolysis alongside increased cytokine receptor components (IL4R, IL10RB, TNFRSF12A).
- **Suggested validation:** Quantify soluble ectodomains (e.g., sIL4R, sIL10RB) in conditioned media; assess ADAM10/17 activity and protease abundance; test effects of metalloprotease inhibitors on shedding and signaling.
- **Priority:** medium

### Hypothesis 5
- **Hypothesis:** Autophagy is suppressed via mTORC1-associated mechanisms.
- **Why interesting:** ULK1 and WDFY3 are decreased while MTORC1 and glycolysis gene sets are enriched.
- **Suggested validation:** Monitor LC3-II and p62 flux with/without bafilomycin; assess ULK1 and S6K/4E-BP1 phosphorylation; metabolic assays for glycolytic rate; evaluate reversibility with mTORC1 inhibitors.
- **Priority:** medium

### Hypothesis 6
- **Hypothesis:** Benztropine perturbs vesicle trafficking/endolysosomal pathways, reflected by increased SV2A, TMEM59L, and NRSN2.
- **Why interesting:** Aligns with neuronal signaling context and observed increases in vesicle/membrane-associated proteins.
- **Suggested validation:** Assay synaptic vesicle recycling/endocytosis; track SV2A trafficking; examine endolysosomal markers (LAMP1, Rab proteins) and extracellular vesicle release.
- **Priority:** medium

## Confidence
medium

## Caution notes
- Pathway ‘activation’ is inferred from protein abundance-based enrichments; without phospho-proteomics or transcriptomics, describe these as enrichments rather than definitive activation.
- Links to primary GPCR targets (mAChR, dopamine, histamine receptors) are indirect; observed changes may reflect downstream or off-target effects.
- Reactome and KEGG yielded no significant pathways; interpretation relies on Hallmark and GO enrichments in this run.
- Cell-type context is unspecified; angiogenesis and EMT signatures can reflect ECM/adhesion remodeling rather than true angiogenic or EMT transitions.
- Several observations hinge on few markers (e.g., autophagy from ULK1/WDFY3; Wnt from DKK1/ZNRF3; ectodomain proteolysis without protease evidence); treat as preliminary.
- Growth arrest signatures could arise from cytostatic or cytotoxic effects; verify dose, time, and viability to distinguish mechanisms.
- Increased lipid/lipoprotein signatures indicate lipid remodeling but do not imply hepatocyte-like identity; confirm with lipidomics and lineage markers if relevant.


---

# Bepotastine

# Drug summary: Bepotastine

## Executive summary
Bepotastine exposure yielded a balanced proteomic response (25 up, 25 down) dominated by two pathway-level signals: (1) significant enrichment of inflammatory/TNFA signaling via NF-κB (Hallmark NES ~1.78, FDR ~0.034) and (2) a hypoxia-like program (Hallmark NES ~1.75, FDR ~0.024). In parallel, there is robust suppression of intermediate filament/keratin organization (GO BP down; NES ≤ −2.27, FDR = 0), supported by decreases in keratin/barrier proteins (e.g., KRT14, FLG2). Additional single- or few-protein changes suggest possible metabolic and signaling adjustments (e.g., mitochondrial/chromatin-associated proteins up such as TOMM5, TK2, TFAP2C, SETD1B; peroxisomal enzymes AMACR, PHYH down; select GPCR/calcium components altered, including F2R up, GNG4 and CALML3 down; PTEN down). These latter features are hypothesis-generating in the absence of pathway-level corroboration. No Reactome/KEGG enrichments were detected, and some pathway sections are empty; conclusions are therefore focused on the strongest Hallmark/GO evidence.

## Findings consistent with expected biology
- Engagement of inflammatory signaling pathways: Hallmark TNFA_SIGNALING_VIA_NFKB is significantly enriched among up-regulated proteins (NES ~1.78, FDR ~0.034), consistent with the drug’s Immunology/Inflammation domain.

## Unexpected but plausible findings
- Robust suppression of intermediate filament/keratin programs (GO BP down; NES ≤ −2.27, FDR = 0), with KRT14 and FLG2 decreased, indicating cytoskeletal/barrier remodeling in this model.
- Activation of a hypoxia-like stress program (Hallmark HYPOXIA; NES ~1.75, FDR ~0.024), suggesting metabolic/stress adaptation alongside inflammatory signaling.
- Putative attenuation of peroxisomal fatty-acid/alpha-oxidation (AMACR and PHYH down); suggestive but based on two proteins and no pathway enrichment.
- Isolated shifts in GPCR and calcium signaling components (F2R up; GNG4 and CALML3 down), implying possible signaling rewiring; tentative without pathway support.
- Single-protein decrease in PTEN, which could impact PI3K/AKT signaling; tentative and requires validation.
- Upregulation of transcriptional/chromatin and developmental regulators (e.g., TFAP2C, SETD1B, SUFU) and select RTK/axon guidance cues (e.g., TYRO3, EFNA1); suggests broader rewiring but currently lacks pathway-level support.

## Follow-up hypotheses
### Hypothesis 1
- **Hypothesis:** Bepotastine exposure induces NF-κB-related inflammatory signaling and a hypoxia-like stress response in this cell context.
- **Why interesting:** Both Hallmark TNFA via NF-κB and Hypoxia are significantly enriched, indicating coordinated stress/inflammatory pathway engagement that could influence downstream phenotypes and drug-response biology.
- **Suggested validation:** Assess NF-κB activity (p65 nuclear translocation, IκBα phosphorylation; qPCR of canonical targets) and HIF pathway engagement (HIF-1α stabilization, VEGFA/CA9 mRNA); perform time course and dose–response; consider oxygen consumption and glycolytic flux measurements.
- **Priority:** high

### Hypothesis 2
- **Hypothesis:** Bepotastine disrupts intermediate filament/keratin network organization and epithelial/barrier features.
- **Why interesting:** GO BP down enrichment is strong (FDR = 0) with decreased KRT14 and FLG2, suggesting a structural phenotype that may affect cell integrity and barrier function.
- **Suggested validation:** Immunofluorescence for keratin networks (K14, K5), assessment of cell morphology, transepithelial electrical resistance (if applicable), and RNA confirmation of keratin/barrier gene expression; include time/dose dependence.
- **Priority:** high

### Hypothesis 3
- **Hypothesis:** Peroxisomal fatty-acid/alpha-oxidation capacity is reduced following exposure.
- **Why interesting:** Downregulation of AMACR and PHYH aligns with the hypoxia-like signature and implies a shift away from oxygen-intensive lipid metabolism.
- **Suggested validation:** Measure peroxisomal alpha-/beta-oxidation activity, targeted lipidomics for phytanic/pristanic acid derivatives, and peroxisomal markers (PEX proteins) by WB/IF; evaluate rescue or enhancement under normoxic vs hypoxic conditions.
- **Priority:** medium

### Hypothesis 4
- **Hypothesis:** GPCR signaling dynamics are altered (e.g., enhanced PAR1/F2R signaling and reduced Gβγ availability via GNG4 down), potentially affecting thrombin/ERK/Ca2+ responses.
- **Why interesting:** F2R up with GNG4 and CALML3 down suggests a shift in GPCR and calcium signaling components, which could influence responsiveness to protease-activated receptor stimuli.
- **Suggested validation:** Test thrombin-induced signaling (Ca2+ flux, ERK phosphorylation), evaluate G-protein subunit levels and coupling, and perform pharmacologic challenge assays; include multiple time points and doses.
- **Priority:** low

### Hypothesis 5
- **Hypothesis:** PTEN reduction increases PI3K/AKT pathway activity in this context.
- **Why interesting:** PTEN is decreased at the protein level, which could relieve suppression of PI3K/AKT signaling and impact survival/growth pathways.
- **Suggested validation:** Confirm PTEN abundance by immunoblot, assess AKT phosphorylation (Ser473/Thr308) and downstream targets, and test for dose–response and reversibility; consider phosphatase activity assays.
- **Priority:** low

## Confidence
medium

## Caution notes
- Pathway enrichment is limited (two Hallmark up and two GO BP down; no Reactome/KEGG), so conclusions beyond these signals should be conservative.
- Several mechanistic inferences rely on one to two proteins; treat these as hypothesis-generating until corroborated.
- Inflammatory (NF-κB) and hypoxia signatures indicate pathway engagement but do not establish on-target causality or direction relative to histamine receptor pharmacology.
- Proteomics reflects abundance, not pathway activity; NF-κB/HIF activation claims require activity markers or transcript-level support.
- The strong keratin/intermediate filament signal may be cell-type specific; avoid broad generalization without additional models.
- Hypoxia hallmark enrichment may reflect a generalized stress response rather than true hypoxia; confirm with HIF targets or oxygen-related measurements.
- Potential off-target or concentration-dependent effects cannot be excluded without dose–response and time-course data.
- Single-protein findings with mechanistic implications (e.g., PTEN) require orthogonal validation.


---

# Betahistine

# Drug summary: Betahistine

## Executive summary
Betahistine exposure produces a robust cytoskeletal signature dominated by suppression of intermediate filament organization (two GO BP sets, strong negative NES, FDR 0). A concurrent enrichment of the Hallmark KRAS_SIGNALING_DN set among down-regulated proteins is present but should be interpreted cautiously and not taken as evidence of KRAS pathway inhibition or activation. Protein-level shifts suggest extracellular matrix and adhesion remodeling (LOXL2, CEMIP, IGFBP5 up; EFEMP1, SULF2 down), alongside indications of mitochondrial adaptation (CHCHD4 down with PET100 and UQCC6 up; MSRA up). Up-regulation of SMPD2 and GAL3ST1 points to a potential shift in sphingolipid/ceramide metabolism, and several ubiquitin/proteostasis factors change in abundance, though without pathway-level corroboration. No up-regulated immune/inflammatory pathways were detected in this dataset. Overall, the data support a primary structural/cytoskeletal remodeling phenotype with secondary metabolic adaptations; additional observations are hypothesis-generating and require validation.

## Findings consistent with expected biology
- Strong pathway-level suppression of intermediate filament organization, indicating cytoskeletal remodeling (GO BP down-regulation with FDR 0).
- Protein-level evidence for ECM and adhesion remodeling accompanying structural changes (e.g., LOXL2 and CEMIP up; EFEMP1 and SULF2 down).

## Unexpected but plausible findings
- Enrichment of the Hallmark KRAS_SIGNALING_DN set among down-regulated proteins (NES approximately −2, FDR 0); association noted but directionality of KRAS pathway activity cannot be inferred.
- Convergent mitochondrial signals suggesting compensatory responses (CHCHD4 down; PET100 and UQCC6 up; MSRA up).
- Up-regulation of sphingolipid-related enzymes (SMPD2 and GAL3ST1), consistent with a possible shift in ceramide/glycosphingolipid metabolism (protein-level only).
- Absence of up-regulated immune/inflammatory pathway enrichment despite histamine/GPCR annotation; within this model, no pathway-level immune activation is evident.
- Changes in select ubiquitin/proteostasis components (e.g., MKRN1, OTUD3, MSL2, DCAF12 down; UBE3D up) without pathway enrichment, suggesting preliminary proteostasis perturbation.
- Single-protein indication potentially related to Wnt regulation (ZNRF3 up), highly preliminary and not sufficient to assign pathway directionality.

## Follow-up hypotheses
### Hypothesis 1
- **Hypothesis:** Betahistine suppresses intermediate filament organization, altering cellular architecture and motility.
- **Why interesting:** Two independent GO BP sets show strong down-regulation (FDR 0), aligning with a dominant cytoskeletal phenotype.
- **Suggested validation:** Quantify vimentin and keratin networks by immunofluorescence and immunoblot; assess filament solubility; measure cell shape, adhesion, and migration (e.g., wound-healing or transwell assays).
- **Priority:** high

### Hypothesis 2
- **Hypothesis:** Betahistine perturbs the mitochondrial protein import/assembly axis and induces compensatory oxidative stress responses.
- **Why interesting:** CHCHD4 is down while PET100 and UQCC6 are up and MSRA increases, suggesting import stress with compensatory respiratory assembly and redox adaptation.
- **Suggested validation:** Measure CHCHD4/MIA40 pathway activity with import reporters; assess respiratory complex assembly (BN-PAGE) and oxygen consumption; quantify ROS and MSRA dependency; evaluate mitochondrial membrane potential.
- **Priority:** high

### Hypothesis 3
- **Hypothesis:** Betahistine shifts sphingolipid metabolism toward increased ceramide/glycosphingolipid turnover that contributes to cytoskeletal remodeling.
- **Why interesting:** SMPD2 and GAL3ST1 are up-regulated, indicating potential changes in ceramide and sulfated glycosphingolipids.
- **Suggested validation:** Perform targeted lipidomics for ceramides, sphingomyelins, and sulfatides; pharmacologically or genetically modulate SMPD2 to test impact on intermediate filament and adhesion phenotypes.
- **Priority:** medium

### Hypothesis 4
- **Hypothesis:** The KRAS_SIGNALING_DN enrichment reflects modulation of RAS–MAPK-adjacent programs rather than direct KRAS inhibition/activation.
- **Why interesting:** Strong hallmark enrichment among down-regulated proteins, yet directionality is ambiguous at the proteome level.
- **Suggested validation:** Evaluate ERK phosphorylation and RAS activity; profile canonical KRAS target transcripts; test whether MEK/ERK pathway modulation alters the observed cytoskeletal and mitochondrial signatures.
- **Priority:** medium

### Hypothesis 5
- **Hypothesis:** The observed structural changes are mediated by histamine-receptor–linked GPCR signaling in this model.
- **Why interesting:** Cytoskeletal remodeling is compatible with GPCR outputs, and the compound is annotated to histamine receptors while immune pathway activation is not detected here.
- **Suggested validation:** Define histamine receptor expression; use selective antagonists or genetic knockdown to determine whether blocking receptor signaling attenuates the cytoskeletal and mitochondrial phenotypes.
- **Priority:** medium

### Hypothesis 6
- **Hypothesis:** Betahistine may weakly tilt Wnt pathway regulation via ZNRF3 up-regulation.
- **Why interesting:** ZNRF3, a Wnt pathway E3 ligase, is increased, but this single-protein observation is insufficient for pathway assignment.
- **Suggested validation:** Run TOPFlash/TCF reporter assays, assess LRP6 phosphorylation, and test sensitivity to RSPO modulation to evaluate Wnt pathway activity.
- **Priority:** low

## Confidence
medium

## Caution notes
- Only down-regulated pathways reached significance; no up-regulated pathway categories were enriched, so inferences about up-programs are based on limited protein-level patterns.
- Many secondary observations (ECM remodeling, sphingolipids, ubiquitin/proteostasis, chromatin factors, Wnt) rely on small protein sets without pathway-level corroboration and should be considered hypothesis-generating.
- Interpretation of KRAS_SIGNALING_DN enrichment is non-directional; do not infer KRAS pathway inhibition or activation without additional data.
- Intermediate filament suppression is strongly supported at the pathway level, but leading-edge proteins are not detailed here, limiting mechanistic specificity.
- Lack of immune/inflammatory enrichment may reflect the cell model and receptor expression context rather than a general property of the compound.
- Conclusions are drawn from steady-state protein abundance; no phosphoproteomic, transcriptomic, or functional assays are provided to link changes to histamine/GPCR signaling.
- Potential off-target effects or indirect adaptations (e.g., mitochondrial and lipid changes) cannot be excluded.


---

# Betamethasone dipropionate

# Drug summary: Betamethasone dipropionate

## Executive summary
Proteomics of betamethasone dipropionate exposure shows clear on‑target engagement of the glucocorticoid pathway, evidenced by a significant reduction of NR3C1 protein. The most coherent program-level signal is induction of metal‑ion response/detoxification pathways (GO: response to zinc and copper; FDR < 0.05), supported by metallothionein MT1E up‑regulation. Xenobiotic metabolism markers are increased (notably CYP1A1 and CYP3A7), indicating activation of drug‑metabolizing capacity, though pathway-level enrichment is not provided. Intermediate filament organization is significantly suppressed (GO down; FDR ~0.038) with reduced KRT17, suggesting cytoskeletal remodeling. Select secreted/ECM factors (DKK1, SPP1, NPNT) are decreased, consistent with an anti-inflammatory/secretory dampening motif but based on a small set. Additional coordinated increases in membrane/caveolae and transport-related proteins (CAVIN2, FXYD1, SLC16A6, SC5D) and a modest PTEN decrease are noteworthy but remain interpretive without pathway support. Overall, the data support GR engagement with stress‑adaptive detoxification and structural remodeling, while some signals (xenobiotic enzymes, membrane/caveolae changes, PTEN reduction) warrant cautious follow‑up.

## Findings consistent with expected biology
- On‑target glucocorticoid receptor engagement: NR3C1 protein decrease (log2FC ~ -0.92; FDR-significant).
- Decreases in select secreted factors (e.g., SPP1, DKK1, NPNT), consistent with anti‑inflammatory/immune‑modulatory effects, noting this is based on a limited protein set.

## Unexpected but plausible findings
- Induction of metal‑ion response programs (GO: response to zinc and copper; NES ~1.96–1.97, FDR < 0.05) with MT1E up‑regulation.
- Up‑regulation of xenobiotic/drug‑metabolizing enzymes (CYP1A1, CYP3A7) without pathway enrichment, suggesting increased metabolic capacity.
- Suppression of intermediate filament organization (GO down; NES ~ -2.01, FDR ~0.038) with reduced KRT17.
- Coordinated increases in membrane/caveolae and transport/lipid‑handling proteins (CAVIN2, FXYD1, SLC16A6, SC5D) suggesting membrane remodeling in the absence of pathway enrichment.
- Modest decrease in PTEN alongside reductions in several ubiquitin/ligase or adaptor proteins (e.g., USP12, OTUD3, ZMIZ2), raising but not establishing altered signaling/turnover.

## Follow-up hypotheses
### Hypothesis 1
- **Hypothesis:** Betamethasone engages GR‑dependent pathways that secondarily induce metallothioneins and metal‑ion detoxification responses.
- **Why interesting:** Strong GO enrichment for zinc/copper responses with MT1E up‑regulation indicates a robust, potentially protective stress‑adaptation program.
- **Suggested validation:** Time‑course qPCR/proteomics for MT1 family (MT1E/MT1X/MT2A); intracellular zinc/copper buffering assays; test GR dependence using RU486 or NR3C1 knockdown; assess promoter activity or chromatin binding if feasible.
- **Priority:** high

### Hypothesis 2
- **Hypothesis:** Xenobiotic enzyme induction (CYP1A1, CYP3A7) arises via nuclear receptor cross‑talk (e.g., AhR for CYP1A1; PXR/CAR for CYP3A7) rather than GR alone.
- **Why interesting:** Differential CYP regulation could impact drug metabolism and safety liabilities in combination settings.
- **Suggested validation:** Measure CYP activities (e.g., EROD for CYP1A1; substrate assays for CYP3A); test receptor involvement using antagonists/siRNA (AhR inhibitor, PXR/CAR modulators); reporter assays for AhR/PXR/CAR; evaluate dose/time dependence.
- **Priority:** medium

### Hypothesis 3
- **Hypothesis:** Betamethasone remodels caveolae/membrane organization and ion/lipid transport capacity (via CAVIN2, FXYD1, SLC16A6, SC5D).
- **Why interesting:** Membrane remodeling could alter signaling compartmentalization, transport, and pharmacokinetics of co‑administered agents.
- **Suggested validation:** Quantify caveolin/cavin complexes (CAV1/CAV2/CAVIN2); assess caveolae abundance (electron microscopy or membrane fractionation); measure ion transport and lactate/monocarboxylate flux; test GR dependence and reversibility.
- **Priority:** medium

### Hypothesis 4
- **Hypothesis:** Suppression of intermediate filament organization reflects a cytoskeletal adaptation to glucocorticoid exposure with potential effects on cell mechanics and migration.
- **Why interesting:** Cytoskeletal changes can influence barrier function, adhesion, and tissue remodeling under steroid therapy.
- **Suggested validation:** Immunofluorescence and western blots for keratins (e.g., KRT17, KRT8/18) and IF regulators; functional assays for cell stiffness/motility; replicate pathway enrichment in larger protein sets/time points.
- **Priority:** medium

### Hypothesis 5
- **Hypothesis:** Modest PTEN reduction shifts signaling toward PI3K/AKT activation in a subset of contexts.
- **Why interesting:** Even small PTEN decreases can modulate survival signaling, potentially affecting steroid responses and combination strategies.
- **Suggested validation:** Assess p‑AKT/p‑S6 by western blot or phospho‑proteomics; rescue with PTEN overexpression; test sensitivity to PI3K/mTOR inhibitors; confirm reproducibility across doses and time points.
- **Priority:** low

## Confidence
medium

## Caution notes
- Several inferences rely on small protein sets (e.g., secreted factors, caveolae/membrane proteins); interpret program‑level conclusions cautiously.
- Mechanistic attribution to GR versus other nuclear receptors is not established by proteomics alone; NR3C1 reduction supports engagement but does not prove downstream causality.
- Xenobiotic metabolism and metal‑ion responses may reflect cell‑type‑specific stress adaptations; orthogonal assays and pathway enrichment would strengthen interpretation.
- Single‑protein changes of modest magnitude (e.g., PTEN, KRT17) should not be over‑interpreted without corroborating pathway or functional evidence.
- Pathway enrichment is limited (no Hallmark/Reactome/KEGG hits; reliance on GO terms), constraining confidence in broad pathway narratives.
- Data are from a single omics layer and likely a single condition; time‑course, dose‑response, and transcriptomic validation are recommended.


---

# Betamethasone disodium phosphate

# Drug summary: Betamethasone disodium phosphate

## Executive summary
Betamethasone disodium phosphate elicited a proteomic program dominated by strong suppression of biosynthetic and proliferative machinery, with broad down-enrichment of rRNA processing, ribosome biogenesis, and RNA splicing/mRNA processing, alongside reduced MYC target and G2/M checkpoint hallmarks. Concomitantly, epithelial–mesenchymal transition and hypoxia hallmarks were up, with concordant increases in extracellular matrix/secreted factors (e.g., SERPINE1, LOXL2, FSTL3, MATN3, IGFBP3), indicating a stress-adaptive mesenchymal remodeling state. Inflammatory-associated hallmarks (TNFA/NFκB, IL6/STAT3) were enriched, suggesting a cytokine/stress-response component rather than definitive pathway activation. Cholesterol homeostasis was up-enriched with increased CAVIN2, consistent with membrane/lipid remodeling. Xenobiotic/steroid-metabolizing enzymes (CYP1A1, CYP3A7) increased and NR3C1 decreased, hinting at receptor/metabolic feedback; however, nuclear receptor mechanism inferences lack pathway-level support. Overall, the profile indicates curtailed biosynthetic capacity with a shift toward mesenchymal/stress programs; mechanistic assertions beyond these core findings should be considered provisional.

## Findings consistent with expected biology
- Robust down-regulation of biosynthetic programs: rRNA processing, ribosome biogenesis, and RNA splicing/mRNA processing are strongly suppressed (GO BP down with large-magnitude NES and FDR≈0).
- Attenuation of proliferative tone: MYC target hallmarks and G2/M checkpoint are down-enriched, consistent with reduced cell-cycle progression.
- Mesenchymal/stress remodeling: EMT and hypoxia hallmarks are up with concordant increases in SERPINE1, LOXL2, FSTL3, MATN3, IGFBP3 and reduced KRT14.
- Lipid/cholesterol remodeling: Hallmark CHOLESTEROL_HOMEOSTASIS is up and CAVIN2 is increased, suggesting changes in membrane/lipid handling.

## Unexpected but plausible findings
- Enrichment of TNFA/NFκB and IL6/STAT3 hallmarks in a suppressed-growth context, indicating a stress/inflammatory signature despite broad anti-biosynthetic effects (without definitive evidence of canonical pathway activation).
- DKK1 down-regulation occurring alongside EMT up-enrichment, hinting at a possible WNT-axis shift; this rests on single-protein evidence and lacks pathway support.
- Marked induction of MIXL1 (log2FC ~4.6) without pathway-level corroboration, suggesting a potential lineage/stemness-related stress response that remains speculative.
- CYP1A1/CYP3A7 up-regulation with decreased NR3C1, suggesting steroid/xenobiotic metabolism engagement and possible receptor feedback, but lacking confirmatory pathway enrichment.

## Follow-up hypotheses
### Hypothesis 1
- **Hypothesis:** The compound suppresses MYC-linked ribosome biogenesis and RNA processing, reducing translational capacity and proliferative drive.
- **Why interesting:** This provides a coherent mechanism for the dominant anti-biosynthetic signature and connects multiple down-enriched programs.
- **Suggested validation:** Assess rRNA synthesis (5-EU incorporation, pre-rRNA qPCR), polysome profiling for global translation, and MYC levels/activity; verify down-regulation of additional ribosome biogenesis factors.
- **Priority:** high

### Hypothesis 2
- **Hypothesis:** Treatment induces an EMT-like, hypoxia-associated stress state with ECM remodeling.
- **Why interesting:** Such a state may alter adhesion, motility, and therapy responses, and is strongly supported by hallmarks and ECM protein increases.
- **Suggested validation:** Measure EMT markers (Vimentin, Fibronectin up; E-cadherin down), HIF target genes, and matrix remodeling enzymes; assess morphology and migration assays.
- **Priority:** high

### Hypothesis 3
- **Hypothesis:** Inflammatory hallmark enrichments reflect a stress-responsive cytokine program rather than canonical NFκB/STAT3 activation.
- **Why interesting:** Distinguishing stress-associated transcriptional programs from true pathway activation informs mechanism and combinatorial strategies.
- **Suggested validation:** Quantify secreted cytokines (e.g., IL-6, SERPINE1) by ELISA; test RELA (p65) nuclear translocation and STAT3 Y705 phosphorylation; profile a broader NFκB/STAT3 target panel.
- **Priority:** medium

### Hypothesis 4
- **Hypothesis:** Cellular lipid/cholesterol handling and membrane microdomain organization are remodeled.
- **Why interesting:** Membrane/lipid changes can influence signaling and drug uptake; current evidence (CAVIN2, hallmark enrichment) suggests this axis.
- **Suggested validation:** Measure cellular cholesterol (filipin staining, enzymatic assays), examine SREBF2 target expression, and assess CAV1/CAV2 levels and caveolae by microscopy.
- **Priority:** medium

### Hypothesis 5
- **Hypothesis:** Steroid/xenobiotic metabolism is engaged with feedback on steroid receptor abundance/activity.
- **Why interesting:** CYP induction with NR3C1 decrease suggests receptor/metabolic adaptation that could modulate response dynamics.
- **Suggested validation:** Evaluate NR3C1 localization and transcriptional activity (GRE-reporter), test GR ligand competition, and measure CYP1A1/3A activity; check additional nuclear receptor targets.
- **Priority:** low

### Hypothesis 6
- **Hypothesis:** A WNT/β-catenin balance shift contributes to the EMT-like state.
- **Why interesting:** DKK1 reduction may relieve WNT inhibition, potentially reinforcing mesenchymal programs.
- **Suggested validation:** Assess β-catenin nuclear localization, TOPFlash reporter activity, and expression of WNT targets (e.g., AXIN2, CCND1).
- **Priority:** low

## Confidence
high

## Caution notes
- Inflammatory hallmark enrichments (TNFA/NFκB, IL6/STAT3) indicate signature similarity and do not alone establish pathway activation; canonical components and phosphorylation/localization evidence are needed.
- Caveolae/membrane remodeling is inferred largely from CAVIN2 plus hallmark enrichment; corroborate with additional lipid/caveolar markers (e.g., CAV1/CAV2) before firm conclusions.
- NR3C1 protein reduction and CYP increases suggest receptor/metabolic feedback but do not confirm nuclear receptor activation status; proteomics lacks subcellular/activity resolution.
- Hallmark MYOGENESIS often reflects cytoskeletal/ECM remodeling in non-muscle cells; avoid interpreting as true myogenic differentiation without orthogonal evidence.
- Hallmark ANGIOGENESIS down may not imply anti-angiogenic activity outside endothelial contexts; interpret in light of cell type and system.
- Mechanistic specificity is limited by reliance on GSEA of a single proteomic dataset with no GO BP up and no Reactome/KEGG enrichments; key conclusions should be validated functionally.
- Proteomic abundance changes do not necessarily reflect TF activity states or post-translational regulation; complement with phosphorylation/localization assays.


---

# Betamethasone valerate

# Drug summary: Betamethasone valerate

## Executive summary
Betamethasone valerate exposure yields a coherent proteomic signature dominated by broad suppression of rRNA processing, ribosome biogenesis, RNA splicing/mRNA processing, and cytoplasmic translation, alongside negative enrichment of MYC target programs and angiogenesis. These features indicate reduced biosynthetic capacity and an anti-angiogenic state, supported by decreased NRP1, SPP1, and EFEMP1. In parallel, cholesterol homeostasis is increased with induction of detox/stress-related proteins (e.g., CYP1A1, CYP3A7, MT1E), and glucocorticoid receptor protein (NR3C1) is strongly downregulated, consistent with ligand-induced receptor feedback. Unexpectedly, IL6–JAK–STAT3 and broader inflammatory response pathways are positively enriched, and estrogen-response signatures (early and late) are also up, suggesting potential context-dependent adaptive signaling or nuclear receptor cross-talk. Additional stress/adaptation programs (hypoxia, EMT, myogenesis) are enriched, DKK1 is decreased, and MIXL1 is markedly induced; these observations are statistically supported but should be interpreted cautiously. Reactome/KEGG enrichments are not detected, so pathway specificity rests on Hallmark and GO categories; overall, the dataset supports GR engagement with strong anti-anabolic and anti-angiogenic features, plus several unexpected adaptive signaling programs that warrant orthogonal validation.

## Findings consistent with expected biology
- Broad suppression of rRNA processing/ribosome biogenesis, RNA splicing/mRNA processing, and cytoplasmic translation (strong negative GO BP enrichment).
- Downregulation of MYC target programs (Hallmark MYC_TARGETS_V2) consistent with reduced biosynthetic drive.
- Anti-angiogenic signature with decreased NRP1, SPP1, and EFEMP1 and negative enrichment of Hallmark ANGIOGENESIS.
- Upregulation of cholesterol/sterol handling and detox/stress proteins (e.g., CYP1A1, CYP3A7, MT1E) consistent with metabolic adaptation.
- Marked downregulation of NR3C1 protein, consistent with ligand-induced glucocorticoid receptor feedback.

## Unexpected but plausible findings
- Positive enrichment of IL6–JAK–STAT3 and INFLAMMATORY_RESPONSE pathways despite glucocorticoid exposure (supported by SERPINE1, JUNB, FSTL3).
- Enrichment of ESTROGEN_RESPONSE (early and late), suggestive of nuclear receptor cross-talk in this context (supported by TFAP2C, ITGB3BP, PAQR8), without direct ER markers in the top proteins.
- Stress/adaptation signatures (HYPOXIA, EMT, MYOGENESIS) positively enriched; EMT inference remains tentative due to limited canonical marker proteins.
- Strong downregulation of DKK1 accompanied by EMT pathway enrichment, potentially hinting at mesenchymal features (cautious interpretation).
- Marked induction of MIXL1, an atypical developmental transcription factor, as a single-protein outlier requiring confirmation.

## Follow-up hypotheses
### Hypothesis 1
- **Hypothesis:** GR activation reduces anabolic capacity by suppressing ribosome biogenesis/translation and MYC-associated programs.
- **Why interesting:** Provides a mechanistic basis for anti-proliferative or cytostatic effects of glucocorticoids in this system and explains broad biosynthetic downshifts.
- **Suggested validation:** Assess global protein synthesis (OPP-puromycin or SUnSET), rRNA transcription (5-EU incorporation), and MYC protein levels; check proliferation markers and cell-cycle readouts in a matched time course.
- **Priority:** high

### Hypothesis 2
- **Hypothesis:** Betamethasone induces an anti-angiogenic state.
- **Why interesting:** Anti-angiogenic effects may contribute to tissue remodeling or anti-tumor contexts; supported by decreased NRP1, SPP1, EFEMP1 and pathway enrichment.
- **Suggested validation:** Measure angiogenic factors (e.g., VEGF secretion) and endothelial tube formation or migration assays using conditioned media from treated cells.
- **Priority:** medium

### Hypothesis 3
- **Hypothesis:** The drug reprograms lipid/xenobiotic metabolism, increasing cholesterol homeostasis and detox capacity.
- **Why interesting:** Metabolic adaptation could influence steroid handling, membrane composition, and drug interactions.
- **Suggested validation:** Quantify cellular cholesterol and lipid efflux, test CYP1A1/3A activity assays, and perform time-course proteomics/transcriptomics for sterol and xenobiotic pathway components.
- **Priority:** medium

### Hypothesis 4
- **Hypothesis:** IL6–JAK–STAT3 signaling is paradoxically activated as a context-dependent adaptive response to glucocorticoid exposure.
- **Why interesting:** Explains positive enrichment of inflammatory/STAT3 signatures under a GR agonist and may impact therapeutic responses or resistance.
- **Suggested validation:** Measure phospho-STAT3 (Y705) and secreted IL6 by Western/ELISA; test reversibility with JAK inhibitors and evaluate time- and dose-dependence.
- **Priority:** high

### Hypothesis 5
- **Hypothesis:** Apparent estrogen-response enrichment reflects nuclear receptor cross-talk or off-target modulation in this cell context.
- **Why interesting:** Nuclear receptor interactions could modulate downstream transcriptional programs and influence combination therapies.
- **Suggested validation:** Use ER reporter assays and ER antagonists to test dependency; profile canonical ER target transcripts/proteins over a time course.
- **Priority:** medium

### Hypothesis 6
- **Hypothesis:** Ligand-induced down-modulation of NR3C1 underlies receptor-level feedback.
- **Why interesting:** Receptor abundance dynamics can shape sensitivity and duration of GR signaling.
- **Suggested validation:** Time-course quantification of NR3C1 protein and mRNA; test proteasome inhibition for rescue to evaluate degradation contribution.
- **Priority:** high

### Hypothesis 7
- **Hypothesis:** EMT/hypoxia signatures and DKK1 loss may indicate a shift toward mesenchymal-like features in a subset of cells.
- **Why interesting:** Phenotypic plasticity could alter adhesion, migration, and drug response.
- **Suggested validation:** Examine canonical EMT markers (e.g., E-cadherin, Vimentin) by immunoblot/immunofluorescence and perform migration/invasion assays; include single-cell analyses if possible.
- **Priority:** low

### Hypothesis 8
- **Hypothesis:** MIXL1 induction represents a cell-state outlier or rare program triggered by steroid exposure.
- **Why interesting:** If reproducible, may reveal unanticipated transcriptional reprogramming with implications for differentiation or stress states.
- **Suggested validation:** Confirm MIXL1 by targeted proteomics and qPCR; assess cell-to-cell heterogeneity (single-cell RNA/protein) and test dose/time dependence.
- **Priority:** low

## Confidence
medium

## Caution notes
- Pathway inferences are based on proteomic rank enrichment (Hallmark/GO); Reactome/KEGG were not significant, limiting pathway specificity.
- Reduced biosynthetic capacity is strongly supported, but explicit proliferation arrest is not demonstrated; cell-cycle hallmark changes were not highlighted.
- Activation of IL6–STAT3, estrogen-response, EMT, and hypoxia signatures should be validated with orthogonal markers (e.g., phospho-STAT3, ER reporters, canonical EMT markers).
- Attribution to GR engagement relies on the drug’s known target and NR3C1 downregulation; direct induction of canonical GR targets was not among the top proteins.
- The pro-inflammatory signature under a GR agonist is counterintuitive and may be cell-type, time- or dose-dependent; confirm with targeted assays.
- Single-protein outliers (e.g., MIXL1) should be interpreted cautiously until independently replicated.


---

# Betazole

# Drug summary: Betazole

## Executive summary
Betazole exposure produced a coherent proteomic signature dominated by strong suppression of intermediate filament organization (GO BP down; FDR=0), alongside extracellular matrix/adhesion remodeling characterized by loss of NPNT, PRG4, and CD164 with concurrent increases in CEMIP and CD9. Protein-level changes (ATF1↑, PDE4D↑, RIC8B↑) are consistent with engagement of histamine receptor–GPCR/Gs–cAMP signaling and feedback control, but this remains inferential in the absence of upregulated pathway enrichments or phospho/cAMP readouts. A prominent induction of TXNIP suggests a metabolic/redox stress component. Additional protein-level decreases in components of autophagy/lysosomal and glycosaminoglycan/glycosylation machinery (ATG14, HPS4, SPPL3, CHPF2, SLC9A8) point to altered proteostasis and trafficking, though these signals are mixed (e.g., HPS1↑) and should be interpreted cautiously. Modest reductions in cytokine receptor markers (IL4R, TNFRSF12A) hint at decreased inflammatory responsiveness but lack pathway corroboration. No upregulated Hallmark or GO BP pathways were detected; therefore, conclusions about upregulated programs rely on coordinated protein patterns and carry moderate confidence.

## Findings consistent with expected biology
- Protein-level indications of GPCR/Gs–cAMP engagement consistent with histamine receptor activation (ATF1↑, PDE4D↑ as feedback phosphodiesterase, RIC8B↑ supporting G-protein coupling).

## Unexpected but plausible findings
- Robust suppression of intermediate filament–based processes and organization (GO BP down; NES ~ -2.1; FDR=0), with keratin KRT79 among downregulated proteins.
- Extracellular matrix/adhesion remodeling: marked NPNT↓ (strongest downregulated protein) with PRG4↓ and CD164↓, together with CEMIP↑ (hyaluronan remodeling) and CD9↑.
- Strong TXNIP induction, indicating a metabolic/redox stress–related response in the absence of pathway-level enrichment.
- Mixed reductions in autophagy/lysosomal and glycosylation-related proteins (ATG14↓, HPS4↓, SPPL3↓, CHPF2↓, SLC9A8↓) suggesting perturbed proteostasis/trafficking, tempered by HPS1↑.
- Decreases in select cytokine/immune receptors (IL4R↓, TNFRSF12A↓, RC3H2↓) suggesting a possible dampening of inflammatory signaling, with mixed context (e.g., GRAP↑); interpretation is provisional.

## Follow-up hypotheses
### Hypothesis 1
- **Hypothesis:** Betazole activates histamine receptor–driven Gs–cAMP/PKA signaling that stabilizes ATF1 activity and induces PDE4D as feedback.
- **Why interesting:** Mechanistic confirmation links target engagement to the observed transcriptional/feedback proteomic changes and clarifies signaling mode.
- **Suggested validation:** Measure intracellular cAMP; assess PKA substrate and ATF1/CREB phosphorylation by immunoblot/PRM; test sensitivity to H2 receptor antagonists and PKA inhibition.
- **Priority:** high

### Hypothesis 2
- **Hypothesis:** Betazole suppresses intermediate filament programs, driving cytoskeletal reorganization that alters cell morphology and adhesion.
- **Why interesting:** This is the strongest pathway-level signal and may underlie downstream adhesion and motility phenotypes.
- **Suggested validation:** Immunofluorescence and immunoblot for keratins/intermediate filament proteins (e.g., KRTs); cell shape and motility assays; qPCR/protein quantification of IF genes.
- **Priority:** high

### Hypothesis 3
- **Hypothesis:** Betazole remodels the extracellular matrix, particularly hyaluronan turnover, via CEMIP induction and NPNT/PRG4 loss, altering cell–matrix interactions.
- **Why interesting:** Coordinated ECM changes can modulate adhesion and signaling, potentially impacting tissue remodeling and inflammatory responses.
- **Suggested validation:** Measure extracellular hyaluronan content/fragmentation and CEMIP activity; quantify NPNT/PRG4 secretion; adhesion assays; surface CD9 profiling by flow cytometry.
- **Priority:** high

### Hypothesis 4
- **Hypothesis:** TXNIP upregulation reflects activation of a metabolic/redox stress pathway that intersects with GPCR/cAMP signaling.
- **Why interesting:** TXNIP is a strong, unexpected signal that could influence glucose/redox homeostasis and downstream signaling.
- **Suggested validation:** Assess TXNIP mRNA/protein kinetics; measure ROS, NADPH/NADP+ ratios, and thioredoxin activity; evaluate glucose uptake and AMPK status; probe modulation by glucose availability.
- **Priority:** medium

### Hypothesis 5
- **Hypothesis:** Betazole perturbs vesicle trafficking/autophagy and glycosaminoglycan/glycosylation pathways, impacting secretory/lysosomal function.
- **Why interesting:** Coordinated decreases in ATG14, HPS4, SPPL3, CHPF2, and SLC9A8 suggest altered proteostasis with potential functional consequences.
- **Suggested validation:** Monitor LC3-II and p62 turnover, LAMP1/lysosomal markers; glycomics to profile chondroitin sulfate and global N-/O-glycan changes; test rescue with autophagy modulators; reconcile HPS1↑ vs HPS4↓.
- **Priority:** medium

### Hypothesis 6
- **Hypothesis:** Betazole reduces responsiveness to specific cytokines (e.g., IL-4, TWEAK) through receptor downregulation.
- **Why interesting:** If confirmed, this could indicate an anti-inflammatory tilt relevant to the immunology/inflammation pathway context.
- **Suggested validation:** Assess IL-4–STAT6 and TWEAK–NF-κB signaling after betazole treatment; quantify surface IL4R and TNFRSF12A by flow cytometry; evaluate cytokine-responsive gene induction.
- **Priority:** low

## Confidence
medium

## Caution notes
- No upregulated Hallmark or GO BP pathways were detected; inferences about upregulated programs (e.g., cAMP/PKA) are based on coordinated protein changes and need orthogonal confirmation.
- Cell type/model context is unspecified; interpretations involving tissue-restricted markers (e.g., keratins, BCAS1, PROM1) may not generalize.
- Autophagy/lysosome and glycosylation interpretations are mixed (e.g., HPS1↑ vs HPS4↓; SPPL3 decrease has context-dependent effects); avoid broad conclusions without functional assays.
- The suggested anti-inflammatory tilt rests on a small set of proteins and lacks pathway-level support; consider it provisional.
- Protein abundance does not directly report signaling activity; phosphoproteomics or biochemical assays are required to confirm pathway activation.
- GO-based suppression of intermediate filament organization is strong, but morphological/functional consequences require validation.
- ECM/hyaluronan remodeling inferences from CEMIP/NPNT/PRG4 changes should be corroborated with extracellular matrix measurements.


---

# Bicyclol

# Drug summary: Bicyclol

## Executive summary
Proteomics after Bicyclol exposure shows a hypoxia-like adaptive response (Hallmark HYPOXIA enrichment; NES 1.84, FDR ~0.028) with concomitant increases in stress-associated proteins, including JUNB, LOXL2, NRP1 and antioxidant/mitochondrial factors (PRXL2C, TOMM5, BOLA1). A pronounced induction of the xenobiotic enzyme CYP1A1 is observed, consistent with activation of a xenobiotic/oxidative stress program. The Hallmark KRAS_SIGNALING_DN set is negatively enriched (NES -1.75, FDR ~0.017), which can be compatible with a MAPK/AP-1 activation-like state, but this remains an inference without corroborating pathway or phospho evidence. Several components of membrane trafficking (SEC16B, PIP5K1C, TMEM208) and translation control/biogenesis (EEF1AKMT1, RSL24D1) are reduced, suggesting selective attenuation of secretory/translation modules under stress. Unexpectedly, the DNA replication kinase CDC7 is strongly upregulated, indicating possible replication stress or S-phase pressure at the protein level. Despite the drug’s annotation to autophagy, no significant autophagy-related pathway enrichment or coordinated changes in core autophagy machinery were detected in this dataset. Overall, the data support a generalized stress-adaptation signature with xenobiotic metabolism induction; more specific claims (e.g., MAPK/AP-1 or mTORC2 engagement) require orthogonal validation.

## Findings consistent with expected biology
- None listed.

## Unexpected but plausible findings
- Significant hypoxia-like enrichment with coordinated increases in stress-associated proteins (JUNB, LOXL2, NRP1, PRXL2C), indicating a generalized adaptive response.
- Robust CYP1A1 upregulation consistent with xenobiotic/oxidative stress activation, potentially impacting metabolic state or drug–drug interaction risk.
- Negative enrichment of Hallmark KRAS_SIGNALING_DN, suggestive of a MAPK/AP-1 activation-like state, but lacking direct pathway/phospho support (interpret with caution).
- Selective decreases in trafficking/translation-associated proteins (SEC16B, PIP5K1C, TMEM208; EEF1AKMT1, RSL24D1), indicating attenuation of secretory/translation components under stress.
- Marked CDC7 increase at the protein level, pointing to possible replication stress or checkpoint adaptation (single-protein observation).
- No detectable autophagy pathway enrichment despite prior annotation, suggesting autophagy engagement is not evident under the tested conditions.

## Follow-up hypotheses
### Hypothesis 1
- **Hypothesis:** Bicyclol induces a hypoxia-like (HIF-associated) adaptive program that contributes to stress tolerance.
- **Why interesting:** Explains the Hallmark HYPOXIA enrichment and elevation of hypoxia/stress-associated proteins; may intersect with antiviral or cytoprotective effects.
- **Suggested validation:** Measure HIF1α/HIF2α stabilization and canonical targets (e.g., VEGFA, CA9) by immunoblot/qPCR; assess oxygen consumption and glycolytic shift (Seahorse). Test sensitivity to HIF pathway inhibitors.
- **Priority:** high

### Hypothesis 2
- **Hypothesis:** CYP1A1 induction reflects activation of the AhR xenobiotic response by Bicyclol.
- **Why interesting:** Clarifies mechanism of xenobiotic metabolism induction and potential metabolic liabilities or interactions.
- **Suggested validation:** AhR reporter assay and nuclear translocation; confirm additional AhR targets (CYP1B1, NQO1) by qPCR/proteomics; test blockade with AhR antagonist (e.g., CH223191) for CYP1A1 reversal.
- **Priority:** high

### Hypothesis 3
- **Hypothesis:** A MAPK/AP-1 pathway contributes to the stress signature and JUNB upregulation.
- **Why interesting:** Provides a mechanistic link for KRAS_SIGNALING_DN negative enrichment and AP-1 component increase.
- **Suggested validation:** Assess p-ERK, p-cJUN, and broader AP-1 components (JUN/FOS family) by immunoblot; test whether MEK inhibition (e.g., trametinib) attenuates JUNB induction.
- **Priority:** medium

### Hypothesis 4
- **Hypothesis:** Cells reduce secretory throughput and translation under Bicyclol via integrated stress response (ISR) signaling.
- **Why interesting:** Accounts for reduced SEC16B/PIP5K1C/TMEM208 and translation modifiers and links stress adaptation to proteostasis.
- **Suggested validation:** Measure eIF2α phosphorylation and ATF4 induction; perform puromycin incorporation (SUnSET) for global translation; test ISR inhibition (ISRIB) for rescue.
- **Priority:** medium

### Hypothesis 5
- **Hypothesis:** CDC7 upregulation indicates replication stress and altered S-phase control.
- **Why interesting:** May unveil a liability to replication stress-targeting agents or checkpoint inhibitors.
- **Suggested validation:** Quantify EdU incorporation, phospho-Chk1 (Ser345) and γH2AX; test sensitivity to CDC7 inhibition (e.g., XL413) and replication stress modulators.
- **Priority:** medium

### Hypothesis 6
- **Hypothesis:** mTORC2 signaling may be modestly engaged (PRR5 increase) to support survival under stress.
- **Why interesting:** Could fine-tune survival pathways during hypoxia-like adaptation.
- **Suggested validation:** Assess AKT Ser473 phosphorylation and RICTOR/PRR5 levels; compare effects of rapamycin vs. Torin-1; look for coordinated changes in other mTORC2 components.
- **Priority:** low

## Confidence
medium

## Caution notes
- Interpret KRAS/MAPK activation cautiously; the inference is based on KRAS_SIGNALING_DN negative enrichment without corroborating pathway or phosphoproteomic evidence.
- AP-1 involvement is suggested primarily by JUNB; broader JUN/FOS family support is not shown.
- mTORC2 involvement (PRR5 increase) is speculative and not supported by coordinated pathway changes.
- ECM/adhesion remodeling suggestions (from LOXL2, NRP1) are limited by the small protein set.
- Mitochondrial/oxidative stress adaptation is indicated by a few proteins (TOMM5, BOLA1, PRXL2C) without pathway enrichment; treat as tentative.
- Autophagy engagement is not supported by pathway analysis in this dataset despite prior annotation; avoid inferring autophagy activation here.
- Hallmark HYPOXIA may reflect generalized cellular stress or metabolic adaptation rather than true hypoxia.
- Several observations (e.g., CDC7, PRR5 changes) are single-protein findings and should be validated.
- Pathway evidence is limited largely to Hallmark sets; GO/Reactome sections were empty, constraining mechanistic resolution.


---

# Binimetinib

# Drug summary: Binimetinib

## Executive summary
Binimetinib proteomic profiling shows a coherent MEK-pathway attenuation signature with strong downstream proliferative arrest. Reciprocal KRAS hallmark behavior, marked MYC protein decrease, and broad suppression of E2F/G2M/mitotic spindle programs alongside reduced CDC6/CDC7/CDC25B/SASS6/SKA1 support a robust anti-proliferative effect. In parallel, there is pronounced enrichment of oxidative phosphorylation and fatty-acid/peroxisomal catabolism with increases in lipid handling and mitochondrial proteins, consistent with metabolic adaptation under MAPK blockade. Pathway-level decreases in inflammatory and angiogenesis hallmarks (with reduced CCL20 and PLAUR) are observed, although EFNA1 is up, indicating gene-level heterogeneity. Stress-response features (ROS pathway up, GPX3, DDIT4/REDD1, TP53INP2 induction) suggest energy/oxidative stress; autophagy engagement is plausible but unproven. Although Reactome and KEGG sections are empty, the hallmark/GO evidence is strong and aligns with MEK inhibition biology.

## Findings consistent with expected biology
- Attenuation of RAS/MAPK proliferative output with reciprocal KRAS hallmarks (KRAS_SIGNALING_UP down; KRAS_SIGNALING_DN up) and marked MYC protein reduction.
- Broad cell-cycle arrest with coordinated downregulation of E2F targets, G2/M checkpoint, mitotic spindle/chromosome segregation processes and decreased CDC6/CDC7/CDC25B/SASS6/SKA1.

## Unexpected but plausible findings
- Strong induction of oxidative phosphorylation and fatty-acid/peroxisomal β-oxidation programs with increased lipid transport/oxidation and mitochondrial import proteins (e.g., SLC27A3, SLC25A42, PNPLA8, TOMM40L, ABHD2, APOA5, CYP4F2).
- Upregulation of bile-acid/xenobiotic metabolism pathways, atypical in this context, supported by CYP4F2 and APOA5 increases.
- Suppression of inflammatory and angiogenesis hallmarks with decreases in CCL20 and PLAUR, but EFNA1 upregulation highlights pathway-level conclusions may mask gene-level heterogeneity.
- Stress-response engagement (ROS pathway up; GPX3, DDIT4/REDD1, TP53INP2 induction) consistent with energy/oxidative stress; potential autophagy activation remains speculative without direct markers.
- PRIMPOL upregulation despite cell-cycle suppression, suggestive of a DNA damage/replication stress tolerance response (uncertain).

## Follow-up hypotheses
### Hypothesis 1
- **Hypothesis:** Binimetinib-treated cells adopt a fatty-acid–supported, OXPHOS-dependent metabolic state as an adaptive survival mechanism.
- **Why interesting:** If cells rely on FAO/OXPHOS under MEK inhibition, co-targeting mitochondrial respiration or fatty-acid entry could enhance efficacy.
- **Suggested validation:** Measure OCR/ECAR and palmitate oxidation; test sensitivity to FAO/mitochondrial inhibitors (e.g., etomoxir, CPT1 knockdown, complex I inhibitors) and assess synergy with binimetinib.
- **Priority:** high

### Hypothesis 2
- **Hypothesis:** PPARα-driven transcription contributes to the induction of fatty-acid/peroxisomal catabolism after MEK inhibition.
- **Why interesting:** Identifying a PPARα axis would provide a mechanistic handle for combination strategies and biomarker development.
- **Suggested validation:** Assess PPARα target gene/protein induction; test effects of PPARα agonists/antagonists (e.g., GW6471) on FAO/OXPHOS signatures and cell viability under binimetinib.
- **Priority:** medium

### Hypothesis 3
- **Hypothesis:** Autophagy is engaged downstream of stress signals (DDIT4/TP53INP2) and supports survival under MEK inhibition.
- **Why interesting:** Autophagy dependency could create a therapeutic vulnerability to autophagy inhibitors in combination with binimetinib.
- **Suggested validation:** Monitor LC3-II processing (± bafilomycin), p62 turnover, and autophagosome formation; test functional impact of ATG gene knockdown or pharmacological autophagy inhibition on binimetinib response.
- **Priority:** medium

### Hypothesis 4
- **Hypothesis:** PRIMPOL upregulation reflects replication stress tolerance, and PRIMPOL is required for survival under MEK pathway suppression.
- **Why interesting:** Targeting replication stress adaptation may sensitize cells to MEK inhibition.
- **Suggested validation:** Quantify γH2AX/RPA foci and PRIMPOL expression; evaluate effects of PRIMPOL knockdown on DNA damage markers and viability in the presence of binimetinib.
- **Priority:** low

## Confidence
high

## Caution notes
- Direct MEK/ERK pathway inhibition (e.g., pERK) was not measured; MAPK attenuation is inferred from hallmarks and downstream protein changes.
- Conclusions derive from a single proteomics layer; no phosphoproteomics, transcriptomics, or metabolomics are available to corroborate pathway activity or metabolic flux.
- Evidence for autophagy is indirect (DDIT4, TP53INP2) without canonical markers or pathway enrichment; interpret autophagy conclusions cautiously.
- Increased OXPHOS and FAO do not establish reduced glycolysis; glycolytic markers/flux were not assessed.
- Angiogenesis and invasion-related interpretations should be cautious given gene-level heterogeneity (e.g., EFNA1 up despite pathway down).
- Several upregulated proteins are secreted (e.g., APOA5, INHBE, GPX3); extracellular/medium contributions and cell-type specificity should be considered.
- Temporal and dose context are unspecified; distinguishing acute pharmacodynamics from adaptive responses requires time-course studies.
- Empty Reactome/KEGG sections do not negate the strong Hallmark/GO evidence but limit pathway-level granularity.
- Cell line or tissue context is not specified, which may influence the relevance of bile-acid/xenobiotic and lipid metabolism changes.


---

# Bromhexine (hydrochloride)

# Drug summary: Bromhexine (hydrochloride)

## Executive summary
Bromhexine (hydrochloride) elicited a dominant inflammatory and stress-associated proteomic phenotype. Hallmark enrichments indicated strong inflammatory response and IL6–JAK–STAT3 programs, along with TNFα–NFκB, accompanied by a hypoxia-like signature and an EMT-like/ECM-remodeling program. Concordant protein increases included LOXL2, MMP14, CEMIP, CD109, TYK2, and actin-associated components (TPM2, AFAP1). Cytoskeletal reorganization was further supported by a robust down-regulation of intermediate filament organization at the pathway level. Secondary, more tentative signals included a possible tilt in cholesterol homeostasis (HMGCR down, NPC1L1 up) and isolated decreases in components related to protein/mitochondrial import (SEC61G, TOMM7, SLC25A39). Notably, despite the drug’s autophagy-related annotation, no autophagy pathway enrichment was observed. These findings should be interpreted conservatively as abundance-based signatures rather than direct pathway activation.

## Findings consistent with expected biology
- None listed.

## Unexpected but plausible findings
- Strong enrichment of inflammatory response, IL6–JAK–STAT3, and TNFα–NFκB programs with protein-level alignment (e.g., TYK2 up), indicating a pronounced pro-inflammatory state.
- Hypoxia-like and EMT-like/ECM-remodeling signatures supported by marked LOXL2 upregulation and increases in MMP14, CEMIP, and CD109.
- Cytoskeletal reprogramming characterized by increased actin-associated components (TPM2, AFAP1) and a significant pathway-level decrease in intermediate filament organization.
- Upregulation of the stress/immune ligand MICA, consistent with heightened cellular stress signaling and potential immune recognition.
- Tentative cholesterol homeostasis shift with HMGCR down and NPC1L1 up, suggesting possible compensation toward uptake over synthesis (no lipid pathway enrichment).
- Isolated decreases in proteins involved in translocation/mitochondrial function (SEC61G, TOMM7, SLC25A39), suggestive but not conclusive for reduced import capacity.

## Follow-up hypotheses
### Hypothesis 1
- **Hypothesis:** Bromhexine induces a pro-inflammatory, stress-associated program that coincides with an EMT-like/ECM-remodeling response.
- **Why interesting:** This may impact epithelial barrier properties and tissue remodeling, with potential relevance to airway biology and infection contexts.
- **Suggested validation:** Quantify secreted cytokines (e.g., IL-6, TNFα), assess STAT3 and NF-κB phosphorylation/nuclear localization, evaluate EMT markers (E-cadherin, vimentin) and ECM remodeling (LOXL2 and MMP14 activity).
- **Priority:** high

### Hypothesis 2
- **Hypothesis:** A hypoxia-like response contributes to the LOXL2/MMP14/ECM signature, potentially via HIF stabilization or metabolic stress.
- **Why interesting:** Links stress metabolism with ECM remodeling and may influence host–pathogen interactions in low-oxygen or inflamed tissues.
- **Suggested validation:** Measure HIF-1α/2α stabilization and canonical targets (e.g., GLUT1, CA9), perform Seahorse assays for bioenergetics, and test HIF pathway inhibition for attenuation of LOXL2/MMP14 induction.
- **Priority:** medium

### Hypothesis 3
- **Hypothesis:** Bromhexine shifts cholesterol homeostasis toward uptake over synthesis.
- **Why interesting:** Cholesterol balance can affect membrane composition and viral entry/replication dynamics.
- **Suggested validation:** Assess SREBP2 processing, cellular cholesterol content, LDL/NPC1L1-mediated uptake, and pharmacologic modulation (e.g., statin or ezetimibe co-treatment) to test functional consequences.
- **Priority:** medium

### Hypothesis 4
- **Hypothesis:** Cellular immunogenicity is enhanced via upregulation of NKG2D ligands (e.g., MICA).
- **Why interesting:** Could alter NK cell recognition and cytotoxicity toward treated cells.
- **Suggested validation:** Flow cytometry for surface MICA/B, NK-cell co-culture cytotoxicity assays, and NKG2D blocking to assess dependence.
- **Priority:** medium

### Hypothesis 5
- **Hypothesis:** Proteostasis and organelle import are subtly perturbed, contributing to stress signatures.
- **Why interesting:** Even mild deficits in ER translocation or mitochondrial import could amplify stress/inflammatory responses.
- **Suggested validation:** Reporter assays for Sec61-dependent translocation and mitochondrial import, markers of UPR/ISR (BiP, ATF4, CHOP), and broader profiling of import machinery.
- **Priority:** low

## Confidence
medium

## Caution notes
- Abundance-based proteomics does not directly measure pathway activation; avoid causal language about signaling without phospho- or secretion data.
- EMT interpretation should be framed as EMT-like/ECM remodeling unless epithelial identity and canonical EMT marker shifts are demonstrated.
- No autophagy-related pathway enrichment was detected despite the drug’s annotation; absence of evidence here does not exclude context-dependent autophagy effects.
- Conclusions about cholesterol metabolism and protein/mitochondrial import are based on a few proteins and lack pathway-level enrichment; treat as preliminary.
- Pathway calls rely mainly on Hallmark enrichments and one GO BP down term; the absence of Reactome/KEGG hits warrants conservative interpretation.
- Inflammatory and hypoxia-like signatures may reflect generalized cellular stress; orthogonal assays (viability, HIF targets, secreted cytokines) are recommended.
- Cell type and context are unspecified, limiting extrapolation to specific tissues or disease settings.


---

# Budesonide

# Drug summary: Budesonide

## Executive summary
Budesonide exposure yields a proteomic profile dominated by strong suppression of RNA processing and protein synthesis machinery (RNA splicing, rRNA processing/ribosome biogenesis, cytoplasmic translation), with coherent GO evidence (multiple terms with NES ~ -1.8 to -2.2, low FDR) and protein-level concordance (e.g., SNRNP48, RSL24D1 down). Concurrently, there is induction of xenobiotic/steroid metabolism and transport (CYP3A7, CYP24A1, SLC16A6, SLCO4A1), redox/stress proteins (TXNIP, MT1E), and select membrane/ion-transport components (CAVIN2, FXYD1). Immediate-early/inflammatory-associated proteins (IER3, TRAF3) are reduced, and NR3C1 (GR) protein is modestly decreased, consistent with adaptation to glucocorticoid exposure. A single GO signal suggests increased phospholipid catabolism, but this rests on one enriched term without clear phospholipase drivers among top proteins and should be considered tentative. No Hallmark/Reactome/KEGG enrichments were detected, so interpretations rely primarily on GO BP and protein-level changes. Overall, the data align with a glucocorticoid-responsive state that dampens biosynthetic capacity and enhances metabolic/transport adaptation, with additional context-dependent features (e.g., CYP24A1, HLA-B) that warrant follow-up.

## Findings consistent with expected biology
- Broad downregulation of RNA processing and translation programs (spliceosome, rRNA processing/ribosome biogenesis, cytoplasmic translation) with strong GO support and protein-level concordance.
- Induction of xenobiotic/steroid metabolism and solute transporters (e.g., CYP3A7, SLC16A6, SLCO4A1), consistent with glucocorticoid-driven metabolic/transport adaptation.
- Upregulation of redox/stress response proteins (TXNIP, MT1E).
- Reduction of immediate-early/inflammatory-associated proteins (IER3, TRAF3), consistent with anti-inflammatory actions.
- Modest decrease in NR3C1 (glucocorticoid receptor) protein, compatible with adaptation to glucocorticoid exposure (interpret with caution).

## Unexpected but plausible findings
- Strong induction of CYP24A1 (vitamin D catabolic enzyme), suggesting possible nuclear receptor crosstalk with vitamin D signaling under budesonide treatment (single-protein evidence).
- Upregulation of HLA-B despite the drug’s immunosuppressive context (single-protein evidence, likely context-dependent).
- Elevated CAVIN2 and FXYD1 indicating potential membrane/caveolae and ion-transport remodeling; pathway-level corroboration is lacking.
- GO signal for phospholipid catabolic process (single enriched term) without obvious phospholipase drivers among top proteins; provisional.

## Follow-up hypotheses
### Hypothesis 1
- **Hypothesis:** The observed repression of RNA processing/translation and induction of metabolic/transport proteins is GR-dependent on-target biology.
- **Why interesting:** Confirms mechanism of action and addresses the absence of canonical GR targets among top changes.
- **Suggested validation:** Test GR dependence using a GR antagonist (e.g., mifepristone) or GR knockdown; assess canonical GR-inducible markers (FKBP5, TSC22D3/GILZ, DUSP1, SGK1, ANXA1) by qPCR/proteomics; perform time course to capture early GR targets.
- **Priority:** high

### Hypothesis 2
- **Hypothesis:** Budesonide induces CYP24A1 via vitamin D receptor (VDR) pathway crosstalk.
- **Why interesting:** Explains robust CYP24A1 upregulation and implicates nuclear receptor interplay affecting steroid/vitamin D metabolism.
- **Suggested validation:** Measure CYP24A1 mRNA/protein kinetics; use VDR knockdown/antagonists or VDR reporter assays; quantify 1,25(OH)2D3/24,25(OH)2D3 levels; test whether CYP24A1 induction persists with GR blockade.
- **Priority:** medium

### Hypothesis 3
- **Hypothesis:** TFEB upregulation reflects activation of lysosomal/clearance programs that support lipid/membrane remodeling.
- **Why interesting:** Provides a mechanistic link between TFEB/CAVIN2 changes and the tentative phospholipid catabolic signal.
- **Suggested validation:** Assess TFEB nuclear localization; profile CLEAR network gene expression; measure lysosomal markers (LAMP1, cathepsins), autophagic flux, and lipid turnover; examine caveolae components (CAV1/CAVIN1) and membrane ultrastructure.
- **Priority:** medium

### Hypothesis 4
- **Hypothesis:** Budesonide increases MHCI antigen presentation in this context, as suggested by HLA-B upregulation.
- **Why interesting:** Contrasts with an immunosuppressive expectation and may be cell-context dependent.
- **Suggested validation:** Measure surface HLA-A/B/C by flow cytometry; evaluate antigen-presentation machinery transcripts/proteins; assess interferon-response status and test JAK/STAT dependency.
- **Priority:** low

## Confidence
medium

## Caution notes
- Upregulated phospholipid catabolic process is supported by a single GO term; identify leading-edge genes and confirm induction of direct lipid-catabolic enzymes before asserting pathway activation.
- Mechanistic attribution to GR is plausible given the drug but should be framed as inferred; canonical GR targets are not prominent among the highlighted proteins.
- TFEB/membrane remodeling inference is tentative without lysosome/autophagy pathway enrichment.
- NR3C1 protein decrease should not be overinterpreted as feedback without time-course or transcriptional evidence; it may reflect broader translational downregulation.
- Claims about broad immunosuppression require pathway-level evidence (e.g., NF-κB/AP-1 target suppression); current support rests on a small number of proteins.
- Pathway enrichment relies primarily on GO BP; the absence of Hallmark/KEGG/Reactome signals limits cross-ontology corroboration.
- Cellular context and timepoint are unspecified; several proteins (e.g., CYP3A7, PGC, HLA-B) are highly context dependent.


---

# Butylphthalide

# Drug summary: Butylphthalide

## Executive summary
Proteomic profiling of butylphthalide treatment reveals a robust hypoxia-like program with a coordinated downshift of RNA splicing/processing and ribosome biogenesis, consistent with a stress-adaptive reduction in biosynthetic load. This is driven by strong Hallmark HYPOXIA enrichment (NES ~2.20, FDR 0) and multiple GO biological processes down for RNA metabolism and ribosome biogenesis (NES ~ -2.10 to -1.87). Protein-level changes (e.g., COQ4 up) are directionally consistent with adaptation but are insufficient to infer specific metabolic rewiring. An EMT-regulatory gene set is significantly enriched (NES ~2.11, FDR 0) alongside increases in adhesion/signaling proteins (CD9, TSPAN13, EFNA1, DLK1), suggesting altered adhesion/cell-state regulation rather than a full EMT program given unknown cellular context. A microtubule-related shift is suggested by down-enrichment of MICROTUBULE_DEPOLYMERIZATION (NES ~ -1.88, FDR ~0.04) and marked STMN3 downregulation, consistent with hypothetical microtubule stabilization. Upregulation of several membrane/organelle remodeling components (e.g., ATL3, RNF167, HPS1, DDHD1, PIGN, SMPDL3A) hints at trafficking/ER–endolysosomal remodeling, but pathway-level corroboration is lacking and this should be treated as tentative. Notable individual decreases (e.g., IGFBP2, CYP1A1, HDAC8) are observed without coherent pathway context. Overall, the most supported conclusions are activation of a hypoxia-like stress response and suppression of RNA/ribosome biogenesis, with medium confidence; cytoskeletal and membrane remodeling signals are hypothesis-generating.

## Findings consistent with expected biology
- Robust hypoxia-like program activation (Hallmark HYPOXIA up; NES ~2.20, FDR 0), consistent with a stress response
- Coordinated suppression of RNA splicing/processing and ribosome biogenesis (multiple GO BP terms down; NES ~ -2.10 to -1.87), indicating biosynthetic downscaling

## Unexpected but plausible findings
- Significant enrichment for regulation of EMT with increased adhesion/signaling proteins (e.g., CD9, TSPAN13, EFNA1, DLK1), suggesting altered cell-state/adhesion rather than a true EMT program
- Evidence consistent with reduced microtubule depolymerization and possible microtubule stabilization (MICROTUBULE_DEPOLYMERIZATION down; STMN3 markedly down), requiring orthogonal confirmation
- Tentative indication of membrane/organelle remodeling (elevated ATL3, RNF167, HPS1, DDHD1, PIGN, SMPDL3A) without pathway-level enrichment

## Follow-up hypotheses
### Hypothesis 1
- **Hypothesis:** Butylphthalide induces a hypoxia-like (potential HIF-driven) stress program that downscales RNA splicing and ribosome biogenesis.
- **Why interesting:** This integrates the strongest pathway signals and provides a unifying explanation for the biosynthetic downshift, aligning with stress adaptation.
- **Suggested validation:** Assess HIF pathway activation (HIF-1α protein stabilization, target gene readouts such as CA9/VEGFA), pimonidazole adducts, and metabolic flux; quantify global translation (puromycin/SUnSET) and rRNA biogenesis (pre-rRNA processing assays).
- **Priority:** high

### Hypothesis 2
- **Hypothesis:** Butylphthalide promotes microtubule stabilization via suppression of stathmin-3 and related depolymerization machinery.
- **Why interesting:** STMN3 downregulation with enrichment against microtubule depolymerization suggests cytoskeletal remodeling that could affect trafficking and cell morphology.
- **Suggested validation:** Tubulin polymerization assays (soluble vs polymerized tubulin fractionation), live-cell EB3 comet imaging, nocodazole challenge, and rescue/overexpression of STMN3 to test reversibility.
- **Priority:** medium

### Hypothesis 3
- **Hypothesis:** Butylphthalide remodels cell adhesion/signaling networks via tetraspanin and ephrin axis changes, altering adhesion without inducing full EMT.
- **Why interesting:** EMT-regulatory enrichment with increased CD9/TSPAN13/EFNA1 suggests adhesion and signaling reprogramming relevant to cell–cell/ECM interactions.
- **Suggested validation:** Measure adhesion/migration (ECM attachment, wound healing), focal adhesion markers (pFAK, vinculin), and EMT marker panel (E-cadherin, vimentin, ZEB1) to distinguish adhesion rewiring from EMT.
- **Priority:** medium

## Confidence
medium

## Caution notes
- Cellular context is unspecified; interpret EMT-related enrichment as altered adhesion/cytoskeletal regulation rather than a bona fide EMT without epithelial context and canonical marker changes.
- Proteomic abundance does not equate to pathway activity or subcellular localization; confirm hypoxia/HIF activation and microtubule dynamics with orthogonal assays.
- The microtubule depolymerization signal is near the FDR threshold and may be driven by few proteins; treat microtubule stabilization as hypothesis-generating.
- Membrane/organelle remodeling inferences lack Reactome/KEGG support; consider targeted pathway analyses or imaging before mechanistic claims.
- Gene set enrichment is based on a limited set of significant proteins (~50); provide leading-edge sizes/overlaps to gauge robustness.
- The absence of significant 'hallmark_down' sets is not a quality issue; conclusions should rely on present, statistically supported signals.


---

# Cabazitaxel

# Drug summary: Cabazitaxel

## Executive summary
Cabazitaxel exposure yields a coherent anti-proliferative proteomic signature characterized by strong suppression of E2F/G2–M programs and DNA replication/repair modules, alongside induction of a p53-consistent checkpoint (CDKN1A/p21, CDKN1B/p27 up) and an apoptosis hallmark. Stress/inflammatory responses are prominent (TNF–NFκB, IL6–STAT3, hypoxia) with increased ROS-handling proteins (e.g., TXNIP, GPX3). Epithelial/cytoskeletal remodeling is evident (keratinocyte/epidermal differentiation, intermediate filament organization up; ACTG1 down), and secretory/wound-associated programs (complement/coagulation; TIMP1, F5 up) are enriched. An EMT hallmark co-occurs with epithelial junction/differentiation markers (DSG3 up), indicating mixed epithelial remodeling rather than a full EMT transition. Calcium-responsive signaling is enriched with TRPC5 up, but functional channel activation is not established. Autophagy-specific enrichment is not detected. Direct microtubule engagement is inferred from the known MOA and the cytostatic pattern rather than pathway-level enrichments. Overall, the data support cell-cycle arrest with stress and secretory responses; some remodeling and calcium-associated features warrant targeted follow-up.

## Findings consistent with expected biology
- Robust suppression of proliferative programs: E2F targets and G2/M checkpoint down with broad decreases in DNA replication/repair regulators (e.g., CDC7, CDC25B, MCM10, RAD54B, EXO1, CLSPN).
- Checkpoint engagement consistent with a p53–p21/p27 cytostatic response and an apoptosis hallmark (TP53, CDKN1A, CDKN1B up).
- Prominent stress/inflammatory signaling (TNF–NFκB, IL6–STAT3, hypoxia) and increased ROS-related metabolism (TXNIP, GPX3 up).
- Cytoskeletal/epithelial remodeling consistent with microtubule-targeting stress: intermediate filament organization and keratinocyte/epidermal differentiation up; ACTG1 down.
- Upregulation of complement/coagulation and wound-response programs (e.g., TIMP1, F5), suggesting a secretory stress milieu.

## Unexpected but plausible findings
- EMT hallmark enrichment coincident with epithelial differentiation/desmosomal markers (DSG3 up), indicating mixed epithelial remodeling rather than a classical EMT shift.
- Strong downregulation of homologous recombination and interstrand crosslink repair gene sets accompanying replication shutdown; likely coupled to proliferation exit but could reduce repair capacity if sustained.
- Enrichment of calcium-responsive signaling with TRPC5 up; functional calcium channel engagement remains uncertain.

## Follow-up hypotheses
### Hypothesis 1
- **Hypothesis:** Cabazitaxel induces a p53–p21/p27–mediated checkpoint leading to cell-cycle arrest and apoptosis in this model.
- **Why interesting:** Aligns with the dominant proteomic and pathway signals and explains the broad suppression of E2F/G2–M and DNA replication modules.
- **Suggested validation:** Time-course immunoblot for p53 (total and Ser15/Ser46), p21, p27; cell-cycle profiling by flow cytometry (EdU/PI or DAPI); Annexin V/cleaved caspase-3/PARP assays to distinguish cytostasis versus apoptosis.
- **Priority:** high

### Hypothesis 2
- **Hypothesis:** Homologous recombination capacity is functionally reduced after treatment, primarily secondary to S-phase depletion but potentially sensitizing to PARP inhibitors if HR suppression persists.
- **Why interesting:** Could inform rational combinations; HR pathway markers are down and HR gene sets are suppressed.
- **Suggested validation:** RAD51 foci formation after ionizing radiation with gating for S/G2; DR-GFP HR reporter in synchronized cells; PARP inhibitor combination studies with cell-cycle control (e.g., partial synchronization or short pulses).
- **Priority:** medium

### Hypothesis 3
- **Hypothesis:** Cabazitaxel triggers NF-κB/IL6–STAT3–driven inflammatory secretome that modulates survival and microenvironmental signaling.
- **Why interesting:** Inflammatory hallmarks are among the strongest enrichments and may influence adaptive responses.
- **Suggested validation:** Assess NF-κB p65 nuclear translocation and STAT3 Tyr705 phosphorylation; measure secreted cytokines (e.g., IL6, IL8) by ELISA/Luminex; test whether NF-κB or STAT3 pathway inhibitors enhance growth inhibition.
- **Priority:** medium

### Hypothesis 4
- **Hypothesis:** Treatment promotes a mixed epithelial remodeling state with strengthened desmosomal adhesion rather than a full EMT.
- **Why interesting:** EMT hallmark co-occurs with DSG3 up and epidermal differentiation, suggesting altered adhesion/migration states.
- **Suggested validation:** Immunofluorescence for DSG3, E-cadherin, vimentin; cell–cell adhesion and migration assays (wound-healing/Transwell); transcript profiling of EMT and differentiation markers.
- **Priority:** medium

### Hypothesis 5
- **Hypothesis:** TRPC5-associated calcium signaling contributes to stress adaptation under Cabazitaxel.
- **Why interesting:** Response to calcium ion is enriched and TRPC5 is upregulated; if functional, calcium signaling could be targetable.
- **Suggested validation:** Live-cell Ca2+ imaging with TRPC5 agonists/antagonists; TRPC5 knockdown or pharmacologic inhibition to test effects on viability and stress markers under treatment.
- **Priority:** low

## Confidence
high

## Caution notes
- Proteomic abundance does not directly prove pathway activation; p53 pathway engagement is inferred and should be confirmed with post-translational readouts and functional assays.
- No clear autophagy-related enrichment is detected despite Cabazitaxel’s association with autophagy in some contexts; avoid implying autophagy engagement here.
- Direct microtubule/tubulin engagement is not reflected by specific pathway enrichments (e.g., Hallmark Mitotic Spindle not observed); the link to microtubule disruption is mechanistic context rather than proteomic proof.
- EMT hallmark can capture wound/ECM/inflammatory programs; DSG3 up argues against a full EMT. Interpret EMT-related signals as mixed epithelial remodeling.
- Suppression of homologous recombination and crosslink repair likely tracks with replication/S-phase exit; claims of repair-specific inhibition should be tempered and functionally tested.
- Evidence for calcium channel activity is limited; TRPC5 up and a broad ‘response to calcium ion’ term do not establish functional engagement.
- Lineage/context dependence: keratinocyte/epidermal differentiation and DSG3 up suggest epithelial context; results may not generalize to other cell types.
- Single time-point proteomics limits inference on dynamics and causality; distinguish cytostatic versus cytotoxic outcomes with time-course and orthogonal assays.
- Reactome/KEGG sections are empty; conclusions rely on Hallmark and GO enrichments.


---

# Camphor

# Drug summary: Camphor

## Executive summary
Camphor exposure elicits a robust pro-inflammatory and hypoxia-like proteomic program (Hallmark IL6/JAK/STAT3, TNFA/NFKB, HYPOXIA) accompanied by strong repression of RNA maturation machinery, including spliceosome function, rRNA processing, and ribosome biogenesis. These coherent pathway-level shifts support a stress-adaptive, growth-attenuated state. A secondary pattern suggests membrane/mechanosensory remodeling, with increased PIEZO2 and vesicle/tetraspanin markers (CD9, PROM1, SYNGR3), though this lacks pathway enrichment and should be considered tentative. Single-protein changes hint at additional biology (e.g., LIG4 decrease suggesting possible NHEJ vulnerability; SHQ1 increase amid global RNA-processing downregulation), but these require further validation. Overall, the strongest, actionable signal is inflammatory/hypoxia activation coupled to suppression of RNA and ribosome biogenesis.

## Findings consistent with expected biology
- Activation of IL6/JAK/STAT3 signaling (Hallmark NES ~2.16; FDR ~0)
- Activation of hypoxia-like program (Hallmark NES ~1.88; FDR ~0.0025)
- Activation of TNFA signaling via NFkB (Hallmark NES ~1.65; FDR ~0.041)
- Global repression of RNA processing modules, including RNA splicing, rRNA processing, and ribosome biogenesis (multiple GO BP sets, NES approximately -2.30 to -2.02; FDR ≤ 0.01)

## Unexpected but plausible findings
- Induction of the mechanosensitive channel PIEZO2 (~+2.05 log2FC) with concurrent upregulation of vesicle/membrane-associated proteins (CD9, PROM1, SYNGR3, ADAM15), suggesting membrane/mechanosensory remodeling (pathway corroboration lacking).
- Upregulation of the snoRNP assembly factor SHQ1 (~+1.07 log2FC) despite broad suppression of RNA/rRNA processing pathways, consistent with a possible compensatory RNP assembly response (speculative).
- Downregulation of DNA Ligase IV (LIG4; ~-1.86 log2FC) potentially indicating reduced non-homologous end joining capacity; broader DNA-repair suppression is not established.

## Follow-up hypotheses
### Hypothesis 1
- **Hypothesis:** Camphor triggers STAT3/NFkB- and hypoxia-associated signaling that drives a stress-adaptive reduction in RNA maturation and ribosome biogenesis.
- **Why interesting:** Unifies the strongest Hallmark and GO signatures and may explain growth attenuation; provides a mechanistic axis for downstream phenotypes relevant to inflammation and cytostasis.
- **Suggested validation:** Measure STAT3 phosphorylation and NFkB nuclear translocation; assess HIF1A stabilization; quantify nascent rRNA synthesis (47S pre-rRNA, EU/BrU incorporation) and nucleolar markers; evaluate global splicing changes by RNA-seq; test dependence on Ca2+ influx/TRP-channel blockade to probe upstream coupling.
- **Priority:** high

### Hypothesis 2
- **Hypothesis:** Camphor promotes membrane/mechanosensory remodeling, partly via PIEZO2 induction, with increased vesicle trafficking/extracellular vesicle markers.
- **Why interesting:** Links observed protein changes to the drug’s ion-channel/membrane activity profile and could affect neuronal signaling or vesicle-mediated communication.
- **Suggested validation:** Confirm PIEZO2 upregulation by immunoblot/RT-qPCR; assess mechanosensitive currents (patch clamp) and Ca2+ responses; quantify extracellular vesicles (NTA/flow) and tetraspanin markers (CD9); perturb PIEZO2 (siRNA/CRISPR) and/or use TRP-channel antagonists to test dependence.
- **Priority:** medium

### Hypothesis 3
- **Hypothesis:** Camphor reduces non-homologous end joining efficiency via LIG4 downregulation, creating a DNA-repair vulnerability.
- **Why interesting:** If confirmed, this could inform combination strategies with DNA-damaging agents or guide safety assessments.
- **Suggested validation:** Use NHEJ reporter assays; measure γH2AX/53BP1 foci kinetics after irradiation; quantify LIG4 and other NHEJ factors (XRCC4, DNA-PKcs); test rescue by LIG4 overexpression.
- **Priority:** medium

### Hypothesis 4
- **Hypothesis:** SHQ1 upregulation represents a compensatory attempt to maintain snoRNP assembly under global RNA-processing suppression.
- **Why interesting:** Would highlight an adaptive node balancing ribosome biogenesis stress.
- **Suggested validation:** Evaluate snoRNP component levels (e.g., dyskerin/NOP10) and pre-rRNA processing by Northern blot/qPCR; assess rRNA modification status; knock down SHQ1 to test sensitization to camphor-induced RNA biogenesis stress.
- **Priority:** low

## Confidence
medium

## Caution notes
- Upregulated pathway evidence is primarily from Hallmark gene sets; GO/Reactome/KEGG up categories are empty, so membrane/mechanotransduction inferences rely on a limited set of proteins.
- No phosphoproteomics or direct transcription-factor activity assays are provided; STAT3/NFkB/HIF activation is inferred from proteome-based enrichment.
- Single-protein observations (e.g., LIG4, SHQ1) should not be generalized without corroborating pathway-level evidence.
- Cell-type context is unspecified; increases in neuronal/glial-associated proteins (e.g., PIEZO2, BCAS1, GAL3ST1) may reflect lineage-specific responses.
- Exposure conditions are not detailed; inflammatory/hypoxia signatures could reflect generalized stress or culture hypoxia rather than specific TRP-channel engagement.
- Interpretations of survival signaling (e.g., from CFLAR/PRR5) are underpowered and not supported by pathway enrichment.


---

# Carbenoxolone (disodium)

# Drug summary: Carbenoxolone (disodium)

## Executive summary
Proteomics after carbenoxolone exposure shows a robust stress-adaptation profile dominated by (i) hypoxia/HIF-like signaling and pH-regulatory features and (ii) TNF/NF-κB–associated inflammatory activation, alongside (iii) strong suppression of intermediate filament organization. Up-regulated proteins include CA9, SERPINE1, IGFBP1, EFNA1, LOXL2, JUNB, and TXNIP, aligning with Hallmark HYPOXIA (NES ~1.90, FDR ~0.008) and TNFA_SIGNALING_VIA_NFKB (NES ~1.82, FDR ~0.011). Down-regulation of intermediate filament programs is strongly supported by GO terms (NES ~-2.12 and ~-1.99; FDR ≤0.005). ECM features are mixed (LOXL2, CEMIP, SERPINE1 up; POSTN down), indicating non-uniform matrix remodeling. Metabolic stress signatures (TXNIP up; NDUFA3, AMACR, SLC13A5 down) are suggestive of hypoxia-associated rewiring but should be considered preliminary. A KRAS_SIGNALING_DN hallmark decrease (NES ~-1.70, FDR ~0.024) and reduced SPRY2 tentatively point to altered RTK/RAS-MAPK tone, lacking direct pathway corroboration. No clear proteomic linkage to annotated targets (11β-HSD or gap-junction proteins) is observed. Up-regulated GO/Reactome/KEGG pathways are not detected, so conclusions rely on Hallmark enrichments and individual protein changes.

## Findings consistent with expected biology
- Robust hypoxia-like response with pH-regulatory features: Hallmark HYPOXIA up (NES ~1.90, FDR ~0.008) and increased CA9, SERPINE1, IGFBP1, EFNA1, LOXL2.
- Activation of inflammatory signaling consistent with TNF/NF-κB: Hallmark TNFA_SIGNALING_VIA_NFKB up (NES ~1.82, FDR ~0.011) with increases in JUNB, SERPINE1, IGFBP1, TXNIP.
- Strong suppression of intermediate filament organization/cytoskeletal programs: GO BP down (INTERMEDIATE_FILAMENT_ORGANIZATION and INTERMEDIATE_FILAMENT_BASED_PROCESS; NES ~-2.12 and ~-1.99; FDR ≤0.005).

## Unexpected but plausible findings
- Mixed extracellular matrix remodeling rather than a uniform fibrotic program: LOXL2, CEMIP, SERPINE1 up with POSTN down.
- Broad repression of intermediate filament organization at the pathway level despite few explicit IF proteins among the top individual changes, indicating a coordinated programmatic shift.
- Suggestive metabolic stress/rewiring consistent with hypoxia: TXNIP up with decreases in NDUFA3, AMACR, and SLC13A5, but evidence is limited and should be considered preliminary.
- Possible elevation of RTK/RAS–MAPK tone inferred from Hallmark KRAS_SIGNALING_DN down and reduced SPRY2; inference is tentative without direct MAPK pathway markers.

## Follow-up hypotheses
### Hypothesis 1
- **Hypothesis:** Carbenoxolone induces HIF-1–driven transcriptional activation that underlies the hypoxia-like proteomic signature.
- **Why interesting:** Explains concurrent up-regulation of CA9, SERPINE1, IGFBP1, LOXL2 and Hallmark HYPOXIA enrichment; would unify stress-adaptation features.
- **Suggested validation:** Assess HIF1A stabilization and nuclear localization; HRE-luciferase reporter; qPCR/proteomics of canonical HIF targets (CA9, SERPINE1, IGFBP1, LOXL2) under normoxia; use HIF-1 inhibition or HIF1A knockdown to test dependency.
- **Priority:** high

### Hypothesis 2
- **Hypothesis:** Carbenoxolone activates NF-κB signaling, contributing to the inflammatory proteomic response.
- **Why interesting:** Aligns with TNFA_SIGNALING_VIA_NFKB enrichment and increased JUNB, SERPINE1, IGFBP1, TXNIP.
- **Suggested validation:** NF-κB reporter assay; p65/RelA nuclear translocation and IκBα phosphorylation by immunoblot/IF; test sensitivity to NF-κB pathway inhibitors (e.g., IKK inhibitors) and quantify cytokine/chemokine outputs.
- **Priority:** high

### Hypothesis 3
- **Hypothesis:** Suppression of intermediate filament networks alters cell architecture and mechanics secondary to hypoxia/inflammatory stress.
- **Why interesting:** GO BP shows strong down-regulation of IF organization, implying structural reprogramming that may affect adhesion and resilience.
- **Suggested validation:** Immunofluorescence and proteomics of vimentin/keratins; live-cell imaging of IF dynamics; cell mechanics/adhesion assays; test reversal with HIF/NF-κB pathway blockade to probe causality.
- **Priority:** medium

### Hypothesis 4
- **Hypothesis:** Carbenoxolone shifts cellular metabolism toward glycolytic/redox adaptation at the expense of mitochondrial β-oxidation/OXPHOS.
- **Why interesting:** TXNIP up with NDUFA3, AMACR, and SLC13A5 down is consistent with hypoxia-associated metabolic remodeling.
- **Suggested validation:** Seahorse OCR/ECAR profiling; lactate and extracellular pH measurements; mitochondrial membrane potential and ROS assays; targeted metabolomics of TCA and fatty acid oxidation intermediates.
- **Priority:** medium

### Hypothesis 5
- **Hypothesis:** Carbenoxolone may increase RTK/RAS–MAPK signaling tone.
- **Why interesting:** Hallmark KRAS_SIGNALING_DN enrichment in the down direction and decreased SPRY2 suggest a potential increase in RTK/RAS activity.
- **Suggested validation:** Measure phospho-ERK/MEK and RAS-GTP levels; test sensitivity to MEK inhibitors; transcript/proteomic profiling of MAPK target genes.
- **Priority:** low

## Confidence
medium

## Caution notes
- Inferences are based on a single proteomic layer; no phosphoproteomic or transcriptomic data are available to confirm signaling activation states (e.g., HIF, NF-κB, MAPK).
- No significant GO/Reactome/KEGG up-regulated pathways were reported; up-program conclusions rely on Hallmark enrichments plus individual protein changes.
- Metabolic and RTK/RAS interpretations are supported by relatively few proteins and should be viewed as preliminary.
- No direct proteomic linkage to annotated targets (11β-HSD or gap junction proteins) is evident; mechanism-of-action connections remain indirect.
- Experimental context (cell type, oxygenation, inflammatory milieu) is not specified; stress signatures should be validated with orthogonal assays.
- Some pathway sections are empty (e.g., GO BP up), limiting breadth of pathway-level support.


---

# Caroverine (hydrochloride)

# Drug summary: Caroverine (hydrochloride)

## Executive summary
Caroverine (hydrochloride) treatment elicited a proteomic response dominated by lipid/cholesterol remodeling and suppression of epithelial/ECM-associated programs. Hallmark cholesterol homeostasis was significantly up-regulated (NES 1.80, FDR 0.006), accompanied by coordinated changes in lipid-handling proteins (e.g., PCSK9 down; multiple transporters up). In parallel, pathway-level down-regulation of Notch signaling, angiogenesis, apical surface, and intermediate filament organization was observed, with concordant decreases in ECM/secreted factors (CCN2, SPP1, NPNT, CEMIP) and keratins (KRT14, KRT17). Epigenetic repression-associated GO terms (negative regulation of gene expression, heterochromatin organization) were also down, with JARID2 reduced. Neuronal/synaptic and cytoskeletal proteins STMN3 and CPLX2 decreased, consistent with attenuated synaptic/cytoskeletal dynamics, but this is supported only at the protein level and may reflect a non-neuronal/epithelial-like context. Glucocorticoid receptor NR3C1 was reduced, suggesting a potential shift in hormonal/stress signaling. No Reactome or KEGG enrichments were detected, and GO_BP_up was empty; interpretations are therefore grounded in Hallmark and GO down signatures and concordant protein changes. Overall, the data support lipid metabolic adaptation and reduced epithelial/ECM/angiogenic signaling as the primary features, with limited evidence for direct neuronal pathway modulation.

## Findings consistent with expected biology
- Modest decreases in synaptic/cytoskeletal regulators (STMN3, CPLX2), directionally consistent with an iGluR-active agent but supported only at the protein level and without neuronal pathway enrichment.

## Unexpected but plausible findings
- Significant up-regulation of cholesterol homeostasis (NES 1.80, FDR 0.006) with concordant lipid-handling changes (PCSK9 down; transporters/metabolic nodes such as SLC13A5, SLC16A6, SLC25A25, ABHD2 up).
- Down-regulation of Notch signaling and angiogenesis (NES ~ -1.6, FDR ~0.03–0.04) with decreased ECM/secreted factors (CCN2, SPP1, NPNT, CEMIP).
- Suppression of epithelial structural features: apical surface program down (NES -1.59, FDR 0.039) and intermediate filament organization down (GO NES -2.05, FDR 0.010) with keratin reductions (KRT14, KRT17).
- Down-regulation of epigenetic repression-associated processes (NEGATIVE_REGULATION_OF_GENE_EXPRESSION_EPIGENETIC; HETEROCHROMATIN_ORGANIZATION) with JARID2 decreased.
- Glucocorticoid receptor NR3C1 reduction, indicating a potential shift in hormonal/stress signaling tone.

## Follow-up hypotheses
### Hypothesis 1
- **Hypothesis:** Caroverine induces a cellular lipid remodeling response characterized by enhanced cholesterol homeostasis signaling.
- **Why interesting:** Membrane lipid composition can influence ion channel function and broader cell physiology; a robust cholesterol signature may represent an adaptive response with implications for drug action and tolerance.
- **Suggested validation:** Quantify cellular cholesterol and lipid species (untargeted lipidomics); assess LDLR abundance and SREBP target genes; measure cholesterol uptake/efflux (e.g., DiI-LDL uptake); filipin staining for free cholesterol; replicate across cell types.
- **Priority:** high

### Hypothesis 2
- **Hypothesis:** Caroverine dampens pro-angiogenic and ECM signaling via reduced secretion/abundance of factors such as CCN2, SPP1, NPNT, and CEMIP alongside Notch pathway suppression.
- **Why interesting:** Shifts in extracellular signaling may alter microenvironmental responses and cell–matrix interactions, potentially affecting tissue remodeling and drug responses.
- **Suggested validation:** Secretome profiling (ELISA/MRM) for ECM/angiogenic factors; endothelial tube formation or migration assays using conditioned media; Notch reporter assays; qPCR of canonical Notch targets.
- **Priority:** medium

### Hypothesis 3
- **Hypothesis:** Caroverine reduces epithelial structural/polarity features, reflected by down-regulated apical surface and intermediate filament programs.
- **Why interesting:** Loss of epithelial structural integrity can impact barrier function, cell adhesion, and drug permeability.
- **Suggested validation:** Immunofluorescence for keratins (KRT14/KRT17), junctional markers (ZO-1, E-cadherin); transepithelial electrical resistance (TEER); cytoskeletal imaging and morphology assays.
- **Priority:** medium

### Hypothesis 4
- **Hypothesis:** Caroverine attenuates glucocorticoid signaling capacity via reduced NR3C1 protein levels.
- **Why interesting:** GR down-regulation could alter stress-response and inflammatory gene programs, influencing pharmacologic effects and side-effect profiles.
- **Suggested validation:** Measure GR protein and nuclear translocation; assess dexamethasone-induced target gene expression (e.g., FKBP5) and GRE reporter activity; rescue experiments with GR agonists.
- **Priority:** medium

### Hypothesis 5
- **Hypothesis:** Caroverine decreases epigenetic repression tone, leading to a relative de-repression of chromatin.
- **Why interesting:** Chromatin state changes can reprogram cellular responses and may underlie sustained proteomic shifts.
- **Suggested validation:** ATAC-seq for chromatin accessibility; ChIP-qPCR/ChIP-seq for repressive marks (H3K27me3, H3K9me3); expression analysis of known Polycomb targets; broader profiling of chromatin regulators.
- **Priority:** low

## Confidence
medium

## Caution notes
- Cell-type context is unspecified and the proteome shows epithelial/secreted protein features; avoid over-attributing changes to neuronal/iGluR mechanisms.
- Inferences about cholesterol uptake based on PCSK9 down-regulation are preliminary without LDLR measurements or functional assays.
- Pathway suppressions for NOTCH_SIGNALING, ANGIOGENESIS, and APICAL_SURFACE are moderate (NES ~ -1.6; FDR ~0.03–0.04); interpret biological impact conservatively.
- Several interpretations rely on small protein sets without pathway enrichment (e.g., trafficking/stress markers); treat as exploratory.
- No Reactome or KEGG enrichments were detected; mechanistic conclusions should rely on Hallmark/GO signals and concordant proteins.
- Changes in extracellular/secreted proteins (e.g., CCN2, SPP1, DKK1) may reflect secretion dynamics as well as abundance; confirm with orthogonal readouts.
- Protein-level decreases in STMN3/CPLX2 are noted but lack neuronal pathway corroboration and may be context-dependent.


---

# Cetirizine (dihydrochloride)

# Drug summary: Cetirizine (dihydrochloride)

## Executive summary
Cetirizine exposure produced a coherent stress/remodeling proteomic profile dominated by Hallmark enrichments for TNFA signaling via NF-κB and IL6–JAK–STAT3 (NES ~1.8), a hypoxia-like program (NES ~1.7), and a strong ECM/remodeling/differentiation signal (Hallmark MYOGENESIS, NES ~1.86). Corresponding protein increases included IGFBP3, CTF1, RNF128, RC3H2, LOXL2 and SOX18; a modest rise in neuronal/synaptic-associated proteins (SYNGR3, FRRS1, PTBP2, NDEL1) was observed without pathway-level corroboration. Downregulated Gene Ontology terms indicated suppression of rRNA processing and SSU rRNA maturation (NES ~ -1.9; near the FDR threshold), accompanied by decreased TAF1D and increased BRF1, suggesting a modest reduction in Pol I-linked ribosome biogenesis and a tentative shift in RNA polymerase program balance. Additional protein-level decreases in sterol/lipid metabolism enzymes (HMGCR, AKR1C1, AMACR) and selected trafficking/ER-shaping components (CLTCL1, REEP2, ZDHHC9) were noted but lacked pathway enrichment. Several pathway sections were empty (e.g., GO BP up, Hallmark down), so mechanistic specificity relies mainly on Hallmark and limited GO-down signals. Overall, the strongest, pathway-supported conclusions are activation of inflammatory cytokine signaling and a stress-associated remodeling state; more granular mechanistic inferences remain preliminary.

## Findings consistent with expected biology
- Robust enrichment of an ECM/remodeling/differentiation program (Hallmark MYOGENESIS up; supported by LOXL2 and SOX18 increases).
- Modest protein-level increases in neuronal/synaptic-associated proteins (SYNGR3, FRRS1, PTBP2, NDEL1) consistent with the drug’s neuronal signaling annotation, without pathway enrichment.

## Unexpected but plausible findings
- Activation of pro-inflammatory cytokine pathways (Hallmark TNFA_SIGNALING_VIA_NFKB and IL6_JAK_STAT3; NES ~1.8) despite antihistamine context; supported by pathway statistics and concordant increases in immune-associated proteins.
- Hypoxia-like stress program (Hallmark HYPOXIA up; NES ~1.7) with accompanying increases in LOXL2 and IGFBP3.
- Downregulation of rRNA processing and SSU rRNA maturation (GO BP down; NES ~ -1.9) with ↓TAF1D and ↑BRF1, suggesting modest suppression of Pol I–linked ribosome biogenesis and a tentative shift toward Pol III activity.
- Decreases in cholesterol/steroid metabolism enzymes (↓HMGCR, ↓AKR1C1, ↓AMACR) observed at the protein level without pathway enrichment, indicating a possible but unconfirmed reduction in sterol/lipid biosynthesis.
- Selective decreases in trafficking/ER-shaping and antigen presentation–related proteins (↓CLTCL1, ↓REEP2, ↓ZDHHC9, ↓TAPBPL) based on individual proteins only; low confidence without pathway support.

## Follow-up hypotheses
### Hypothesis 1
- **Hypothesis:** Cetirizine induces NF-κB and STAT3 pathway activation in this cell context, driving the observed inflammatory Hallmark enrichments.
- **Why interesting:** Directionally unexpected for an H1 antagonist; if confirmed, indicates context- or off-target–dependent pro-inflammatory signaling.
- **Suggested validation:** Assess NF-κB p65 nuclear translocation and STAT3 Tyr705 phosphorylation by immunoblot/IF; quantify IL6/TNF secretion; perform time-course and dose–response; verify with RNA-seq for NF-κB/STAT3 target gene induction.
- **Priority:** high

### Hypothesis 2
- **Hypothesis:** Cetirizine modestly suppresses nucleolar Pol I activity and rRNA biogenesis with a compensatory tilt toward Pol III programs.
- **Why interesting:** Aligns with GO rRNA processing down and the ↓TAF1D/↑BRF1 pattern; suggests a stress-driven rebalancing of translational capacity.
- **Suggested validation:** Measure 47S/45S pre-rRNA by qRT-PCR; visualize nucleolar markers (fibrillarin, NPM1); ChIP or CUT&RUN for Pol I occupancy; quantify tRNA and Pol III targets; time-course to confirm causality.
- **Priority:** medium

### Hypothesis 3
- **Hypothesis:** A hypoxia-like response contributes to metabolic rewiring, including reduced sterol/lipid biosynthesis.
- **Why interesting:** Connects Hallmark HYPOXIA with protein-level decreases in HMGCR/AKR1C1/AMACR; would implicate HIF/SREBP cross-talk under drug exposure.
- **Suggested validation:** Test HIF-1α stabilization and HRE-reporter activity; profile SREBP target genes by qPCR/RNA-seq; assess cholesterol synthesis flux (13C-acetate incorporation) and cellular lipidomics.
- **Priority:** medium

### Hypothesis 4
- **Hypothesis:** Cetirizine perturbs membrane trafficking/endocytosis in a subset of pathways.
- **Why interesting:** Consistent with decreases in CLTCL1, REEP2, and ZDHHC9; could influence receptor signaling and cargo routing.
- **Suggested validation:** Track transferrin/receptor endocytosis and recycling; assess ER/endomembrane morphology by microscopy; test palmitoylation status of known ZDHHC9 substrates.
- **Priority:** low

## Confidence
medium

## Caution notes
- Pathway interpretations rely predominantly on Hallmark enrichments; Reactome/KEGG and GO BP up sections are empty, limiting mechanistic specificity.
- The rRNA processing suppression signal is near the FDR threshold and supported by few proteins; Pol I suppression and Pol III shift should be considered preliminary.
- Inferences about sterol/lipid metabolism, trafficking, and antigen processing are based on small protein sets without pathway-level support and should be treated as hypotheses.
- Proteomic abundance does not establish signaling activation; phosphorylation and transcriptional readouts (e.g., NF-κB/STAT3) are needed for confirmation.
- Cell-type context is unspecified; interpretation of lineage-biased markers (e.g., CD68, neuronal proteins) should be cautious and not overgeneralized.
- No direct link to histamine receptor engagement is evident in this proteomic snapshot; observed effects may reflect indirect or off-target responses.


---

# Chloroprocaine (hydrochloride)

# Drug summary: Chloroprocaine (hydrochloride)

## Executive summary
Proteomic profiling of Chloroprocaine (hydrochloride) exposure reveals a robust stress program dominated by hypoxia-like and inflammatory (TNFα/NF-κB) signaling with a pathway-level shift toward glucose catabolism. Concordant increases in SERPINE1 and IGFBP1 support these signatures. A strong epithelial/ECM remodeling signal is present (GO: lung alveolus development) with SOX18, BMPR2, CEMIP, and CLEC3B up and EFEMP1 down, consistent with generalized tissue remodeling rather than lung-specific biology. Intermediate filament processes are strongly suppressed, with keratin KRT79 decreased. Isolated lipid-related changes—most notably marked HMGCR downregulation, alongside NEU3 and ZDHHC17 decreases—suggest potential membrane/cholesterol remodeling, although lipid pathways were not enriched. No direct evidence of target engagement or ion homeostasis disruption is available in this dataset. Reactome/KEGG sections were not enriched, which does not detract from the strongly supported Hallmark/GO signals.

## Findings consistent with expected biology
- Robust hypoxia-like response with increased SERPINE1 and IGFBP1 (Hallmark HYPOXIA up).
- Inflammatory activation including TNFα/NF-κB signaling (Hallmark INFLAMMATORY_RESPONSE and TNFA_SIGNALING_VIA_NFKB up).
- Pathway-level shift toward glucose catabolism/glycolysis (GO: glucose catabolic process up).
- Suppression of intermediate filament organization (GO down), exemplified by decreased KRT79.

## Unexpected but plausible findings
- Enrichment of a lung alveolus development program indicating generalized epithelial/ECM remodeling (SOX18, BMPR2, CEMIP, CLEC3B up; EFEMP1 down) outside a lung-specific context.
- Marked HMGCR reduction with concurrent NEU3 and ZDHHC17 decreases, suggesting potential membrane lipid/cholesterol remodeling despite no dedicated lipid pathway enrichment.

## Follow-up hypotheses
### Hypothesis 1
- **Hypothesis:** Chloroprocaine elicits HIF- and NF-κB–mediated stress signaling that drives glycolytic reprogramming and induction of SERPINE1 and IGFBP1.
- **Why interesting:** Integrates the dominant hypoxia and inflammatory pathway signals with concordant protein changes, offering a unifying mechanism for the observed stress phenotype.
- **Suggested validation:** Assess HIF-1α stabilization and NF-κB p65 nuclear translocation; quantify HIF/NF-κB target transcripts; measure extracellular acidification/lactate (Seahorse glycolytic rate); confirm SERPINE1/IGFBP1 secretion by ELISA.
- **Priority:** high

### Hypothesis 2
- **Hypothesis:** The ‘lung alveolus development’ enrichment reflects a broader epithelial/ECM remodeling program involving BMPR2/SOX18/CEMIP that may alter barrier and matrix dynamics.
- **Why interesting:** Explains consistent ECM/vascular-associated protein shifts and may impact cell–matrix interactions relevant to drug response and tissue stress adaptation.
- **Suggested validation:** Profile ECM/remodeling genes by qPCR/proteomics; verify BMPR2, SOX18, CEMIP, EFEMP1 changes by immunoblot; assess matrix deposition (e.g., collagen staining) and cell migration/adhesion assays.
- **Priority:** medium

### Hypothesis 3
- **Hypothesis:** Chloroprocaine reduces cholesterol biosynthetic capacity and select membrane lipid modifications, altering membrane composition and signaling.
- **Why interesting:** HMGCR downregulation (with NEU3 and ZDHHC17 decreases) could modulate membrane properties and receptor/ion channel function, potentially influencing drug responses.
- **Suggested validation:** Measure total/free cholesterol and mevalonate pathway intermediates (lipidomics); test HMGCR/SREBP2 axis activity; rescue experiments with mevalonate; assess global palmitoylation (acyl-biotin exchange) and glycosphingolipid profiles.
- **Priority:** medium

### Hypothesis 4
- **Hypothesis:** Intermediate filament network suppression accompanies the stress response and may shift cytoskeletal architecture toward altered mechanics and trafficking.
- **Why interesting:** Coherent with strong GO downregulation; cytoskeletal changes can affect signaling, adhesion, and secretion under stress.
- **Suggested validation:** Immunofluorescence for keratins/intermediate filaments; quantify additional IF proteins; measure cell mechanical properties (e.g., traction force, deformability) and trafficking assays.
- **Priority:** low

## Confidence
high

## Caution notes
- Conclusions are largely pathway-driven; verify broader marker sets (e.g., canonical glycolytic enzymes) across the ranked list where possible.
- No direct evidence of Na+/K+ ATPase target engagement or ion flux changes is provided; avoid causal attribution to membrane transport without orthogonal assays.
- ‘Lung alveolus development’ likely reflects generic epithelial/ECM remodeling and should not be interpreted as lung-specific biology absent contextual data.
- Lipid/cholesterol remodeling inferences are tentative given lack of pathway-level enrichment; confirm with additional enzyme changes or lipidomics.
- Secreted stress proteins (SERPINE1, IGFBP1) can be generalized stress markers; interpret in the context of the strong hypoxia/inflammatory pathways.
- Empty Reactome/KEGG sections do not negate the observed Hallmark/GO signatures but indicate limited enrichment in those libraries.


---

# Chloropyramine hydrochloride

# Drug summary: Chloropyramine hydrochloride

## Executive summary
Proteomics of chloropyramine hydrochloride exposure shows a dominant induction of inflammatory/cytokine programs, with strong enrichment of IL6–JAK–STAT3, TNFA–NFkB, IL2–STAT5, and Complement hallmarks and increased levels of cytokine/JAK–STAT adapters (IL4R, TYK2, TRAF3, ZC3H12A, CTF1). Concurrently, adhesion and extracellular matrix programs shift, with GO enrichment for extracellular matrix disassembly and homophilic cell–cell adhesion and increases in LOXL2, CEMIP, LAMC3, CLEC3B, and CD164. A coordinated suppression of ribosome biogenesis, rRNA processing/Pol I transcription, and RNA splicing is evident across multiple GO terms (NES ~ −2, very low FDR), accompanied by loss of intermediate filament organization and decreased keratins (KRT14, KRT79), consistent with a stress-adaptive reduction in biosynthetic and cytoskeletal programs. Hallmark cholesterol homeostasis is up modestly (with GAL3ST1 increased), and mitochondrial carrier proteins SLC25A38 (heme biosynthesis–linked) and SLC25A39 (mitochondrial glutathione import) are decreased, suggesting metabolic adjustments. These patterns align with the drug’s Immunology/Inflammation context and are compatible with perturbation of adhesion/RTK/GPCR signaling, though upstream causality (e.g., FAK, VEGFR, histamine receptor engagement) remains inferential. No Hallmark downregulated sets were returned, but numerous GO downregulated pathways indicate robust repression of RNA biogenesis/processing.

## Findings consistent with expected biology
- Activation of cytokine/inflammatory signaling: Hallmark IL6–JAK–STAT3, TNFA–NFkB, IL2–STAT5 enrichments with increased IL4R, TYK2, TRAF3, ZC3H12A, and CTF1.
- Engagement of adhesion and extracellular matrix remodeling: GO up for extracellular matrix disassembly and homophilic cell–cell adhesion with increases in LOXL2, CEMIP, LAMC3, CLEC3B, and CD164.
- Complement pathway engagement consistent with an inflammatory milieu (Hallmark Complement up).

## Unexpected but plausible findings
- Strong suppression of ribosome biogenesis, rRNA processing/Pol I transcription, and RNA splicing (multiple GO BP down with NES ~ −2 and FDR ≈ 0), suggesting a broad downshift of biosynthetic/RNA-processing programs.
- Marked repression of intermediate filament organization with decreased keratins KRT14 and KRT79, indicating cytoskeletal remodeling.
- Upregulation of Hallmark cholesterol homeostasis with limited direct protein-level support (GAL3ST1 increased), pointing to modest lipid/cholesterol program engagement.
- Downregulation of mitochondrial carriers SLC25A38 (heme biosynthesis–related) and SLC25A39 (mitochondrial glutathione import), suggesting altered mitochondrial heme/redox homeostasis.
- Enrichment of osteoclast differentiation, and borderline Hallmark estrogen response early and myogenesis programs; likely secondary to inflammatory/ECM signaling rather than primary on-target effects.

## Follow-up hypotheses
### Hypothesis 1
- **Hypothesis:** Chloropyramine activates JAK/STAT and NFkB inflammatory signaling cascades in treated cells.
- **Why interesting:** Strong Hallmark enrichment (IL6–JAK–STAT3, TNFA–NFkB, IL2–STAT5) with increased IL4R, TYK2, TRAF3, ZC3H12A, and CTF1 indicates coordinated cytokine pathway activation aligned with the drug’s immunology context.
- **Suggested validation:** Measure STAT3 Tyr705 phosphorylation and NFkB p65 nuclear translocation; run STAT/NFkB reporter assays; quantify canonical targets (e.g., SOCS3, CCL2) by qPCR; test reversibility with JAK or IKK inhibitors.
- **Priority:** high

### Hypothesis 2
- **Hypothesis:** The compound induces a stress-adaptive repression of ribosome biogenesis and RNA processing, reducing translational capacity.
- **Why interesting:** Multiple GO terms for rRNA processing, Pol I transcription, and splicing are strongly downregulated (NES ~ −2), consistent with a coordinated reduction of biosynthetic machinery.
- **Suggested validation:** Assess nucleolar integrity (fibrillarin/UBF immunofluorescence), 47S pre‑rRNA levels (qPCR), nascent RNA synthesis (EU incorporation), and protein synthesis (puromycin SUnSET); correlate with cell proliferation indices.
- **Priority:** high

### Hypothesis 3
- **Hypothesis:** Chloropyramine perturbs cell–ECM and cell–cell adhesion dynamics, promoting ECM disassembly.
- **Why interesting:** GO enrichment for extracellular matrix disassembly and homophilic adhesion with increased LOXL2, CEMIP, LAMC3, CLEC3B, and CD164 suggests altered adhesion/ECM turnover.
- **Suggested validation:** Perform adhesion and detachment assays, collagen/gelatin degradation assays, and gelatin zymography; image focal adhesion markers (pFAK Y397, paxillin) and cadherin localization; test whether FAK pathway inhibitors modulate the phenotype.
- **Priority:** medium

### Hypothesis 4
- **Hypothesis:** Mitochondrial redox/heme homeostasis is perturbed via decreased SLC25A39 (mitochondrial glutathione import) and SLC25A38 (heme biosynthesis transporter).
- **Why interesting:** Concurrent downregulation of these carriers suggests decreased mitochondrial glutathione and altered heme synthesis, consistent with broader stress and metabolic adaptation.
- **Suggested validation:** Quantify mitochondrial GSH pools (biochemical assays), cellular heme content, and ROS; assess mitochondrial respiration; rescue experiments by expressing SLC25A39 or supplementing GSH precursors.
- **Priority:** medium

### Hypothesis 5
- **Hypothesis:** Cholesterol homeostasis pathways are modestly engaged downstream of inflammatory signaling.
- **Why interesting:** Hallmark cholesterol homeostasis is enriched and GAL3ST1 is increased, indicating lipid pathway involvement albeit with limited breadth at the protein level.
- **Suggested validation:** Measure SREBP2 processing and LDLR/ABCA1 levels; assess cellular cholesterol by filipin staining or mass spectrometry; test sensitivity to statins or cholesterol loading.
- **Priority:** low

## Confidence
medium

## Caution notes
- Pathway enrichments indicate coordinated patterns but do not establish upstream causality; attribution to FAK/VEGFR or histamine receptor signaling remains inferential and requires phospho/signaling assays.
- The lipid/cholesterol inference is based on one Hallmark set and limited protein-level support; treat as modest and confirm with orthogonal lipid measurements.
- Interpretation of SLC25A38/39 should distinguish their roles: SLC25A38 is linked to heme biosynthesis, whereas SLC25A39 mediates mitochondrial glutathione import; avoid conflating their functions.
- The conclusion of a ‘growth-suppressive’ state is suggested by repression of RNA biogenesis/processing but lacks direct proliferation or cell-cycle markers.
- Osteoclast differentiation, estrogen response early, and myogenesis enrichments may reflect cross-talk with inflammatory/ECM programs; consider them secondary until validated.
- Absence of Hallmark downregulated sets does not imply lack of repression; GO downregulated pathways are numerous and robust, indicating strong suppression of RNA biogenesis/processing.


---

# Chlorpheniramine (maleate)

# Drug summary: Chlorpheniramine (maleate)

## Executive summary
Proteomic profiling of chlorpheniramine (maleate) shows a robust pro-inflammatory activation state characterized by IL6–JAK–STAT3 (Hallmark NES ~2.19, FDR ~0) and TNF–NFκB signaling (NES ~1.64), accompanied by hypoxia and angiogenesis programs (HYPOXIA NES ~1.87; ANGIOGENESIS NES ~1.71) and EMT/TGF-β–associated remodeling (EMT NES ~1.63; TGF_BETA_SIGNALING NES ~1.62). Concomitantly, epithelial/intermediate filament organization is strongly suppressed (GO down: INTERMEDIATE_FILAMENT_ORGANIZATION NES ~-2.50; INTERMEDIATE_FILAMENT_BASED_PROCESS NES ~-2.41), with additional decreases in rRNA processing and protein localization to chromatin. Protein-level changes align with these pathway signals: ECM modifiers and stress/angiogenic modulators are increased (LOXL2, CEMIP, IGFBP1, IGFBP5), while epithelial/adhesion-associated proteins are decreased (KRT79, CALML3, S100A14, LIMS2). A KRAS-like gene-set pattern is present (KRAS_SIGNALING_UP enriched; KRAS_SIGNALING_DN depleted) but lacks direct RAS/MAPK proteomic support and is best interpreted as a convergent stress/inflammatory transcriptional signature. KEGG/Reactome and GO BP up enrichments were not detected; thus, conclusions rely chiefly on Hallmark sets integrated with targeted protein changes. The pro-inflammatory, EMT/hypoxia direction is unexpected for a histamine receptor–targeting antihistamine and may reflect context- or dose-dependent off-target or stress responses in this model.

## Findings consistent with expected biology
- Engagement of immunology/inflammation-related programs is plausible for a histamine receptor–targeting compound, although the observed direction here is activation rather than suppression.
- ECM/stromal remodeling signatures (e.g., LOXL2, CEMIP up) alongside inflammatory activation are biologically coherent as part of a generalized stress/adaptive response.

## Unexpected but plausible findings
- Strong activation of IL6–JAK–STAT3 and TNF–NFκB inflammatory programs despite antihistamine targeting, supported by multiple Hallmark enrichments.
- Up-regulation of hypoxia and angiogenesis programs (HYPOXIA, ANGIOGENESIS), consistent with a stress-adaptive state.
- EMT/TGF-β–associated remodeling with suppression of epithelial/intermediate filament organization (EMT and TGF-β hallmarks up; GO down for intermediate filament processes; KRT79, CALML3, S100A14, LIMS2 down) and ECM modifiers up (LOXL2, CEMIP).
- ‘KRAS-like’ gene-set activation (KRAS_SIGNALING_UP enriched with reciprocal KRAS_SIGNALING_DN suppressed) without direct MAPK/RAS protein support; likely a convergent transcriptional program rather than canonical KRAS pathway activation.
- Marked increases in IGFBP1 and IGFBP5, indicating IGF-axis modulation under stress/angiogenic conditions; direction of IGF signaling remains indeterminate.
- Modest decrease in PTEN (single-protein, log2FC ~ -0.62) noted as tentative and not mechanistically interpretable without additional evidence.

## Follow-up hypotheses
### Hypothesis 1
- **Hypothesis:** Chlorpheniramine triggers IL6–STAT3 and NFκB activation via context-specific off-target or compensatory mechanisms in this cell type, driving downstream EMT/hypoxia programs.
- **Why interesting:** Pro-inflammatory activation is unexpected for an H1 antagonist and could explain the coordinated EMT, angiogenesis, and epithelial cytoskeleton suppression signatures.
- **Suggested validation:** Time- and dose-response assays for STAT3 (Y705) and NFκB p65 (Ser536) phosphorylation; measure secreted IL6/TNFα; compare to other H1 antagonists and H1 receptor knockdown; test pathway dependency by JAK inhibition or IL6R neutralization to see whether EMT/angiogenesis markers (e.g., LOXL2, IGFBP1/5) are attenuated.
- **Priority:** high

### Hypothesis 2
- **Hypothesis:** EMT/TGF-β pathway engagement mediates ECM remodeling and loss of epithelial cytoskeletal programs in response to drug-induced stress.
- **Why interesting:** EMT and TGF-β hallmarks are up, ECM modifiers (LOXL2, CEMIP) are increased, and epithelial/intermediate filament components are suppressed.
- **Suggested validation:** Assess SMAD2/3 phosphorylation and TGF-β1 levels; quantify EMT markers (E-cadherin, Vimentin) and migration; test reversal using a TGF-β receptor inhibitor (e.g., SB431542) and evaluate rescue of epithelial markers and reduction of LOXL2/CEMIP and IGFBP1/5.
- **Priority:** medium

### Hypothesis 3
- **Hypothesis:** HIF-1–driven hypoxia signaling is engaged as part of a stress-adaptive response, contributing to the angiogenesis signature and IGFBP up-regulation.
- **Why interesting:** HYPOXIA and ANGIOGENESIS hallmarks are enriched; several stress/ECM-related proteins are up-regulated.
- **Suggested validation:** Measure HIF-1α stabilization and HIF target genes (VEGFA, CA9) under normoxia; evaluate sensitivity to HIF pathway inhibition (e.g., echinomycin) and compare with prolyl hydroxylase inhibitor controls; assess mitochondrial/respiratory stress (Seahorse OCR/ECAR).
- **Priority:** medium

### Hypothesis 4
- **Hypothesis:** IGF signaling is modulated by increased IGFBP1/5, altering IGF availability and downstream signaling in this setting.
- **Why interesting:** Substantial increases in IGFBP1 and IGFBP5 suggest altered IGF ligand sequestration or presentation during stress.
- **Suggested validation:** Measure free IGF1/IGF2 levels and IGF1R phosphorylation with/without exogenous IGF in the presence of drug; assess AKT/ERK readouts; perturb IGFBPs (neutralization or knockdown) to test causality.
- **Priority:** medium

### Hypothesis 5
- **Hypothesis:** The ‘KRAS-like’ Hallmark signature reflects downstream STAT3/NFκB-driven transcription rather than canonical RAS–MAPK activation.
- **Why interesting:** KRAS up/down Hallmark pattern is present without direct proteomic evidence for MAPK activation, suggesting convergent transcriptional outputs.
- **Suggested validation:** Profile ERK1/2 phosphorylation and RAS activity; test whether MEK inhibition reduces the KRAS_UP signature; evaluate whether JAK/STAT3 or NFκB inhibition abrogates the KRAS-like enrichment.
- **Priority:** low

## Confidence
medium

## Caution notes
- Pathway inferences rely predominantly on Hallmark enrichments applied to proteomic data; transcription factors/cytokines are underrepresented in proteomics, so signals reflect correlated programs rather than direct measurement.
- Cell type, dose, and timepoint are not provided; the pro-inflammatory, hypoxia/angiogenesis, and EMT-like signatures may reflect context-dependent or off-target/stress responses rather than on-target H1 receptor antagonism.
- KRAS-like gene-set activation should not be equated with oncogenic RAS/MAPK pathway activation without phospho-ERK/RAS activity evidence.
- Angiogenesis enrichment indicates an angiogenic program at the proteome level and does not imply functional angiogenesis in vitro.
- IGFBP1/IGFBP5 increases indicate IGF-axis modulation but do not specify whether IGF signaling is increased or decreased; functional assays are required.
- Single-protein observations (e.g., PTEN decrease) are insufficient for mechanistic claims without convergent pathway evidence.
- GO BP up, KEGG, and Reactome pathway enrichments were not detected; conclusions emphasize Hallmark sets and selected protein changes.
- Suppression of rRNA processing and protein localization to chromatin is based on single GO BP down sets and should be considered tentative.
- Use of TXNIP as hypoxia evidence is limited; it more broadly reflects redox/metabolic stress.


---

# Chlorpropamide

# Drug summary: Chlorpropamide

## Executive summary
Chlorpropamide induced a coherent growth-suppression signature characterized by strong negative enrichment of E2F targets, G2/M checkpoint, and mitotic spindle programs, with concordant decreases in key cell-cycle proteins (e.g., CDC25B, CDT1, PTTG1, SKA1, BIRC5). In parallel, lipid/cholesterol and lipoprotein metabolism programs were robustly upregulated (cholesterol homeostasis; sterol biosynthesis), accompanied by increased nutrient transport signatures (amino acid and organic ion transport; import across the plasma membrane). Inflammatory/IL6–JAK–STAT3 signaling and a coagulation-leaning profile were prominent, with upregulation of IL10RB, IL4R, IL32, and complement C9. Stress-associated features (HYPOXIA hallmark, DDIT4 up) co-occurred with an up MTORC1 hallmark, indicating nuanced or time-dependent mTOR pathway behavior. Additional enrichment of EMT/adipogenesis/myogenesis hallmarks and increased APOA5, ANGPTL8, DLK1, and FSTL3 point to a lipid-handling/secretory shift accompanying growth suppression. Notably, metallothioneins (MT1E, MT1X) decreased despite stress/inflammatory signals. Reactome and KEGG sections were empty, but Hallmark and GO enrichments with multiple concordant protein-level changes support these conclusions. No direct proteomic evidence links the response to Na+/K+ ATPase subunits, so mechanistic connection to the nominal target remains to be established.

## Findings consistent with expected biology
- Coordinated suppression of proliferation programs (E2F targets, G2/M checkpoint, mitotic spindle) with decreases in CDC25B, CDT1, PTTG1, SKA1, BIRC5, CDKN3.
- Robust induction of cholesterol/sterol and fatty acid/bile acid metabolism (cholesterol homeostasis and sterol biosynthesis hallmarks), with APOA5 and ANGPTL8 up.
- Upregulation of membrane transport processes (amino acid and organic anion/cation transport; import across plasma membrane), exemplified by SLC16A13 increases.
- Activation of inflammatory IL6–JAK–STAT3 signaling and a coagulation/secretory tilt, with IL10RB, IL4R, IL32, and complement C9 up.
- Engagement of stress-associated programs (HYPOXIA hallmark) and DDIT4 upregulation consistent with cellular stress signaling.

## Unexpected but plausible findings
- Mixed mTOR-related signals: HYPOXIA hallmark and DDIT4 up alongside an up MTORC1 hallmark, suggesting context-dependent mTORC1 behavior.
- Downregulation of metallothioneins (MT1E, MT1X) despite stress/inflammatory pathway engagement.
- Concomitant enrichment of EMT/adipogenesis/myogenesis hallmarks with growth suppression.
- Upregulation of lipid-handling/secreted factors (APOA5, ANGPTL8, DLK1, FSTL3) not typically elevated in proliferative states.

## Follow-up hypotheses
### Hypothesis 1
- **Hypothesis:** Chlorpropamide primarily induces cell-cycle arrest (cytostasis) rather than overt cytotoxicity.
- **Why interesting:** Distinguishing cytostasis from cell death informs dosing, combinations, and biomarker selection.
- **Suggested validation:** Time-course viability and apoptosis assays (Annexin V/PI, caspase 3/7), cell-cycle profiling by flow cytometry (DNA content; pH3 for mitosis), and clonogenic recovery assays.
- **Priority:** high

### Hypothesis 2
- **Hypothesis:** The drug drives lipid/cholesterol biosynthesis and lipoprotein handling via stress/inflammatory (STAT3-linked) signaling.
- **Why interesting:** Defines a metabolic adaptation that could be targetable or serve as a pharmacodynamic biomarker.
- **Suggested validation:** Measure pSTAT3 (Y705) and IL6 levels; perturb with JAK/STAT inhibitors; quantify SREBP2/HMGCR and sterol biosynthesis flux (13C-acetate tracing); assess secreted APOA5/ANGPTL8 by ELISA.
- **Priority:** high

### Hypothesis 3
- **Hypothesis:** mTORC1 activity is dynamically modulated (transient activation followed by DDIT4-mediated restraint under stress).
- **Why interesting:** mTOR dynamics may dictate combination strategies with nutrient or mTOR pathway inhibitors.
- **Suggested validation:** Time-resolved phospho-S6 and phospho-4EBP1 immunoblotting; DDIT4 induction kinetics; rapamycin/torin sensitivity assays.
- **Priority:** medium

### Hypothesis 4
- **Hypothesis:** Perturbation of Na+/K+ ATPase function contributes to the observed stress and transport signatures.
- **Why interesting:** Links the proteomic response to the nominal target and may explain IL6/transport induction.
- **Suggested validation:** Assess Na+/K+ ATPase activity, intracellular Na+ and K+ (ion-selective probes), and ATP1A/B subunit abundance; test competition/phenocopy with ouabain; monitor downstream ion-stress markers.
- **Priority:** medium

### Hypothesis 5
- **Hypothesis:** Chlorpropamide induces a secretory phenotype with complement/coagulation factor production.
- **Why interesting:** Paracrine outputs could modulate microenvironmental responses and safety profiles.
- **Suggested validation:** Secretome analysis of conditioned media (ELISA or proteomics) for C9, IL32, and coagulation factors; test sensitivity to JAK pathway inhibition.
- **Priority:** medium

### Hypothesis 6
- **Hypothesis:** Reduced metallothionein expression alters metal/oxidative stress handling.
- **Why interesting:** Could affect redox homeostasis and sensitivity to metal or oxidative stresses.
- **Suggested validation:** Measure cellular Zn2+ and ROS; challenge with zinc/cadmium or oxidants; quantify MT1/MT2 transcripts and proteins after treatment.
- **Priority:** low

## Confidence
high

## Caution notes
- Proteomic suppression of cell-cycle programs cannot by itself distinguish cytostasis from cytotoxicity; dedicated viability and cell-cycle assays are required.
- mTOR pathway status cannot be resolved from protein abundance and hallmark scores alone; phospho-specific readouts are needed.
- Interpretation of a secretory/coagulation phenotype should be corroborated with conditioned-media measurements; cell lysate data alone may be confounded.
- Claims of broad transport upregulation are driven by pathway enrichment; only a subset of transporters (e.g., SLC16A13) are individually highlighted.
- No direct evidence links the response to Na+/K+ ATPase modulation; ATP1A/B subunits and ion homeostasis should be examined to connect to the nominal target.
- Reactome and KEGG sections were empty; conclusions rely on robust Hallmark and GO BP enrichments with concordant protein-level changes.


---

# Ciclesonide

# Drug summary: Ciclesonide

## Executive summary
Ciclesonide treatment yields a coherent glucocorticoid receptor–like proteomic response characterized by strong suppression of proliferation and biosynthetic machinery (E2F/G2M programs; ribosome biogenesis, rRNA processing, RNA splicing, chromatin organization) and a shift toward catabolic metabolism. Lipid and organic acid catabolic processes are enriched, with marked induction of CYP24A1 and CYP3A7 and reduced PLIN2, consistent with decreased lipid storage and increased detoxification capacity. Stress/growth-restraint markers TXNIP and DDIT4 are up, supporting a growth-inhibitory state, though mTOR attenuation remains inferential. Unexpectedly, type I interferon–like and coagulation/acute inflammatory signatures are induced (ISG20, HLA-B; Hallmark IFN-α and Coagulation), suggesting context-dependent innate immune activation under GR agonism. Reactome/KEGG sections are empty, but Hallmark and GO evidence is robust; immune and nuclear receptor cross-talk interpretations should be considered preliminary.

## Findings consistent with expected biology
- Suppression of proliferation programs: Hallmark E2F_TARGETS and G2M_CHECKPOINT are strongly down, consistent with GR-linked anti-proliferative effects.
- Broad repression of biosynthetic and RNA-processing machinery: significant downregulation of ribosome biogenesis, rRNA processing, ribonucleoprotein biogenesis, RNA splicing, and chromatin organization.
- Metabolic shift toward catabolism: enrichment of lipid, glycerolipid, monocarboxylate, and organic acid catabolic processes.
- Induction of steroid/xenobiotic metabolism enzymes: robust upregulation of CYP24A1 and CYP3A7.
- Stress/growth restraint markers up: TXNIP and DDIT4 are increased, aligning with an anti-growth, stress-adaptive state.
- Reduced lipid storage: PLIN2 is down, consistent with fewer lipid droplets.

## Unexpected but plausible findings
- Type I interferon–like activation under a GR agonist: Hallmark INTERFERON_ALPHA_RESPONSE is enriched with ISG20 and HLA-B up.
- Pro-coagulation/acute inflammatory module: Hallmark COAGULATION and GO acute inflammatory response are enriched despite GR targeting.
- Potential nuclear receptor cross-talk involving vitamin D metabolism: very strong CYP24A1 induction suggests enhanced vitamin D catabolism, but VDR pathway activation is not established.
- ECM remodeling signals are mixed and tentative (LOXL2 up with MMP14 and COL7A1 down) without pathway-level enrichment.
- Strong MIXL1 upregulation (developmental transcription factor) lacks pathway corroboration and may be cell-type specific or off-target.

## Follow-up hypotheses
### Hypothesis 1
- **Hypothesis:** Ciclesonide-induced growth restraint is mediated by GR-dependent repression of E2F/G2M programs and reduced translational capacity.
- **Why interesting:** Explains the dominant anti-proliferative phenotype and integrates multiple downregulated biosynthetic pathways.
- **Suggested validation:** Test GR dependence with RU486 or NR3C1 knockdown; assess proliferation (EdU/BrdU, Ki-67), cell-cycle markers (phospho-RB), and nascent protein synthesis (puromycin incorporation).
- **Priority:** high

### Hypothesis 2
- **Hypothesis:** mTOR signaling is attenuated downstream of DDIT4/TXNIP induction, contributing to reduced protein synthesis.
- **Why interesting:** Provides a mechanistic link between stress markers and broad suppression of ribosome biogenesis and translation.
- **Suggested validation:** Measure phospho-S6K, phospho-4E-BP1, and global translation rates; compare to rapamycin; evaluate rescue by DDIT4 or TXNIP knockdown.
- **Priority:** medium

### Hypothesis 3
- **Hypothesis:** Ciclesonide triggers a context-dependent type I interferon–like program via autocrine/paracrine signaling or GR–STAT/IRF cross-talk.
- **Why interesting:** Accounts for unexpected IFN-α enrichment under GR agonism and has implications for antiviral and immunomodulatory effects.
- **Suggested validation:** Quantify ISG panel (e.g., IFITs, MX1), STAT1/2 and IRF3/7 activation; measure IFN-α/β in supernatant; block with IFNAR-neutralizing antibody or JAK inhibitors and assess reversal.
- **Priority:** high

### Hypothesis 4
- **Hypothesis:** CYP24A1 upregulation reduces active vitamin D levels, altering VDR signaling and contributing to metabolic reprogramming.
- **Why interesting:** Links pronounced CYP24A1 induction to potential nuclear receptor cross-talk and metabolic outcomes.
- **Suggested validation:** Measure 1,25(OH)2D and 25(OH)D metabolites; assess VDR target proteins/pathway enrichment; test CYP24A1 inhibition or VDR agonism for modulation of the response.
- **Priority:** medium

### Hypothesis 5
- **Hypothesis:** The proteome favors reduced lipid storage and increased fatty acid oxidation under ciclesonide.
- **Why interesting:** Aligns PLIN2 down and catabolic GO enrichments with functional metabolic consequences.
- **Suggested validation:** Assess lipid droplets (BODIPY staining), triglyceride content, and FAO rates (Seahorse palmitate oxidation assay); profile acylcarnitines.
- **Priority:** medium

### Hypothesis 6
- **Hypothesis:** Apparent pro-coagulation/inflammatory signature reflects secreted factor changes rather than intracellular proteome alone.
- **Why interesting:** Clarifies the basis of the Hallmark Coagulation and acute inflammatory enrichments under GR agonism.
- **Suggested validation:** Secretome proteomics or ELISAs for coagulation/inflammatory proteins (e.g., SERPINA1, FGA/FGB if relevant cell type); pathway enrichment on secreted proteins.
- **Priority:** low

## Confidence
medium

## Caution notes
- Cell type, dose, and time-point are unspecified; GR and interferon responses are highly context dependent.
- mTOR pathway inhibition and ‘quiescence’ are inferred and lack direct signaling readouts; phosphoproteomics or functional assays are needed.
- Antigen presentation inference from HLA-B alone is limited; broader antigen-processing components were not shown upregulated.
- Vitamin D receptor cross-talk remains speculative without additional VDR targets or pathway enrichment.
- Generalization to broad xenobiotic metabolism induction is not supported beyond CYP24A1 and CYP3A7.
- ECM remodeling interpretation is tentative and based on few proteins without pathway-level support.
- Reactome and KEGG sections are empty; conclusions rest on Hallmark and GO enrichments and selected protein-level changes.


---

# Clobetasone butyrate

# Drug summary: Clobetasone butyrate

## Executive summary
Proteomics profiling of clobetasone butyrate shows a glucocorticoid receptor–consistent signature dominated by anti-proliferative and anti-remodeling effects. Hallmark cell-cycle programs are robustly suppressed (E2F_TARGETS NES -1.85, FDR 0.005; G2M_CHECKPOINT NES -1.68, FDR 0.034), and a metallothionein-centered metal-ion response is induced (GO: response to copper ion NES 2.06, FDR 0.047) with coordinated increases in MT1B/MT1E/MT1H/MT2A. Several extracellular matrix/adhesion and wound-repair proteins (EFEMP1, COL6A1, SPP1, CCN1, CLDN1) decrease, consistent with anti-remodeling. NR3C1 protein is modestly but significantly reduced, consistent with receptor feedback, though this is single-protein evidence. Additional signals—CYP3A7/CYP4F2/CYP4F22 induction, a small set of stress/innate markers (S100A7, TRIM21, TNFRSF12A), and a possible metabolic downshift (PFKM, NME4, BNIP3 decreases)—are plausible but pathway under-supported and should be considered tentative. A marked and unexpected MIXL1 increase stands out and warrants follow-up. No KEGG/Reactome enrichments were detected; empty pathway sections likely reflect thresholding rather than absence of biology. The KRAS hallmark change (KRAS_SIGNALING_DN decreased) is noted but not interpreted due to set semantics.

## Findings consistent with expected biology
- Strong suppression of cell-cycle programs (Hallmark E2F_TARGETS NES -1.847, FDR 0.005; G2M_CHECKPOINT NES -1.681, FDR 0.034), indicative of an anti-proliferative state.
- Induction of a metallothionein-driven metal-ion response (GO: response to copper ion NES 2.056, FDR 0.047) with coordinated upregulation of MT1B, MT1E, MT1H, and MT2A.
- Protein-level downregulation of extracellular matrix/adhesion and wound-repair factors (e.g., EFEMP1, COL6A1, SPP1, CCN1, CLDN1), consistent with anti-remodeling effects.
- Modest NR3C1 (glucocorticoid receptor) protein decrease (log2FC ~ -1.02, highly significant), consistent with receptor feedback; interpret cautiously as single-protein evidence.

## Unexpected but plausible findings
- Very strong and highly significant MIXL1 induction (log2FC ~ 3.95), atypical for canonical GR programs and merits follow-up.
- Upregulation of S100A7 with increases in TRIM21 and TNFRSF12A suggests a limited stress/innate-response component despite overall anti-proliferative signaling (protein-level only, no pathway enrichment).
- Coordinated increases in CYP3A7, CYP4F2, and CYP4F22 suggest possible xenobiotic/lipid-metabolism activation via nuclear receptor cross-talk; tentative without pathway support.
- Signals consistent with a metabolic downshift (decreased PFKM, NME4, BNIP3) are observed but lack pathway-level support and should be treated as preliminary.

## Follow-up hypotheses
### Hypothesis 1
- **Hypothesis:** MIXL1 induction is a GR-dependent transcriptional event that contributes to the observed state change.
- **Why interesting:** MIXL1 is a transcription factor not typically associated with GR agonism; its strong upregulation could reflect an unrecognized regulatory axis or cell-state transition.
- **Suggested validation:** Confirm by qPCR and Western time course; test GR dependence via GR antagonism (e.g., mifepristone) or NR3C1 knockdown/CRISPR; assess GR occupancy at MIXL1 regulatory regions by ChIP-qPCR; evaluate downstream transcriptional effects by RNA-seq.
- **Priority:** high

### Hypothesis 2
- **Hypothesis:** Metallothionein induction is mediated by MTF1 and/or GR, conferring cytoprotective metal/oxidative stress buffering.
- **Why interesting:** Mechanistic attribution clarifies whether this response is a direct GR effect or involves metal-responsive transcription factor cross-talk, with implications for stress resilience.
- **Suggested validation:** qPCR/Western for MT1/MT2 isoforms and MTF1; MTF1 knockdown or inhibitor where applicable; luciferase reporters for metal response elements; GR antagonism to test dependence; challenge with copper/zinc to assess functional buffering.
- **Priority:** medium

### Hypothesis 3
- **Hypothesis:** CYP3A7/CYP4F2/CYP4F22 upregulation arises from PXR/CAR cross-talk rather than direct GR activation.
- **Why interesting:** Defines potential drug–drug interaction liabilities and nuclear receptor cross-regulation under clobetasone exposure.
- **Suggested validation:** Measure CYP mRNA/protein across time; use PXR/CAR agonists/antagonists or siRNA/CRISPR to test dependence; perform PXR/CAR reporter assays; compare with a selective GR agonist and with PXR-null or CAR-deficient models if available.
- **Priority:** medium

### Hypothesis 4
- **Hypothesis:** Reduced ECM/adhesion protein abundance translates into functional anti-remodeling (reduced migration/contraction).
- **Why interesting:** Links proteomic anti-remodeling signals to phenotypic outcomes relevant to tissue repair and inflammation.
- **Suggested validation:** Scratch-wound or transwell migration assays; collagen gel contraction; ELISA/Western for EFEMP1/SPP1/CCN1; rescue with exogenous matrix factors to test causality.
- **Priority:** medium

### Hypothesis 5
- **Hypothesis:** NR3C1 downregulation reflects receptor feedback/desensitization kinetics after agonist exposure.
- **Why interesting:** Receptor autoregulation influences duration and magnitude of GR responses.
- **Suggested validation:** Time-course qPCR/Western for NR3C1; assess GR nuclear translocation and degradation; test reversibility after washout; evaluate dependence on proteasome activity.
- **Priority:** low

## Confidence
medium

## Caution notes
- Beyond cell-cycle suppression and the copper/metal-ion response, other thematic inferences (xenobiotic metabolism, metabolic downshift, innate/stress signaling) rely on a small number of proteins without pathway enrichment and should be treated as tentative.
- KRAS hallmark sets are context-dependent; the decrease in KRAS_SIGNALING_DN is not interpreted as reduced KRAS/pro-growth signaling without corroboration.
- NR3C1, MIXL1, and S100A7 interpretations are based on proteomics only; orthogonal validation (transcript-level assays, targeted protein assays) is recommended.
- CYP induction may reflect nuclear receptor cross-talk (e.g., PXR/CAR) and could be cell-type specific; attribute cautiously.
- ECM/adhesion downregulation is supported by multiple proteins but lacks pathway enrichment; treat as protein-level evidence pending broader confirmation.
- Cell type and exposure conditions are not specified; responses such as metallothionein and S100A7 induction can be highly context dependent.
- This is a single-layer proteomics snapshot without phosphoproteomics or transcriptomics, limiting mechanistic attribution.
- Empty pathway sections (e.g., KEGG/Reactome, Hallmark up) reflect significance thresholds rather than absence of biology.


---

# Clomiphene (citrate)

# Drug summary: Clomiphene (citrate)

## Executive summary
Clomiphene (citrate) elicits a robust anti-proliferative proteomic response characterized by down-regulation of E2F/G2M cell-cycle programs, DNA replication, and homologous recombination, with concordant decreases in key regulators (e.g., CDC25B, ZWINT, SASS6, RAD54B, RAD51AP1, CDCA7, USP1). In parallel, there is a strong induction of epithelial/keratinocyte differentiation and intermediate filament organization, accompanied by an antimicrobial peptide signature (e.g., S100A7, DCD). The treatment also activates a pronounced secretory/inflammatory program, including complement/coagulation and angiogenesis-related factors (e.g., F5, C9, POSTN, ANGPTL8, NRP2, TNFRSF12A), and enriches apoptosis pathways, suggesting a predominantly cytostatic response with a potential cytotoxic component. Stress/hypoxia signaling is engaged (HYPOXIA hallmark; DDIT4 up), alongside metabolic pathway enrichment for oxidative phosphorylation, TCA cycle, and lipid/adipogenic and xenobiotic metabolism. The IGF axis appears markedly remodeled with strong up-regulation of IGFBP1/2/3/5, consistent with altered IGF signaling tone. Notably, mTORC1 signaling is enriched at the pathway level despite REDD1 (DDIT4) induction, indicating potentially complex or temporal mTOR regulation. Classical estrogen response signatures are not conspicuous, so direct linkage to on-target ER modulation is not established by these data alone. Overall, the profile supports growth restraint with epithelial differentiation and a broad secretory/metabolic reprogramming, with key mechanistic aspects requiring orthogonal validation.

## Findings consistent with expected biology
- Anti-proliferative program with suppression of E2F/G2M modules and DNA replication/HR pathways, supported by decreases in CDC25B, ZWINT, SASS6, RAD54B, RAD51AP1, CDCA7, USP1
- Remodeling of the IGF axis with strong up-regulation of IGFBP1/2/3/5 and enrichment of IGF receptor signaling-related processes

## Unexpected but plausible findings
- Robust keratinocyte/epidermal differentiation and intermediate filament reorganization with antimicrobial peptide induction (e.g., S100A7, DCD)
- Pronounced complement/coagulation and inflammatory signaling with EMT/angiogenesis hallmarks and increased secreted/stromal factors (e.g., F5, C9, POSTN, ANGPTL8, NRP2, TNFRSF12A, PLA2G2D)
- Concurrent enrichment of HYPOXIA and OXIDATIVE_PHOSPHORYLATION/TCA programs, suggesting a mixed stress–metabolic activation state
- mTORC1 pathway hallmark enrichment despite strong DDIT4 (REDD1) induction, implying noncanonical or dynamic mTOR regulation
- Lipid/adipogenic and xenobiotic metabolism program activation with increases in lipid-handling proteins (e.g., APOC3, ANGPTL8)

## Follow-up hypotheses
### Hypothesis 1
- **Hypothesis:** Clomiphene enforces a G1/S blockade leading to cytostasis, with a potential apoptotic component.
- **Why interesting:** Broad suppression of E2F/G2M and DNA replication/HR with apoptosis hallmark enrichment suggests growth arrest with possible cell death contribution.
- **Suggested validation:** Time-course flow cytometry for cell-cycle phase distribution (PI/DAPI), EdU/BrdU incorporation, pRB and E2F target immunoblots; Annexin V/PI staining, cleaved PARP/caspase assays; viability measurements.
- **Priority:** high

### Hypothesis 2
- **Hypothesis:** Up-regulated IGFBP1/2/3/5 attenuate IGF1R signaling, contributing to growth restraint.
- **Why interesting:** Coordinated IGFBP induction and enrichment of IGF-related processes plausibly reduce bioavailable IGF and downstream AKT/ERK signaling.
- **Suggested validation:** Measure secreted IGFBPs (ELISA) and IGF1R phosphorylation; assess pAKT/pERK; rescue experiments with excess IGF-I/II or IGFBP neutralization; evaluate proliferation with/without IGF supplementation.
- **Priority:** high

### Hypothesis 3
- **Hypothesis:** Treatment induces a keratinocyte-like epithelial differentiation program.
- **Why interesting:** Strong enrichment of keratinocyte/epidermis development and intermediate filament processes with up-regulated S100A7 and DCD indicates differentiation toward a squamous phenotype.
- **Suggested validation:** qPCR/IF for keratin markers (e.g., KRT1/10/14), involucrin/filaggrin; morphological assessment; barrier/differentiation assays; test reversibility upon drug washout.
- **Priority:** medium

### Hypothesis 4
- **Hypothesis:** A pro-inflammatory/coagulation secretory program is induced and may exert paracrine effects on stromal/endothelial cells.
- **Why interesting:** Elevated complement/coagulation and matrix/angiogenic factors (F5, C9, POSTN, ANGPTL8, TNFRSF12A) with TNFA/NF-kB enrichment suggest a remodeling secretome.
- **Suggested validation:** Secretome profiling (conditioned media proteomics/ELISA); NF-κB reporter assays; co-culture with fibroblasts/endothelial cells to test migration/angiogenic responses; EMT marker panel and migration assays.
- **Priority:** medium

### Hypothesis 5
- **Hypothesis:** Clomiphene triggers hypoxia/DDIT4 signaling while maintaining or enhancing mitochondrial respiration.
- **Why interesting:** HYPOXIA hallmark and DDIT4 up-regulation co-occur with OXPHOS/TCA enrichment, implying a mixed stress–metabolic state.
- **Suggested validation:** Seahorse assays (OCR/ECAR), HIF1α stabilization, mitochondrial membrane potential; quantify OXPHOS complex subunits; time-course to resolve dynamics.
- **Priority:** medium

### Hypothesis 6
- **Hypothesis:** mTORC1 activity is dynamically modulated despite REDD1 induction.
- **Why interesting:** mTORC1 hallmark enrichment conflicts with REDD1 up-regulation, suggesting temporal or feedback regulation of mTOR signaling.
- **Suggested validation:** Immunoblots for p-S6 (S235/236) and p-4EBP1 over a time course; rapamycin/torin sensitivity; assess Rheb/TSC signaling status.
- **Priority:** medium

## Confidence
medium

## Caution notes
- Conclusions are based on proteomics at a single time point; key mechanistic inferences (mTOR activity, EMT status, cytostatic vs cytotoxic balance) require orthogonal validation (phospho-signaling, apoptosis/proliferation and migration assays).
- Many elevated proteins are secreted/plasma-associated (e.g., IGFBPs, F5, C9, APOC3, HBA1); interpretation of ‘secretory programs’ should consider sample preparation, potential serum/extracellular carryover, and distinguish cell-associated proteome from secretome.
- Hallmark EMT enrichment may reflect extracellular matrix/secreted components rather than a true EMT transition; dedicated adhesion/migration markers are not provided.
- Classical estrogen-response signatures are not apparent; on-target ER engagement by clomiphene is not established by these data alone.
- Co-enrichment of hypoxia and oxidative phosphorylation/TCA pathways may reflect temporal dynamics or mixed stress states; directionality of metabolic effects is uncertain without functional assays.
- The prominent keratinocyte/skin differentiation signature indicates a context-specific epithelial model; generalization to other cell types should be cautious.


---

# Cobicistat

# Drug summary: Cobicistat

## Executive summary
Cobicistat exposure produced a coherent proteomic shift from proliferation toward a metabolically adaptive state. Proliferative programs (E2F/MYC targets, G2M checkpoint) and biosynthetic machinery (RNA processing/splicing, rRNA/ribosome biogenesis, chromatin/heterochromatin organization) were strongly repressed. In parallel, sterol/cholesterol homeostasis and steroid/sterol biosynthesis were robustly induced, consistent with the drug’s metabolic enzyme/CYP classification. Coordinated increases in lipid handling and nutrient transport (amino acids, fatty acids, monocarboxylates, organic anions) were observed. An inflammatory/secretory tilt was suggested by IL6–JAK–STAT3 and coagulation hallmarks, with AGR2 and FGB up. Secondary features included EMT/angiogenesis and Hedgehog hallmark enrichment. Key upregulated proteins supporting these programs included SC5D, NPC1L1, CYP4F2, CYP19A1, TXNIP, SLC16A13, NRP1, CEMIP, DLK1; notable downregulated proteins included ID3, AMD1, SHQ1, FANCM, and UBE2B. mTORC1 enrichment was modest and should be considered suggestive given concurrent suppression of ribosome biogenesis. Reactome/KEGG sections were uninformative here; conclusions rely on strong Hallmark/GO evidence and significant proteins.

## Findings consistent with expected biology
- Activation of sterol/cholesterol metabolism and lipid-handling programs (e.g., SC5D, NPC1L1, CYP4F2 up; strong CHOLESTEROL_HOMEOSTASIS and sterol biosynthesis enrichments).
- Induction of steroid metabolism enzymes with estrogen-response features (CYP19A1 up with ESTROGEN_RESPONSE_LATE hallmark).

## Unexpected but plausible findings
- Marked suppression of proliferation (E2F/MYC targets, G2M) with broad downregulation of RNA processing and ribosome biogenesis (e.g., ID3, AMD1, SHQ1, FANCM down).
- Induction of IL6–JAK–STAT3 signaling and a coagulation-leaning/secretory shift (AGR2 and FGB up), suggesting stress-responsive paracrine changes (moderate evidence).
- Enhanced nutrient transport capacity across multiple modalities (amino acids, fatty acids, monocarboxylates, organic anions), with SLC16A13 up and strong GO transport enrichments.
- EMT/angiogenesis and Hedgehog hallmark enrichment with limited direct markers (NRP1, CEMIP, DLK1 up; LGR4 and ZNRF3 up), considered secondary/context-dependent.
- Concurrent elevation of cholesterol uptake and biosynthesis (NPC1L1 and SC5D up), indicating a coordinated cholesterol demand state.
- Metabolic stress indicators (TXNIP up) with a shift in fatty acid/peroxisomal metabolism (AMACR down).

## Follow-up hypotheses
### Hypothesis 1
- **Hypothesis:** Cobicistat triggers SREBP-driven activation of cholesterol biosynthesis and uptake, increasing cellular cholesterol demand and steroidogenic enzyme expression.
- **Why interesting:** Links the drug’s metabolic enzyme/CYP classification to membrane lipid homeostasis and transporter remodeling; may influence drug transport, signaling microdomains, and cell adaptation.
- **Suggested validation:** Assess SREBP1/2 cleavage and target gene/protein levels; quantify cellular cholesterol and sterol intermediates; test sensitivity to NPC1L1 inhibition (e.g., ezetimibe) and statins; perform 13C-acetate/cholesterol flux tracing.
- **Priority:** high

### Hypothesis 2
- **Hypothesis:** The treatment enforces cell-cycle exit with a stress-associated secretory phenotype partly mediated by IL6–JAK–STAT3 signaling.
- **Why interesting:** Explains the strong anti-proliferative signature and coagulation/secretory markers; may impact combination strategies and cell-state transitions independent of the drug’s enhancer role.
- **Suggested validation:** Measure cell-cycle markers (EdU incorporation, pRB phosphorylation, Cyclin E/A), STAT3 phosphorylation and target genes, and secreted factors (IL6, fibrinogen) by ELISA; test reversibility after washout.
- **Priority:** high

### Hypothesis 3
- **Hypothesis:** Upregulated amino acid/monocarboxylate transport reflects enhanced nutrient uptake under metabolic stress rather than canonical mTORC1 activation.
- **Why interesting:** Reconciles transporter enrichment with suppressed ribosome biogenesis; has implications for metabolic dependencies and sensitivity to nutrient limitation.
- **Suggested validation:** Profile phospho-S6K and phospho-4EBP1; quantify amino acid and lactate/pyruvate uptake; assess TXNIP–thioredoxin redox state; test response to rapamycin/torin and amino acid starvation.
- **Priority:** medium

### Hypothesis 4
- **Hypothesis:** Increased CYP19A1 contributes to an estrogen-response program in lipid-rewired cells, potentially altering paracrine signaling.
- **Why interesting:** Connects steroidogenesis to observed estrogen-response enrichment and could modulate downstream transcriptional networks.
- **Suggested validation:** Measure estradiol and related steroids; assess ER target gene expression and ER phosphorylation; test sensitivity to aromatase inhibition.
- **Priority:** medium

## Confidence
high

## Caution notes
- mTORC1 and Hedgehog conclusions are primarily pathway-level and lack multiple direct effector proteins in the top hits; treat as suggestive pending targeted or phospho-proteomic validation.
- mTORC1 hallmark up co-occurs with strong suppression of ribosome biogenesis and RNA processing, arguing against assuming canonical mTORC1 activation.
- Inflammatory interpretations beyond IL6–JAK–STAT3 enrichment (e.g., ‘antiviral-like’ patterns) are weakly supported by individual proteins; confirm with cytokine/STAT target panels.
- Transporter activation is largely pathway-driven with few individual transporters among top changes; validate with broader transporter assays or transcriptomics.
- EMT/angiogenesis signals have modest FDRs and limited marker support; consider context-dependent and secondary.
- Lack of Reactome/KEGG enrichment here does not negate the biology; primary evidence is from strong Hallmark and GO signals and protein-level changes.


---

# Cromolyn (sodium)

# Drug summary: Cromolyn (sodium)

## Executive summary
Proteomic profiling of Cromolyn (sodium) indicates a robust structural reprogramming characterized by strengthened cell–cell adhesion programs and suppression of intermediate filament/keratin processes. This is supported by strong GO enrichments (cell–cell adhesion up; intermediate filament organization down) and coherent protein changes (EFNB1, CD9, TSPAN15 up; KRT79, FLG2, CLDN12, S100A7 down). Hallmark enrichments for HYPOXIA and IL6/JAK/STAT3 are significant but have limited direct protein corroboration (e.g., NDUFA3 down; IGFBP3 up), suggesting a secondary stress-like response rather than confirmed pathway activation. A cholesterol homeostasis Hallmark signal is present without prominent lipid-metabolism proteins among top hits, implying possible membrane/lipid remodeling accompanying the adhesion shift. Notably, multiple histone H2A variants and macroH2A1/2 are increased, consistent with altered chromatin state or cell-cycle/nuclear content changes. The IGF axis appears rebalanced (IGFBP3 up; IGFBP2 down). No clear enrichment is observed for PI3K/Akt/mTOR or Wnt pathways despite the drug’s annotated context. Reactome/KEGG sections are uninformative and one Hallmark-down section is empty; conclusions are therefore based on the strongest GO/Hallmark evidence and differentially abundant proteins.

## Findings consistent with expected biology
- Enhanced cell–cell adhesion programs with concurrent attenuation of intermediate filament/keratin networks, consistent with downstream consequences of membrane/ion channel perturbation (GO: adhesion up; intermediate filament organization down; EFNB1, CD9, TSPAN15 up; KRT79, FLG2 down).
- Reduced epithelial barrier-associated and select ECM/glycocalyx components (e.g., FLG2, CLDN12, S100A7, COL6A1 down) aligning with the broader structural shift.

## Unexpected but plausible findings
- Significant Hallmark enrichments for HYPOXIA and IL6/JAK/STAT3 signaling with modest protein-level support (NDUFA3 down; IGFBP3 up), indicative of a stress-like response.
- Marked increases in histone H2A variants and macroH2A1/2, suggesting altered chromatin state or cell-cycle/nuclear content changes at the protein level.
- Reciprocal IGF-binding protein changes (IGFBP3 up; IGFBP2 down), pointing to a shift in IGF-axis regulation concurrent with stress signatures.
- Cholesterol homeostasis Hallmark enrichment without strong direct lipid-metabolism protein corroboration, implying possible membrane/lipid remodeling.
- MYOGENESIS Hallmark enrichment at modest significance, likely reflecting overlap with cytoskeletal/ECM programs and requiring cautious interpretation.
- Upregulation of ZNRF3 (a Wnt antagonist) in the absence of Wnt pathway enrichment; this single-protein observation suggests but does not establish Wnt tone dampening.

## Follow-up hypotheses
### Hypothesis 1
- **Hypothesis:** Cromolyn increases junctional cell–cell adhesion while weakening keratin/intermediate filament networks.
- **Why interesting:** Strongest and most coherent signature in the dataset (GO adhesion up; intermediate filament organization down) with multiple concordant proteins.
- **Suggested validation:** Measure cell–cell adhesion (cell aggregation assays, TEER if epithelial); immunostain and quantify junctional markers (e.g., cadherins) and keratins; assess barrier function and cytoskeletal organization.
- **Priority:** high

### Hypothesis 2
- **Hypothesis:** Cromolyn elicits a secondary hypoxia-like and IL6/STAT3 stress response.
- **Why interesting:** Significant Hallmark enrichments with supportive but limited protein evidence (NDUFA3 down; IGFBP3 up) suggest a coordinated stress program.
- **Suggested validation:** Assess HIF1α stabilization and HIF target proteins; oxygen consumption and glycolytic flux; STAT3 phosphorylation and nuclear localization; cytokine measurements (e.g., IL6).
- **Priority:** medium

### Hypothesis 3
- **Hypothesis:** Cromolyn alters chromatin composition, increasing histone H2A variants and macroH2A content.
- **Why interesting:** Multiple histone variants are up, indicating a shift in chromatin packaging or cell-cycle state.
- **Suggested validation:** Chromatin fractionation and immunoblotting for H2A variants/macroH2A; ATAC-seq or nuclease sensitivity; monitor proliferation markers to distinguish cell-cycle effects.
- **Priority:** medium

### Hypothesis 4
- **Hypothesis:** Cromolyn induces membrane/lipid remodeling consistent with increased cholesterol homeostasis signaling.
- **Why interesting:** Cholesterol homeostasis Hallmark enrichment and increases in tetraspanins (CD9, TSPAN15) suggest changes in membrane microdomains.
- **Suggested validation:** Untargeted lipidomics and filipin staining for free cholesterol; expression of SREBP target proteins; membrane order assays (e.g., Laurdan GP).
- **Priority:** medium

### Hypothesis 5
- **Hypothesis:** Cromolyn rebalances the IGF axis toward IGFBP3 dominance, modulating IGF bioavailability.
- **Why interesting:** Reciprocal IGFBP3 up and IGFBP2 down align with stress signatures and could impact growth signaling.
- **Suggested validation:** Quantify IGFBP2/3 in media and cells (ELISA/immunoblot); assess IGF pathway readouts (pAKT) with and without exogenous IGF; monitor proliferation/survival.
- **Priority:** medium

### Hypothesis 6
- **Hypothesis:** Cromolyn dampens basal Wnt signaling via ZNRF3 upregulation.
- **Why interesting:** ZNRF3 is a Wnt antagonist; its increase may reduce Wnt tone, although support is currently limited to a single protein.
- **Suggested validation:** Wnt reporter assays (TOPFlash), β-catenin levels/localization, and Wnt target protein measurements under basal and Wnt-stimulated conditions.
- **Priority:** low

## Confidence
medium

## Caution notes
- Pathway enrichments from proteomic data indicate coordinated trends but do not prove pathway activation; interpret HYPOXIA and IL6/STAT3 signals as stress-like unless validated functionally.
- Increased histone variants may reflect cell-cycle state or nuclear content changes rather than targeted epigenetic remodeling; additional chromatin-focused assays are required.
- Cholesterol homeostasis enrichment is pathway-driven with limited direct lipid-protein support; corroboration with lipidomics is recommended.
- Conclusions about ECM/glycocalyx reduction and mitochondrial/vesicular components are based on small protein sets and should be considered low-confidence until replicated.
- Labeling the adhesion shift as ‘homophilic’ derives from GO gene sets; canonical cadherins/protocadherins are not prominent among top changes in this dataset.
- No clear enrichment for PI3K/Akt/mTOR or Wnt pathways despite the drug’s annotated context; avoid over-connecting findings to these pathways without broader evidence.
- The MYOGENESIS Hallmark enrichment likely reflects overlap with structural programs and should be interpreted cautiously.
- Cell-type context is not specified; proteins such as FLG2, KRT79, and S100A7 can be tissue-specific, which may affect generalizability.
- Some pathway sections are empty (e.g., Hallmark down; Reactome/KEGG), limiting pathway-level interpretability but not negating the observed GO/Hallmark signals.


---

# Cyclothiazide

# Drug summary: Cyclothiazide

## Executive summary
Cyclothiazide exposure produced a proteomic profile dominated by up-regulation of intermediate filament organization (GO BP enrichment: NES ~2.23; FDR 0). In parallel, multiple membrane trafficking/modification components (SNX29, VPS26C, NDFIP1, ZDHHC3, PIGW) and transcriptional/chromatin regulators (JUNB, SETD1B, HDAC10, PRDM2, CNOT6) increased, consistent with adaptive remodeling of membrane protein handling and gene regulation. Modest increases in ERN1 (IRE1) and PPP1R15B suggest a potential proteostasis/ER stress–related adjustment, but this is not supported by broader pathway enrichment or canonical markers and should be considered provisional. Down-regulated proteins were limited and focal (e.g., IL10RB, TNFRSF12A, AQP11, SLC35F5, SGCE) without coherent pathway-level enrichment. Overall, the strongest and most reliable signal is cytoskeletal reorganization centered on intermediate filaments, with supportive but tentative evidence for enhanced membrane processing and mild ER stress/ISR engagement.

## Findings consistent with expected biology
- Elevated membrane trafficking/modification machinery (SNX29, VPS26C, NDFIP1, ZDHHC3, PIGW), consistent with adaptive handling of membrane/ion channel proteins.
- Up-regulation of transcriptional/chromatin regulators (JUNB, SETD1B, HDAC10, PRDM2, CNOT6), indicative of broader transcriptional remodeling.

## Unexpected but plausible findings
- Strong enrichment for intermediate filament organization and intermediate filament–based processes (GO BP NES ~2.23; FDR 0), indicating pronounced cytoskeletal remodeling biased toward intermediate filaments.
- Concurrent increases in ERN1 (IRE1) and PPP1R15B, suggestive of mild ER stress/ISR-related proteostasis adjustments despite absence of pathway-level UPR enrichment.
- Decreases in select cytokine/transport-associated proteins (IL10RB, TNFRSF12A, AQP11, SLC35F5, SGCE) without coherent down-regulated pathway enrichment.

## Follow-up hypotheses
### Hypothesis 1
- **Hypothesis:** Cyclothiazide induces intermediate filament remodeling that stabilizes the membrane–cytoskeleton interface during altered ion channel activity.
- **Why interesting:** Links an iGluR-targeting compound to structural adaptations that could influence receptor localization, surface stability, and cellular mechanics.
- **Suggested validation:** Identify and quantify specific intermediate filament proteins driving the enrichment (e.g., vimentin, keratins, GFAP as appropriate); perform time-course immunofluorescence and detergent-solubility assays to assess filament organization; test dependence on iGluR activity with antagonists or ion flux blockers.
- **Priority:** high

### Hypothesis 2
- **Hypothesis:** Cyclothiazide engages a mild IRE1/ISR-mediated adaptive proteostasis response that adjusts secretory/membrane protein handling.
- **Why interesting:** Proteostasis engagement could modulate receptor maturation/trafficking and may have implications for efficacy and safety.
- **Suggested validation:** Assess canonical UPR/ISR markers (HSPA5/BiP, XBP1 splicing, ATF4, DDIT3/CHOP, phospho-eIF2α); evaluate ERN1 phosphorylation/activation; probe PPP1R15A/B dynamics; use ER stress modulators to test reversibility.
- **Priority:** medium

### Hypothesis 3
- **Hypothesis:** Cyclothiazide enhances endosomal/retromer-dependent sorting and palmitoylation processes that regulate surface abundance of membrane proteins, including iGluRs.
- **Why interesting:** Coordinated increases in VPS26C, SNX29, NDFIP1, ZDHHC3, and PIGW suggest adjusted trafficking and post-translational processing that could directly affect receptor surface levels.
- **Suggested validation:** Monitor retromer/SNX complex localization and abundance (VPS35/26, SNX29); knock down VPS26C/SNX29 and assess effects on surface levels of target receptors by biotinylation or flow cytometry; quantify palmitoylation status of relevant substrates.
- **Priority:** medium

### Hypothesis 4
- **Hypothesis:** Cyclothiazide modestly attenuates selected cytokine receptor nodes (e.g., IL10RB, TNFRSF12A) as a secondary effect.
- **Why interesting:** May alter inflammatory responsiveness or crosstalk with neuronal signaling in specific cell contexts.
- **Suggested validation:** Confirm protein and mRNA changes for IL10RB and TNFRSF12A; perform ligand-stimulated signaling assays (e.g., STAT3, NF-κB readouts); assess cell-type specificity and off-target dependence.
- **Priority:** low

## Confidence
medium

## Caution notes
- Intermediate filament enrichment is robust, but the specific proteins driving the signal are not listed among top hits; identifying them is important for mechanistic clarity.
- UPR/ISR inference relies on ERN1 and PPP1R15B without pathway-level enrichment or canonical markers; interpret as provisional.
- Characterizing an immediate-early–like response based primarily on JUNB is under-supported; avoid strong claims without additional IEGs or enrichment.
- Down-regulated proteins are few and lack coherent pathway enrichment; avoid broad functional claims regarding cytokine or transport pathways.
- Proteomics alone cannot assess activation states (e.g., phosphorylation) or transcriptional dynamics; complementary assays are needed.
- No Hallmark/Reactome/KEGG enrichments were detected; conclusions are based on GO BP and individual protein changes.
- Links to neuronal signaling context are indirect in this dataset; absence of explicit neuronal pathway enrichment should be acknowledged.


---

# Cytidine

# Drug summary: Cytidine

## Executive summary
Cytidine exposure produced a robust and coherent suppression of epidermal/keratinocyte differentiation and intermediate filament organization programs at the proteome level. Multiple GO biological processes related to keratin networks and epidermal development were significantly downregulated, concordant with decreases in keratin-associated proteins (e.g., HRNR, KRT9, KRT79, DCD). On the up side, enrichment was limited: a modest Hallmark KRAS_SIGNALING_UP signal was detected alongside increased abundance of select receptor/adaptor and extracellular milieu–associated proteins (e.g., SPRED1, LOXL2, IGFBP5, LGR4, GRAP, CEMIP). Several signaling and transcriptional regulators (PTEN, BCL2L2, KLF6, HIVEP1) were reduced, but without pathway-level enrichment. No significant up-enriched GO terms or down-enriched Hallmark sets were observed. Expected cell cycle/DNA damage or nucleotide metabolism signatures were not detected under these conditions, suggesting context-, dose-, or time–dependence for canonical nucleoside antimetabolite effects. Overall, the evidence strongly supports a shift away from an epidermal/keratinocyte-like differentiated state, with only tentative indications of upstream growth factor/MAPK involvement.

## Findings consistent with expected biology
- None listed.

## Unexpected but plausible findings
- Strong, coordinated repression of keratin/epidermal differentiation and intermediate filament organization programs, supported by multiple significant GO BP down-regulations and decreases in keratin-associated proteins (HRNR, KRT9, KRT79, DCD).
- Modest enrichment of a KRAS/MAPK-related signature (Hallmark KRAS_SIGNALING_UP) together with increased SPRED1 and selected receptor/ECM-associated proteins (LOXL2, IGFBP5, LGR4, GRAP, CEMIP); interpreted as a tentative signal rather than definitive pathway activation.
- Decreases in several regulators (PTEN, BCL2L2, KLF6, HIVEP1) suggest broader regulatory shifts, but without corroborating pathway-level enrichment; these changes should be considered supportive observations only.

## Follow-up hypotheses
### Hypothesis 1
- **Hypothesis:** Cytidine shifts cells away from an epidermal/keratinocyte-like differentiated state toward a less differentiated phenotype.
- **Why interesting:** Explains the concordant suppression of keratin/intermediate filament processes and epidermal differentiation terms with matching protein decreases, indicating a state change with potential functional consequences for barrier/cytoskeletal properties.
- **Suggested validation:** Measure mRNA/protein markers of keratinocyte differentiation (e.g., KRT1/9/10, IVL, FLG) by qPCR and immunoblot/immunofluorescence; assess keratin network organization via microscopy; perform time-course proteomics to determine onset and reversibility.
- **Priority:** high

### Hypothesis 2
- **Hypothesis:** A weak engagement of growth factor–RAS/MAPK signaling contributes to the observed state shift.
- **Why interesting:** Consistent with modest KRAS hallmark enrichment and upregulation of SPRED1 and selected receptor/ECM-associated proteins that can interface with MAPK signaling, suggesting upstream cueing may be involved.
- **Suggested validation:** Phospho-proteomics or immunoblot time courses for p-ERK, p-MEK, and p-EGFR; test sensitivity to MEK/ERK inhibitors and assess whether inhibition attenuates the keratin differentiation suppression; evaluate feedback markers (SPRED1) dynamics.
- **Priority:** medium

### Hypothesis 3
- **Hypothesis:** Cytidine induces extracellular matrix remodeling that accompanies or facilitates the phenotypic shift.
- **Why interesting:** Upregulation of LOXL2, CEMIP, IGFBP5, and LGR4 suggests altered cell–matrix interactions that can impact differentiation and signaling.
- **Suggested validation:** Secretome analysis; assays for collagen cross-linking (LOXL2 activity) and hyaluronan turnover (CEMIP); adhesion/migration assays; evaluate whether ECM-targeted inhibitors or knockdowns modulate the observed proteomic changes.
- **Priority:** medium

### Hypothesis 4
- **Hypothesis:** Modest PTEN reduction biases signaling cross-talk toward MAPK/PI3K pathways without overt pathway activation.
- **Why interesting:** PTEN decrease co-occurs with a KRAS hallmark signal and other regulatory changes, raising the possibility of altered signaling tone even in the absence of strong pathway enrichment.
- **Suggested validation:** Quantify p-AKT/p-S6 and p-ERK levels; rescue PTEN via overexpression or stabilize PTEN and assess impact on differentiation markers; probe sensitivity to PI3K/mTOR inhibitors alone and in combination with MEK inhibitors.
- **Priority:** low

## Confidence
medium

## Caution notes
- Protein abundance changes do not establish pathway activity; inferences about MAPK/PI3K engagement require phospho-proteomic or functional activity readouts.
- Cell-type context is not specified; the strong repression of keratinocyte/epidermal programs suggests a keratinocyte-like baseline, but generalization to other cell types is uncertain.
- Upstream signaling and ECM remodeling interpretations are based on a limited set of upregulated proteins without corroborating GO/Reactome/KEGG up-enrichment and should be considered tentative.
- Single-protein changes (e.g., PTEN) should not be used to anchor pathway-level conclusions, especially with modest effect sizes.
- No enrichment for expected cell cycle/DNA damage pathways was observed; this absence may reflect dose, timing, or model context rather than a lack of effect.
- Empty up-enrichment categories (GO_BP_up, Hallmark_down) indicate a lack of statistically significant findings in those sets and should not be over-interpreted.


---

# Danthron

# Drug summary: Danthron

## Executive summary
Proteomics after Danthron exposure shows a robust hypoxia-like response and pro-inflammatory activation. Hallmark HYPOXIA (NES 2.17, FDR 0) and TNFA_SIGNALING_VIA_NFKB/INFLAMMATORY_RESPONSE (NES ~1.98/1.76, FDR ~6e-4/0.012) are strongly enriched, with protein support from PFKM and IGFBP1 (hypoxia/energy stress) and TRAF3, ISG20, CD68 (inflammatory). Moderate enrichment of TGF-β/EMT accompanies increases in ECM/hyaluronan remodeling enzymes (ADAM15, CEMIP, CHSY1, HS6ST1), suggesting partial mesenchymal/matrix-remodeling features but lacking canonical EMT marker shifts. Decreases in HMGCR and selected mitochondrial components (e.g., NDUFA3, DNAJC30) point to reduced sterol synthesis and possible respiratory attenuation, though coverage is limited. CYP1A1 is strongly repressed, consistent with perturbed xenobiotic signaling within an inflammatory context. Selective reductions in HLA-B, IL4R, and TJP3 indicate potential immunomodulatory and junctional changes that diverge from the broader inflammatory signature. Overall, the pattern supports energy- and inflammation-driven reprogramming compatible with AMPK–mTOR/autophagy crosstalk, but direct autophagy/AMPK readouts are not evident. Pathway interpretation relies primarily on Hallmark sets; other pathway resources did not return significant results.

## Findings consistent with expected biology
- Robust hypoxia-like program (Hallmark HYPOXIA NES 2.17, FDR 0) with supporting protein increases (PFKM, IGFBP1), consistent with energy-stress biology.
- Pro-inflammatory signaling enrichment (TNFA_SIGNALING_VIA_NFKB and INFLAMMATORY_RESPONSE up; TRAF3, ISG20, CD68 increased) indicating activation of inflammatory pathways.
- Repression of xenobiotic metabolism marker CYP1A1 (strongly down), compatible with an inflammatory milieu and altered biotransformation.
- Evidence consistent with suppressed mevalonate/cholesterol synthesis (HMGCR down), although broader pathway coverage is limited.

## Unexpected but plausible findings
- Moderate TGF-β and EMT signatures (borderline FDRs) with ECM/hyaluronan remodeling proteins up (ADAM15, CEMIP, CHSY1, HS6ST1) despite absence of canonical EMT marker shifts; interpret as partial/early remodeling.
- Selective downregulation of immune/junction components (HLA-B, IL4R, TJP3) concurrent with inflammatory hallmark activation, suggesting pathway-specific immune modulation.
- Increases in centrosome/kinetochore-associated proteins (CEP97, CEP290, SKA2) without cell-cycle hallmark enrichment, indicating a noncanonical structural or mitotic-checkpoint–related signal.
- Decreases in selected mitochondrial-associated proteins (NDUFA3, DNAJC30), suggestive of respiratory attenuation but not demonstrative of a global OXPHOS suppression.

## Follow-up hypotheses
### Hypothesis 1
- **Hypothesis:** Danthron induces an energy-stress axis that stabilizes HIF and shifts metabolism toward glycolysis while curtailing sterol synthesis via AMPK–mTOR crosstalk.
- **Why interesting:** Integrates strong HYPOXIA enrichment, PFKM/IGFBP1 increases, and HMGCR downregulation with the drug’s putative AMPK/autophagy linkage; provides a unifying mechanism for metabolic reprogramming.
- **Suggested validation:** Measure p-AMPK (Thr172), p-ACC (Ser79), and mTORC1 readouts (p-S6K, p-4E-BP1); assess HIF stabilization; perform Seahorse ECAR/OCR and lactate/glucose uptake assays; quantify sterol biosynthesis (SREBP2 processing, lathosterol/cholesterol levels); evaluate autophagy markers (LC3-II, p62) and flux.
- **Priority:** high

### Hypothesis 2
- **Hypothesis:** NF-κB pathway activation contributes to suppression of AhR-driven xenobiotic genes, explaining CYP1A1 downregulation.
- **Why interesting:** Links strong TNF/NF-κB hallmarks with pronounced CYP1A1 decrease, clarifying the interplay between inflammatory signaling and biotransformation.
- **Suggested validation:** Test NF-κB activation (p65 nuclear translocation, IκBα degradation, NF-κB reporter); measure cytokine/NF-κB target transcripts; assess AhR activity (XRE reporter) and CYP1A1 mRNA/protein ± NF-κB inhibitors or TNF blockade; evaluate rescue with an AhR agonist.
- **Priority:** medium

### Hypothesis 3
- **Hypothesis:** Danthron promotes ECM remodeling and a partial EMT-like state via TGF-β signaling.
- **Why interesting:** Explains enrichment of TGF-β/EMT hallmarks and upregulation of ADAM15, CEMIP, CHSY1, and HS6ST1; may impact adhesion and invasiveness.
- **Suggested validation:** Assess p-SMAD2/3 and TGF-β reporter activity; profile EMT markers (E-cadherin, vimentin, SNAI1/ZEB1); perform migration/invasion assays; quantify hyaluronan content/fragmentation; test sensitivity to TGF-β receptor inhibition.
- **Priority:** medium

### Hypothesis 4
- **Hypothesis:** Inflammatory activation coexists with selective downregulation of antigen presentation and epithelial junction components.
- **Why interesting:** The combination of HLA-B, IL4R, and TJP3 decreases alongside inflammatory hallmarks suggests pathway-specific immunomodulation with potential relevance to immune visibility and barrier function.
- **Suggested validation:** Measure surface HLA class I and IL4R by flow cytometry and immunoblot; confirm transcripts; examine barrier properties (TEER) where applicable; test cytokine responsiveness (e.g., IFN-γ, IL-4) and evaluate cell-type dependence across models.
- **Priority:** medium

## Confidence
medium

## Caution notes
- Pathway interpretation relies primarily on Hallmark gene sets; GO/Reactome/KEGG sections were not significantly enriched and are absent.
- Glycolytic shift is suggested by HYPOXIA enrichment and PFKM/IGFBP1 but lacks broader enzyme coverage; metabolomics is recommended.
- Mevalonate/cholesterol suppression inference is based mainly on HMGCR; additional enzymes and sterol measurements are needed.
- Mitochondrial attenuation is inferred from a small number of proteins (e.g., NDUFA3, DNAJC30) and does not establish global OXPHOS changes.
- TGF-β/EMT signals have borderline FDRs and lack canonical EMT marker changes; interpret as moderate confidence.
- NF-κB activation is inferred from enrichment rather than direct activity measurements; phospho- and localization assays are needed.
- Single-protein decreases (HLA-B, IL4R, TJP3) should be validated across replicates and conditions; effects may be cell-type specific.
- CYP1A1 repression may reflect interplay with inflammatory signaling or culture conditions and is not necessarily a drug-specific effect.
- Cellular context (cell type, dose, time) is not specified; proteins like CD68 and HLA-B can be context-dependent.


---

# Daprodustat

# Drug summary: Daprodustat

## Executive summary
Proteomics after Daprodustat exposure shows clear on‑target HIF pathway engagement with a robust Hallmark HYPOXIA signature (NES ~2.12, FDR 0) and upregulation of canonical hypoxia-associated proteins (e.g., CA9, JUNB, EFNA1). Pathway-level enrichment supports a shift toward glycolysis (Hallmark GLYCOLYSIS), consistent with hypoxic metabolic adaptation. An additional enrichment of cholesterol/lipid homeostasis is observed, suggesting lipid remodeling under HIF activation, though direct protein-level coverage is limited. In parallel, multiple GO terms indicate broad downregulation of RNA splicing, rRNA processing, and ribosome biogenesis, as well as intermediate filament/cytoskeletal and epidermal differentiation programs. Notable single-protein changes include strong TXNIP upregulation and decreases in the citrate transporter SLC13A5 and ECM component COL6A1, which plausibly align with the metabolic and structural remodeling indicated by pathway results. Sections with no reported enrichments (e.g., GO BP up, Hallmark down) are treated as absence of significant signal rather than contradiction. Overall, the data are most consistent with HIF stabilization driving hypoxic adaptation, accompanied by reduced RNA processing/translation machinery and potential lipid metabolic adjustments.

## Findings consistent with expected biology
- On‑target HIF pathway activation: Hallmark HYPOXIA enrichment with upregulation of CA9 and other hypoxia-associated proteins (e.g., JUNB, EFNA1).
- Metabolic adaptation toward glycolysis at the pathway level (Hallmark GLYCOLYSIS enrichment).

## Unexpected but plausible findings
- Upregulation of cholesterol/lipid homeostasis at the pathway level (Hallmark CHOLESTEROL_HOMEOSTASIS), suggesting lipid remodeling under HIF activation, but with limited direct protein exemplars.
- Broad suppression of RNA splicing, rRNA processing, and ribosome biogenesis programs, indicating a decrease in RNA processing/translation machinery.
- Downregulation of intermediate filament organization and epidermal/keratinocyte differentiation gene sets, likely reflecting cytoskeletal/differentiation state changes rather than specific cell identity.
- Marked TXNIP upregulation and single-protein decreases in SLC13A5 and COL6A1, consistent with nuanced glucose/redox control, altered citrate handling, and ECM remodeling, respectively (tentative at the single-protein level).

## Follow-up hypotheses
### Hypothesis 1
- **Hypothesis:** Daprodustat increases glycolytic flux downstream of HIF stabilization.
- **Why interesting:** Confirms functional metabolic reprogramming underlying Hallmark GLYCOLYSIS enrichment and links pathway signal to flux.
- **Suggested validation:** Measure extracellular acidification rate and lactate production; quantify key glycolytic enzymes/transporters (e.g., HK2, PFKP, LDHA, SLC2A1) by targeted proteomics or immunoblot; perform time-course assays.
- **Priority:** high

### Hypothesis 2
- **Hypothesis:** HIF activation reduces RNA splicing/rRNA processing and ribosome biogenesis as part of an adaptive downshift in translational capacity.
- **Why interesting:** Explains coordinated downregulation of multiple RNA processing and ribosome biogenesis GO terms and may impact protein synthesis.
- **Suggested validation:** Assess global protein synthesis (e.g., puromycin/SUnSET), rRNA processing markers and nucleolar proteins, and spliceosome component abundance; corroborate with transcriptomics or ribosome profiling.
- **Priority:** medium

### Hypothesis 3
- **Hypothesis:** Daprodustat perturbs cellular cholesterol/lipid metabolism under HIF, altering sterol handling and membrane lipid composition.
- **Why interesting:** Hallmark CHOLESTEROL_HOMEOSTASIS enrichment and upregulation of select lipid-related proteins suggest coordinated lipid remodeling.
- **Suggested validation:** Perform untargeted and targeted lipidomics; quantify SREBP pathway components and cholesterol biosynthesis/uptake proteins (e.g., HMGCR, LDLR); test sensitivity to statins or lipid depletion.
- **Priority:** medium

### Hypothesis 4
- **Hypothesis:** Reduced SLC13A5 lowers citrate import, modulating cytosolic acetyl‑CoA availability and linking to observed lipid metabolism signals.
- **Why interesting:** Connects a strong single-protein decrease to potential changes in acetyl‑CoA and lipid biosynthesis.
- **Suggested validation:** Measure citrate uptake and intracellular citrate/acetyl‑CoA; test whether extracellular citrate supplementation or SLC13A5 overexpression rescues lipidomic changes; evaluate ACLY activity.
- **Priority:** low

## Confidence
high

## Caution notes
- Pathway-level enrichments, particularly GO BP down terms, should be corroborated by examining the breadth of contributing proteins beyond the top hits.
- Cell type and treatment time/dose are not specified; GO terms referencing epidermal/keratinocyte programs may reflect gene-set labels rather than true cell identity.
- Cholesterol/lipid remodeling conclusions require additional protein-level support and orthogonal lipidomics before being considered coordinated.
- TXNIP regulation is context- and time-dependent; its upregulation here should be validated in independent conditions or time courses.
- Single-protein observations (e.g., SLC13A5, COL6A1) are hypothesis-generating and should not be overgeneralized without corroborating pathway or multi-protein evidence.
- These are proteomics-only data; functional assays and/or transcriptomics are needed to confirm reduced biosynthetic load or shifts in metabolic flux.
- Empty pathway sections (e.g., GO BP up, Hallmark down) indicate no significant results were reported and should not be interpreted as contradictions.


---

# Deflazacort

# Drug summary: Deflazacort

## Executive summary
Deflazacort (glucocorticoid receptor agonist) produced a robust pathway-level suppression of RNA splicing, mRNA processing, and rRNA processing (multiple GO terms; NES ≈ -2; FDR < 0.05), indicating coordinated downregulation of RNA maturation machinery. Consistent with GR engagement, NR3C1 protein decreased (log2FC ~ -0.86). In contrast, no upregulated pathway enrichments were detected; however, individual proteins suggest a mitochondrial/redox adaptation (e.g., TOMM5, PET100, TK2, MSRA, MT1E) and selected transporter changes (FXYD1, SLC16A6, SLCO4A1). Extracellular signaling/ECM factors were mixed (CD109, FSTL3 up; DKK1, CCN2, SULF2, ITIH5 down), implying remodeling without a single coherent direction. Notable single-protein changes include PIEZO2 up and PTEN down, alongside strong induction of NAGS and MIXL1. These protein-level increases should be treated as hypotheses pending pathway-level validation. Overall, the data support GR activation with prominent suppression of RNA processing, plus suggestive metabolic and ECM/signaling adaptations, with medium confidence.

## Findings consistent with expected biology
- Feedback downregulation of the glucocorticoid receptor NR3C1 following GR agonism (log2FC ~ -0.86; FDR ~ 5e-06).

## Unexpected but plausible findings
- Broad suppression of RNA splicing and mRNA/rRNA processing programs (multiple GO BP terms, NES ~ -2.0; FDR < 0.05).
- Protein-level increases suggest mitochondrial import/respiratory assembly and redox adaptation (e.g., TOMM5, PET100, TK2, MSRA, MT1E), without pathway enrichment support.
- Mixed remodeling of extracellular signaling/ECM regulators (CD109, FSTL3 up; DKK1, CCN2, SULF2, ITIH5 down), indicating altered Wnt/TGF-beta/ECM tone without coherent enrichment.
- Upregulation of the mechanosensitive channel PIEZO2 (log2FC ~ 1.86; FDR ~ 0.008).
- Decrease of PTEN protein (log2FC ~ -0.62; FDR ~ 0.029).
- Strong induction of NAGS and MIXL1 (log2FC ~ 2.8 and ~ 2.7; significant FDR), unusual in many contexts.

## Follow-up hypotheses
### Hypothesis 1
- **Hypothesis:** Deflazacort suppresses core RNA splicing machinery, leading to reduced splicing capacity and altered isoform usage.
- **Why interesting:** Explains the dominant pathway-level downregulation and could impact broad gene expression and drug response phenotypes.
- **Suggested validation:** Perform RNA-seq with differential splicing analysis (e.g., rMATS), assess spliceosome component levels by immunoblot, and use splicing reporter assays.
- **Priority:** high

### Hypothesis 2
- **Hypothesis:** Treatment induces a mitochondrial/redox adaptation to maintain metabolic and oxidative homeostasis.
- **Why interesting:** Would link GR activation to metabolic resilience and potential liabilities under stress.
- **Suggested validation:** Measure mitochondrial respiration (OCR/ECAR), mitochondrial membrane potential, and ROS levels; confirm upregulation of TOMM5/PET100/TK2/MSRA/MT1E by qPCR/immunoblot; test sensitivity to mitochondrial stressors.
- **Priority:** medium

### Hypothesis 3
- **Hypothesis:** PTEN reduction increases PI3K/AKT signaling as a cross-talk with GR signaling.
- **Why interesting:** Could reshape survival and metabolic pathways under glucocorticoid exposure.
- **Suggested validation:** Assess AKT and downstream substrate phosphorylation, compare PTEN mRNA vs protein to distinguish transcriptional vs post-translational effects, and test proteasome inhibition or GR antagonism effects on PTEN levels.
- **Priority:** medium

### Hypothesis 4
- **Hypothesis:** Upregulation of PIEZO2 enhances mechanotransduction, potentially modulating GR responses to mechanical cues.
- **Why interesting:** Would reveal a noncanonical, context-dependent link between GR signaling and mechanosensitivity.
- **Suggested validation:** Patch-clamp or calcium imaging under mechanical stimulation; validate PIEZO2 up by qPCR/immunoblot; assess functional impact using PIEZO2 knockdown or pharmacological modulation.
- **Priority:** low

### Hypothesis 5
- **Hypothesis:** Deflazacort remodels ECM/Wnt/TGF-beta signaling tone via opposing changes in regulators (e.g., DKK1, CCN2, ITIH5 down; CD109, FSTL3 up).
- **Why interesting:** Could influence cell–matrix interactions and growth factor signaling in a model-dependent manner.
- **Suggested validation:** Measure SMAD and β-catenin/TCF reporter activities, analyze secreted protein levels, and perform phospho-profiling of TGF-beta/Wnt pathways.
- **Priority:** low

## Confidence
medium

## Caution notes
- Up-regulated programs are inferred from individual proteins without pathway enrichment and should be considered hypothesis-generating.
- Chromatin-related conclusions are limited (single GO term and few proteins); avoid inferring global chromatin state changes.
- Single-protein findings (e.g., PTEN, PIEZO2, NAGS, MIXL1) require orthogonal validation before assigning mechanistic significance.
- RNA processing pathway suppression at the proteome level does not by itself establish functional splicing consequences; transcript-level analyses are needed.
- Cell type/context is unspecified; ECM and secreted factor changes may be model-dependent and should not be overgeneralized.
- NR3C1 decrease magnitude is modest; time-course studies are recommended to confirm GR feedback dynamics.


---

# Delavirdine (mesylate)

# Drug summary: Delavirdine (mesylate)

## Executive summary
In host-cell proteomics, delavirdine (an HIV reverse transcriptase inhibitor) produced a coherent pattern of lipid and stress adaptation with cytoskeletal reorganization. The most robust signal was cholesterol homeostasis upregulation (Hallmark NES 2.40, FDR 0), accompanied by proteostasis stress signatures including the integrated stress response and response to misfolded proteins (GO NES ~1.95). Enrichment for positive regulation of small GTPase–mediated signaling (GO NES ~1.95) aligned with increases in membrane/cell-surface remodeling markers (NRP1/NRP2, MMP14) and a pronounced downregulation of intermediate filament organization (GO down NES −2.29/−2.20) with decreased keratin/cytoskeletal proteins (e.g., KRT14, CNN1). Hallmark Hedgehog (NES 1.92, FDR ~0.006) and mTORC1 (NES 1.58, FDR ~0.045) enrichments were detected but lack canonical readouts among top proteins and should be considered tentative. Together, the data point to off-target or adaptive host responses involving lipid remodeling and proteostasis stress, with concurrent cytoskeletal changes. Mechanistic attributions (e.g., SREBP-driven sterol programs, ER stress origin, autophagy) remain hypotheses pending targeted validation.

## Findings consistent with expected biology
- None listed.

## Unexpected but plausible findings
- Strong activation of cholesterol homeostasis programs (Hallmark NES 2.40, FDR 0), supported by increased lipid/sterol-related proteins (e.g., ABHD2, DDHD1, COQ4).
- Upregulation of proteostasis stress and integrated stress response gene sets (GO NES ~1.95), with increases in ER/protein quality-control markers (e.g., AGR2, HID1).
- Enrichment of positive regulation of small GTPase–mediated signaling (GO NES ~1.95), with upregulation of MCF2L and RGL1 and increased NRP1/NRP2 and MMP14, consistent with membrane/cytoskeletal remodeling.
- Marked downregulation of intermediate filament/keratin biology (GO down NES −2.29/−2.20) with decreased KRT14 and CNN1, indicating cytoskeletal reorganization.
- Hallmark Hedgehog (NES 1.92) and mTORC1 (NES 1.58) enrichments at the protein-abundance level, although canonical pathway components are not prominent among top hits and functional activation is unconfirmed.
- Suppression of selected epithelial/secretory/metabolic markers (e.g., IGFBP2, HNF4G) suggesting a shift away from epithelial programs, without corroborating pathway-level enrichment.

## Follow-up hypotheses
### Hypothesis 1
- **Hypothesis:** Delavirdine triggers SREBP-driven sterol/lipid remodeling to adapt membrane/ER homeostasis.
- **Why interesting:** Explains the dominant cholesterol homeostasis signature and could mechanistically couple lipid remodeling to observed proteostasis stress.
- **Suggested validation:** Assess SREBP1/2 cleavage and nuclear translocation; quantify canonical targets (HMGCR, SQLE, LDLR) by proteomics/qPCR; perform lipidomics (cholesterol/sterol esters) and filipin staining; test sensitivity to SREBP pathway inhibition.
- **Priority:** high

### Hypothesis 2
- **Hypothesis:** Delavirdine induces an integrated stress response/ER stress program.
- **Why interesting:** Accounts for ISR and misfolded-protein response enrichments and AGR2/HID1 increases; provides a unifying driver for adaptive proteome changes.
- **Suggested validation:** Measure eIF2α phosphorylation, ATF4 and CHOP protein levels, and XBP1 splicing; evaluate rescue with ISRIB or PERK inhibition; confirm AGR2 upregulation and ER chaperone induction.
- **Priority:** high

### Hypothesis 3
- **Hypothesis:** Hedgehog and/or mTORC1 pathways are engaged secondarily to stress and lipid remodeling.
- **Why interesting:** Could represent pro-survival/anabolic adaptation to stress, aligning with Hallmark enrichments.
- **Suggested validation:** Test GLI1/GLI2 target induction and sensitivity to SMO antagonists (Hedgehog); assess S6K/S6 phosphorylation and rapamycin/torin sensitivity (mTORC1).
- **Priority:** medium

### Hypothesis 4
- **Hypothesis:** Cytoskeletal remodeling reduces keratin/intermediate filament networks and may enhance cell plasticity.
- **Why interesting:** Integrates strong downregulation of intermediate filament organization with increases in membrane remodeling proteins.
- **Suggested validation:** Immunofluorescence for keratin networks (e.g., KRT14); measure migration/invasion (wound healing, transwell); assess EMT-associated markers; test functional impact of NRP1/NRP2 or MMP14 inhibition.
- **Priority:** low

### Hypothesis 5
- **Hypothesis:** Autophagy/lysosomal flux is modestly engaged as an adaptive response.
- **Why interesting:** ATG12 upregulation hints at autophagy involvement alongside proteostasis stress.
- **Suggested validation:** Monitor LC3-II and p62 turnover with and without bafilomycin; LysoTracker staining; assess autophagosome markers and autophagic flux assays.
- **Priority:** low

## Confidence
medium

## Caution notes
- Pathway inferences are based on protein abundance; activation versus compensatory responses cannot be distinguished without activity/phosphoproteomic readouts.
- Hedgehog and mTORC1 enrichments lack canonical activity markers among top proteins; treat as tentative until validated (e.g., GLI targets, S6K/S6 phosphorylation).
- Cholesterol homeostasis interpretation would be strengthened by direct evidence of SREBP activation and canonical targets; current support rests on gene set enrichment and a subset of lipid-related proteins.
- Autophagy/lysosome engagement is underpowered (e.g., ATG12, CD68) and requires flux assays for confirmation.
- Small GTPase and motility interpretations are broad; there is no direct EMT/migration pathway enrichment and conclusions should be conservative.
- The GO term 'VISUAL_BEHAVIOR' reflects annotation noise and is not mechanistically informative here.
- No Reactome/KEGG enrichment was detected, reducing redundancy support for pathway calls.
- Observed host-cell changes are likely off-target or adaptive and may be cell-type–specific, not directly related to the antiviral mechanism.


---

# Desloratadine

# Drug summary: Desloratadine

## Executive summary
Desloratadine exposure produced a coherent proteomic shift characterized by strong induction of cholesterol/sterol and lipoprotein programs and concurrent enrichment of cytokine/JAK–STAT–linked inflammatory, complement, and coagulation signatures. ECM/tissue-remodeling and angiogenesis signatures were also enriched, while cell-cycle, DNA replication/chromosome organization, and ribosome biogenesis programs were broadly suppressed, consistent with growth attenuation. Notably, a cluster of neuronal/vesicular trafficking proteins increased despite no dedicated pathway enrichment, and Wnt-modulatory nodes showed opposing changes, indicating morphogen pathway crosstalk with unclear net direction. These conclusions are based on abundance-driven pathway enrichments and selected protein changes; signaling activation and causal linkage to histamine receptor antagonism are not established.

## Findings consistent with expected biology
- Robust upregulation of cholesterol/sterol metabolism and lipoprotein handling (e.g., CHOLESTEROL_HOMEOSTASIS and sterol-related GO terms up; SC5D, APOA5 increased).
- Enrichment consistent with cytokine/JAK–STAT–associated inflammatory, complement, and coagulation programs (IL6_JAK_STAT3_SIGNALING, IL2_STAT5_SIGNALING, INFLAMMATORY_RESPONSE, COMPLEMENT, COAGULATION; increased IL4R, IL10RB, RNF128, NDFIP1).
- Coordinated suppression of proliferation programs and biosynthetic capacity (Hallmarks E2F_TARGETS and G2M_CHECKPOINT down; GO DNA replication/chromosome organization and ribosome biogenesis down; decreased RPL29, RPL39).
- ECM/tissue remodeling and angiogenesis signatures enriched, including EMT- and TGF-β–related sets, supported by proteins such as CEMIP, NRP2, DLK1, TYRO3 and CELSR2.

## Unexpected but plausible findings
- Marked increase in neuronal/vesicular trafficking–associated proteins (TMEM59L, NRSN2, SV2A, SGCE, TMC7) without parallel pathway enrichment.
- Opposing regulation of Wnt modulators (ZNRF3 up; DKK1 down) alongside EMT/TGF-β and HEDGEHOG hallmark enrichment, suggesting complex morphogen crosstalk with unclear net Wnt direction.
- Tentative indications of altered autophagy–lysosome axis (TP53INP2 up with TFEB and CLN5 down) without pathway-level enrichment support.

## Follow-up hypotheses
### Hypothesis 1
- **Hypothesis:** Desloratadine induces a sterol/cholesterol biosynthesis and transport program, increasing cellular lipid handling and membrane sterol content.
- **Why interesting:** Membrane lipid remodeling can broadly influence receptor signaling, trafficking, and drug response; it is a dominant proteomic signature here.
- **Suggested validation:** Quantify cellular cholesterol and sterols (filipin staining, targeted lipidomics), assess SREBP processing and expression of key cholesterol biosynthetic enzymes by immunoblot/qPCR, and confirm APOA5/SC5D protein changes in independent replicates.
- **Priority:** high

### Hypothesis 2
- **Hypothesis:** Desloratadine suppresses cell-cycle progression leading to growth arrest with decreased DNA replication and ribosome biogenesis.
- **Why interesting:** Broad downregulation of E2F/G2M and replication/ribosome programs suggests a major functional impact on proliferation.
- **Suggested validation:** Measure DNA synthesis (EdU incorporation), mitotic markers (phospho–histone H3), cell-cycle distribution (flow cytometry), and colony growth; assess apoptosis versus cytostatic arrest (Annexin V/PI, cleaved PARP).
- **Priority:** high

### Hypothesis 3
- **Hypothesis:** Cytokine/JAK–STAT–related signaling is engaged at the level of pathway components, contributing to inflammatory, complement, and coagulation signatures.
- **Why interesting:** Receptor upregulation (IL4R, IL10RB) and pathway enrichments suggest cytokine responsiveness that could shape downstream remodeling and immune-related effects.
- **Suggested validation:** Assess STAT3/STAT5 phosphorylation and nuclear translocation, JAK dependency using small-molecule inhibition, and cytokine secretion profiles; corroborate target gene induction by qPCR.
- **Priority:** medium

### Hypothesis 4
- **Hypothesis:** Desloratadine promotes ECM/tissue remodeling with angiogenic features rather than a full EMT transition.
- **Why interesting:** Enrichment of EMT/TGF-β/angiogenesis hallmarks and increased CEMIP, NRP2, DLK1, TYRO3 suggest microenvironmental remodeling that may impact tissue responses.
- **Suggested validation:** Quantify canonical EMT and ECM markers (E-cadherin, vimentin, fibronectin, collagen), SMAD2/3 phosphorylation, migration/invasion assays, and secreted ECM protein measurements.
- **Priority:** medium

### Hypothesis 5
- **Hypothesis:** Desloratadine alters vesicle trafficking or secretory pathways, reflected by increased neuronal/vesicular proteins.
- **Why interesting:** A coordinated increase in vesicle-associated proteins could indicate changes in exocytosis/endocytosis that intersect with receptor signaling and lipid handling.
- **Suggested validation:** Track endo-lysosomal and secretory trafficking (transferrin uptake/recycling, VAMP/SV2A localization), quantify extracellular vesicle release, and assess synaptic vesicle protein abundance across conditions.
- **Priority:** medium

### Hypothesis 6
- **Hypothesis:** Morphogen pathway crosstalk is perturbed, with Wnt regulatory nodes remodeled without a clear net direction.
- **Why interesting:** ZNRF3 up and DKK1 down alongside EMT/TGF-β and Hedgehog enrichment imply pathway rewiring that may influence remodeling and proliferation.
- **Suggested validation:** Measure Wnt/β-catenin activity (TOPFlash reporter, β-catenin localization), assess canonical Wnt target gene expression, and test sensitivity to Wnt pathway modulators.
- **Priority:** medium

## Confidence
high

## Caution notes
- Pathway enrichments are based on protein abundance changes and do not by themselves prove signaling activation; phosphoproteomic or functional readouts are needed for confirmation.
- Cell type/context is unspecified; EMT, myogenesis, and angiogenesis terms may reflect generalized ECM remodeling rather than true lineage transitions.
- Several observations (neuronal/vesicular protein cluster, Wnt modulators, autophagy–lysosome hints) rely on small protein subsets and lack pathway-level corroboration; treat as tentative.
- Causality to histamine H1 receptor antagonism is not established; observed changes may be downstream or off-target effects.
- Suppression of proliferation may reflect growth arrest rather than cytotoxicity; functional assays are needed to distinguish outcomes.
- KEGG and Reactome sections are empty; conclusions are drawn from Hallmark and GO results and should be interpreted within this scope.
- mTORC1 and KRAS hallmark enrichments were detected but lack supporting effector-level protein changes here; any inference of activation should be cautious.


---

# Dexpramipexole (dihydrochloride)

# Drug summary: Dexpramipexole (dihydrochloride)

## Executive summary
Proteomics after Dexpramipexole (dihydrochloride) exposure shows a modest but coherent induction of an inflammatory/innate immune program and a significant downregulation of intermediate filament organization. The inflammatory signal is supported by Hallmark INFLAMMATORY_RESPONSE enrichment (NES ~1.68, FDR < 0.05) and increased proteins such as JUNB, ISG20, HLA-B, TRIM8, GMEB1, and SP140L. The cytoskeletal signal is supported by GO down-regulation of INTERMEDIATE_FILAMENT_ORGANIZATION (NES ~−2.04, FDR < 0.01) with decreased KRT79. Additional protein-level shifts suggest membrane/lipid–ECM remodeling (SMPDL3A, NEU3, CEMIP, TLCD3A), possible GPCR/cAMP feedback (PDE4D up; ADGRL2 up), and increased mechanosensory-channel proteins (PIEZO2, TMC7), but these lack pathway corroboration and should be treated as tentative. No broad pathway enrichment beyond the two terms above was detected. Overall, the profile indicates an acute stress/inflammatory response with cytoskeletal remodeling, with medium confidence given the limited pathway breadth and unknown cellular context.

## Findings consistent with expected biology
- Induction of immediate-early/transcriptional regulators (e.g., JUNB up), consistent with an acute cellular response to stimulation.

## Unexpected but plausible findings
- Inflammatory/innate immune activation with increased MHC class I component levels (HLA-B), supported by Hallmark INFLAMMATORY_RESPONSE enrichment.
- Down-regulation of intermediate filament organization, supported by a significant GO BP term and decreased KRT79.
- Protein-level increases in mechanosensory/neuronal-associated channels (PIEZO2, TMC7) without pathway-level support.
- Tentative evidence for membrane lipid/ECM remodeling (SMPDL3A, NEU3, CEMIP, TLCD3A up) without corroborating pathway enrichment.
- Protein-level suggestion of GPCR/cAMP feedback (PDE4D up; ADGRL2 up) without pathway enrichment.

## Follow-up hypotheses
### Hypothesis 1
- **Hypothesis:** Dexpramipexole elicits a pro-inflammatory/innate immune response that includes increased MHC I component abundance.
- **Why interesting:** An immune activation signature is unexpected for a dopaminergic-class agent and could reflect off-target or stress-related effects relevant to efficacy or tolerability.
- **Suggested validation:** Time-course and dose-response qPCR/proteomics for inflammatory genes; NF-κB/IRF reporter or phospho-assays; cytokine/chemokine secretion profiling; surface HLA class I quantification by flow cytometry.
- **Priority:** high

### Hypothesis 2
- **Hypothesis:** Intermediate filament organization is reduced following treatment, indicating cytoskeletal remodeling.
- **Why interesting:** Cytoskeletal changes may impact cell morphology, resilience, and signaling, with potential relevance to neuronal processes.
- **Suggested validation:** Immunofluorescence imaging and Western blot of intermediate filament proteins appropriate to the model (e.g., keratins, vimentin); assessment of cell morphology and mechanical properties; rescue or perturbation studies of IF components.
- **Priority:** medium

### Hypothesis 3
- **Hypothesis:** Membrane lipid and hyaluronan/sialic acid processing are altered (via SMPDL3A, NEU3, CEMIP, TLCD3A), secondarily coupling to the inflammatory state.
- **Why interesting:** Cell-surface lipid/glycocalyx remodeling can modulate receptor signaling and immune interactions.
- **Suggested validation:** Targeted lipidomics (sphingolipids, ceramides, gangliosides); sialidase activity assays; hyaluronan content/turnover measurements; knockdown or pharmacologic modulation of NEU3/SMPDL3A followed by proteomic/functional readouts.
- **Priority:** medium

### Hypothesis 4
- **Hypothesis:** Protein-level changes reflect compensatory GPCR/cAMP pathway adaptation (PDE4D up; ADGRL2 up).
- **Why interesting:** Could indicate feedback to GPCR signaling consistent with the drug class, though not yet supported at the pathway level.
- **Suggested validation:** Cellular cAMP measurements (basal and upon GPCR agonism); PDE activity assays; PDE4 inhibition challenge to test reversibility; PKA substrate phosphorylation readouts.
- **Priority:** low

### Hypothesis 5
- **Hypothesis:** Upregulation of mechanosensory channels (PIEZO2, TMC7) alters mechanotransduction or membrane tension sensing.
- **Why interesting:** May link membrane remodeling to sensory signaling; relevance depends on cell type and baseline channel expression.
- **Suggested validation:** Channel expression confirmation (qPCR/Western); mechanosensitive current recordings or Ca2+ imaging under mechanical stimulation; loss-of-function studies to assess functional contribution.
- **Priority:** low

## Confidence
medium

## Caution notes
- Pathway enrichment is limited to one Hallmark up term and one GO BP down term; broader conclusions should be restrained.
- Proteomic abundance changes do not establish pathway activity; functional assays are required for GPCR/cAMP, immune signaling, or cytoskeletal dynamics.
- Increased HLA-B indicates higher MHC I component levels but does not by itself confirm antigen presentation activity.
- Interpretation of intermediate filament suppression should be cautious; the GO signal is strong but supported by a limited set of detected proteins (e.g., KRT79).
- Membrane/lipid–ECM remodeling and GPCR/cAMP feedback are inferred from few proteins without pathway corroboration and should be treated as tentative.
- Cell type and experimental context are not specified; ‘neuronal-associated’ and immune interpretations may vary with baseline expression; replication with dose/time series is advised.


---

# DFO

# Drug summary: DFO

## Executive summary
DFO induces a robust hypoxia-like stress program with a metabolic shift toward glycolysis and away from mitochondrial respiration, accompanied by p53-pathway engagement and broad cell-cycle repression. Proteomics shows strong upregulation of canonical hypoxia/stress and glycolytic effectors (PFKFB3, NDRG1, BHLHE40, SERPINE1, LOXL2, GDF15, TXNIP, IGFBP1) with pathway-level suppression of oxidative phosphorylation and electron transport. Cell-cycle and DNA replication programs are repressed (G2/M and E2F down), concordant with increased CDKN1A (p21) and TP53INP2 and decreased CDC25B, CDT1, and CDKN3. Extracellular matrix remodeling and collagen biosynthesis/organization are prominently induced (EMT hallmark with LOXL2, COL7A1, COL9A3, CTHRC1, CEMIP, SERPINE1), alongside pro-inflammatory signaling (TNF–NFkB/inflammatory response). A heme metabolism hallmark is enriched and several epidermis/keratinocyte differentiation GO terms are up; both are interpreted cautiously as they are primarily pathway-driven. Notably, HMGCR is strongly decreased, suggesting possible suppression of cholesterol/mevalonate metabolism. Overall, findings are coherent with a hypoxia-mimetic, stress-adaptive response that halts proliferation, reprograms metabolism, and remodels the extracellular milieu.

## Findings consistent with expected biology
- Robust hypoxia-like program with glycolysis up and mitochondrial oxidative phosphorylation/electron transport pathways down (Hallmark HYPOXIA up; GLYCOLYSIS up; OXIDATIVE_PHOSPHORYLATION down).
- p53-pathway engagement with p21 induction and broad cell-cycle repression (P53_PATHWAY up; G2M_CHECKPOINT and E2F_TARGETS down; CDKN1A up; CDC25B, CDT1, CDKN3 down).
- Extracellular matrix remodeling with increased collagen biosynthesis/organization (EMT hallmark up; collagen-related GO terms up; LOXL2, COL7A1, COL9A3, CTHRC1, CEMIP, SERPINE1 up).
- Pro-inflammatory stress signaling consistent with TNF–NFkB/inflammatory response enrichment (TNFA_SIGNALING_VIA_NFKB and INFLAMMATORY_RESPONSE up; JUNB, SERPINE1, GDF15, TXNIP up).

## Unexpected but plausible findings
- Enrichment of epidermis/keratinocyte differentiation and skin development programs despite the absence of clear core keratin upregulation among top proteins.
- Heme metabolism hallmark up alongside strong pathway-level downregulation of oxidative phosphorylation.
- Marked HMGCR downregulation, suggesting potential suppression of cholesterol/mevalonate biosynthesis.
- IGF receptor signaling pathway enrichment appears driven largely by IGFBP1 upregulation and should be interpreted cautiously.

## Follow-up hypotheses
### Hypothesis 1
- **Hypothesis:** DFO triggers HIF-mediated hypoxia signaling that drives glycolytic reprogramming and p21-dependent cell-cycle arrest.
- **Why interesting:** Integrates the dominant pathway signals (HYPOXIA, GLYCOLYSIS up; OXPHOS down) with coherent protein changes (PFKFB3, NDRG1, BHLHE40, CDKN1A), providing a unifying mechanism for the stress-adaptive phenotype.
- **Suggested validation:** Measure HIF1A/HIF2A stabilization and target gene induction; test dependency by HIF1A/EPAS1 knockdown or HIF transcriptional inhibitor and assess effects on PFKFB3, NDRG1, CDKN1A; perform Seahorse assays (OCR/ECAR) to confirm metabolic shift.
- **Priority:** high

### Hypothesis 2
- **Hypothesis:** Mitochondrial respiration is functionally suppressed with compensatory glycolysis increase under DFO.
- **Why interesting:** Consistent pathway-level suppression of ETC/OXPHOS with glycolysis up suggests a bioenergetic reprogramming that may underlie growth arrest and stress resistance.
- **Suggested validation:** Quantify basal/maximal OCR and ECAR (Seahorse), assess ETC complex protein levels/activities, and test whether glycolysis inhibitors exacerbate DFO sensitivity.
- **Priority:** high

### Hypothesis 3
- **Hypothesis:** DFO elicits a wound-healing/ECM remodeling program rather than full EMT.
- **Why interesting:** EMT hallmark and collagen biosynthesis are strongly enriched with increases in LOXL2, SERPINE1, and collagens, but canonical epithelial–mesenchymal marker shifts are not evident among top proteins.
- **Suggested validation:** Assess epithelial/mesenchymal markers (E-cadherin, Vimentin, ZEB1/2), cell morphology/migration assays, and collagen cross-linking (LOXL2 activity).
- **Priority:** medium

### Hypothesis 4
- **Hypothesis:** DFO suppresses the cholesterol/mevalonate pathway, as indicated by decreased HMGCR.
- **Why interesting:** A pronounced HMGCR decrease may reveal an additional metabolic axis affected by DFO with implications for membrane synthesis and signaling.
- **Suggested validation:** Profile mevalonate pathway enzymes (MVK, MVD, SQLE) by proteomics/qPCR, measure cellular sterol intermediates, and test rescue with mevalonate supplementation.
- **Priority:** medium

### Hypothesis 5
- **Hypothesis:** Pro-inflammatory NFkB signaling is activated downstream of DFO-induced stress.
- **Why interesting:** TNFA_SIGNALING_VIA_NFKB and INFLAMMATORY_RESPONSE are enriched, aligning with upregulation of stress-responsive factors (JUNB, TXNIP, SERPINE1, GDF15).
- **Suggested validation:** Measure NFkB p65 nuclear translocation, IκBα phosphorylation/degradation, and NFkB reporter activity; quantify cytokine/chemokine secretion.
- **Priority:** medium

### Hypothesis 6
- **Hypothesis:** DFO perturbs heme/iron handling pathways in this system.
- **Why interesting:** HEM_METABOLISM hallmark is enriched, suggesting altered heme/iron dynamics that could interact with metabolic and stress responses.
- **Suggested validation:** Assess HMOX1, ALAS1/2, FECH, ferritin (FTH1/FTL) levels; measure labile iron pool and cellular heme content.
- **Priority:** medium

## Confidence
high

## Caution notes
- Several conclusions (e.g., NFkB activation, heme metabolism, IGF signaling) rely primarily on pathway enrichment rather than multiple concordant individual protein changes; interpret these with caution.
- Use 'p53-pathway engagement' rather than 'p53-driven' without direct measures of p53 activation.
- The EMT signal may reflect ECM remodeling/wound-response; canonical EMT marker shifts (e.g., CDH1 down, VIM up) are not highlighted among top changes.
- Epidermis/keratinocyte differentiation GO enrichments should be tempered in the absence of clear keratin upregulation.
- Do not attribute mismatch repair suppression based on MUTYH (base excision repair); the MMR pathway down signal is pathway-centric, and changes in MSH/MLH/PMS proteins were not highlighted.
- ETC/OXPHOS suppression is supported at the pathway level; few specific ETC subunits appear among the top downregulated proteins.
- IGF-axis engagement is tentative, largely driven by IGFBP1 increase; additional IGF pathway components were not prominent.
- Reactome and KEGG sections are empty; conclusions are based on Hallmark and GO evidence and protein-level changes.


---

# Diacerein

# Drug summary: Diacerein

## Executive summary
Diacerein exposure produced a stress-adaptive proteomic state dominated by a robust hypoxia-like program (HALLMARK HYPOXIA up; NES 2.08, FDR 0.0019) and a coordinated reduction in RNA splicing/spliceosome-associated processes (multiple GO terms down; NES ~ -1.9 to -2.0). A modest, suggestive engagement of inflammatory TNF/NF-kB signaling was observed (HALLMARK TNFA_SIGNALING_VIA_NFKB up; NES 1.73, FDR 0.0498) with LTBR increased, but canonical NF-kB effectors were not prominent, so this should be treated as tentative. Several stress/secreted proteins (IGFBP1, CEMIP, MT1B, ADA) increased, consistent with an inflammatory/stress milieu. Down-regulation of a GO process annotated as negative regulation of macroautophagy supports a possible increase in autophagic activity, though direct autophagy markers were not among top changes and this remains provisional. Decreases in TP53 and PTEN were noted without accompanying proliferation/cell-cycle pathway enrichment, indicating a context-dependent observation rather than a proliferative shift. No Reactome/KEGG enrichments were detected and some pathway sections were empty; interpretation therefore relies primarily on Hallmark and GO evidence and should be considered with appropriate caution.

## Findings consistent with expected biology
- Suggestive engagement of inflammatory signaling consistent with an immunology/inflammation context (HALLMARK TNFA_SIGNALING_VIA_NFKB up near the FDR threshold; LTBR up-regulated).
- Increased stress/ECM-remodeling and secreted factors (e.g., IGFBP1, CEMIP, MT1B, ADA), fitting a stress/inflammatory milieu.

## Unexpected but plausible findings
- Robust hypoxia-like signature at the pathway level despite limited classic HIF targets among the top individual proteins.
- Coordinated down-regulation of RNA splicing/spliceosome-associated processes (multiple GO terms significantly down).
- Suppression of an estrogen-responsive program (GO: response to estrogen) at the enrichment level.
- Decreases in TP53 and PTEN without concurrent proliferation/cell-cycle pathway activation.
- Potential derepression of autophagy inferred from down-regulation of 'negative regulation of macroautophagy' (tentative).

## Follow-up hypotheses
### Hypothesis 1
- **Hypothesis:** Diacerein induces hypoxia/HIF-driven signaling in this context.
- **Why interesting:** HALLMARK HYPOXIA is strongly enriched (NES 2.08, FDR 0.0019), indicating a dominant program that may shape downstream stress and metabolic responses.
- **Suggested validation:** Assess HIF1A/HIF2A stabilization and nuclear localization; measure canonical HIF targets (e.g., CA9, VEGFA) by proteomics/qPCR; evaluate oxygen consumption and glycolytic shift (Seahorse).
- **Priority:** high

### Hypothesis 2
- **Hypothesis:** Diacerein broadly suppresses spliceosome components and RNA splicing capacity.
- **Why interesting:** Multiple splicing-related GO terms are significantly down, suggesting a proteome-wide reduction in core post-transcriptional processing.
- **Suggested validation:** Survey a broader panel of spliceosome proteins beyond the top hits; perform RNA-seq to quantify intron retention/exon skipping; immunoblot core factors (e.g., SF3B1, SNRNP proteins).
- **Priority:** high

### Hypothesis 3
- **Hypothesis:** TNF/NF-kB signaling is engaged but modest at the proteome level.
- **Why interesting:** HALLMARK TNFA_SIGNALING_VIA_NFKB is enriched near the FDR cutoff and LTBR is up-regulated, aligning with an inflammatory mechanism.
- **Suggested validation:** Monitor RELA (p65) nuclear translocation and IκBα (NFKBIA) degradation; measure NF-kB target transcripts/proteins (e.g., IL6, CCL2) by qPCR/ELISA; include orthogonal phospho- or reporter assays.
- **Priority:** medium

### Hypothesis 4
- **Hypothesis:** Autophagic flux increases as part of a stress-adaptive response.
- **Why interesting:** Down-regulation of a GO term for negative regulation of macroautophagy is consistent with derepression of autophagy, potentially coordinating with reduced RNA splicing.
- **Suggested validation:** Assess LC3-II turnover with/without bafilomycin; quantify p62/SQSTM1 dynamics; image LC3 puncta; profile ATG proteins.
- **Priority:** medium

### Hypothesis 5
- **Hypothesis:** Reductions in TP53 and PTEN reflect context-dependent modulation of stress checkpoints without inducing proliferation.
- **Why interesting:** Both proteins decrease at the proteome level but no proliferation/cell-cycle enrichment is observed, implying a noncanonical or transient effect.
- **Suggested validation:** Measure p53 transcriptional activity (e.g., CDKN1A, MDM2), p53 stabilization, and PTEN levels/activity; evaluate cell-cycle markers and viability under treatment.
- **Priority:** low

## Confidence
medium

## Caution notes
- The TNF/NF-kB signal is modest (hallmark FDR ~0.05) and lacks broad canonical target representation; interpret as suggestive rather than definitive activation.
- The hypoxia hallmark is robust but classic HIF targets are not among the most changed proteins; exclude culture-condition artifacts and confirm with orthogonal assays.
- Splicing suppression is inferred from gene-set enrichment; few spliceosome components show large individual fold-changes among top hits.
- Autophagy inference is based on a regulatory GO term without direct autophagy effectors among leading proteins and requires dedicated validation.
- No Reactome/KEGG enrichments were detected and some pathway sections are empty; conclusions rely primarily on Hallmark/GO evidence from proteomics alone.
- Protein-level decreases in TP53/PTEN are context dependent and not accompanied by proliferation/cell-cycle pathway signatures; avoid functional overinterpretation.


---

# Digitoxin

# Drug summary: Digitoxin

## Executive summary
Proteomic profiling of Digitoxin exposure reveals a coherent stress-adaptive epithelial response consistent with Na+/K+ ATPase inhibition. Strong GO enrichment (NES >2, FDR≈0) supports induction of epithelial differentiation/adhesion programs (keratinocyte/epidermal differentiation, intermediate filament organization, tight junction assembly) and engagement of actin/lamellipodium/ARP2/3-associated processes. Ion/energy homeostasis adaptation is indicated by enrichment for regulation of potassium ion transport and mitochondrial electron transport, with concordant increases in ATP5IF1, TXNIP, GDF15 and AP-1/small Maf factors (JUNB, FOSL1, MAFF, MAFG). Protein-level decreases in cell-cycle/replication factors (e.g., CDC25B, CDT1, MCM9, USP1, CDCA7) support proliferative arrest, although no downregulated pathway sets were reported. Unexpected features include a strong enrichment for U2-type prespliceosome assembly, coordinated upregulation of IGFBP2/3/5, and marked suppression of metallothioneins (MT1E/G/X) despite redox stress markers. Together, the data indicate a shift toward an epithelial, stress-adapted, less proliferative state. Interpretations of function (e.g., junction reinforcement, IGF signaling changes) should be considered hypotheses pending targeted validation.

## Findings consistent with expected biology
- Ion and energy-stress adaptation consistent with Na+/K+ ATPase inhibition: enrichment for regulation of potassium ion transport and mitochondrial electron transport with increased ATP5IF1, TXNIP, GDF15 and AP-1/small Maf factors (JUNB, FOSL1, MAFF, MAFG).
- Engagement of epithelial cytoskeletal and junctional programs: strong up-enrichment for intermediate filament organization, tight junction assembly, and lamellipodium/ARP2/3-related processes with concordant proteins (e.g., DSG3, ESAM, CST5).
- Suppression of proliferation-associated proteins: robust decreases in cell-cycle/replication factors (CDC25B, CDT1, MCM9, USP1, CDCA7), consistent with reduced proliferation (protein-driven inference).
- Epithelial differentiation-like signature: enrichment for keratinocyte/epidermal differentiation terms with supporting proteins (S100A7, DSG3, CST5), indicating a shift toward a more differentiated epithelial state.

## Unexpected but plausible findings
- Strong up-enrichment of U2-type prespliceosome assembly (NES≈2.35, FDR≈0), suggesting splicing machinery engagement not directly anticipated from Na+/K+ ATPase inhibition.
- Coordinated induction of IGFBP2, IGFBP3, and IGFBP5, consistent with an IGF-modulatory secretome; functional impact on IGF signaling remains uncertain.
- Marked suppression of metallothioneins (MT1X, MT1G, MT1E) despite concurrent redox/mitochondrial stress markers (TXNIP, GDF15), diverging from canonical MT induction under oxidative stress.
- TRPC5 upregulation alongside potassium transport regulatory enrichment, implicating nonselective cation/Ca2+ channel involvement within a broader ion-homeostasis response.

## Follow-up hypotheses
### Hypothesis 1
- **Hypothesis:** Digitoxin induces an epithelial differentiation and adhesion program that contributes to proliferative arrest.
- **Why interesting:** Aligns strong pathway enrichment (keratinocyte/epidermal differentiation, junction assembly) with decreased cell-cycle proteins, suggesting a coordinated cell-state shift relevant to efficacy and epithelial toxicities.
- **Suggested validation:** Assess junction/differentiation markers (DSG3, claudins, ZO-1, involucrin/filaggrin) by immunofluorescence and immunoblot; measure TEER and desmosome/tight junction morphology; quantify proliferation markers (Ki-67, pRB) and cell-cycle profiles.
- **Priority:** high

### Hypothesis 2
- **Hypothesis:** Na+/K+ ATPase inhibition elicits a mitochondrial/energy-stress axis involving ATP5IF1 induction and AP-1/small Maf activation, culminating in increased GDF15 and TXNIP.
- **Why interesting:** Provides a mechanistic link between ion pump perturbation and metabolic/redox signaling that may underlie tolerability and therapeutic windows.
- **Suggested validation:** Measure ATP/AMP and AMPK activation; assess OCR/ECAR and mitochondrial membrane potential; evaluate ATP5IF1 levels and phosphorylation; quantify ROS and GDF15 secretion (ELISA); probe AP-1/sMaf target gene induction.
- **Priority:** high

### Hypothesis 3
- **Hypothesis:** Digitoxin modulates pre-mRNA splicing as part of the stress response.
- **Why interesting:** Strong prespliceosome enrichment suggests post-transcriptional regulation that could reshape stress and proliferation programs.
- **Suggested validation:** Perform RNA-seq to profile alternative splicing changes; assess spliceosome component abundance and phosphorylation; test sensitivity to splicing modulators for synergy or additive effects.
- **Priority:** medium

### Hypothesis 4
- **Hypothesis:** IGFBP2/3/5 induction reduces IGF bioavailability, contributing to growth arrest.
